# Syntaxin 5 determines Weibel-Palade body size and Von Willebrand factor secretion by controlling Golgi architecture

**DOI:** 10.1101/2021.12.10.472050

**Authors:** Marije Kat, Ellie Karampini, Arie Johan Hoogendijk, Petra Bürgisser, Aat A. Mulder, Floris van Alphen, Jenny Olins, Dirk Geerts, Maartje van den Biggelaar, Coert Margadant, Jan Voorberg, Ruben Bierings

**Affiliations:** Molecular Hematology, Sanquin Research and Landsteiner Laboratory, Amsterdam University Medical Center, University of Amsterdam, The Netherlands; Vascular Biology, Royal College of Surgeons, Dublin, Ireland; Hematology, Erasmus University Medical Center, Rotterdam, The Netherlands; Molecular Cell Biology, Leiden University Medical Center, Leiden, The Netherlands; Medical Biology, Amsterdam University Medical Center, location AMC, University of Amsterdam, The Netherlands; Angiogenesis laboratory, Cancer Center Amsterdam, Amsterdam University Medical Center, location VUmc, The Netherlands; Experimental Vascular Medicine, Amsterdam University Medical Center, University of Amsterdam, The Netherlands

## Abstract

Von Willebrand factor (VWF) is a multimeric hemostatic protein primarily synthesized in endothelial cells (ECs). VWF is stored in endothelial storage organelles, the Weibel-Palade bodies (WPBs), whose biogenesis strongly depends on VWF anterograde trafficking and Golgi architecture. Elongated WPB morphology is correlated to longer VWF strings with better adhesive properties. We previously identified the SNARE SEC22B, which is involved in anterograde ER-to-Golgi transport, as a novel regulator of WPB elongation. To elucidate novel determinants of WPB morphology we explored endothelial SEC22B interaction partners in a mass spectrometrybased approach, identifying the Golgi SNARE Syntaxin 5 (STX5). We established STX5 knockdown in ECs using shRNA-dependent silencing and analyzed WPB and Golgi morphology, using confocal and electron microscopy. STX5-depleted ECs exhibited extensive Golgi fragmentation and decreased WPB length, which was associated with reduced intracellular VWF levels, and impaired stimulated VWF secretion. However, the secretion-incompetent organelles in shSTX5 cells maintained WPB markers such as Angiopoietin 2, P-selectin, Rab27A, and CD63. Taken together, our study has identified SNARE protein STX5 as a novel regulator of WPB biogenesis.

## Introduction

Von Willebrand factor (VWF) is a hemostatic glycoprotein that is primarily synthesized in endothelial cells (ECs) and acts as a factor VIII chaperone as well as an adhesive grid for thrombus formation.^1^ Decreased VWF plasma levels or mutations in the *VWF* gene can cause Von Willebrand disease (VWD), the most common bleeding disorder.^2^ VWF undergoes a multistep maturation process that involves dimerization in the endoplasmic reticulum (ER), followed by multimerization and proteolytic processing in the Golgi.^1^ Smaller VWF multimers are continuously secreted (primarily) at the basolateral surface via the constitutive pathway, whilst larger VWF multimers are condensed into storage organelles emerging from the *trans*-Golgi network (TGN): the Weibel-Palade bodies (WPBs).^3^

WPB biogenesis is tightly linked to VWF synthesis, which is highlighted by the absence of WPBs in VWF knockout ECs, and their de novo formation in non-ECs by ectopic VWF expression.^4–6^ Besides VWF, WPBs contain inflammatory and angiogenic proteins, and recruit essential transport and exocytotic machinery.^7,8^ WPB exocytosis occurs via basal (continuous) or regulated (stimulus-induced) secretion pathways, both predominantly targeting the apical surface facing the blood vessel lumen.^3^ WPBs play an important role during primary hemostasis as their release ensures the immediate delivery of VWF (and other molecules) in the vessel lumen in response to injury, whereupon VWF tubules unfurl into long VWF strings on the apical surface, which subsequently become decorated by platelets.^8^

WPBs have a distinct, elongated morphology: the cigar-shaped structure is composed of densely packed helical tubules of VWF multimers running along the length of the organelle enwrapped by a tightly fitted endomembrane.^9,10^ Both the length and adhesive properties of VWF strings correlated with WPB length; shorter WPBs generate shorter VWF strings, with lower adhesive capacity for platelets and plasma VWF.^11,12^ However, what drives their distinct morphology is still largely unknown. The range in WPB size was defined by the VWF quanta model, which describes how during biogenesis VWF nanoclusters of a discrete length (i.e. quanta) are co-packaged in variable numbers at the Golgi, ultimately determining the length of the WPB.^13^ Although WPB length is known to be determined by Golgi ribbon architecture as well as by levels of VWF synthesis,^13,14^ only recently has the control of VWF progression through the early secretory pathway been appreciated as a determinant of WPB length.^15–17^

In anterograde transport vesicles bud off at ER exit sites, containing specific cargo *en route* to the Golgi. During this process soluble N-ethylmaleimide-sensitive fusion protein attachment protein receptors (SNAREs) are incorporated in the vesicle membrane (i.e. v-SNAREs), which can form complexes with Golgi-associated t-SNAREs to facilitate membrane fusion and cargo release in the Golgi lumen.^18,19^ One study showed that the ARF guanine-nucleotide exchange factor (GEF) Golgi Brefeldin A resistant GEF 1 (GBF1) modulates vesicle fission at the ER and TGN, impacting WPB size by controlling anterograde VWF transport and WPB segregation from the TGN.^15^

Exocytic SNARE proteins play a key role in WPB exocytosis,^20–24^ and some have also been associated with VWF plasma levels and severity of VWD.^25,26^ The role of SNARE proteins in WPB biogenesis and VWF trafficking, however, remains elusive. We have recently characterized the v-SNARE SEC22B as a novel WPB size regulator through its role in anterograde VWF transport and support of elongated Golgi morphology.^16^ In the current study, we aimed at identifying novel determinants of VWF trafficking through mapping ER-Golgi fusion machinery in ECs by elicidating the SEC22B interactome. The proteomic screen revealed a plethora of potential interactors, including the SNARE protein syntaxin-5 (STX5). Knockdown of STX5 in ECs resulted in extensive fragmentation of Golgi architecture, VWF retention in the ER, and significantly shorter and fewer WPBs. In addition, both intracellular VWF levels and regulated WPB exocytosis were significantly suppressed, highlighting STX5 as an essential component of the machinery driving WPB biogenesis and release.

## Methods

### Mass spectrometry analysis

Sample preparation, data acquisition and data analysis were performed as previously described.^27^ Detailed experimental procedures are described in the supplemental material.

### VWF string assay

VWF string assays were essentially performed as described using 75,000 shCTRL or shSTX5 transduced cells per channel in gelatin-coated 6-channel IBIDI μ-slides VI 0.4.^28^ Strings were visualized by supplementing perfusion mix with AF488-conjugated anti-VWF antibody (DAKO) at 2 μM concentration. Further experimental detail can be found in the supplemental material.

### Secretion assay

ECs were transduced and grown in 6-well plates and cultured for 7 days prior to the experiment with regular medium replacement. Basal VWF release was determined as unstimulated secretion over 48 hours. For histamine-induced secretion cells were starved for 30 minutes in M-199 with 1% BSA and subsequently stimulated with 100 μM histamine (Sigma-Aldrich; H7125) for 1 hour. Lysates were obtained in lysis buffer (PBS, 1% Triton X-100) supplemented with Halt protease and Phosphatase inhibitor cocktail. VWF levels were determined by ELISA as described.^21^

Antibodies and DNA constructs are listed in **Table S1** and **Table S2**, respectively. Additional methods can be found in the supplemental material.

## Results

### The SEC22B interactome in ECs contains SNARE proteins involved in anterograde, retrograde and intra-Golgi protein trafficking

To map the composition of the ER-to-Golgi SNARE networks that control VWF trafficking and WPB biogenesis, we employed unbiased affinity purification-mass spectrometry (MS) in ECs using mEGFP-tagged SEC22B as bait. A total of 841 proteins were significantly enriched in mEGFP-SEC22B compared to the mEGFP control (**Figure 1A,B, Table S3**). Gene ontology (GO) enrichment analysis revealed that the most prominent complexes within the cellular components ontology were ‘membrane protein’ ‘inner mitochondrial membrane protein’, and ‘SNARE’ (**Figure 1C**). Furthermore, SEC22B and its potential interacting partners collaborate in 235 enriched biological processes, including anterograde and retrograde ER-Golgi trafficking, intra-Golgi trafficking as well as Golgi organization (**Table S4**).

**Figure 1.**
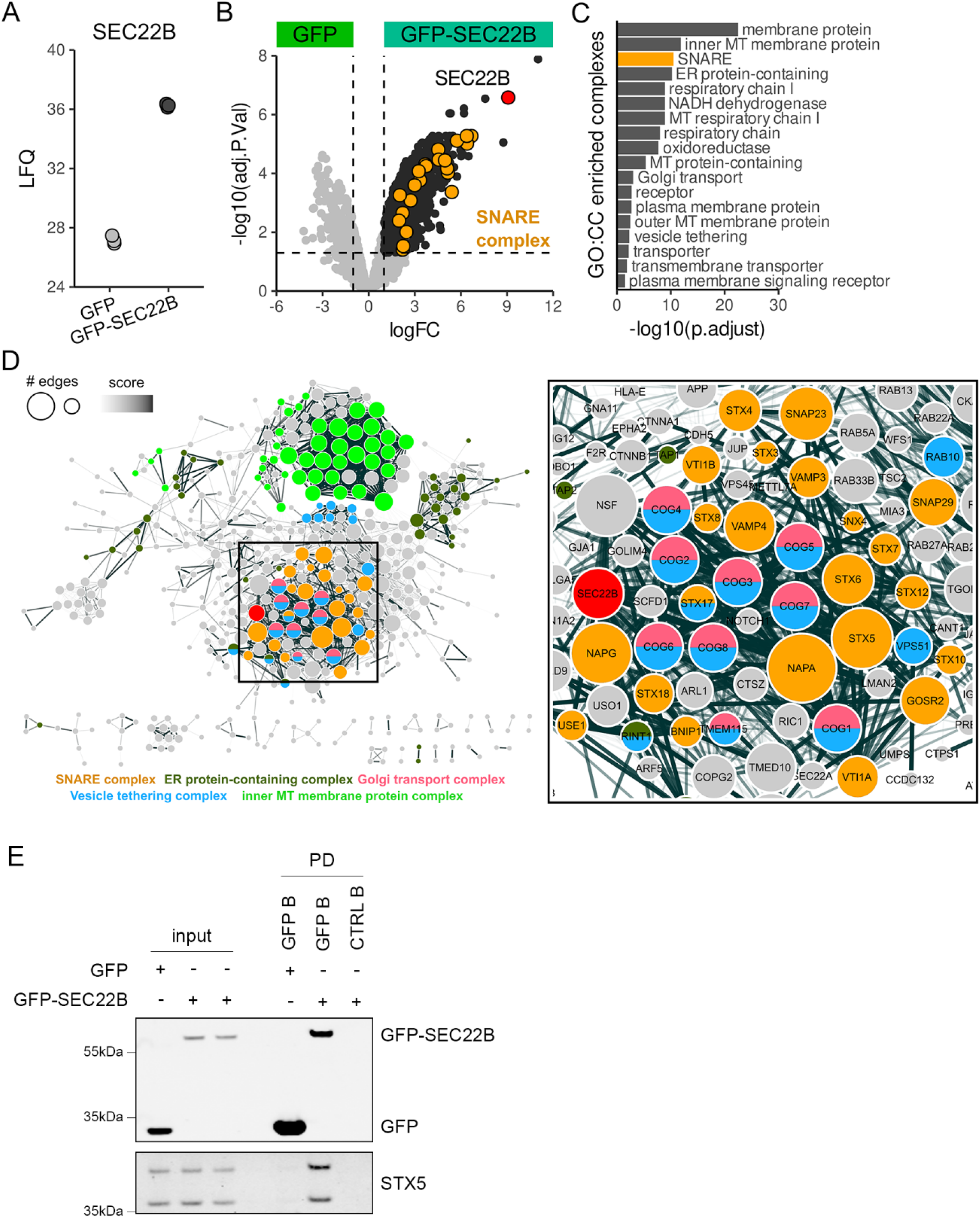
STX5 is part of the SEC22B interactome in endothelial cells. A) Label-free quantification (LFQ) of SEC22B proteins levels in pulldown samples determined by mass spectrometry analysis. B) Volcano plot of significantly enriched proteins in the mEGFP and mEGFP-SEC22B pulldown samples. Red dot represents SEC22B used as bait. Orange dots represent SNARE complex proteins based on GO:0031201 annotation. Dotted lines indicate significance thresholds (P.adjust<0.05 and |LFC|>1). C) Bar plot of enriched GO:CC protein complexes. D) STRING-DB analysis of the enriched proteins in mEGFP-SEC22B showing high confidence interactors (combined STRING-DB scores >0.9). Colors represent enriched complexes: SNARE complex (orange), ER protein-containing complex (dark green), Golgi transport complex (pink), Vesicle tethering complex (blue) and inner MT membrane protein complex (green). Red node indicates SEC22B. Disconnected nodes are not shown. Right panel shows zoom of area with high SNARE complex protein density. E) Western blot analysis of STX5 in input and pulldown (PD) samples from HUVECs expressing mEGFP or mEGFP-SEC22B. B: beads.

To visualize protein interactions a STRING network was generated based on high confidence interactions, showing two major clusters representing proteins that are part of the inner MT membrane protein complex or a SNARE complex, with SEC22B as part of the latter (**Figure 1D**). Among the significant hits were many SNAREs, including STX5 and GOSR2 (Golgi SNAP Receptor Complex Member 2), which are part of the anterograde SNARE complex, and STX18, which facilitates retrograde trafficking.^29^ Another protein complex represented in the STRING analysis comprises components of the vesicle-tethering NBAS, RINT1, ZW10 (NRZ) complex (i.e. NAPA, NBAS, SCFD1, SCFD2, ZW10, C19orf25), which regulates SNARE complex formation of incoming vesicles on the ER membrane.^30^ In addition, members of the conserved oligomeric Golgi (COG) complex were identified (i.e. COG1-6, COG8), suggesting a link between SEC22B and the intra-Golgi vesicle membrane tethering complex.^31,32^ A last notable hit in the SEC22B interactome screen is GBF1, which was previously implicated in ER-Golgi transport of VWF.^15^

The SEC22B interactome analysis has uncovered a large protein network containing both known and novel candidates of protein trafficking in the endothelial biosynthetic pathway. Based on interaction score and the number of edges directed to SEC22B in the STRING analysis, we selected one of the most prominent hits within the SNARE cluster, STX5, for further follow-up. STX5 takes part in both anterograde ER-to-Golgi trafficking and in retrograde intra-Golgi trafficking.^29^ We validated the interaction between STX5 and GFP-SEC22B by immunoblotting for co-precipitated endogenous STX5 (**Figure 1E**), which is expressed as a long (~40 kDa; STX5L) and a short (~34 kDa; STX5S) isoform, resulting from an alternative start codon.^33^ Furthermore, immunofluorescent STX5 staining in HUVECs showed a perinuclear localization, overlapping with the *trans*-Golgi network marker TGN46 (**Figure S1A**). SEC22B also concentrated in the Golgi area, but was additionally observed in small punctae localized in a wider area around the nucleus (**Figure S1B**), most likely representing ER exit sites and trafficking vesicles.^34^ Collectively, these observations suggest that STX5 and SEC22B interact at the Golgi apparatus.

### STX5 and SEC22B depletion in ECs induces unique and shared whole proteome alterations

To study role of STX5 in ECs, we silenced its expression by stable expression of shRNAs targeting STX5. We determined knockdown efficiency of five short hairpin (sh)RNAs targeting STX5 in comparison to a non-targeting shRNA control (shCTRL). Two shRNAs (shSTX5_59826 and shSTX5_59827) efficiently targeted both STX5 isoforms (**Figure S2A,B**).

To assess the impact of STX5 silencing in an unbiased manner, we explored differences on the proteomic level between shSTX5 and shSEC22B compared to shCTRL and untransduced cells, to determine the similarities and differences between silencing either of the two interacting partners in ECs. Samples from the same condition clustered together in the principle component analysis (PCA), showing minimal variability between replicates (**Figure 2A**). After confirming knockdown of SEC22B and STX5 (**Figure 2B, Figure S3**), we analyzed the significantly changed proteins (**Table S5**). Analysis of proteomic alterations between the shSTX5 and shSEC22B conditions revealed 48 overlapping proteins, which included VWF and angiopoietin-2 (Ang-2), suggesting that STX5- and SEC22B-dependent intracellular trafficking regulate the transport of multiple WPB cargo proteins (**Figure 2C**). Moreover, SEC22B depletion rendered a total of 176 unique significantly changed proteins, whereas in shSTX5 cells only 59 proteins were altered.

**Figure 2.**
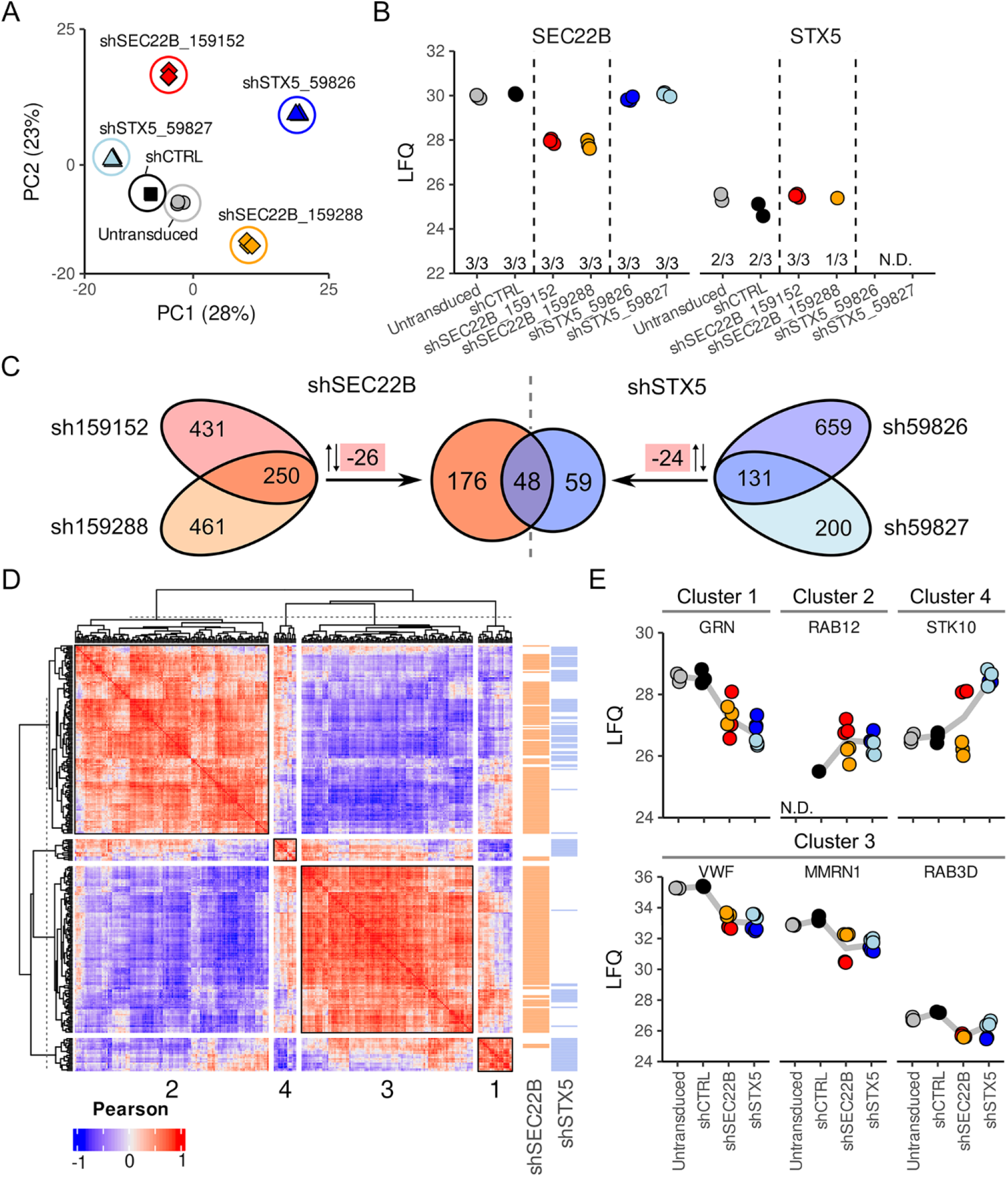
Whole proteome analysis reveals clusters of differentially regulated proteins in shSTX5 and shSEC22B endothelial cells. A) Principle component analysis (PCA) of analyzed samples. B) Label-free quantification (LFQ) plot showing MS-based SEC22B and STX5 protein levels. C) Venn diagrams showing overlap between individual shRNAs and number of shared proteins between shSEC22B and shSTX5 within differentially regulated proteins compared to shCTRL and untransduced conditions. Red boxes show the amount of shared proteins of which the direction of regulation were opposed (up vs. down). D) Correlation heatmap showing Pearson coefficients of regulated proteins separated in 4 clusters. Row annotation indicates if a protein was regulated by SEC22B (orange), STX5 (blue) or both. E) LFQ plots of cluster members. N.D.: not detected.

To further examine these hits and assess protein co-regulation, we generated a coexpression heatmap based on Pearson’s correlations, which visualized 4 differentially regulated protein clusters (**Figure 2D**). These clusters revealed proteins that were mainly driven by STX5 (clusters 1 and 4) or shared between SEC22B and STX5 (cluster 2). Further inspection of cluster 3 highlighted the presence of WPB proteins, as illustrated by the decrease in intracellular VWF levels in both shSTX5 and shSEC22B, and in addition a reduction in Ang-2 and multimerin 1 (MMRN1) (**Figure 2E**). Rab3D and VAMP3 that have been previously implicated in WPB formation and exocytosis^23,35^ were only significantly downregulated in shSEC22B cells, but their expression levels appeared to be lower in shSTX5_59826 as well.

In summary, STX5 and SEC22B depletion cause overlapping proteomic changes, however, STX5 knockdown also induces unique alterations in the proteome, highlighting the importance of STX5-mediated protein transport.

### STX5 silencing results in altered WPB length and loss of the Golgi architecture

To address a potential role of STX5 in WPB biogenesis we investigated WPB and Golgi morphology in shSTX5 and shCTRL HUVECs. STX5 knockdown was validated by Western blot and (nearly) absent immunofluorescent staining upon STX5 knockdown (**Figure 3A,B**). Since shSTX5_59826 yields the most efficient and consistent knockdown, we selected this shRNA for further experiments. To confirm that the effects were specific for STX5 knockdown, key experiments were also performed with shSTX5_59827. VWF staining revealed that upon STX5 silencing the characteristic elongated shape of WPBs was lost, whilst the VWF that was present concentrated in spherical granules mostly found in the perinuclear area (**Figure 3C**). Assessment of immunogold-labeled VWF by electron microscopy (EM) confirmed that these VWF-positive structures were indeed short WPBs (**Figure S4**). Quantitative analysis of the size of VWF positive structures revealed that WPB length was drastically decreased after depletion of STX5 (**Figure 3D**). We also observed a concomitant disintegration of the Golgi; while TGN46 staining showed a compact, continuous ribbon structure in shCTRL cells, shSTX5 cells showed TGN46 staining on dispersed, unlinked structures (**Figure 3C**). Quantification revealed that in the shCTRL condition approximately 70% of the cells contained a continuous, compact Golgi, but the vast majority (~90%) of Golgi’s in shSTX5 cells appeared fragmented and dispersed (**Figure 3E;** examples of Golgi states shown below). Co-staining of TGN46 with the *cis*-Golgi marker GM130 revealed a similar dispersed phenotype for the *cis*-Golgi in shSTX5 cells, suggesting that the entire Golgi architecture is affected by STX5 depletion (**Figure S5**). These dispersed Golgi’s in shSTX5 cells produced shorter WPBs, indicating that WPB length is dependent on extended Golgi ribbon organization (**Figure 3F**), in agreement with previous literature.^13,16^ Interestingly, SEC22B was no longer apparent on the dispersed Golgi’s, and instead a prominent staining surrounding the nucleus was observed (**Figure S6**). Together these results indicate that STX5 is needed for the formation of elongated WPBs, by controlling the maintenance of extended Golgi stacks that allow for loading of multiple VWF quanta into newly forming WPBs.

**Figure 3.**
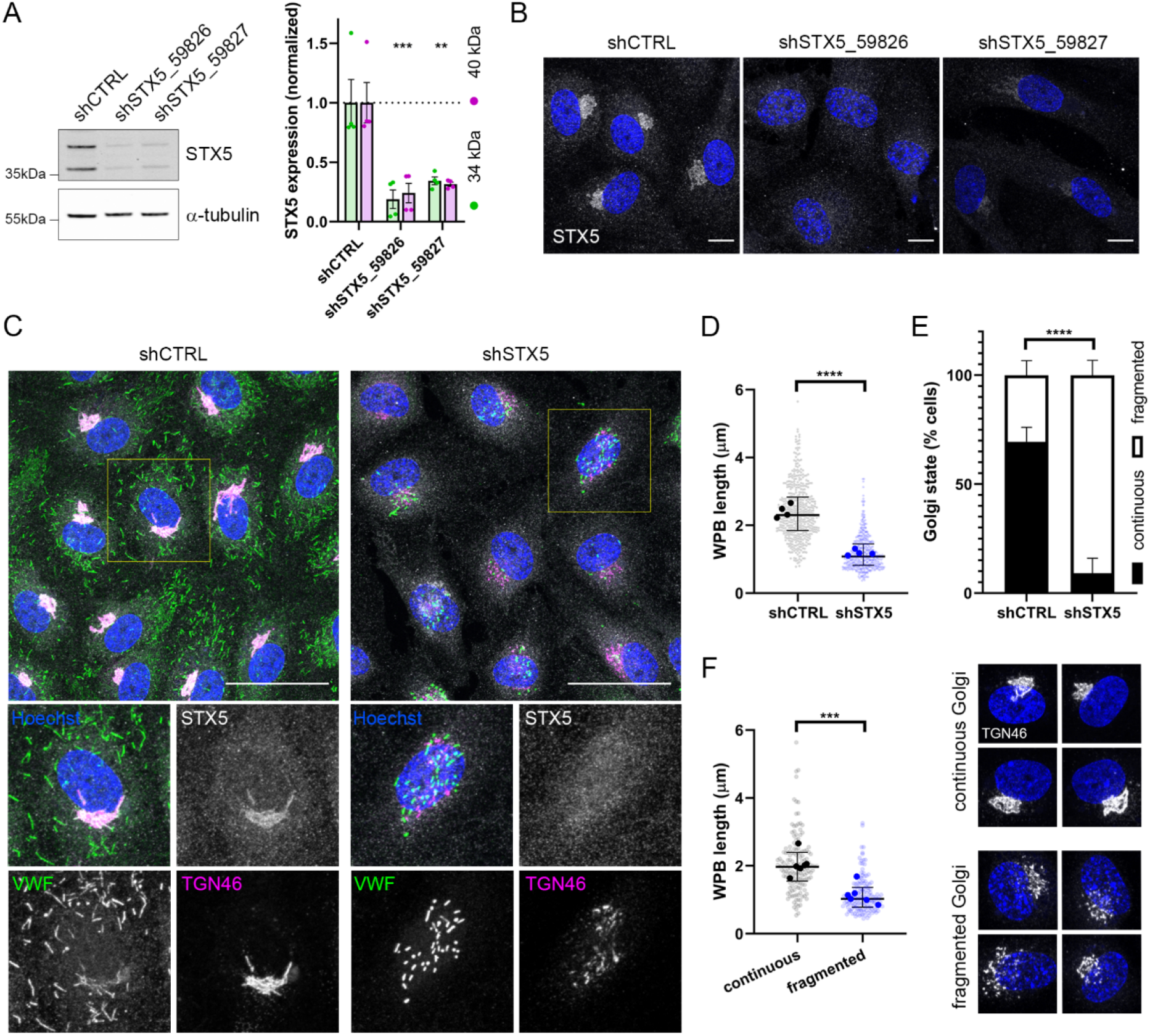
STX5 downregulation results in decreased WPB length and extensive Golgi fragmentation. A) Western blot analysis of STX5 in shCTRL and shSTX5 transduced HUVECs (α-tubulin as loading control) and quantification of STX5 knockdown efficiency (mean±SEM, n=4, t-test, **P<0.01, ***P<0.001). B) Immunofluorescent staining of STX5 (gray) and nuclei (blue) in shCTRL and shSTX5 cells (scale bar: 10 μm). C) VWF (green), TGN46 (magenta), STX5 (gray), and nuclei (Hoechst; blue) in shCTRL and shSTX5 cells (scale bar: 40 μm). Boxed areas are magnified below (individual channels in grayscale). D) WPB length in shCTRL and shSTX5 cells measured in micrometers (μm). Each transparent dot represents a WPB, average length of each biological replicate is shown in fill color (median±IR, n=4, t-test, ****P<0.0001). E) The percentage of cells containing a continuous or fragmented Golgi in shCTRL and shSTX5 cells (median±IR, shCTRL n=5, shSTX5 n=7, t-test, ****P<0.0001). Examples of continuous and fragmented Golgi’s stained with TGN46 (gray) staining are shown below. F) WPB length in shSTX5 cells with continuous versus fragmented Golgi (median±IR, n=6, t-test, ***P<0.001).

### No rough ER dilation due to VWF retention upon STX5 silencing

We used transmission electron microscopy (TEM) to further evaluated the morphology of WPBs, Golgi, and the ER (**Figure 4A**). Similar as earlier noted by light microscopy, the length of the WPBs (**Figure 4A**; WPBs indicated by cyan overlay) was reduced upon STX5 silencing (**Figure 4B**). Golgi ribbon structures arranged in compact stacks were clearly identifiable in shCTRL cells, whereas shSTX5 cells contained widely dispersed Golgi fragments (**Figure 4A**; Golgi indicated by green overlay), which were a challenge to locate despite using the centriole as a reference point (**Figure 4A**; centrioles indicated by magenta overlay). As a consequence, the majority of shSTX5 cells that were analyzed by TEM seemingly lacked elongated Golgi stacks entirely (**Figure 4C**). These observations, along with IF microscopy images showing an altered morphology of the Golgi apparatus in STX5 deficient cells (**Figure 3C, Figure S5**), support the emerging concept that formation of elongated WPBs depends on the presence of a TGN with extended ribbons to package multiple VWF quanta in nascent WPBs.^13,14^

**Figure 4.**
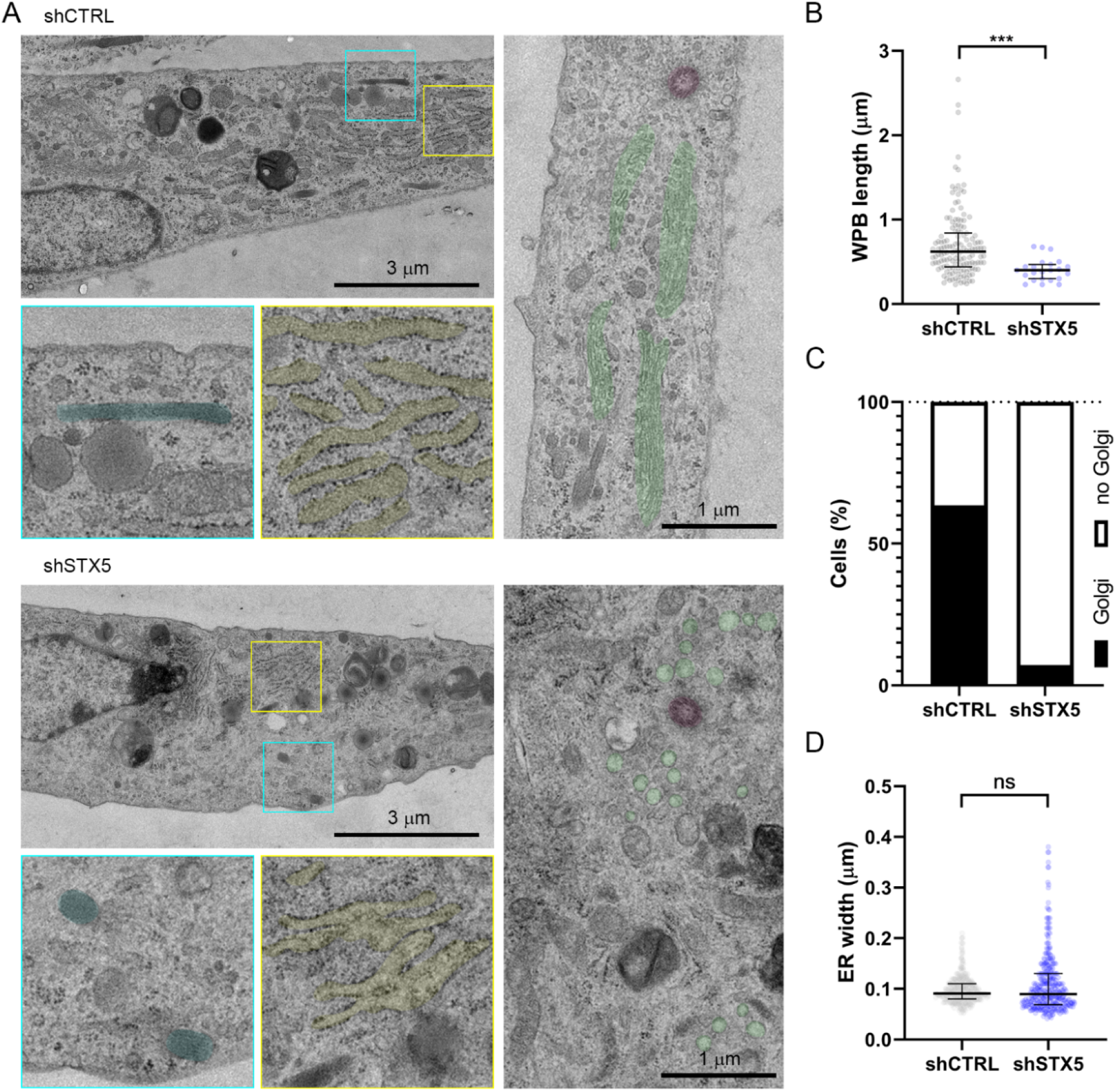
STX5 depletion induces morphological changes in Golgi and rough ER. A) Transmission electron microscopy images of shCTRL and shSTX5 cells. Boxed regions are magnified below outlined with corresponding colors. WPBs and rER sheets are highlighted with cyan and yellow overlays, respectively. In the image on the right, Golgi segments are highlighted in green and centrioles in magenta. Scale bars represent 3 μm and 1 μm as indicated. B) WPB length measured in TEM images (median±IR, t-test, ***P<0.001). C) The percentage of cells with Golgi ribbons versus no Golgi ribbons. D) The rER sheet width measured in shCTRL and shSTX5 cells (median±IR, n=3, t-test, ns: not significant).

We previously uncovered that SEC22B depletion causes retention of VWF inside the ER lumen and massive dilation of the rough ER (rER) as the SEC22B-dependent anterograde trafficking pathway to the Golgi is blocked.^16^ STX5 depletion also caused some retention of VWF in the ER as judged by the overlap of a perinuclear pool of VWF with the rER marker Protein Disulfide Isomerase A3 (PDI) (**Figure S7**). Morphological analysis of the ER revealed that STX5 silencing did not cause VWF retention to the same extent as observed previously in SEC22B depleted cells,^16^ as no large round dense (VWF positive) structures were detected inside the ER lumen (**Figure 4A**; rER sheets indicated with yellow overlay). Modest dilation of the rER was occasionally observed upon STX5 knockdown (**Figure S8**; rER sheets indicated with yellow overlay), but on average the luminal width of rER cisternae was not significantly altered in the shSTX5 cells (**Figure 4D**). Overall, these results indicate that while STX5 and SEC22B depletion have a comparable impact on WPB and Golgi morphology, the increase of ER volume to accommodate accumulating (secretory) proteins is unique to SEC22B knockdown.

### Characteristic WPB markers colocalize with VWF positive structures in shSTX5 cells

Since WPBs contain an array of proteins besides VWF,^8^ we addressed whether their localization to WPBs depends on STX5 by analyzing several WPB cargo and WPB membrane associated proteins using immunofluorescence microscopy. The soluble angiogenic mediator Ang-2 and transmembrane adhesion receptor P-selectin (also referred to as CD62P) are sorted to the WPB during biogenesis at the TGN.^36–38^ Confocal imaging showed that Ang-2 and P-selectin both localized at WPBs in both shCTRL and shSTX5 cells, implying that trafficking of these proteins from the Golgi to their storage compartment continues despite the loss of elongated Golgi architecture upon STX5 silencing (**Figure 5A,B**). We also investigated localization of two late-stage WPB markers: Rab27A and CD63, each representative of a separate route for post-Golgi protein arrival to WPBs. Rab27A, an established maturation marker that is mobilized from the cytosol and recruits essential exocytosis machinery to the WPB membrane,^39–41^ was present on the small WPBs in STX5 depleted ECs (**Figure 6A**). CD63, an integral membrane protein that reaches the WPBs from late endosomes via an AP-3-dependent mechanism,^20,42,43^ was observed on spherical compartments, likely endosomes, and elongated WPBs in shCTRL cells as well as rounded WPBs in shSTX5 cells, indicating that its recruitment from endosomes was not impaired by STX5 depletion (**Figure 6B**). Thus, localization of a selection of established WPB markers remains unchanged after STX5 silencing, indicating that trafficking of cytosolic-, Golgi- and endosome-derived cargo is not impaired.

**Figure 5.**
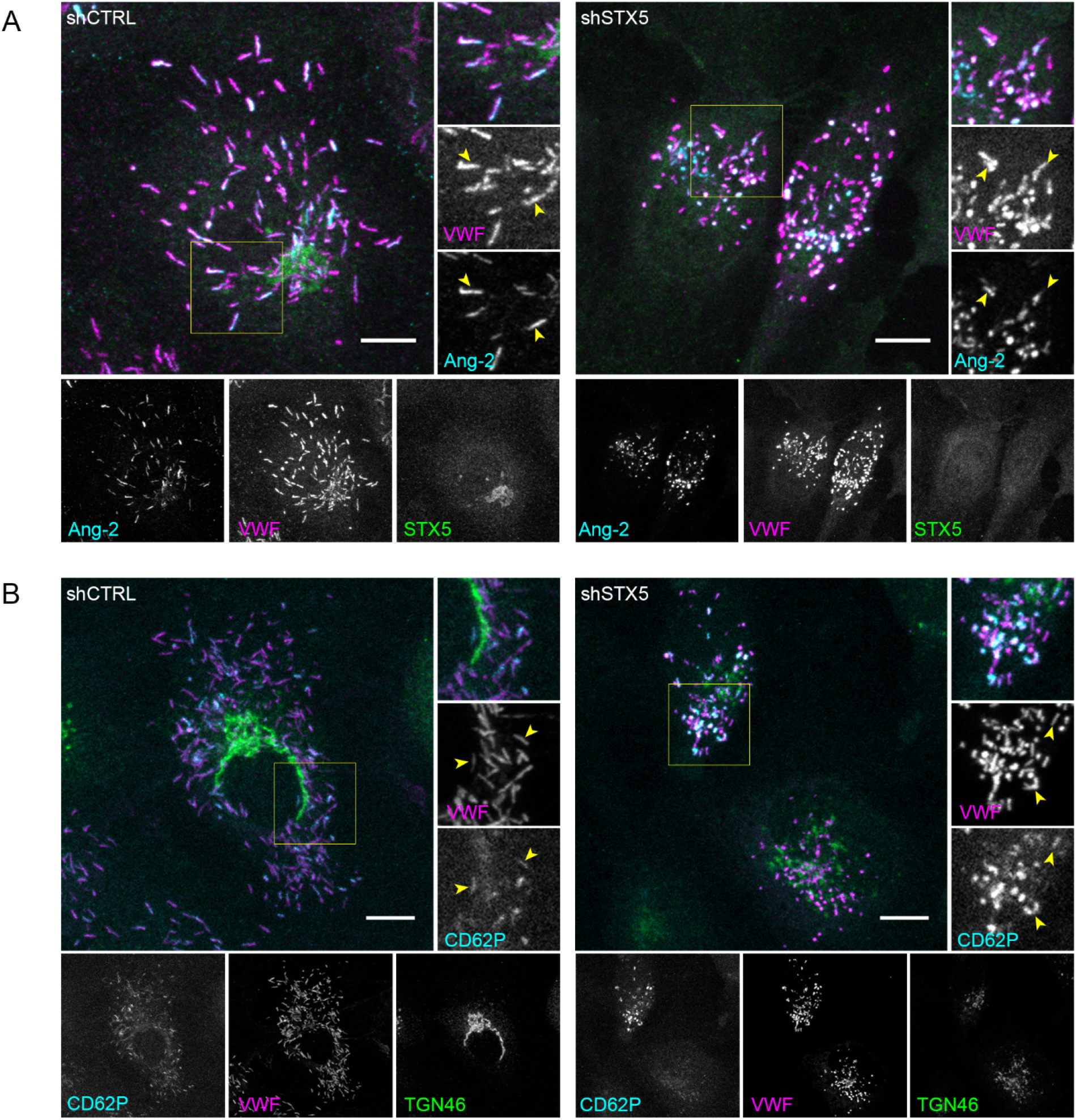
WPB cargo proteins Ang-2 and P-selectin localize to small WPBs in STX5 depleted cells. A) Immunofluorescent staining of Angiopoietin 2 (Ang-2, cyan), VWF (magenta), and STX5 (green) and B) P-selectin (CD62P, cyan), VWF (magenta), and TGN46 (green) in shCTRL and shSTX5 HUVECs (scale bar: 10 μm). Individual channels are shown below in gray scale. Boxed areas are magnified on the right. Yellow arrowheads indicate Ang-2 and P-selectin positive WPBs in (A) and (B) respectively.

**Figure 6.**
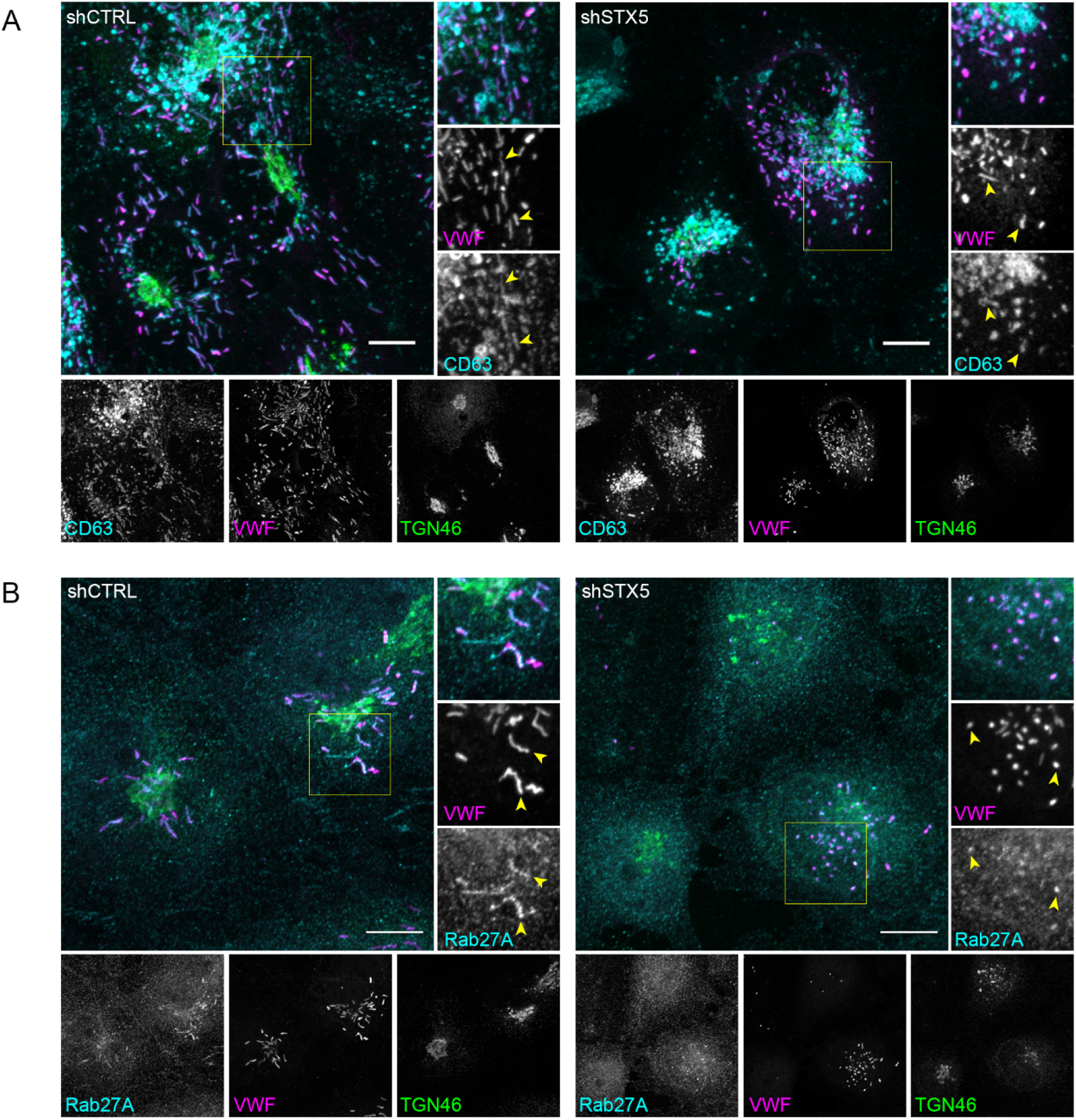
Normal recruitment of maturation marker Rab27A and endosome-derived CD63 to small WPBs upon STX5 knockdown. A) Immunofluorescent staining of Rab27A (cyan), VWF (magenta), and STX5 (green) and B) P-CD63 (cyan), VWF (magenta), and TGN46 (green) in shCTRL and shSTX5 HUVECs (scale bar: 10 μm). Individual channels are shown below in gray scale. Boxed areas are magnified on the right. Yellow arrowheads indicate Rab27A and CD63 positive WPBs in (A) and (B), respectively.

### Short WPBs in STX5 depleted cells are secretion-incompetent

A previous report has shown that WPB size is correlated to the length and adhesive properties of VWF strings.^11^ Since loss of STX5 expression results in shorter WPBs, we hypothesized that stimulated release of these WPBs results in formation of shorter VWF strings. To test this we perfused ECs under 2.5 dynes/cm^2^ flow conditions with histamine to induce WPB exocytosis and a fluorescent VWF antibody to detect VWF strings. A large number of VWF strings with varying lengths were formed on the surface of shCTRL cells, but strikingly no VWF string formation was seen for shSTX5 cells (**Figure 7A**). Confocal analysis of unstimulated conditions revealed that small WPBs were present in shSTX5 cells, predominantly localized in the perinuclear area (**Figure S9**). However, the absence of VWF string formation indicates these WPBs are stimulated secretion-incompetent.

**Figure 7.**
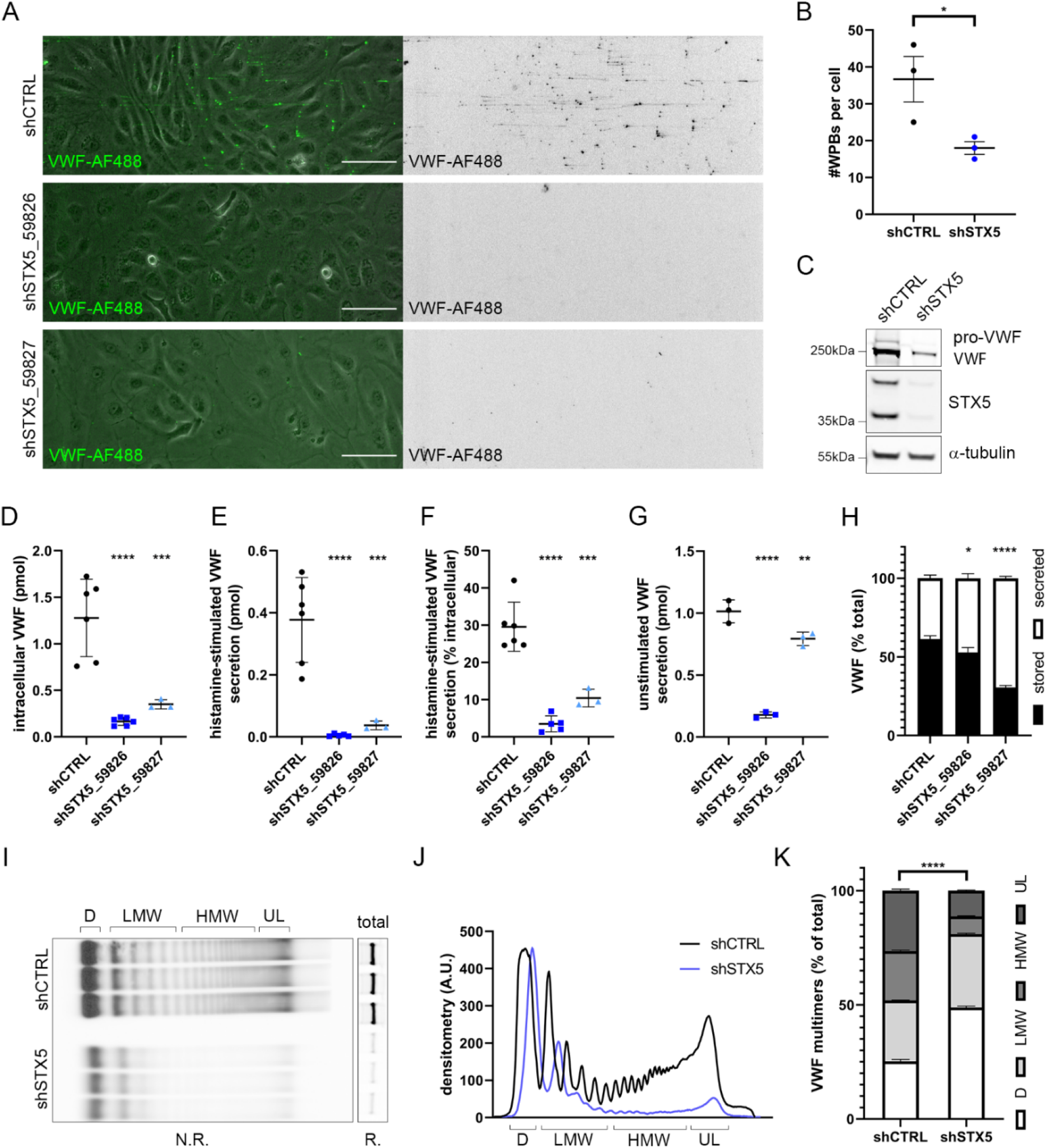
STX5 silencing impairs stimulated VWF secretion. A) VWF string assay with shCTRL and shSTX5-transduced HUVECs following stimulation with 100 μM histamine. Extracellular VWF is shown in green in differential interference contrast (DIC) and fluorescence overlay images and black in inverted fluorescence images below (scale bar: 100 μm). B) The average number of WPBs per cell in shCTRL and shSTX5 cells (mean±SEM, n=3, t-test, *P<0.05). C) Western blot analysis of VWF expression in shCTRL and shSTX5 cells (α-tubulin as loading control). D) Intracellular VWF levels in picomoles (pmol) as measured by ELISA in shCTRL and shSTX5 cells. E) 1-hour histamine-induced VWF secretion in pmol and F) calculated percentage of intracellular VWF (of unstimulated cells). G) 48-hour unstimulated VWF release in pmol and H) calculated proportions of secreted versus stored VWF (mean±SEM, n=3-6, one way ANOVA, *P<0.05, **P<0.01, ***P<0.001, ****P<0.0001). I) VWF multimer analysis on media secreted over 24 hours secreted from shCTRL and shSTX5 cells. D: dimer, LMW: Low-molecular weight, HMW: high molecular weight, UL: ultra large. N.R.: non-reducing conditions, R.: reducing conditions. J) Normalized densitometry analysis and (K) quantification of the area under the curve (AUC) of D, LMW, HMW, and UL multimers as a percentage of total AUC (mean±SEM, n=3, twoway ANOVA, ****P<0.0001).

Immunofluorescent analysis of WPBs in shSTX5 and shCTRL cells showed that in addition to size, the average number of WPBs per cell was significantly reduced upon STX5 knockdown (**Figure 7B**). To examine whether the observed decrease in WPB numbers was due to downregulated VWF expression in STX5 depleted cells, we quantified intracellular VWF levels by Western blot and ELISA (**Figure 7C,D**). In line with the reduction of WPBs, intracellular VWF levels were significantly reduced upon STX5 silencing, which could indicate a decreased expression, or an increased degradation or secretion. Consistent with the VWF string assay data histamine-stimulated secretion of VWF was sharply reduced following STX5 silencing, emphasizing that the small WPBs in STX5 knockdown cells are stimulus-insensitive (**Figure 7E,F**). Unstimulated VWF secretion, which is a composite of basal, WPB-derived secretion and constitutive secretion, was decreased, but not proportionate to stored VWF (**Figure 7G,H**). Subsequent VWF multimer analysis of the unstimulated releasates from shCTRL and shSTX5 cells revealed a remarkable change in multimer size composition, with relatively more VWF dimers (D) and low molecular weight (LMW) multimers, but less high molecular weight (HMW) and ultra large (UL) multimers secreted by STX5 depleted cells (**Figure 7I-K**). Notably, shSTX5-derived VWF dimers and LMW multimers showed a slightly reduced mobility in gel electrophoresis, possibly indicating differences in post-translational modification.

Collectively, these data indicate that loss of STX5 has critical consequences for WPB biogenesis, VWF string formation, secretion, and multimerization, indicating that this SNARE protein is a crucial player in the endothelial secretory pathway.

## Discussion

SNARE proteins constitute the key machinery that promotes membrane fusion during vesicle transport.^19^ We have previously identified several SNARE proteins involved in WPB maturation and exocytosis, while recently we have shown the critical role of SEC22B, an ER-to-Golgi SNARE, in VWF trafficking and WPB formation.^16,20,21^ In this study, a MS-based approach elucidated the SEC22B interactome in ECs and identified potential novel regulators of WPB biogenesis. We focused on STX5, a cognate SNARE protein that is primarily found in the Golgi membrane and has been shown to facilitate ER-to-Golgi and intra-Golgi protein trafficking.^29^ By silencing STX5 expression, we discovered a severe defect in WPB biogenesis, accompanied by fragmentation of the Golgi and abrogation of secretagogue-induced VWF release. Our data point to a crucial role for STX5-containing SNARE complexes in the ability of ECs to efficiently store and secrete VWF.

The importance of VWF transport in WPB biogenesis is underlined by previous studies showing that WPB size depends on anterograde VWF delivery and Golgi morphology, and ultimately determines their functionality in hemostasis.^11–13,44^ Recently, novel players have been identified that modulate biosynthetic pathways crucially involved in WPB biogenesis.^15,16^ The Arf GTPases Arf1 and Arf4 and their GEF GBF1 are essential for the formation of WPBs through regulation of membrane fission.^15^ GBF1 deficiency was accompanied by the formation of giant, secretion-incompetent WPBs. Contrarily, v-SNARE SEC22B regulates ER-to-Golgi transport by facilitating vesicle fusion and depletion of SEC22B in ECs blocks anterograde VWF transport and results in fragmented Golgi’s and shorter WPBs.^16^ To this we can now add STX5, the depletion of which shows considerable phenotypic overlap with that of SEC22B deficient cells. These studies indicate that regulators of vesicle fission and fusion are essential components of the trafficking pathways utilized for VWF multimer assembly as well as WPB biogenesis. Interestingly, GBF1 was also a hit in our SEC22B interactome screen, suggesting that there are opportunities for crosstalk between these two pathways.

STX5 is an integral member of the Golgi apparatus and can form complexes with SEC22B, which is localized on ER-derived vesicles, facilitating their fusion with the Golgi.^18,29^ While STX5L localizes predominantly to the ER, STX5S has been shown to be crucial for intra-Golgi traffic, but not Golgi morphology.^45^ We observed that STX5 was mainly present in the Golgi, whereas SEC22B exhibited a broader localization. Knockdown of both STX5 isoforms caused fragmentation of Golgi stacks into dispersed mini-stacks, similar to SEC22B depletion.^16^ Therefore, we hypothesize that Golgi fragmentation is caused by the reduced delivery of membrane or structural proteins that maintain Golgi architecture from anterograde ER-derived vesicle transport. Alternatively, as STX5 has been reported to participate in Golgi reassembly after mitosis,^46^ reduced STX5 levels in ECs could potentially interfere with the post-mitotic Golgi restoration, leading to permanent Golgi dispersal.

Golgi architecture has a large impact on WPB biogenesis and elongation. According to the model first proposed by Ferraro continuous, extended Golgi ribbons allow for the integration of more VWF quanta, resulting in elongated WPBs, while Golgi mini-stacks give rises to shorter WPBs.^13^. In keeping with the VWF quanta model in STX5 depleted cells with dispersed Golgi, similar to shSEC22B knockdown cells, WPB length is significantly reduced.^16^

VWF release can be separated into three modes: constitutive, basal and regulated (stimulus-induced) VWF secretion.^3^ The WPB compartment ensures a rapid delivery of VWF to form strings and initiate hemostasis.^7^ Interestingly, histamine-stimulated VWF secretion was severely affected in STX5 depleted cells, as evidenced by the (nearly) absent string formation and secretion. This could be explained by the strongly depleted stimulus-sensitive compartment, i.e. reduced number of peripheral WPBs, decrease in WPB length and reduced overall VWF expression levels. Recently, it has been shown that ECs control functional responses by size selection of WPBs during exocytosis, with a preference for longer WPBs to undergo exocytosis following stimulation with a subset of secretagogues, in order to control functional responses.^44^ Thus, the relative unresponsiveness to histamine could also be a consequence of the reduced length of WPBs in shSTX5 cells.

Unstimulated VWF release was also decreased upon STX5 knockdown, however, the amount of secreted VWF relative to stored VWF was proportionally increased compared to the control, suggesting that basal secretion continues and is even slightly increased. Moreover, the secreted VWF contained fewer larger multimers, indicating a multimerization defect. The observed shift in size of VWF dimers could perhaps be explained by incomplete glycosylation of VWF as a result of mislocalized glycosylation enzymes that normally reside in the different Golgi stacks of the Golgi.^47^ Therefore, we can hypothesize that an essential role of STX5 is to maintain a continuous Golgi from which properly multimerized VWF with post-translational modifications can be stored in elongated organelles, which have the capacity to mature into a stimulus-sensitive WPB.

Partial quantitative VWF deficiency, known as VWD1, is the most prevalent VWD subtype.^2^ A sizeable number of pathogenic *VWF* mutations in VWD1 lead to reduced plasma levels due to retention within ECs,^48–50^ which is often accompanied by lower WPB numbers and loss of their characteristic elongated morphology.^51,52^ In ~25% of VWD1 patients and in ~60% of individuals with “low VWF” no causal mutation in their *VWF* gene can be detected. It has been postulated that the genetic regulation of VWF levels largely depends on an interplay of several quantitative trait loci that influence VWF synthesis and clearance.^53^ While a large portion of the heritable variation in VWF is still unaccounted for, some of these modifiers include SNARE proteins that can influence WPB exocytosis.^25,26,53,54^ Given that a substantial portion of quantitative VWF deficiencies in the group with *VWF* mutations are characterized by defects in intracellular trafficking of VWF, it is plausible that VWD and low VWF pathogenesis in cases without damaging *VWF* mutations may also be driven by genetic variations in cellular components that control correct progression of VWF through the endothelial secretory pathway. The SEC22B interactome determined in this study uncovered an extensive network consisting of SNAREs and their regulators and components of a number of vesicle tethering complexes that operate at the ER (NRZ complex), Golgi (COG complex) and post-Golgi (exocyst). Further cellular studies are needed to elucidate their importance in VWF trafficking. Potentially, data from such “trafficome” studies can be used to inform genetic and genomic studies that focus on identification of modifiers VWF in patients that suffer from bleeding or thrombosis.

## Acknowledgements

We would like to thank Dr. R.I. Koning and Dr. C.R. Jost for critical reading of the manuscript. Work in our laboratory was funded by grants from the Landsteiner Stichting voor Bloedtransfusie Research (LSBR-1707) and the Dutch Thrombosis Foundation (TSN 2017-01).

## Authorship Contributions

MK, EK, AJH, PB, AAM, FvA, and JO performed research and analyzed data; DG provided vital reagents; MvdB provided vital expertise; MK, EK, CM, JV, and RB designed the research and wrote the paper.

## References

1. Sadler JE. von Willebrand factor assembly and secretion. J Thromb Haemost 2009;7 Suppl 124–7.

2. Leebeek FWG, Eikenboom JCJ. Von Willebrand’s Disease. N Engl J Med 2016;375(21):2067–2080.

3. Lopes da Silva M, Cutler DF. von Willebrand factor multimerization and the polarity of secretory pathways in endothelial cells. Blood 2016;128(2):277–285.

4. Wagner DD, Marder VJ. Biosynthesis of von Willebrand Protein by Human Endothelial Cells. J Biol Chem 1983;258(4):2065–2067.

5. Voorberg J, Fontijn R, Calafat J, Janssen H, van Mourik JA, Pannekoek H. Biogenesis of von Willebrand factor-containing organelles in heterologous transfected CV-1 cells. EMBO J 1993;12(2):749–758.

6. Schillemans M, Kat M, Westeneng J, et al. Alternative trafficking of Weibel-Palade body proteins in CRISPR/Cas9-engineered von Willebrand factor-deficient blood outgrowth endothelial cells. Res Pract Thromb Haemost 2019;3(4):718–732.

7. Schillemans M, Karampini E, Kat M, Bierings R. Exocytosis of Weibel–Palade bodies: how to unpack a vascular emergency kit. J Thromb Haemost 2019;17(1):6–18.

8. McCormack JJ, Lopes da Silva M, Ferraro F, Patella F, Cutler DF. Weibel-Palade bodies at a glance. J Cell Sci 2017;130(21):3611–3617.

9. Valentijn KM, Valentijn JA, Jansen KA, Koster AJ. A new look at Weibel-Palade body structure in endothelial cells using electron tomography. J Struct Biol 2008;161(3):447–458.

10. Berriman JA, Li S, Hewlett LJ, et al. Structural organization of Weibel-Palade bodies revealed by cryo-EM of vitrified endothelial cells. Proc Natl Acad Sci U S A 2009;106(41):17407–17412.

11. Ferraro F, Lopes-da-Silva M, Grimes W, et al. Weibel-Palade body size modulates the adhesive activity of its von Willebrand Factor cargo in cultured endothelial cells. Sci Rep 2016;6(1):32473.

12. Nightingale TD, McCormack JJ, Grimes W, et al. Tuning the endothelial response: differential release of exocytic cargos from Weibel-Palade bodies. J Thromb Haemost 2018;16(9):1873–1886.

13. Ferraro F, Kriston-Vizi J, Metcalf DJ, et al. A two-tier Golgi-based control of organelle size underpins the functional plasticity of endothelial cells. Dev Cell 2014;29(3):292–304.

14. Mourik MJ, Faas FGA, Zimmermann H, Voorberg J, Koster AJ, Eikenboom J. Content delivery to newly forming Weibel-Palade bodies is facilitated by multiple connections with the Golgi apparatus. Blood 2015;125(22):3509–3516.

15. Lopes-da-Silva M, McCormack JJ, Burden JJ, Harrison-Lavoie KJ, Ferraro F, Cutler DF. A GBF1-Dependent Mechanism for Environmentally Responsive Regulation of ER-Golgi Transport. Dev Cell 2019;49(5):786–801.e6.

16. Karampini E, Bürgisser PE, Olins J, et al. Sec22b determines Weibel-Palade body length by controlling anterograde ER-Golgi transport. Haematologica 2021;106(4):1138–1147.

17. Karampini E, Bierings R, Voorberg J. Orchestration of Primary Hemostasis by Platelet and Endothelial Lysosome-Related Organelles. Arterioscler Thromb Vasc Biol 2020;40(6):1441–1453.

18. Brandizzi F, Barlowe C. Organization of the ER–Golgi interface for membrane traffic control. Nat Rev Mol Cell Biol 2013;14(6):382–392.

19. Jahn R, Scheller RH. SNAREs--engines for membrane fusion. Nat Rev Mol Cell Biol 2006;7(9):631–43.

20. Karampini E, Schillemans M, Hofman M, et al. Defective AP-3-dependent VAMP8 trafficking impairs Weibel-Palade body exocytosis in Hermansky-Pudlak Syndrome type 2 blood outgrowth endothelial cells. Haematologica 2019;104(10):2091–2099.

21. Schillemans M, Karampini E, van den Eshof BL, et al. Weibel-Palade Body Localized Syntaxin-3 Modulates Von Willebrand Factor Secretion From Endothelial Cells. Arterioscler Thromb Vasc Biol 2018;38(7):1549–1561.

22. Zhu Q, Yamakuchi M, Lowenstein CJ. SNAP23 Regulates Endothelial Exocytosis of von Willebrand Factor. PLoS One 2015;10(8):e0118737.

23. Pulido IR, Jahn R, Gerke V. VAMP3 is associated with endothelial weibel-palade bodies and participates in their Ca(2+)-dependent exocytosis. Biochim Biophys Acta 2011;1813(5):1038–44.

24. van Breevoort D, Snijders AP, Hellen N, et al. STXBP1 promotes Weibel-Palade body exocytosis through its interaction with the Rab27A effector Slp4-a. Blood 2014;123(20):3185–3194.

25. van Loon JE, Sanders Y V., de Wee EM, Kruip MJH a, de Maat MPM, Leebeek FWG. Effect of genetic variation in STXBP5 and STX2 on von Willebrand factor and bleeding phenotype in type 1 von Willebrand disease patients. PLoS One 2012;7(7):e40624.

26. Van Loon J, Dehghan A, Weihong T, et al. Genome-wide association studies identify genetic loci for low von Willebrand factor levels. Eur J Hum Genet 2016;24(7):1035–1040.

27. Schillemans M, Karampini E, Hoogendijk AJ, et al. Interaction networks of Weibel-Palade body regulators syntaxin-3 and syntaxin binding protein 5 in endothelial cells. J Proteomics 2019;205103417.

28. Ercig B, Graça NAG, Kangro K, et al. N-glycan–mediated shielding of ADAMTS13 prevents binding of pathogenic autoantibodies in immune-mediated TTP. Blood 2021;137(19):2694–2698.

29. Linders PTA, van der Horst C, Ter Beest M, van den Bogaart G. Stx5-Mediated ER-Golgi Transport in Mammals and Yeast. Cells 2019;8(8):780.

30. Tagaya M, Arasaki K, Inoue H, Kimura H. Moonlighting functions of the NRZ (mammalian Dsl1) complex. Front Cell Dev Biol 2014;2(June):25.

31. Willett R, Kudlyk T, Pokrovskaya I, et al. COG complexes form spatial landmarks for distinct SNARE complexes. Nat Commun 2013;4(1):1553.

32. Kudlyk T, Willett R, Pokrovskaya ID, Lupashin V. COG6 Interacts with a Subset of the Golgi SNAREs and Is Important for the Golgi Complex Integrity. Traffic 2013;14(2):194–204.

33. Hui N, Nakamura N, Sönnichsen B, Shima DT, Nilsson T, Warren G. An isoform of the Golgi t-SNARE, syntaxin 5, with an endoplasmic reticulum retrieval signal. Mol Biol Cell 1997;8(9):1777–87.

34. Mancias JD, Goldberg J. The Transport Signal on Sec22 for Packaging into COPII-Coated Vesicles Is a Conformational Epitope. Mol Cell 2007;26(3):403–414.

35. Knop M, Aareskjold E, Bode G, Gerke V. Rab3D and annexin A2 play a role in regulated secretion of vWF, but not tPA, from endothelial cells. EMBO J 2004;23(15):2982–92.

36. Fiedler U, Scharpfenecker M, Koidl S, et al. The Tie-2 ligand Angiopoietin-2 is stored in and rapidly released upon stimulation from endothelial cell Weibel-Palade bodies. Blood 2004;103(11):4150–4156.

37. Bonfanti R, Furie BC, Furie B, Wagner DD. PADGEM (GMP140) is a component of Weibel-Palade bodies of human endothelial cells. Blood 1989;73(5):1109–12.

38. McEver RP, Beckstead JH, Moore KL, Marshall-Carlson L, Bainton DF. GMP-140, a platelet alpha-granule membrane protein, is also synthesized by vascular endothelial cells and is localized in Weibel-Palade bodies. J Clin Invest 1989;84(1):92–9.

39. van Breevoort D, van Agtmaal EL, Dragt BS, et al. Proteomic Screen Identifies IGFBP7 as a Novel Component of Endothelial Cell-Specific Weibel-Palade Bodies. J Proteome Res 2012;11(5):2925–2936.

40. Hannah MJ, Hume AN, Arribas M, et al. Weibel-Palade bodies recruit Rab27 by a content-driven, maturation-dependent mechanism that is independent of cell type. J Cell Sci 2003;116(Pt 19):3939–3948.

41. Kat M, Bürgisser PE, Janssen H, et al. GDP/GTP exchange factor MADD drives activation and recruitment of secretory Rab GTPases to Weibel-Palade bodies. Blood Adv 2021;5(23):5116–5127.

42. Kobayashi T, Vischer UM, Rosnoblet C, et al. The Tetraspanin CD63/lamp3 Cycles between Endocytic and Secretory Compartments in Human Endothelial Cells. Mol Biol Cell 2000;11(5):1829–1843.

43. Vischer U, Wagner D. CD63 is a component of Weibel-Palade bodies of human endothelial cells. Blood 1993;82(4):1184–1191.

44. McCormack JJ, Harrison-Lavoie KJ, Cutler DF. Human endothelial cells size-select their secretory granules for exocytosis to modulate their functional output. J Thromb Haemost 2020;18(1):243–254.

45. Linders PTA, Gerretsen ECF, Ashikov A, et al. Congenital disorder of glycosylation caused by starting site-specific variant in syntaxin-5. Nat Commun 2021;12(1):6227.

46. Rabouille C, Kondo H, Newman R, Hui N, Freemont P, Warren G. Syntaxin 5 is a common component of the NSF- and p97-mediated reassembly pathways of Golgi cisternae from mitotic golgi fragments in vitro. Cell 1998;92(5):603–610.

47. Linders PTA, Peters E, Ter Beest M, Lefeber DJ, van den Bogaart G. Sugary Logistics Gone Wrong: Membrane Trafficking and Congenital Disorders of Glycosylation. Int J Mol Sci 2020;21(13):4654.

48. Goodeve A. Diagnosing von Willebrand disease: genetic analysis. Hematol Am Soc Hematol Educ Progr 2016;2016(1):678–682.

49. de Jong A, Eikenboom J. Von Willebrand disease mutation spectrum and associated mutation mechanisms. Thromb Res 2017;159(July):65–75.

50. Eikenboom J, Hilbert L, Ribba AS, et al. Expression of 14 von Willebrand factor mutations identified in patients with type 1 von Willebrand disease from the MCMDM-1VWD study. J Thromb Haemost 2009;7(8):1304–12.

51. Wang J-W, Bouwens EAM, Pintao MC, et al. Analysis of the storage and secretion of von Willebrand factor in blood outgrowth endothelial cells derived from patients with von Willebrand disease. Blood 2013;121(14):2762–72.

52. Starke RD, Paschalaki KE, Dyer CEF, et al. Cellular and molecular basis of von Willebrand disease: studies on blood outgrowth endothelial cells. Blood 2013;121(14):2773–84.

53. Swystun LL, Lillicrap D. Genetic regulation of plasma von Willebrand factor levels in health and disease. J Thromb Haemost 2018;16(12):2375–2390.

54. Smith NL, Chen M-H, Dehghan A, et al. Novel Associations of Multiple Genetic Loci With Plasma Levels of Factor VII, Factor VIII, and von Willebrand Factor. Circulation 2010;121(12):1382–1392.

