## Supplemental Material for "Syntaxin 5 determines Weibel-Palade body size and Von Willebrand factor secretion by controlling Golgi architecture"

#### Supplementary Methods

##### *Cell culture, transfection and lentiviral transduction*

Pooled, cryo-preserved primary human umbilical vein endothelial cells (HUVECs) obtained from Promocell (Heidelberg, Germany) were cultured in endothelial cell growth medium (EGM-2, Promocell) supplemented with 18% fetal calf serum (FCS, Bodinco, Alkmaar, The Netherlands), 100 U/mL penicillin, and 100 ug/mL streptomycin (P/S, Gibco), henceforth this medium will be referred to as EGM-18. Lentiviral transduction was performed as previously described.[1] Lentiviral vectors were produced in human embryonic kidney 293T (HEK293T) cells (ATCC) using a 3$^{rd}$ generation lentiviral packaging system.

##### *DNA constructs*

The lentiviral LVX-mEGFP-LIC vector has been described.[2] A synthetic human SEC22B fragment was generated by gene synthesis containing all 214 codons of the SEC22B coding sequence, flanked by *Bsr*GI and *Not*I sites, to facilitate cloning in frame behind mEGFP in LVX-mEGFP-LIC. The non-targeting, SEC22B-targeting, and STX5-targeting pLKO.1-puro shRNA vectors were obtained from the MISSION® shRNA library developed by TRC at the Broad Institute (**Table S2**).

##### *Mass spectrometry analysis*

*SEC22B interactome*

HUVECs were seeded in 3 separate wells of 6-well culture plates and lentivirally transduced with mEGFP or mEGFP-SEC22B to enable sample preparation in triplicate for each pulldown condition. After 3 days of confluency, cells were rinsed 2x in PBS and subsequently lysed in mass spectrometry (MS) grade lysis buffer (10 mM Tris-HCl pH 7.5, 150 mM NaCl, 0.5 mM EDTA, 0.5% NP40 (v/v)) supplemented with Halt protease and phosphatase inhibitor cocktail (Thermo Scientific). Lysates were centrifuged for 10 minutes at 16,000g and supernatants were transferred to fresh tubes. Cleared lysates were incubated with GFP-Trap magnetic agarose beads or control (CTRL) beads (ChromoTEK) by rotation for 2 hours at room temperature (RT). For interactome analysis by MS, beads were collected on a magnetic stand and were washed 3 times with wash buffer (10 mM Tris-HCl pH 7.5, 150 mM NaCl, 0.5 mM EDTA) supplemented with Halt protease and phosphatase inhibitor cocktail (Thermo Scientific, Breda, The Netherlands) and two times with 1 ml PBS. Sample preparation, data acquisition and data analysis were performed according to.[3]

*shSEC22B and shSTX5 whole proteome*

HUVECs were seeded in 3 separate wells of 6-well plates and lentivirally transduced with shRNAs targeting SEC22B as described,[4] STX5, or non-targeting control shRNA. Untransduced cells were also included. After reaching confluency cells were lysed, samples were processed for mass spectrometry and analyzed as previously described in.[5]

*MS Data analysis*

MS raw files were processed with MaxQuant 1.6.2.10 using the human Uniprot database (downloaded February 2019).[6] MaxQuant output tables were analyzed using R/Bioconductor (version 4.1.0/3.13),[7] 'reverse', 'potential contaminants' and 'only identified by site' peptides were filtered out and label free quantification values were $\log_2$ transformed. Proteins quantified in 2/3 of an experimental group were selected for further analysis. Missing values were imputed by a normal distribution (width=0.3, shift = 1.8), assuming these proteins were close to the detection limit. Statistical analyses were performed using moderated t-tests in the LIMMA package.[8] A Benjamini-Hochberg adjusted P value <0.05 and absolute $\log_2$ fold change >1 was considered statistically significant and relevant. SNARE-complex annotation were obtained from GO-term 'SNARE complex' (GO:0031201). Overrepresentation analyses were performed using clusterprofiler.[9] Protein interactions with STRING-DB combined scores >0.9 were visualized in Cytoscape v3.8.2.The .raw MS files and search/identification files obtained with MaxQuant have been deposited in the ProteomeXchange Consortium[10] via the PRIDE partner repository with the dataset identifier PXD027516.

***Fluorescence microscopy***

Immunofluorescence (IF) staining and imaging was performed as previously described[5] using a Leica SP8 confocal microscope (Leica, Wetzlar, Germany). Super resolution imaging of ER and Golgi stainings was performed on a Zeiss Airyscan (Zeiss, Breda, The Netherlands). All antibodies used for IF stainings have been specified in Table S1. WPBs were visualized using mouse monoclonal anti-VWF CLB-Rag20[11] or rabbit polyclonal anti-VWF (DAKO, A0058) as indicated in Figure legends. Nuclei were stained using Hoechst nuclear dye (Life Technologies; H-1399). Images were processed and analyzed using ImageJ (https://imagej.nih.gov/ij/). WPB length (major axis of cigar-shaped VWF positive structures) and TGN area (periphery of TGN46 staining) were measured in pixels and automatically converted to actual size in ImageJ.

***VWF string assay***

ECs were first transduced with shCTRL or shSTX5 and 75,000 cells per channel were seeded on day 0 in gelatin-coated 6-channel µ-slides VI 0.4 (#80606, IBIDI). The growth medium was replaced every morning and afternoon. After 4 days the VWF string assay was performed under flow conditions using 2.5 dynes/cm$^2$ and at 37°C. The cells were starved for 5 minutes using M-199 medium (Gibco) supplemented with 1% bovine serum albumin (BSA, Fraction V) and subsequently stimulated with 100 µM histamine (Sigma-Aldrich; H7125) added to the starvation medium for 5 minutes. To visualize VWF strings, an Alexa Fluor 488 (AF488)-conjugated anti-VWF antibody (DAKO), prepared using the Alexa Fluor 488 protein labeling kit (ThermoFisher Scientific, A10235), was added at 2 µM concentration to the medium for 5-10 minutes. Live imaging was performed on an Axiovert widefield microscope (Zeiss) using a 10x objective. Unstimulated conditions were fixed with 4% PFA for 5 minutes at RT. IF stainings for VWF, TGN46, Vascular Endothelial (VE) cadherin, and Hoechst, and confocal imaging were performed as previously described.[5]

##### *Western blotting and VWF multimer analysis*

Western blotting was performed as described.[4] For VWF multimer analysis EC media and lysates, prepared as described in the ''secretion assay'' section above, were loaded on 0.9% ultrapure agarose gels run at 75 V for 5 hours. VWF was reduced using β-mercaptoethanol and transfered onto a nitrocellulose membrane (GE Healthcare) overnight by cappilary suction. After blocking for 1 hour in 10% non-fat dry milk (Blotto, Chem Cruz) in PBS-0.05% Tween-20 (Sigma) and incubation with VWF-HRP (DAKO, #P0226, 1:1000) for 3 hours, the blot was visualized by enhanced chemiluminesence according to manufacturer's protocol (Supersignal West Pico Plus chemilumiescent substrate, ThermoFisher Scientific). Densitometry analysis was performed in ImageJ.

##### *Electron microscopy*

*Transmission electron microscopy*

ECs were processed for transmission electron microscopy essentially as described.[4] Briefly, HUVECs cultured in 10 cm petridishes were fixed (TEM: 1,5% glutaraldehyde (GA), 0,1 M cacodylate buffer, pH 7.4;) for 2 hours at RT, washed three times (0,1 M cacodylate), scraped and pelleted in 2% agar solution. Solidified pellets were cut into 1 x 1 $mm^3$ pieces, dehydrated in an ethanol series followed by a series of EPON (LX11, Leadd) to pure EPON. After polymerizing for 2 days at 70°C, ultrathin sections (80 µm) were cut parallel to the surface of the beem capsules using a Reichert Ultracut S (Leica Microsystems) and contrasted with uranylacetate and lead citrate. Sections were examined on a Fei Tecnai Twin transmission electron microscope (FEI, Eindhoven, Netherlands) at 120 kV, with a Gatan Oneview camera (Gatan, Pleasonton) on binning 2. Overlapping images were stitched together as previously described.[12] Image analysis was performed using Aperio Imagescope (Leica) and colored overlays were made in Adobe Illustrator.

*Immuno electron microscopy*

After fixation (2% PFA, 0.2% GA, 0.1 M PHEM) for 2 hours at RT HUVECs were rinsed with 0.1M PHEM buffer, scraped and pelleted in 10% gelatin. Pieces of the pellet (~0.5-1 $mm^3$) were cryoprotected by immersion in PBS / 2.3 M sucrose solution for 1 hour, mounted on a stub, and snapfrozen in liquid nitrogen. Ultrathin cryosections (90 nm) made with a Leica EM Ultracryotome were collected on grids and incubated with rabbit-anti-VWF (DAKO, 1:1000) followed by 10 nm Protein A gold particles (1:300) in 1% BSA in PBS as described.[13] The sections were embedded (methylcellulose, 0.3 % uranylacetate solution) and examined as described above.

***Data statistical analysis***

Student's t-test or one-way ANOVA with Dunnett's multiple comparisons test comparing to shCTRL were performed using GraphPad Prism 8 (Graphpad, La Jolla, CA, USA). Significance values are specified in the Figures and in Figure legends. Data are shown as mean ± SEM, median ± IR, or as contingency-stacked bar graphs.

#### References


1. Schillemans M, Karampini E, Van Den Eshof BL, et al. Weibel-palade body localized syntaxin-3 modulates von willebrand factor secretion from endothelial cells. Arterioscler Thromb Vasc Biol 2018;38(7):1549–1561.

2. van Breevoort D, Snijders AP, Hellen N, et al. STXBP1 promotes Weibel-Palade body exocytosis through its interaction with the Rab27A effector Slp4-a. Blood 2014;123(20):3185–3194.

3. Schillemans M, Karampini E, Hoogendijk AJ, et al. Interaction networks of Weibel-Palade body regulators syntaxin-3 and syntaxin binding protein 5 in endothelial cells. J Proteomics 2019;205103417.

4. Karampini E, Bürgisser PE, Olins J, et al. Sec22b determines Weibel-Palade body length by controlling anterograde ER-Golgi transport. Haematologica 2020;106(4):1138–1147.

5. Karampini E, Schillemans M, Hofman M, et al. Defective AP-3-dependent VAMP8 trafficking impairs Weibel-Palade body exocytosis in Hermansky-Pudlak Syndrome type 2 blood outgrowth endothelial cells. Haematologica 2019;104(10):2091–2099.

6. Tyanova S, Temu T, Cox J. The MaxQuant computational platform for mass spectrometry-based shotgun proteomics. Nat Protoc 2016;11(12):2301–2319.

7. Huber W, Carey VJ, Gentleman R, et al. Orchestrating high-throughput genomic analysis with Bioconductor. Nat Methods 2015;12(2):115–121.

8. Ritchie ME, Phipson B, Wu D, et al. limma powers differential expression analyses for RNA-sequencing and microarray studies. Nucleic Acids Res 2015;43(7):e47–e47.

9. Yu G, Wang L-G, Han Y, He Q-Y. clusterProfiler: an R Package for Comparing Biological Themes Among Gene Clusters. Omi A J Integr Biol 2012;16(5):284–287.

10. Vizcaíno JA, Deutsch EW, Wang R, et al. ProteomeXchange provides globally

coordinated proteomics data submission and dissemination. Nat Biotechnol 2014;32(3):223–226.

11. Stel H, Sakariassen K, de Groot P, van Mourik J, Sixma J. Von Willebrand factor in the vessel wall mediates platelet adherence. Blood 1985;65(1):85–90.

12. Faas FGA, Avramut MC, M. van den Berg B, Mommaas AM, Koster AJ, Ravelli RBG. Virtual nanoscopy: Generation of ultra-large high resolution electron microscopy maps. J Cell Biol 2012;198(3):457–469.

13. Peters PJ, Bos E, Griekspoor A. Cryo- Immunogold Electron Microscopy. Curr Protoc Cell Biol 2006;30(1):Chapter 4, unit 4.7.

14. Bierings R, Hellen N, Kiskin N, et al. The interplay between the Rab27A effectors Slp4-a and MyRIP controls hormone-evoked Weibel-Palade body exocytosis. Blood 2012;120(13):2757–2767.

**Supplementary Figures**

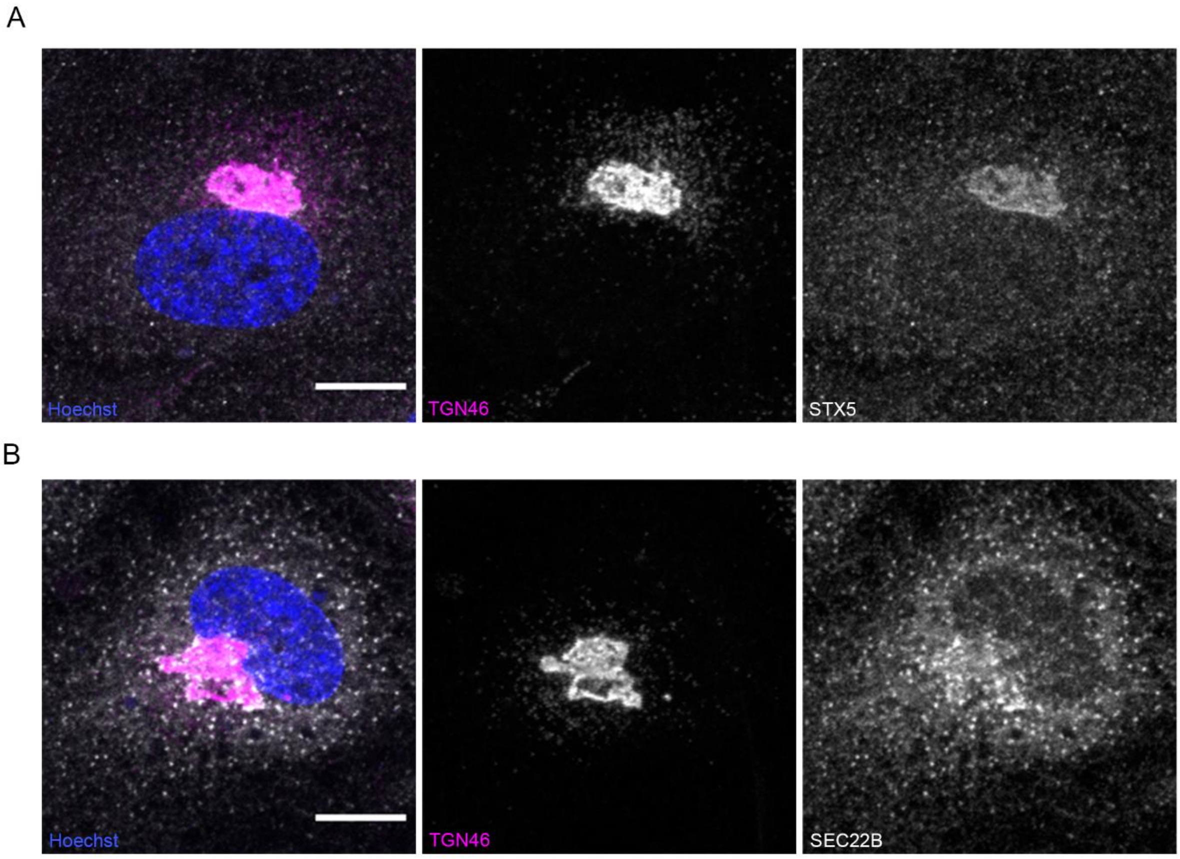


**Figure S1. Endogenous SEC22B and STX5 localization.** Immunofluorescent stainings of (A) STX5 and (B) SEC22B (both in gray) in HUVECs combined with *trans*-Golgi marker TGN46 (magenta) and nuclear staining Hoechst (blue). Scale bars indicate 10 µm.

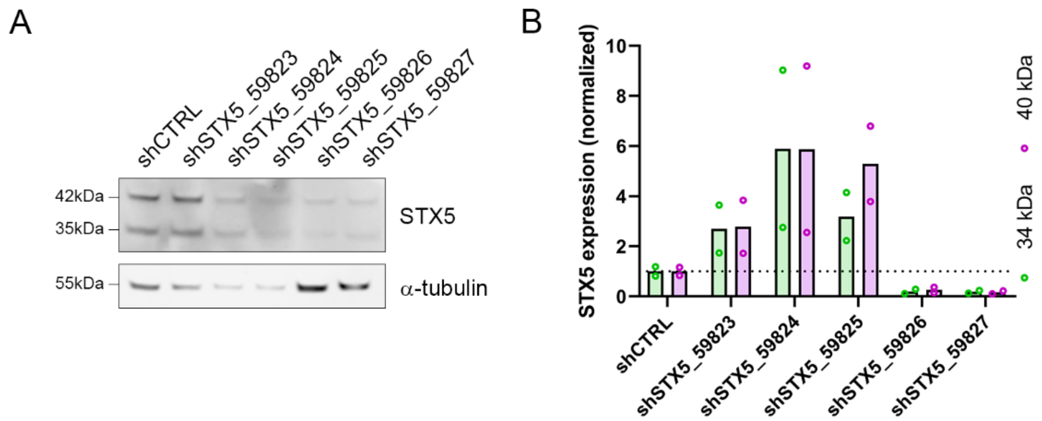


**Figure S2. STX5 knockdown efficiency of single shRNAs.** A) Western blot analysis of STX5 in shCTRL and shSTX5-transduced HUVECs (α-tubulin as loading control). B) Densitometry analysis of 34 and 40 kDa STX5 isoforms normalized to α-tubulin expression (mean, n=2 technical replicates).

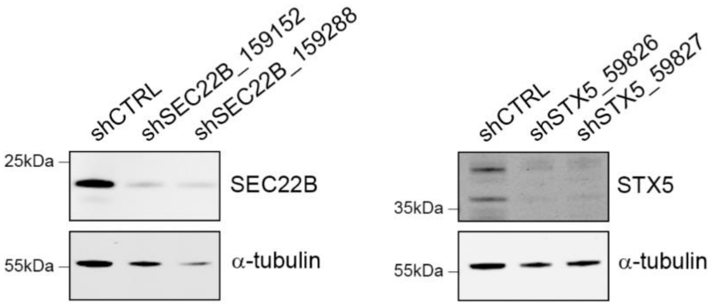


**Figure S3. Efficient STX5 and SEC22B knockdown.** Western blot analysis to confirm SEC22B and STX5 knockdown in SEC22B, respective STX5 shRNA-transduced HUVECs compared to shCTRL (α-tubulin as loading control).

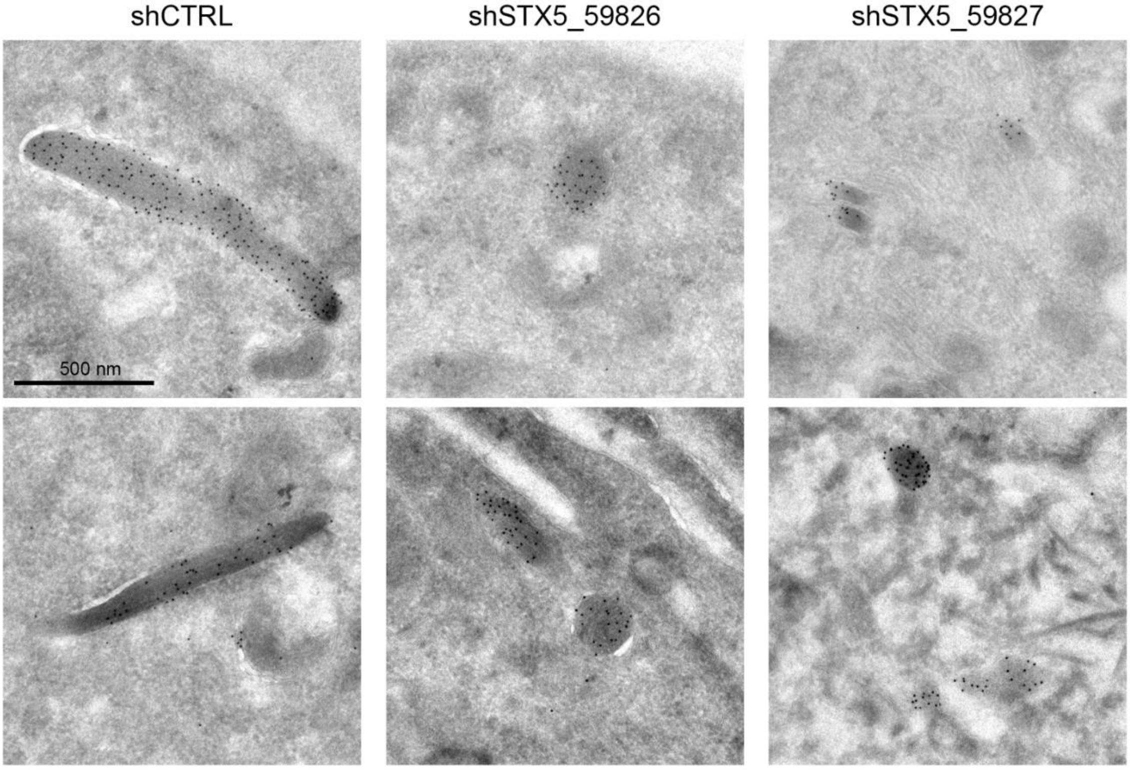


**Figure S4. VWF positive structures in STX5 depleted cells resemble WPBs**. Electron microscopy images of structures containing VWF immuno-gold labeling in shCTRL and shSTX5-transduced HUVECs. (scale bar: 500 nm).

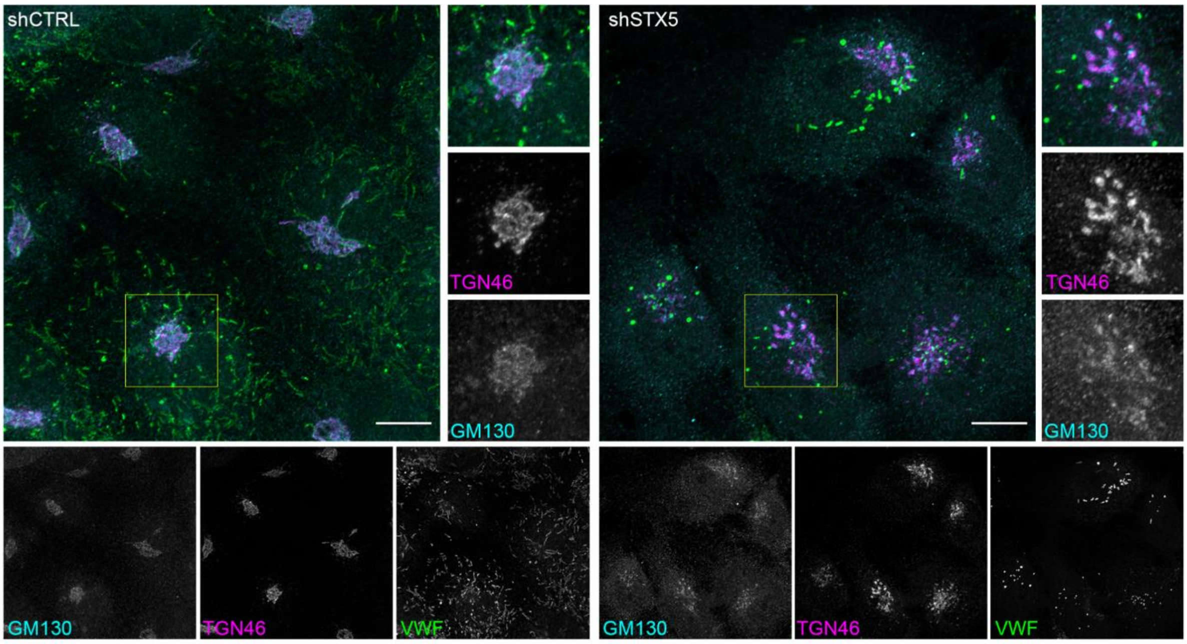


**Figure S5. STX5 knockdown causes fragmentation of *cis*-Golgi.** Immunofluorescent stainings of GM130 (cyan), TGN46 (magenta), and VWF (green) in shCTRL and shSTX5 transduced HUVECs (scale bar: 10 μm). Individual channels are shown below in gray scale. Boxed areas are magnified on the right.

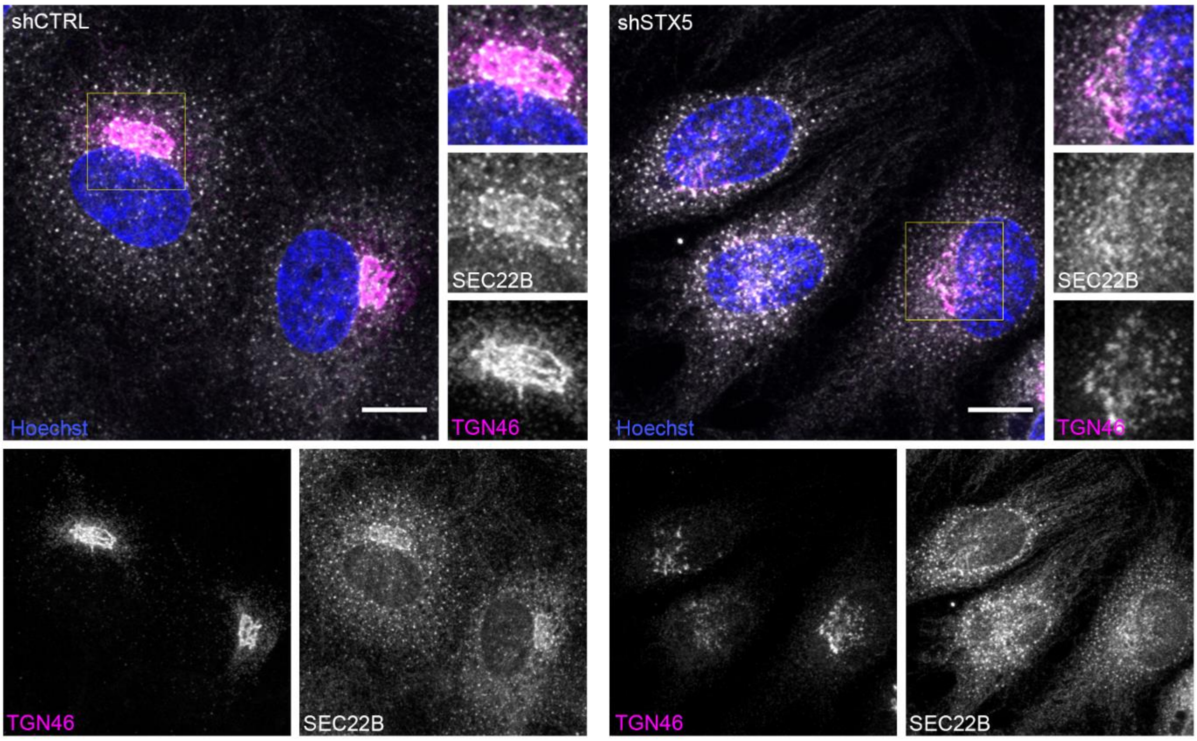


**Figure S6. SEC22B localization is dispersed in STX5 depleted cells.** Immunofluorescent stainings of TGN46 (magenta), SEC22B (gray), and nuclei (Hoechst, blue) in shCTRL and shSTX5 transduced HUVECs (scale bar: 10 μm). Individual channels are shown below in gray scale. Boxed areas are magnified on the right.

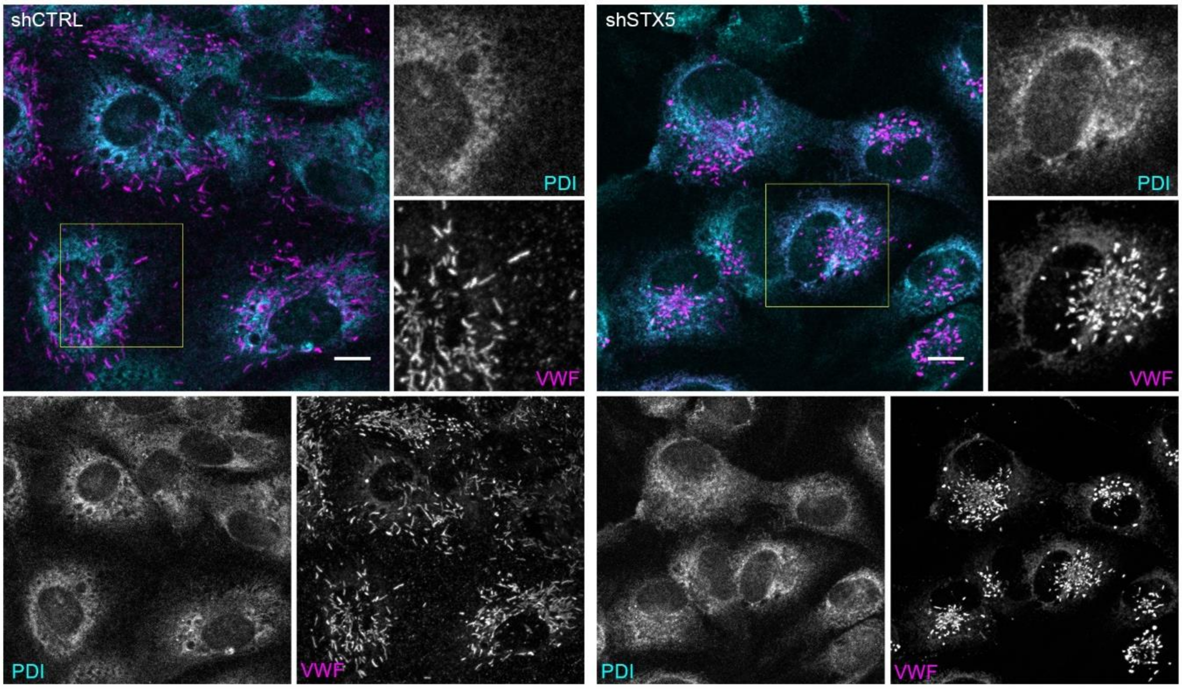


**Figure S7. VWF retention in rough ER.** Immunofluorescent staining of VWF (magenta) and PDI (cyan) in shCTRL and shSTX5 (scale bar: 10 µm). Individual channels are shown below and boxed area are magnified on the right.

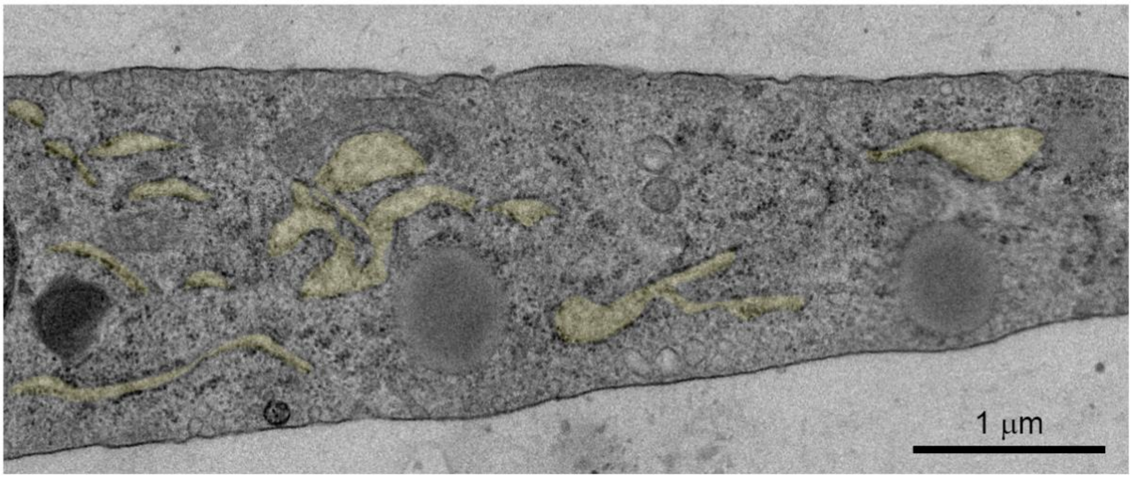


**Figure S8. ER dilation upon STX5 knockdown.** Transmission electron microscopy image of a shSTX5 cell with dilated rER sheets highlighted in yellow (scale bar: 1 µm).

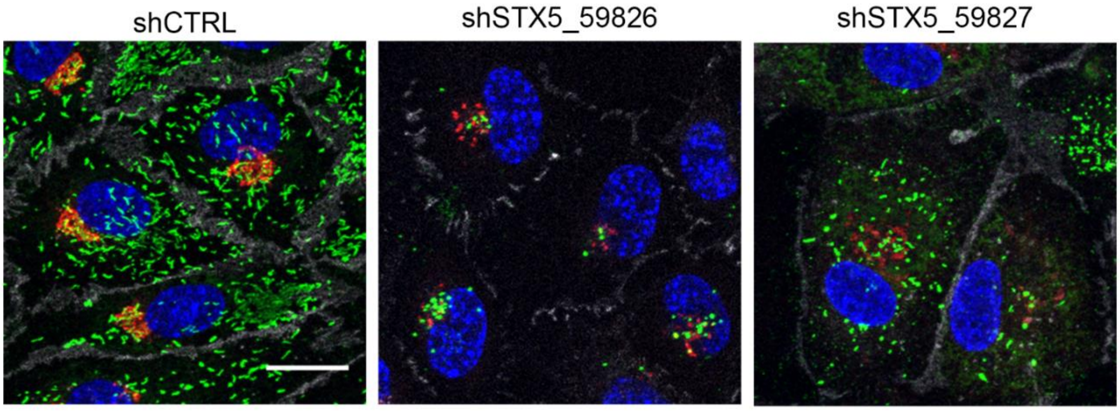


**Figure S9. WPB content in unstimulated cells cultured for VWF string assay.** Immunofluorescent stainings of VWF (green), vascular endothelial (VE) cadherin (gray), TGN46 (red), and nuclei (Hoechst, blue) in shCTRL and shSTX5 transduced HUVECs fixed in IBIDI channel (scale bar: 20 µm).

#### Supplementary Tables

**Table S1.** Antibody list.

| **Target** | **Species (isotype)** | **Company/article** | **Clone/Cat#** | **Application [dilution/conc]** |
|---|---|---|---|---|
| STX5 | Rabbit | Synaptic Systems | 110 053 | IF [1:100]<br>WB [1:1000] |
| SEC22B | Rabbit | Synaptic Systems | 186 003 | IF [1:100] |
| TGN46 | Sheep | Bio-rad | AHP500GT | IF [1:500] |
| GM130 | Rabbit | Cell signaling | D6B1 | IF [1:1000] |
| Calnexin | Rabbit | Abcam | ab22595 | IF [1:100] |
| PDIA3 | Mouse IgG1 | Sigma-Aldrich | CL2444 | IF [1:1000] |
| VE cadherin-AF647 | Mouse IgG1 | BD Pharmigen | 55-7H1 | IF [1:200] |
| Ang-2 | Goat | R&D Systems | AF623 | IF [1:200] |
| CD63 | Mouse IgG1 | CLB | CLB-gran/12 | IF [1:100] |
| CD62P-AF488 | Mouse IgG1 | AbD Serotec | AK-6 | IF [1:100] |
| Rab27A | Rabbit | Bierings et al.[14] | B2423 | IF [1:50] |
| VWF | Rabbit | DAKO | A0085 | IF [1:5000]<br>WB [1:5000]<br>ELISA [6 µg/ml] |
| VWF | Mouse IgG2B | CLB | CLB-Rag20 | IF [1:5000] |
| VWF-HRP | Rabbit | DAKO | A0085 | ELISA [2 µg/ml] |
| α-tubulin | Mouse IgG1 | Sigma | DM1A | WB [1:10000] |
| GFP | Mouse IgG1 | Clontech | JL-8 | WB [1:2500] |

Fluorophore-conjugated secondary antibodies (Molecular Probes) used for immunofluorescence microscopy were purchased from Invitrogen. Secondary antibodies conjugated with infrared dyes (680LT and 800CW) used for immunoblotting were from LI-COR.

**Table S2.** pLKO-shRNA sequences.

| Target | TRCN | Short hairpin sequence (5'→3') |
|---|---|---|
| Non-targeting | shCTRL-C002 | CAACAAGATGAAGAGCACCAA |
| SEC22B | TRCN0000159152 | GCCATCAATGAGATTTAACTT |
| SEC22B | TRCN0000159288 | GCCACAATTTGCTAACATTTA |
| STX5 | TRCN0000059823 | CCAGACAAATAAGCCAGCTTT |
| STX5 | TRCN0000059824 | GCAGTCGAAACTGGCTTCTAT |
| STX5 | TRCN0000059825 | CCATTCAGAGATCCTCAAGTA |
| STX5 | TRCN0000059826 | CCTTAGCAACACATTTGCCAA |
| STX5 | TRCN0000059827 | GCAGAACATTGAGTCGACAA |

### Table S3. Significant hits in endothelial SEC22B interactome screen.

| Genes | LFQ intensity EGFP_A | LFQ intensity EGFP_B | LFQ intensity EGFP_C | LFQ intensity EGFP-SEC22B_A | LFQ intensity EGFP-SEC22B_B | LFQ intensity EGFP-SEC22B_C | adjusted P-value |
|---|---|---|---|---|---|---|---|
| PCDH1 | NA | NA | NA | 32,17465327 | 32,02696559 | 32,07794632 | 1,2974E-08 |
| SEC22B | 26,93988513 | 27,10599496 | 27,47119269 | 36,12694078 | 36,36647366 | 36,25501042 | 2,62637E-07 |
| ATP2B4 | 20,07300136 | 20,52238613 | 21,74345057 | 29,67099479 | 30,05749858 | 28,89844889 | 8,83709E-06 |
| CHCHD3 | 20,91232509 | 20,86166246 | 20,45030512 | 28,49564183 | 28,2008823 | 28,27350985 | 2,8779E-07 |
| ITGA6 | 21,02803284 | 21,22911218 | 21,21477919 | 27,83647409 | 28,2292644 | 27,88221194 | 5,31367E-06 |
| STX18 | 21,50013581 | NA | 21,54703195 | 27,94720663 | 27,78269293 | 28,20820481 | 5,3174E-06 |
| NBAS | 24,6056902 | 23,15343019 | 23,60557718 | 30,66517712 | 30,55004584 | 29,63792644 | 2,59233E-05 |
| ATP2B1 | NA | NA | NA | 26,57402467 | 26,54608702 | 26,45241508 | 9,65323E-06 |
| SNAP29 | NA | NA | NA | 27,26820902 | 27,61224455 | 27,49868806 | 9,65323E-06 |
| BNIP1 | 22,54578369 | 22,20484573 | NA | 28,37633501 | 28,37951963 | 28,45865455 | 5,3174E-06 |
| SPNS1 | NA | NA | NA | 27,08080761 | 27,23343054 | 27,06907209 | 9,04492E-07 |
| ATP1A1 | 24,28362083 | 24,2584654 | 24,07136547 | 30,37368269 | 30,46031568 | 30,37160717 | 6,20406E-07 |
| SCFD2 | 21,19321973 | 21,80313627 | 23,29151135 | 27,95143725 | 28,33636953 | 28,37807295 | 2,21687E-05 |
| PLXND1 | 20,90470223 | NA | 21,64803276 | 27,1773114 | 26,86466904 | 27,77950286 | 2,59233E-05 |
| ANO6 | NA | NA | 23,29389818 | 28,47823156 | 28,51133705 | 28,37276983 | 7,70088E-06 |
| RINT1 | NA | NA | NA | 27,17054645 | 27,07152914 | 27,13613971 | 1,61928E-05 |
| NDUFA8 | 22,19700547 | 21,8347997 | 23,06223591 | 28,19568271 | 28,34375732 | 28,19591733 | 8,83709E-06 |
| TMX1 | NA | NA | NA | 26,46699948 | 26,65799111 | 26,66451608 | 7,70088E-06 |
| NAPA | NA | NA | NA | 26,54364088 | 26,72741343 | 26,74624716 | 7,70088E-06 |
| RMDN3 | 22,83190408 | 21,60720387 | NA | 27,14209381 | 28,61754375 | 27,24417929 | 0,000127272 |
| TM9SF4 | NA | NA | NA | 27,00077664 | 27,53875091 | 26,90770304 | 8,83709E-06 |
| ABCD3 | 21,70476317 | 21,13257504 | 21,98492323 | 27,61203359 | 27,24590256 | 26,93044102 | 8,99292E-06 |
| C19orf25 | NA | NA | 23,74108591 | 27,82765874 | 27,75015075 | 27,84685044 | 6,13362E-05 |
| NDUFB8 | NA | NA | NA | 26,10403608 | 26,3965635 | 26,39610615 | 7,70088E-06 |
| TBC1D9B | 20,57283713 | 21,91440703 | 20,36858319 | 26,17656976 | 27,00699424 | 26,10203448 | 2,33537E-05 |
| ITGB1 | NA | 21,12993699 | 21,90179643 | 26,62212535 | 26,81311084 | 27,28859266 | 5,29522E-06 |
| USE1 | NA | NA | 23,27241565 | 27,33449535 | 26,9627873 | 26,2114484 | 0,000427673 |
| ATP13A1 | 22,34835428 | 22,6484278 | 22,23928852 | 27,76033976 | 27,82559809 | 27,70596032 | 5,39835E-05 |
| CMTM6 | NA | NA | NA | 26,2247367 | 26,4482614 | 26,00688726 | 9,04492E-07 |
| ABHD12 | 21,30452067 | 21,27832694 | 21,17292066 | 26,47924974 | 27,07060824 | 26,10133326 | 4,3446E-05 |
| PPAP2B | NA | NA | NA | 25,36623015 | 25,66152913 | 25,49710482 | 9,04492E-07 |
| NDUFA2 | 22,33123187 | NA | NA | 26,74675971 | 26,95836863 | 26,86573119 | 9,65323E-06 |
| HMOX2 | NA | NA | NA | 25,2435802 | 25,59246297 | 25,33425664 | 5,3174E-06 |
| PKD2 | NA | NA | NA | 25,82902684 | 25,86926606 | 25,80727492 | 2,54086E-05 |
| ARMC10 | NA | 22,25465036 | 22,66641907 | 26,74432348 | 27,66709075 | 26,90919195 | 3,46655E-05 |
| DNAJB14 | 21,51335364 | 20,74984652 | 20,62140445 | 26,10615475 | 26,03848216 | 26,35136025 | 7,70088E-06 |
| STX17 | NA | 22,40509575 | NA | 25,40327629 | 27,11366492 | 26,79066846 | 9,49271E-05 |
| SLC38A10 | 20,39560944 | 21,37581156 | NA | 25,90524872 | 26,08925908 | 25,93400107 | 6,86269E-05 |
| ATP2A2 | 24,92926744 | 24,89698348 | 25,04097092 | 30,03983164 | 30,10674383 | 30,17080907 | 1,2672E-05 |
| UBAC2 | NA | NA | NA | 26,32565393 | 26,52731091 | 26,29972577 | 8,99292E-06 |
| VTI1B | 23,50978273 | 23,62131501 | NA | 28,07362456 | 27,99242659 | 27,81994653 | 6,75479E-05 |
| ST6GALNAC3 | NA | NA | NA | 25,35736353 | 25,50342737 | 25,51968352 | 2,03895E-05 |
| TRAM2 | NA | NA | NA | 26,18447995 | 26,28022591 | 26,30556471 | 0,000154763 |
| TAP1 | NA | NA | NA | 25,62692216 | 26,30274406 | 26,07097667 | 5,3174E-06 |
| AFG3L2 | 22,06746456 | 22,55257717 | 22,48951775 | 27,0456232 | 27,4996767 | 27,59341791 | 6,96968E-05 |
| STX8 | NA | NA | NA | 26,24586631 | 25,87712946 | 26,00896113 | 3,58644E-05 |
| PREB | 21,8172991 | 21,85144253 | 20,37761301 | 26,16019489 | 26,32434964 | 26,43141446 | 2,59233E-05 |
| NDUFV2 | 23,07561494 | 23,82201526 | 23,72598444 | 28,18777726 | 28,93072299 | 28,31189975 | 3,99891E-05 |
| NDUFB5 | NA | NA | NA | 26,17396148 | 26,65635525 | 26,38520138 | 0,000146997 |
| FAM49A | 22,20260573 | 21,83834374 | 22,92134339 | 27,04531074 | 27,63013474 | 27,01722691 | 1,2672E-05 |
| APMAP | NA | NA | 23,10296757 | 26,55916082 | 26,57542476 | 26,5167715 | 0,000198146 |
| UFL1 | 23,37345038 | 22,2600604 | 21,92379317 | 27,49906838 | 27,37558954 | 27,35581886 | 1,29104E-05 |
| STOML2 | 24,32525933 | 23,20157416 | NA | 27,4182025 | 27,96162871 | 27,59981537 | 0,000913355 |
| TREX1 | NA | NA | NA | 25,45892046 | 25,76524299 | 25,39950012 | 2,81005E-05 |
| ARFGEF2 | 20,56866869 | 22,56153129 | NA | 26,2566851 | 26,02831534 | 26,2881878 | 6,95264E-05 |
| LMAN2 | 22,59727968 | NA | 22,46968454 | 26,40374762 | 26,76837638 | 26,68765782 | 0,00014362 |
| TMEM165 | 23,01680625 | NA | 22,89648467 | 26,9291997 | 27,15798076 | 27,04072015 | 0,000116936 |
| JAGN1 | NA | NA | NA | 25,75228366 | 25,87527896 | 25,70706692 | 9,49271E-05 |
| SLC39A7 | 24,66286044 | 24,32003367 | 24,49356671 | 29,23317908 | 29,20445491 | 29,3143871 | 1,2672E-05 |
| GOLM1 | NA | 22,71707727 | NA | 25,74604209 | 25,99170244 | 25,54005258 | 0,000308298 |
| SSR4 | NA | NA | NA | 25,94228128 | 26,23686426 | 25,87366076 | 0,00017757 |
| NDUFS5 | 22,77591405 | 22,2299043 | 23,08268052 | 27,24517723 | 27,65812735 | 27,28956035 | 9,65323E-06 |
| NDUFS4 | 22,45480431 | 22,01174867 | 22,26318781 | 27,19102939 | 27,47490978 | 26,12648298 | 4,3446E-05 |
| GJA1 | NA | 24,24662754 | 24,12597086 | 27,45057572 | 27,73582868 | 27,5408212 | 0,00821971 |
| ABCC1 | NA | NA | NA | 25,41693099 | 25,67960036 | 25,33977091 | 6,8711E-05 |
| SPARC | NA | NA | NA | 24,40256088 | 25,37348357 | 25,40856583 | 0,000138678 |
| DERL2 | 25,45331033 | NA | NA | 27,27435441 | 27,43316694 | 26,86041261 | 0,005941123 |
| DDRGK1 | 23,23947791 | NA | NA | 26,40542041 | 26,62575231 | 26,45406379 | 0,000248739 |
| SLC1A1 | NA | NA | NA | 25,44401671 | 25,52402561 | 25,25017464 | 2,59233E-05 |
| RHBDF1 | NA | NA | NA | 25,07867171 | 25,25111425 | 25,11127865 | 7,70088E-06 |
| ATP9A | NA | NA | 22,25965822 | 26,0984045 | 26,70134031 | 26,10265527 | 0,00012759 |
| BSG | NA | NA | NA | 25,36906266 | 25,26738765 | 25,12892289 | 0,000163874 |
| NDUFV1 | 25,12289433 | 25,24045363 | 25,25194493 | 29,67226007 | 29,94572856 | 29,75303963 | 3,15854E-05 |
| MARC2 | 22,38874257 | 22,63862947 | 22,47427236 | 26,93393355 | 27,42486364 | 26,83593224 | 8,99292E-06 |
| TAPT1 | NA | NA | NA | 25,02542449 | 25,29309131 | 25,11016371 | 7,70088E-06 |
| NAPG | NA | NA | NA | 24,71523475 | 24,97136498 | 24,88371573 | 3,39091E-05 |
| SLC1A5 | NA | NA | NA | 25,57873915 | 25,76065701 | 25,6227677 | 3,56703E-05 |
| CAV1 | 27,13349643 | 26,52127214 | 26,75926059 | 31,39043653 | 31,2456986 | 31,42492369 | 5,3174E-06 |
| TMED2 | NA | NA | NA | 25,30945581 | 25,58694902 | 25,85016947 | 3,50073E-05 |
| DPY19L1 | 23,15403312 | 23,17056937 | 23,72889289 | 27,29333693 | 28,53997126 | 27,82705297 | 6,86269E-05 |
| STX12 | 21,34808407 | 21,14836486 | 21,45843866 | 25,57715496 | 26,11836622 | 25,8355227 | 1,54532E-05 |
| F2R | NA | NA | NA | 25,01307864 | 25,28513888 | 25,10825038 | 1,94901E-05 |
| ZW10 | 23,93606996 | 24,04818285 | 23,98102509 | 28,70777128 | 28,24386162 | 28,47545106 | 7,71869E-05 |
| ATP11C | NA | NA | NA | 25,42828656 | 25,42764738 | 25,26792338 | 3,88583E-05 |
| TMEM192 | NA | NA | NA | 24,49466567 | 24,69991171 | 24,68421013 | 6,09575E-05 |
| CHCHD6 | NA | NA | NA | 25,37099224 | 25,61616292 | 25,60724338 | 7,70088E-06 |
| BCS1L | 22,3105795 | 22,80516525 | 22,18434752 | 26,03328167 | 27,3659466 | 27,15817343 | 2,98081E-05 |
| RHBDF2 | NA | NA | NA | 25,77690807 | 26,1491289 | 25,90334202 | 8,83709E-06 |
| WFS1 | 23,35890656 | 23,21129243 | NA | 27,1580771 | 27,28542113 | 27,26579715 | 2,80912E-05 |
| TM9SF2 | NA | NA | NA | 25,38475688 | 25,61974893 | 25,58792192 | 0,000247691 |
| TIMMDC1 | NA | NA | NA | 24,98294067 | 25,82905104 | 25,09579169 | 2,59233E-05 |
| CLN6 | 21,56555768 | 22,05198687 | NA | 25,92304506 | 26,0818439 | 25,90239922 | 1,2672E-05 |
| CANT1 | NA | NA | NA | 25,15657351 | 25,01380279 | 25,13018059 | 0,000287566 |
| SLC30A7 | NA | NA | NA | 25,17816667 | 25,2465913 | 25,33203819 | 2,98081E-05 |
| SEPN1 | NA | NA | NA | 24,92429169 | 25,29186261 | 25,00735788 | 6,44802E-06 |
| MLH1 | 23,07331197 | NA | NA | 25,69285384 | 25,68273768 | 25,7675179 | 0,002335535 |
| CHP1 | 20,91473549 | NA | 21,60891895 | 25,35712858 | 25,90499617 | 25,79634901 | 3,52786E-05 |
| TMEM161A | NA | NA | NA | 24,68613544 | 24,69407685 | 24,63616392 | 1,47761E-05 |
| ATP2C1 | 23,45274497 | 23,19859491 | 22,33821408 | 26,98174373 | 27,52902525 | 27,33577347 | 3,88583E-05 |
| GGCX | 22,48792441 | 22,26562191 | 22,05321398 | 26,74765623 | 26,10567533 | 26,80045284 | 5,18896E-05 |
| PIGS | NA | NA | 22,00020499 | 25,29077345 | 25,45676048 | 25,56340042 | 6,13362E-05 |
| GALNT7 | NA | NA | NA | 24,98563559 | 25,15553164 | 25,55338602 | 0,000173246 |

| | | | | | | | |
|---|---|---|---|---|---|---|---|
| UNC50 | NA | NA | NA | 24,8406696 | 25,08151887 | 25,56917882 | 0,000235541 |
| PTDSS2 | NA | NA | NA | 25,25659513 | 25,34499341 | 25,27579345 | 6,43093E-05 |
| TTYH3 | NA | NA | NA | 24,53296796 | 24,58583224 | 24,79084244 | 5,32202E-05 |
| CCNYL1 | NA | NA | NA | 25,04117987 | 25,4427203 | 25,20108802 | 2,35359E-05 |
| GPR180 | NA | NA | NA | 25,02850505 | 25,51893356 | 24,90559303 | 0,000141298 |
| TCIRG1 | NA | NA | NA | 25,14697537 | 25,08933991 | 25,41670554 | 3,50073E-05 |
| AGPAT9 | NA | NA | NA | 25,00045431 | 25,44780439 | 25,2399076 | 1,53802E-05 |
| SPCS1 | NA | NA | NA | 25,25688302 | 25,0822907 | 24,69975289 | 7,473E-05 |
| ABCD1 | NA | NA | NA | 25,30598212 | 25,49771396 | 25,29505507 | 5,3174E-06 |
| ANO10 | 23,47963522 | 23,52157169 | 24,02757735 | 27,58835093 | 28,12908016 | 27,71587624 | 7,70088E-06 |
| BCAP29 | 22,6478791 | 22,33388145 | 22,80595243 | 26,58348131 | 27,36711383 | 26,20810246 | 2,98081E-05 |
| TMEM55B | NA | NA | NA | 24,97066308 | 25,05759154 | 24,96018192 | 0,000458611 |
| TUBA4A | NA | NA | NA | 24,50051271 | 24,99312878 | 24,27440773 | 0,000308298 |
| ABCA3 | NA | NA | NA | 24,70835689 | 24,88049642 | 24,53201675 | 4,61931E-05 |
| IGF2R | NA | NA | NA | 24,52235021 | 24,81494501 | 24,47326925 | 0,000226096 |
| ROBO4 | NA | NA | NA | 24,87004257 | 24,80774168 | 24,37504261 | 4,48908E-05 |
| SCD5 | 23,59252 | 23,29277546 | 23,68485219 | 27,47189038 | 27,70734344 | 27,62100752 | 6,32615E-05 |
| LEMD2 | 24,56964241 | 24,4612646 | 24,04243289 | 28,3328321 | 28,65693483 | 28,23917924 | 2,70002E-05 |
| TRAM1 | 24,15259494 | 23,70524848 | 23,9826796 | 27,79197275 | 28,22080363 | 27,98239673 | 9,30193E-06 |
| NDUFS1 | 26,95537836 | 26,07379819 | 26,71122227 | 30,466066 | 30,86127082 | 30,55416953 | 0,006449597 |
| ELOVL5 | 23,73034492 | NA | NA | 25,50542779 | 26,25614518 | 25,6763486 | 0,002042266 |
| TMEM87B | NA | NA | NA | 24,75877788 | 25,11227341 | 24,767139 | 3,41404E-05 |
| MTX1 | 22,43905878 | NA | 23,18099874 | 26,22014861 | 25,91428841 | 25,7040877 | 0,001244259 |
| B3GAT3 | NA | NA | NA | 24,74627279 | 24,99295596 | 24,89666019 | 0,000531359 |
| CLPTM1 | 21,67158944 | 22,18752243 | 22,13561536 | 26,06381637 | 25,80850292 | 26,09250892 | 6,71513E-06 |
| CDKAL1 | NA | NA | NA | 24,45450313 | 24,63306218 | 24,65020419 | 9,48922E-06 |
| SOAT1 | 23,4290532 | 23,35863832 | 23,55672214 | 27,45630616 | 27,39050828 | 27,39034427 | 5,00488E-05 |
| AGPAT6 | 23,32498477 | NA | 23,0054402 | 26,09488615 | 26,61329892 | 26,59063684 | 0,000936831 |
| LEMD3 | 24,3461777 | 25,26713757 | 24,85856424 | 28,70186906 | 28,78078599 | 28,83195239 | 9,85415E-06 |
| RNF126 | NA | NA | NA | 24,94181132 | 25,27095541 | 25,0534543 | 9,49271E-05 |
| EXT2 | 22,44336549 | NA | NA | 26,11213418 | 26,25751258 | 25,05693038 | 0,000467954 |
| GPR89B;GPR89A | NA | NA | NA | 24,65730973 | 25,21737764 | 25,02850505 | 0,000190725 |
| FLNA | NA | NA | NA | 24,98702455 | 24,76339515 | 24,75241127 | 7,48015E-05 |
| LBR | 27,36218838 | 27,20815829 | 26,82432387 | 30,85675209 | 31,06599489 | 31,14643406 | 0,006990481 |
| PHB | 27,22692426 | 26,93055382 | 27,05185829 | 30,84692514 | 31,3009478 | 30,73141679 | 9,12318E-06 |
| GPR126 | 22,1318218 | NA | 22,31617103 | 25,85640482 | 24,49509282 | 26,40608573 | 0,000623321 |
| GALNT1 | 22,50554894 | 23,00111608 | 22,75293184 | 26,38348849 | 26,88145363 | 26,63574889 | 8,06141E-06 |
| PIGK | 24,18820186 | 23,88016942 | NA | 27,35178146 | 26,94022128 | 27,2141937 | 0,000350077 |
| TMEM120A;TMPIT | NA | 23,53183832 | NA | 25,29967338 | 26,13502425 | 25,97978295 | 0,001228015 |
| ATP5J | NA | 23,17834146 | 23,59616518 | 25,63430921 | 26,35228675 | 26,20114412 | 0,004816722 |
| PNKD | NA | NA | NA | 24,53771465 | 24,9361149 | 24,71219363 | 1,55072E-05 |
| LNP | NA | NA | NA | 24,86759397 | 25,21437901 | 24,77881388 | 0,000136249 |
| SLC30A5 | NA | NA | NA | 24,95947457 | 24,76526829 | 24,8488571 | 7,54399E-05 |
| STX16-NPEPL1;STX16 | 23,55145105 | NA | NA | 25,14324275 | 26,18616261 | 25,40606951 | 0,004686347 |
| ACVR1 | NA | NA | NA | 23,93598007 | 23,97281155 | 24,08099054 | 0,000112945 |
| NDUFA12 | 24,26040503 | 24,21859753 | 24,35507951 | 28,07673651 | 28,11381393 | 28,08542456 | 2,59233E-05 |
| NDUFA7 | 24,30806734 | 24,21467545 | 24,40807987 | 28,01243938 | 28,29 | 28,04786054 | 2,21687E-05 |
| SCFD1 | 23,03303376 | NA | 23,12784512 | 27,21576804 | 25,40840386 | 25,65447171 | 0,002492848 |
| VPS45 | 22,91380467 | 23,0849686 | 22,99014483 | 26,83086506 | 26,73233907 | 26,73169192 | 1,23737E-05 |
| SCAMP4 | NA | 21,83487684 | 22,16204829 | 25,30012741 | 25,85459887 | 25,49243634 | 3,46655E-05 |
| SLC27A3 | NA | 22,46495749 | NA | 25,46488273 | 25,57323019 | 25,41218915 | 7,8555E-05 |
| VAMP4 | NA | NA | NA | 24,25997423 | 24,62962032 | 24,3357905 | 5,62903E-05 |
| DOCK1 | 23,14349968 | 23,15644235 | NA | 25,34112922 | 25,94775285 | 26,10389606 | 0,003598848 |
| ATAD1 | 23,60184241 | NA | 22,99078539 | 25,84668309 | 26,68752434 | 25,41995504 | 0,003613003 |
| BCAP31 | 26,01297211 | 25,25853728 | 26,15096945 | 29,41554553 | 29,62488128 | 29,54491604 | 7,48015E-05 |
| USP30 | NA | NA | NA | 24,23680954 | 24,62811887 | 24,51074872 | 2,59233E-05 |
| ATRIP | NA | NA | NA | 24,70197479 | 24,23724731 | 24,37052672 | 6,75479E-05 |
| CERS2 | 23,67260403 | 24,0548209 | NA | 25,94900059 | 27,36427749 | 26,48585043 | 0,003046206 |
| NOTCH1 | 22,96352831 | NA | NA | 24,61924518 | 25,17157753 | 25,11382386 | 0,00254478 |
| UBE2G2 | NA | NA | NA | 24,74493934 | 24,84311635 | 25,15730622 | 0,000239397 |
| ACVRL1 | 23,20749936 | NA | 23,09611352 | 26,10495589 | 26,19044549 | 25,99168082 | 0,000507025 |
| MAOA | 22,46948582 | 23,00521745 | 22,06998988 | 25,65146387 | 27,17569451 | 25,78835512 | 0,000123897 |
| VKORC1 | 25,31333645 | NA | NA | 25,40308122 | 26,1147573 | 25,4388558 | 0,02745435 |
| SNAP23 | NA | NA | NA | 24,6348908 | 25,08602216 | 25,04701804 | 4,89712E-05 |
| GPRC5B | NA | NA | NA | 24,81680125 | 25,31575656 | 25,38426283 | 2,21687E-05 |
| TMEM173 | 24,33585863 | 24,12675866 | 24,19768045 | 27,60463025 | 28,04749656 | 28,0126525 | 3,46655E-05 |
| ASPH | 26,61470355 | 26,04497737 | 26,10359597 | 29,95129828 | 30,00423714 | 29,78026974 | 0,000290741 |
| LMF2 | 24,26656581 | 24,0997292 | 24,5318978 | 27,77089835 | 28,18333499 | 27,91193667 | 6,47356E-05 |
| COMTD1 | NA | NA | NA | 23,95881111 | 24,31699961 | 24,10813071 | 7,48015E-05 |
| TOR4A | 22,64976578 | NA | 22,71311713 | 25,36111759 | 26,07093574 | 25,15356162 | 0,001139638 |
| BAD | NA | NA | NA | 24,20970204 | 24,95269291 | 24,4790184 | 3,0743E-05 |
| MFF | 22,28571038 | NA | NA | 25,80269759 | 24,63721478 | 24,52748989 | 0,000622345 |
| PIGG | NA | NA | NA | 25,31281733 | 25,67551432 | 25,54857312 | 0,001427927 |
| TMED9 | 24,44063096 | 24,45550683 | 24,68158544 | 28,00158215 | 28,3271027 | 28,09677707 | 0,001708648 |
| FAM134C | NA | NA | 21,89385846 | 24,62689431 | 25,16146407 | 23,81616649 | 0,001378838 |
| CCDC127 | 23,11592822 | NA | NA | 24,89841429 | 26,43315102 | 25,30577343 | 0,000941659 |
| ABCC4 | NA | NA | NA | 24,54763214 | 24,67033641 | 24,56022553 | 0,000396224 |
| DNAJC30 | NA | 24,39728189 | NA | 26,35338095 | 25,81034295 | 25,92107093 | 0,003198705 |
| PON2 | 21,85014562 | NA | NA | 24,58669138 | 25,17721634 | 24,84775856 | 0,000112945 |
| NDUFS6 | 24,71816493 | 25,00774283 | 24,82036052 | 28,19192361 | 28,70658618 | 28,42342167 | 0,000142246 |
| PIEZO1 | NA | NA | NA | 24,7943175 | 24,37345038 | 24,92134339 | 8,34567E-05 |
| YME1L1 | 23,08898423 | 22,24350757 | 23,29417872 | 26,13241814 | 26,7693857 | 26,44156521 | 5,622E-05 |
| ATP5D | NA | NA | NA | 24,36105063 | 25,17972386 | 24,31181316 | 2,59233E-05 |
| PTPRF | NA | NA | NA | 24,2155644 | 24,06595382 | 24,22515977 | 0,000136866 |
| DAB2IP | 23,50675986 | 22,66687411 | 22,81005851 | 26,45740255 | 26,24119953 | 26,95004697 | 6,95264E-05 |
| ZDHHC13 | 22,33521805 | NA | 22,48608375 | 24,97801597 | 26,1239403 | 25,41125184 | 0,000214686 |
| LETM1 | 25,1945654 | 24,89684494 | 25,27629056 | 28,5164935 | 28,7933999 | 28,66715844 | 7,87842E-05 |
| COG1 | 24,11545203 | 23,73634495 | 23,05382714 | 27,23954347 | 27,1890507 | 27,08415764 | 5,32202E-05 |
| GOLIM4 | 21,62377256 | NA | 20,7019325 | 24,72821823 | 25,0252555 | 24,71460608 | 5,04595E-05 |
| GNA11 | NA | NA | 23,94073656 | 25,49704389 | 25,95155953 | 25,24630134 | 0,003297664 |
| SLC16A3 | 23,34894855 | 23,44758372 | 23,7060921 | 27,24417929 | 27,40641017 | 26,40863061 | 0,000652051 |
| TMEM35 | 22,81850885 | NA | NA | 24,73283071 | 25,26001013 | 24,7456575 | 0,002641913 |
| ADAM9 | 22,34063359 | 22,20746957 | 21,98561822 | 24,82143145 | 26,64223727 | 25,60127569 | 0,000345619 |
| FAF2 | 25,01252463 | 25,27643257 | 25,35860479 | 28,47950417 | 28,989073 | 28,6412383 | 8,99292E-06 |
| GIMAP2 | 21,80691615 | NA | NA | 24,05133973 | 24,18381769 | 24,30089545 | 0,001139823 |
| MPZL1 | NA | NA | NA | 24,00152846 | 24,09393976 | 24,14600427 | 0,000502241 |
| COG5 | 23,76283781 | NA | 22,52921892 | 26,11721711 | 25,91624905 | 25,42191406 | 0,002045225 |
| DNAJC11 | 24,16545663 | 24,22736505 | 24,16392234 | 27,17178748 | 27,87507971 | 27,90604052 | 1,90859E-05 |
| DHRS7 | 24,15120178 | NA | 23,8196786 | 26,98609149 | 26,78031668 | 27,04218237 | 0,000364099 |
| MGARP | NA | NA | NA | 25,24561247 | 24,48376117 | 24,69842871 | 0,000156632 |
| MT-CO2 | NA | NA | NA | 24,02225266 | 24,84570252 | 24,52175138 | 3,86828E-05 |
| NOP9 | NA | NA | 22,84132248 | 24,29094917 | 24,03816815 | 24,44278357 | 0,003297664 |
| TGOLN2 | NA | NA | 21,73888227 | 24,60365441 | 24,74990788 | 24,57646313 | 0,000136249 |

| | | | | | | | |
|---|---|---|---|---|---|---|---|
| FADS2 | 24,07242901 | NA | 24,28869826 | 26,42452731 | 26,54002301 | 26,28280819 | 0,008364382 |
| YIF1B | 23,27341203 | 23,47122371 | 24,06603596 | 26,95847926 | 27,21038969 | 26,95958511 | 0,001835084 |
| KIAA0100 | 21,46906344 | 21,85502204 | 23,06844898 | 25,66372921 | 25,3431304 | 25,69317299 | 0,000173754 |
| TOMM22 | 23,09342401 | NA | NA | 25,3138208 | 25,78139273 | 25,56880204 | 0,000287566 |
| CASC4 | NA | NA | NA | 24,21615673 | 23,92533349 | 24,10461603 | 2,21687E-05 |
| RCN1 | NA | NA | NA | 24,7234868 | 24,47673385 | 24,83672689 | 0,000236932 |
| FKBP8 | 23,60580322 | 23,45211654 | 23,59866592 | 26,83906016 | 27,17835666 | 26,917069 | 0,000214533 |
| GJA5 | 22,64956845 | 22,28599252 | 22,56950334 | 26,01150129 | 25,91298744 | 25,85590603 | 7,44581E-05 |
| CD93 | 22,90398552 | 23,52695288 | 21,85593454 | 25,94877786 | 26,2824017 | 26,31333645 | 0,000214686 |
| PTK7 | NA | NA | NA | 24,70392934 | 25,14125587 | 24,96736841 | 9,65323E-06 |
| HSDL1 | 22,62520049 | 22,33750649 | NA | 24,59246297 | 25,89647543 | 24,94337726 | 0,001548194 |
| EIF2B3 | 22,13755531 | 21,89211735 | 21,76971357 | 25,00200083 | 25,37878407 | 25,59263405 | 2,59233E-05 |
| PTTG1IP | NA | 21,97938612 | 21,79673517 | 25,27156105 | 25,1215117 | 25,22177177 | 0,000732731 |
| COX20 | 25,26474181 | NA | 25,07956714 | 27,47297499 | 28,00512106 | 27,44118524 | 0,003397624 |
| TUBB2A;TUBB2B | 25,74170315 | NA | NA | 26,22315371 | 26,72559447 | 26,240581 | 0,01651765 |
| RNF121 | NA | NA | NA | 24,38119597 | 25,02660685 | 25,00127074 | 9,35679E-05 |
| HLA-E | 21,30067207 | 22,0573519 | NA | 24,66990407 | 24,91145672 | 24,64691286 | 0,000107597 |
| PTPRM | NA | NA | NA | 23,33087641 | 23,40353636 | 23,61548955 | 0,001769795 |
| TMPPE | NA | NA | NA | 23,5234873 | 24,23921567 | 23,52037314 | 9,56871E-05 |
| EFNB1 | NA | NA | NA | 24,33313078 | 24,62811887 | 24,62639306 | 0,000451805 |
| SEC61G | NA | NA | NA | 23,84708947 | 23,82269608 | 24,97197885 | 0,002509307 |
| STX4 | 21,86876228 | 21,61463616 | 21,79986024 | 24,57968883 | 25,45851375 | 25,26513542 | 0,000169963 |
| CDK5RAP3 | NA | NA | NA | 24,21955792 | 24,23680954 | 24,01256726 | 0,000299615 |
| TMED10 | 25,89864493 | 25,89087506 | 25,93129804 | 29,24372546 | 29,43199222 | 29,02207801 | 0,000177853 |
| SLC27A4 | 24,65545473 | 24,22560109 | 24,14328169 | 27,57304235 | 27,86390145 | 27,54833794 | 2,88271E-05 |
| PTDSS1 | 23,59593763 | 24,14125587 | NA | 26,57945866 | 26,30007503 | 26,36806359 | 0,002194698 |
| ASPHD2 | NA | NA | NA | 24,59576694 | 24,2278057 | 24,30096525 | 0,00083458 |
| DNAJB12 | 23,98979847 | 24,36666372 | 24,5542647 | 27,58770737 | 27,6007371 | 27,54421595 | 3,46655E-05 |
| WLS | NA | 23,84143766 | 24,0129082 | 25,51953356 | 27,32018865 | 25,82992205 | 0,008383831 |
| TMEM115 | 22,16063346 | NA | 22,61488325 | 24,74468276 | 25,72061961 | 25,21637879 | 0,000247802 |
| VTI1A | NA | NA | NA | 24,52826523 | 24,49216126 | 24,69040472 | 8,39203E-05 |
| ATP6V1A | NA | NA | NA | 23,16761721 | 23,21089128 | 23,48296154 | 0,001110994 |
| TMED4 | NA | NA | NA | 24,48818191 | 24,96723646 | 24,49344455 | 0,001183864 |
| DOCK1 | NA | NA | NA | 24,5048219 | 24,49996556 | 24,53854371 | 0,000346197 |
| SCO1 | NA | NA | NA | 23,46338667 | 24,17066105 | 23,51484694 | 0,003628764 |
| AGPAT3 | 22,56162451 | NA | NA | 24,92406511 | 25,21874532 | 25,18495281 | 0,00117407 |
| TMCO1 | 24,11028321 | 23,4287977 | 22,41668621 | 26,60272038 | 26,6699311 | 26,33963501 | 0,000176286 |
| RTN4 | NA | NA | 24,21837581 | 25,3859748 | 26,17237898 | 25,87178231 | 0,011990461 |
| STT3B | 25,36649698 | 25,44783591 | 25,41128417 | 28,62278865 | 28,75983202 | 28,48734639 | 1,23737E-05 |
| ALG1 | 23,93183905 | 23,85993889 | 24,26248545 | 27,05558646 | 27,46435305 | 27,16777415 | 0,000414255 |
| UBE3B | NA | NA | NA | 24,07952645 | 24,19640522 | 24,05944943 | 0,002140357 |
| CHST14 | NA | 23,51605009 | NA | 24,71659593 | 25,15325235 | 25,07451291 | 0,002520668 |
| NSF | 25,40314625 | 25,45422071 | 25,02795694 | 28,40104759 | 28,5653267 | 28,53760358 | 0,00025333 |
| RABL3 | NA | 24,13758657 | 24,11275065 | 26,42276429 | 25,79988493 | 26,76331916 | 0,007123838 |
| PTRH2 | 24,89905999 | 24,12904085 | 25,00341703 | 27,39280243 | 28,2115041 | 28,02559346 | 0,000167832 |
| ARMCX2 | 23,21043057 | NA | 23,59593763 | 26,2859502 | 25,31444329 | 25,00341703 | 0,014956473 |
| TBC1D10B | 23,13254366 | NA | 23,25200992 | 26,30328424 | 26,3900326 | 25,03006393 | 0,009941659 |
| RPN2 | 26,51320604 | 26,20133111 | 26,6056902 | 29,45122448 | 29,83856436 | 29,54423437 | 0,00025017 |
| SRPRB | 25,98906217 | 25,44975747 | 26,09902686 | 28,61150604 | 29,31921975 | 29,11473249 | 0,001503284 |
| SLC27A1 | 23,8438352 | 23,72910041 | 23,71061812 | 26,76850258 | 27,24163598 | 26,77580356 | 0,000123897 |
| FADS3 | 24,2740522 | 24,01103202 | 23,21261394 | 26,32522502 | 27,33951609 | 27,31518647 | 0,00044422 |
| CAPN5 | NA | NA | NA | 24,03380676 | 24,46413489 | 24,21970562 | 0,000324601 |
| CERS5;LASS5 | NA | NA | NA | 23,66893085 | 24,06085159 | 23,44834018 | 0,00042248 |
| MMP14 | 25,94637017 | 25,81159283 | 26,22387179 | 28,90227845 | 29,25739119 | 29,20464147 | 0,000237713 |
| NDUFB4 | 25,43323063 | 25,62335396 | 25,4378087 | 28,43591088 | 28,92753353 | 28,50184174 | 3,2107E-05 |
| GRAMD4 | NA | NA | NA | 23,83489612 | 24,21822797 | 23,93111766 | 0,000115479 |
| FLRT2 | 23,85889617 | 23,55227227 | 23,67271192 | 26,71502522 | 26,82723473 | 26,90506505 | 0,000142352 |
| HAX1 | NA | NA | NA | 24,08586013 | 24,7575068 | 24,17081384 | 9,02393E-05 |
| TMEM126B | NA | NA | NA | 23,54545378 | 24,02021902 | 23,66752392 | 3,2107E-05 |
| NDUFS8 | 23,77996617 | 23,98311469 | 23,07443941 | 26,85078946 | 26,5869204 | 26,70477373 | 0,000198146 |
| LCLAT1 | 23,30813679 | 23,18322708 | 23,60636816 | 26,37273662 | 26,69325276 | 26,32978211 | 0,000112945 |
| CAMLG | 22,91850733 | NA | NA | 25,03019025 | 24,79357354 | 25,2339973 | 0,00071295 |
| MAVS | 23,56010889 | 23,20879459 | 23,73376179 | 26,31920683 | 26,84756742 | 26,62296315 | 5,61599E-05 |
| ATM | 24,70672449 | 24,50573064 | 24,55853329 | 27,63588725 | 27,61526502 | 27,80513572 | 8,63978E-05 |
| DOCK5 | NA | NA | NA | 23,96965352 | 23,68495917 | 23,92397447 | 0,000141481 |
| XXYLT1 | 23,63754647 | NA | 23,58141392 | 25,65780035 | 25,77160371 | 25,76895682 | 0,006428887 |
| TOMM70A | 24,31554928 | 23,90591432 | 24,51761269 | 27,01966775 | 27,56985966 | 27,3740145 | 3,86828E-05 |
| FLNA | NA | 23,00545734 | NA | 24,79535838 | 24,93490095 | 24,68997836 | 0,003668647 |
| CERS6 | NA | NA | NA | 23,42201034 | 23,7293079 | 23,58772167 | 0,000732731 |
| EPHA2 | 24,75501224 | 24,80798729 | 25,36816353 | 27,86879524 | 28,41836338 | 27,84720897 | 0,000123897 |
| CLCN3 | NA | NA | NA | 24,71502522 | 24,89573613 | 24,49984394 | 0,00037833 |
| BRI3 | 22,6909588 | NA | NA | 24,39728189 | 24,85071793 | 24,61363616 | 0,000818879 |
| FKBP11 | 24,73572541 | 24,15228546 | 24,41796116 | 27,37276983 | 27,82268392 | 27,28674345 | 3,46655E-05 |
| PHB2 | 26,85970198 | 26,87829943 | 27,06095463 | 29,83202786 | 30,27253133 | 29,85816108 | 3,50073E-05 |
| ABCB10 | 22,41933787 | 22,42388325 | NA | 24,29151135 | 25,43415377 | 25,68359456 | 0,001037392 |
| MTX2 | NA | NA | 23,90921485 | 24,60969683 | 25,28651433 | 24,84062159 | 0,024411824 |
| SSR1 | 22,54035416 | 22,8498403 | 22,98025814 | 25,99520093 | 26,05113225 | 25,43990214 | 0,000308298 |
| ERLIN1 | NA | NA | NA | 24,84076563 | 24,91013032 | 24,91825254 | 0,022574487 |
| SCAMP3 | 23,17250869 | 23,52097254 | 23,59616518 | 26,55296114 | 26,53431939 | 26,30667753 | 0,0002364 |
| IQGAP3 | 21,41281584 | 21,82694875 | NA | 25,25439807 | 24,47704278 | 23,53860292 | 0,002657859 |
| TMX3 | NA | NA | NA | 23,59968769 | 23,72961908 | 23,75572541 | 0,002256636 |
| TECR | 25,46326193 | 25,45556954 | 25,70292598 | 28,48703965 | 28,7252369 | 28,50028476 | 2,11966E-05 |
| TRADD | 23,46201392 | NA | 23,82648332 | 25,72502232 | 25,261338 | 25,39047548 | 0,023222926 |
| EVI5L | NA | NA | 23,97867066 | 25,07021924 | 25,03657609 | 25,59086524 | 0,031914555 |
| JAG1 | NA | NA | NA | 24,79852599 | 24,82972854 | 24,57945866 | 3,18309E-05 |
| AMFR | NA | NA | NA | 24,08391426 | 24,16092577 | 24,0709562 | 0,000328472 |
| CXCR4 | NA | NA | NA | 23,78954956 | 24,10980515 | 23,25059393 | 0,000777061 |
| ALG2 | 23,52575879 | NA | 23,02408048 | 25,63353341 | 26,17839466 | 24,92129798 | 0,002256636 |
| CYB5R1 | 25,46241964 | 24,62366094 | 24,780567 | 27,77303857 | 28,2341618 | 27,86260151 | 5,15001E-05 |
| TEK | 23,21408251 | 23,51929359 | 22,94595957 | 26,64416435 | 26,55672214 | 25,47035502 | 0,00190568 |
| UQCRH;UQCRHL | 25,58980862 | 24,82919624 | 25,35009587 | 27,91467619 | 28,6719565 | 28,16451707 | 0,000142352 |
| STX5 | 25,780567 | 25,97939048 | 26,07612485 | 28,76002244 | 29,00747552 | 29,04280859 | 0,000251605 |
| RAB35 | 23,93580027 | 23,66003333 | 23,1856032 | 26,43676084 | 26,69750106 | 26,58806494 | 0,000131103 |
| ALG5 | 22,57706273 | 23,26155322 | 23,6462537 | 26,18954092 | 25,75350833 | 26,46461791 | 0,000116213 |
| SSFA2 | 21,28076397 | NA | 21,94520948 | 24,20687366 | 24,29907942 | 24,15946364 | 0,000328472 |
| PIGQ | NA | NA | NA | 24,4388558 | 24,67244218 | 24,76324317 | 0,000202666 |
| FAM162A | 24,1918201 | NA | 24,03069542 | 26,43288032 | 26,2459932 | 25,5517444 | 0,009633647 |
| ABCB8 | NA | NA | NA | 23,83441395 | 24,3688629 | 24,00178614 | 0,000364053 |
| TMPO | NA | NA | NA | 23,63920379 | 23,90802386 | 23,87502111 | 5,37201E-05 |
| CCDC47 | 22,68649882 | 22,58950577 | 22,40054281 | 25,17843265 | 26,10193433 | 25,21107701 | 0,000123747 |
| NDUFA6 | 25,60585972 | 25,56917882 | 25,31219413 | 28,39050828 | 28,58992289 | 28,29994407 | 0,000433049 |
| TMEM2 | NA | NA | NA | 23,63754647 | 23,94788659 | 23,83749705 | 0,000199159 |
| GBF1 | 24,12020686 | 24,35944288 | 24,51544864 | 27,25124069 | 27,27444328 | 27,25015657 | 0,0002245 |

| | | | | | | | |
|---|---|---|---|---|---|---|---|
| GNG12 | 24,24691742 | NA | 23,98485371 | 25,53079708 | 25,9856573 | 25,65984285 | 0,031726265 |
| IMMT | NA | NA | NA | 23,74314235 | 24,07585972 | 23,66600724 | 0,000262856 |
| BCAM | NA | NA | NA | 23,74232013 | 23,81009775 | 24,07054682 | 0,002003026 |
| ESAM | NA | NA | NA | 23,52444415 | 23,60896405 | 23,57276776 | 9,02393E-05 |
| DNAJC1 | 23,98407141 | 23,50348803 | 23,78476582 | 26,68418337 | 26,63075926 | 26,72168895 | 0,000125733 |
| NDC1 | NA | NA | 24,19437754 | 25,31699961 | 25,66106696 | 25,22681404 | 0,017999846 |
| TMEM41A | NA | NA | NA | 23,52838448 | 24,10980515 | 23,61335514 | 0,000188716 |
| HMOX1 | 23,84642007 | 23,71114348 | 24,33504082 | 26,79054418 | 26,89214978 | 26,93954891 | 2,98081E-05 |
| CKAP4 | 30,30834514 | 30,42031252 | 30,22366009 | 33,20844903 | 33,20868158 | 33,24968766 | 8,63978E-05 |
| EMC3 | NA | NA | 23,46898889 | 25,28943723 | 25,433931 | 25,46372965 | 0,000778967 |
| TMEM209 | 22,5415835 | 22,49806715 | 23,0792009 | 25,14693654 | 26,21297009 | 25,45434624 | 0,000169963 |
| YIPF6 | NA | NA | NA | 24,32759102 | 24,58583224 | 24,07700148 | 0,00068567 |
| GNAI2 | 23,83151756 | 23,76850258 | 24,36372626 | 26,7494476 | 26,93977307 | 26,92660076 | 0,000926761 |
| EI24 | NA | NA | NA | 24,14685887 | 24,36125148 | 24,27234445 | 0,002689207 |
| CCDC109B | NA | NA | NA | 23,99057767 | 24,26047682 | 24,08537391 | 0,000321963 |
| COG3 | NA | 23,41454589 | NA | 25,15676636 | 24,65053291 | 25,36813021 | 0,0018531 |
| GALNT2 | 25,92503915 | 26,0650499 | 25,90120259 | 28,81537264 | 28,89951528 | 28,7444518 | 3,36549E-05 |
| AUP1 | 24,66654909 | 24,24836598 | 24,50221365 | 27,25682904 | 27,46193589 | 27,25565909 | 7,28248E-05 |
| ROMO1 | 23,29838034 | NA | NA | 24,24641008 | 24,84172557 | 24,34516265 | 0,009780605 |
| SLC2A1 | NA | NA | 22,01665325 | 24,34062001 | 23,99808835 | 24,03178936 | 0,002692804 |
| SLC30A6 | NA | NA | NA | 23,6156018 | 23,65654619 | 23,78935056 | 0,000142352 |
| PRAF2 | NA | NA | NA | 23,74919183 | 24,25068066 | 23,75796452 | 0,000157144 |
| FUT8 | NA | 21,70282032 | 22,16432141 | 24,49551984 | 24,52957639 | 24,60907681 | 0,000652051 |
| PTPRK | NA | 22,06121428 | NA | 24,32608271 | 24,71999328 | 24,23666358 | 0,002224731 |
| TMTC3 | 25,20579295 | 25,54801449 | 25,95575525 | 28,41025731 | 28,52309092 | 28,25741367 | 0,000103829 |
| ARFGEF1 | 24,60557718 | 24,38198804 | 25,03753992 | 27,45175506 | 27,54877888 | 27,47220035 | 3,18309E-05 |
| EMD | 26,31573929 | 26,15990628 | 26,29886974 | 28,94355053 | 29,19540112 | 29,0673035 | 0,005591359 |
| SLC16A1 | 26,85138534 | 26,70714593 | 26,80980345 | 29,49707055 | 29,76872341 | 29,52411904 | 0,00129615 |
| SMPD4 | 23,78476582 | 23,75154324 | 23,49728759 | 25,7533553 | 26,98272312 | 26,69218873 | 0,000178564 |
| GPR124 | 24,70872524 | 24,27625506 | 22,88859165 | 26,92942548 | 26,52994866 | 26,78793184 | 0,000649381 |
| MCAM | 24,2985901 | 23,26927966 | 23,3253966 | 26,17847065 | 26,37712976 | 26,69922337 | 0,000433049 |
| GPX8 | 26,82310927 | 26,55539155 | 26,85340951 | 29,43037789 | 29,56795755 | 29,58461067 | 6,86269E-05 |
| SRPR | 26,68070054 | 27,11406224 | 26,93663165 | 29,35741387 | 30,36807192 | 29,33112422 | 0,034483071 |
| EVI5 | 21,89877407 | NA | 22,36900939 | 24,49368886 | 23,67206444 | 24,53362156 | 0,003480559 |
| RFTN1 | NA | 22,81930828 | NA | 23,57945866 | 24,06365182 | 23,6530506 | 0,008175036 |
| SLC29A1 | NA | NA | NA | 24,274621 | 24,07585972 | 23,94583458 | 0,000694442 |
| NDUFS3 | 25,68592164 | 25,510628 | 25,77848806 | 28,28467128 | 28,50233507 | 28,47772992 | 0,000198146 |
| GIMAP1-GIMAP5 | NA | NA | NA | 23,62800759 | 23,61526502 | 23,73334805 | 0,00033803 |
| TMEM87A | NA | NA | NA | 23,6612301 | 24,16161783 | 23,71030281 | 0,00017814 |
| CHMP6 | NA | 21,97683678 | NA | 24,36992796 | 23,63333939 | 24,33864934 | 0,001318654 |
| TP53I11 | NA | 22,43824667 | 22,1285847 | 24,94140838 | 24,12801824 | 25,05920185 | 0,003833175 |
| CYP27A1 | NA | NA | NA | 23,90664842 | 24,30089545 | 23,88604408 | 0,001571837 |
| NDUFS2 | 23,94975762 | 24,37145762 | 24,99701163 | 27,07907879 | 27,22995195 | 27,19788663 | 0,000407503 |
| STX6 | NA | NA | NA | 24,12155122 | 24,10245505 | 24,09522831 | 0,000829919 |
| CTSZ | NA | NA | NA | 23,46213877 | 23,47914178 | 23,4230369 | 0,000289241 |
| USO1 | NA | NA | NA | 23,74683657 | 24,01954051 | 24,28503302 | 0,002689409 |
| TMEM11 | NA | NA | NA | 24,11330723 | 24,80395411 | 23,96895079 | 0,000913355 |
| ITGA3 | NA | NA | NA | 23,22999228 | 23,29949871 | 23,40249583 | 0,000184933 |
| SYNJ2BP-COX16;COX16 | NA | NA | NA | 22,93266821 | 23,44670068 | 23,59160726 | 0,003493435 |
| HADHB | 26,49743988 | 26,54995407 | 26,42579215 | 28,68311263 | 29,78045749 | 29,1360419 | 0,000112945 |
| RNF5 | NA | NA | NA | 23,33415433 | 24,13602234 | 23,87605224 | 0,000423088 |
| TMEM30A | NA | NA | NA | 23,5406143 | 23,42700791 | 23,54049606 | 0,002074831 |
| GLT8D1 | 22,96246961 | 23,11638838 | 23,29487984 | 25,62800759 | 26,26262881 | 25,55414757 | 9,02393E-05 |
| HADHA | 27,48008226 | 26,72974872 | 27,2127104 | 29,72535068 | 30,16350972 | 29,60052445 | 0,023613438 |
| UBE3C | 23,96560816 | 23,05963096 | 23,44215078 | 26,0889155 | 26,32771093 | 26,10777164 | 9,35414E-05 |
| TMEM245 | NA | NA | NA | 24,35937585 | 24,6087949 | 24,22398224 | 0,003627469 |
| COA3 | 23,04478564 | 22,91031334 | 22,4525439 | 25,17291303 | 25,78915152 | 25,47766045 | 0,003100361 |
| SEC22A | 23,1084817 | 23,53516524 | 23,33905729 | 25,82065267 | 26,26209112 | 25,92910938 | 0,000178304 |
| RRAS | 22,4767462 | NA | NA | 24,07103806 | 24,40021705 | 24,08529285 | 0,001540242 |
| | 21,72621317 | NA | 21,53331262 | 24,27127608 | 24,36979487 | 24,1467812 | 0,000144042 |
| GPR107 | NA | NA | NA | 24,0282521 | 24,17073745 | 24,20247122 | 0,001096343 |
| RPN1 | 28,56358225 | 28,46244305 | 28,54679357 | 31,12743287 | 31,26718223 | 31,17218104 | 5,17469E-05 |
| ERMP1 | 22,92784884 | NA | NA | 24,32154831 | 24,32086003 | 24,85533578 | 0,002137932 |
| ACTA1;ACTC1;ACTG2;A( | 26,04703885 | 25,37202251 | 26,16802292 | 28,35085463 | 28,82602258 | 28,36202112 | 0,000326129 |
| ZMPSTE24 | 22,84149525 | 23,17180656 | 23,16614653 | 25,67984194 | 25,73347736 | 25,71248229 | 0,002520668 |
| CLU | 23,32827609 | NA | 23,16822956 | 25,59880787 | 25,51857344 | 24,65638253 | 0,00426223 |
| NDUFA13 | 25,32213308 | 25,51358256 | 25,54415698 | 27,86814762 | 28,51949607 | 27,90173199 | 0,000422986 |
| TACSTD2 | NA | NA | NA | 24,50209222 | 24,2137118 | 24,0796892 | 0,000710653 |
| ANKLE2 | 24,09514781 | 24,25031924 | 23,19413703 | 26,37638469 | 26,69630749 | 26,34685401 | 0,00027115 |
| VAPA | 24,73474393 | 24,83354564 | 24,8632989 | 27,25754855 | 27,61400141 | 27,43864961 | 0,00033405 |
| RDH14 | 24,02799911 | NA | 24,05944943 | 25,62948136 | 25,26706612 | 25,34384201 | 0,033654786 |
| UBE4A | NA | NA | NA | 22,86101873 | 23,83759329 | 22,881869 | 0,000556695 |
| MBOAT7 | 27,49891627 | 27,2622166 | 27,3600125 | 30,0077161 | 30,05504337 | 29,91612944 | 4,15196E-05 |
| COG2 | 24,01503726 | 23,22864257 | 24,14172362 | 26,25582114 | 26,49556559 | 26,484714 | 0,003246836 |
| CDS2 | NA | NA | NA | 23,63167532 | 24,07716451 | 23,77011701 | 0,002193416 |
| DHRS7B | NA | 22,87702645 | NA | 24,27099104 | 24,75948919 | 24,68859182 | 0,003903027 |
| TMCO3 | NA | NA | NA | 24,00795664 | 23,59627895 | 23,59023707 | 0,00306744 |
| LRBA | NA | NA | NA | 23,28743058 | 23,05715355 | 23,55870844 | 0,000483387 |
| SOS1 | NA | NA | NA | 23,37238792 | 23,07246172 | 23,03800065 | 0,000245999 |
| CALCRL | 22,77237418 | 22,8346647 | 23,33183324 | 25,60501193 | 25,66589884 | 25,46173297 | 0,000734242 |
| TIE1 | 25,17039364 | 25,13218277 | 25,2005643 | 27,63878964 | 27,83827882 | 27,81616649 | 0,000160894 |
| OCIAD2 | 24,75174753 | 24,72530842 | 24,9060061 | 27,43101587 | 27,5724642 | 27,1669127 | 0,000202666 |
| SH3TC1 | NA | NA | NA | 23,17613601 | 23,59160726 | 23,99196188 | 0,001926119 |
| ICAM2 | 25,96684056 | 25,18635155 | 25,68029814 | 28,06486494 | 28,48143023 | 28,06886715 | 0,000123897 |
| TBC1D15 | NA | NA | NA | 23,50675986 | 23,89208028 | 23,55520136 | 0,000328472 |
| SACM1L | NA | NA | NA | 23,19973344 | 23,38152605 | 22,98653857 | 0,000214686 |
| MAN1A2 | 23,48738472 | 23,65512713 | 22,71845762 | 26,17094752 | 25,4639167 | 25,90449093 | 0,001950308 |
| FLT4 | 23,82036052 | 23,85861167 | 23,52288894 | 26,46726386 | 26,24081753 | 26,17148209 | 0,000903738 |
| SYVN1 | NA | NA | NA | 23,59809795 | 23,27198841 | 23,20466666 | 0,000599391 |
| RABGAP1 | NA | NA | 21,974458 | 23,11824343 | 23,72046305 | 23,60410706 | 0,002509307 |
| PNPLA2 | NA | NA | 24,06208765 | 24,11632492 | 24,48848841 | 24,23855985 | 0,03855599 |
| ATP5B | 26,18569015 | 25,81941061 | 26,44173932 | 28,40932684 | 28,93258261 | 28,70865947 | 7,39215E-05 |
| HK1 | 24,08529285 | 23,22642454 | 23,90646493 | 26,08737851 | 26,76141813 | 25,96569623 | 0,000422986 |
| TBL2 | 27,50650263 | 27,49358198 | 27,20787912 | 29,59573138 | 30,5388064 | 29,63633681 | 0,000507025 |
| TMEM205 | NA | NA | NA | 23,68131735 | 23,66438044 | 23,4093754 | 0,000287566 |
| ZDHHC5 | 22,16432141 | NA | NA | 24,27305626 | 24,47004465 | 23,68923192 | 0,007776862 |
| MARCH6 | NA | NA | NA | 22,53071971 | 22,65883558 | 22,8931919 | 0,000328472 |
| GPD2 | NA | NA | NA | 22,68694757 | 23,27540274 | 23,35002841 | 0,000648325 |
| CUX1 | NA | NA | NA | 23,69816373 | 23,68303229 | 23,71030281 | 0,002128441 |
| LRPPRC | NA | NA | NA | 23,00035974 | 23,86551883 | 22,9624343 | 0,000469469 |
| DNAJC25;hCG_1994888 | 23,18974072 | NA | 22,91801591 | 24,64960134 | 24,90839042 | 25,01546269 | 0,003606109 |
| SLC3A2 | NA | NA | NA | 24,05573125 | 23,83045408 | 23,6946614 | 0,000136249 |
| TPST2 | NA | 22,20338266 | NA | 24,1435931 | 23,33456354 | 23,41763931 | 0,002024124 |

| | | | | | | | |
|---|---|---|---|---|---|---|---|
| RDH11 | 24,3970207 | 24,45173935 | 24,41873331 | 26,87537271 | 26,94726238 | 26,91752432 | 0,001215017 |
| FAM20B | 22,29504806 | NA | NA | 23,74345057 | 23,93183905 | 23,67594498 | 0,001863161 |
| RALA | 25,34621153 | 26,14024189 | 25,67529895 | 27,85335001 | 28,56804819 | 28,18517972 | 0,002094389 |
| FRMD8 | NA | 22,27580055 | NA | 23,44265704 | 23,48369968 | 23,57922846 | 0,004203224 |
| TIMM21 | 22,468094 | 22,65186893 | NA | 24,9355755 | 25,00906795 | 24,6863492 | 0,001368744 |
| TFRC | NA | NA | NA | 22,9263474 | 22,87425197 | 23,08074663 | 0,001694286 |
| NDUFA9 | 25,24103583 | 25,40905163 | 25,29803067 | 27,67754535 | 28,03280892 | 27,61659764 | 0,000299615 |
| NDUFB10 | 22,61288289 | NA | 23,38007312 | 24,52037314 | 25,33316491 | 24,69524571 | 0,003301955 |
| STX7 | 23,5167715 | 23,38798049 | 23,14960573 | 25,30931703 | 25,76986488 | 26,34372344 | 0,009799821 |
| REEP4 | 22,60867084 | 21,95659736 | 22,51390779 | 24,61009125 | 24,94194561 | 24,88613714 | 0,000160894 |
| GOLGA5 | 23,14368651 | 22,84571209 | 23,79441666 | 25,74098301 | 25,98015353 | 25,4193443 | 0,000292446 |
| ITPRIPL2 | NA | NA | NA | 23,05817806 | 23,85661853 | 23,2479026 | 0,002627986 |
| C2CD2 | 22,04216567 | 21,88114085 | 22,21676362 | 24,79099154 | 24,40308122 | 24,2690656 | 0,00033803 |
| STT3A | 25,3859748 | 25,26810191 | 25,60441817 | 27,77599189 | 27,79699255 | 27,97415819 | 3,88583E-05 |
| FAM198B | 23,08856376 | 23,28008429 | 22,84674525 | 25,48664847 | 25,46472696 | 25,51815319 | 0,000198146 |
| RIC1 | 23,26026145 | NA | NA | 24,01546269 | 24,83778575 | 24,22574817 | 0,013042526 |
| SLC33A1 | NA | NA | NA | 23,31962031 | 23,60025504 | 23,15539267 | 0,011832855 |
| MARCH5 | 21,55407729 | 21,18894132 | 22,68583611 | 24,16384558 | 24,41415874 | 24,08456317 | 0,00257168 |
| GNB2 | NA | NA | 23,19340023 | 24,4630748 | 25,15502973 | 24,45381268 | 0,002568671 |
| NBEAL1 | 23,53160039 | NA | NA | 24,29831042 | 24,26313048 | 24,05647565 | 0,011228974 |
| CMIP | NA | 22,1577688 | NA | 23,42624018 | 24,11949464 | 24,1434374 | 0,00186348 |
| TMEM109 | 26,95881111 | 26,60016995 | 27,19338143 | 29,39774702 | 29,37191869 | 29,18824904 | 0,022934101 |
| SCARF2 | NA | NA | NA | 22,99834664 | 23,56546472 | 23,23317085 | 0,001178155 |
| NLRX1 | 22,7293079 | NA | NA | 23,78366731 | 24,60331484 | 24,30959459 | 0,001531387 |
| IARS2 | NA | NA | NA | 22,98071139 | 23,0380174 | 23,09168202 | 0,001599822 |
| EFNB2 | NA | NA | NA | 21,98471467 | 22,71776761 | 21,97547296 | 0,000178564 |
| RAB5C | 23,93219961 | 24,2790925 | 24,078631 | 26,6073845 | 26,50476129 | 26,34507803 | 0,000295799 |
| ERLIN2 | NA | NA | NA | 23,85014562 | 23,80360929 | 23,92813809 | 0,001096343 |
| TMEM43 | 23,04799987 | 23,1965553 | 23,18171158 | 25,50787906 | 25,5584749 | 25,5187235 | 0,002779591 |
| RAB33B | 22,19787538 | NA | NA | 23,40431626 | 24,33463174 | 23,82522201 | 0,002116125 |
| RALB | 23,08563324 | 23,88660233 | 23,13523958 | 25,74142028 | 25,81426052 | 25,68048594 | 0,000724666 |
| CD46 | NA | NA | NA | 22,94148898 | 22,67040124 | 22,67191332 | 0,001139823 |
| DNAJC19 | 25,12159074 | 23,76172246 | 24,07545173 | 26,71554898 | 26,78456616 | 26,5791709 | 0,003834596 |
| ZDHHC3 | NA | NA | NA | 23,16260154 | 23,28362083 | 23,19628514 | 0,00162019 |
| CISD1 | 23,46612825 | 22,62927568 | NA | 24,3383093 | 25,64298088 | 24,56871508 | 0,022485163 |
| SLC25A1 | 27,20359176 | 26,83665467 | 26,50218329 | 28,91846298 | 29,73410109 | 28,99792689 | 0,003046206 |
| ITPRIP | NA | NA | NA | 24,08423875 | 24,63682771 | 23,90113353 | 0,006466495 |
| NNT | NA | NA | NA | 23,03768234 | 23,47060327 | 23,34921859 | 0,00113699 |
| PPP2R5A | NA | NA | NA | 23,89365482 | 24,39381725 | 24,15081455 | 0,001873908 |
| SUN2 | 26,27120482 | 26,20017139 | 26,25131294 | 28,5280938 | 28,58132772 | 28,63304832 | 0,002841251 |
| YIPF3 | NA | NA | NA | 22,35466262 | 22,57380804 | 22,65696073 | 0,001629727 |
| UBE2J1 | NA | NA | NA | 24,02909509 | 23,3357905 | 23,20396508 | 0,006466495 |
| NUP188 | NA | NA | NA | 24,46954792 | 24,42265202 | 24,13868053 | 0,000106203 |
| NUP93 | 26,15562814 | 26,15288501 | 26,25988446 | 28,4229006 | 28,63114102 | 28,51690672 | 8,54054E-05 |
| LPCAT1 | 22,87775668 | 23,24851076 | 23,20176857 | 25,10245505 | 25,69840222 | 25,52429469 | 0,000142661 |
| SLC9A1 | NA | NA | NA | 23,26942235 | 23,36706382 | 23,42175359 | 0,001244259 |
| TMEM57 | 24,58388296 | 24,4671239 | 24,79585378 | 26,92388382 | 26,91627184 | 26,98032788 | 0,000467954 |
| ATP5I | 24,16783156 | 23,93615985 | 24,14024189 | 26,11288981 | 26,78967393 | 26,30591256 | 0,002074831 |
| PPAPDC2 | NA | NA | NA | 22,91247129 | 23,11072925 | 22,94984665 | 0,000904019 |
| RAB2A | NA | NA | NA | 24,1662998 | 24,37934611 | 23,79233277 | 0,00113699 |
| CYC1 | 22,99734765 | 23,23641542 | 23,56174103 | 25,24506839 | 25,95969565 | 25,5499247 | 0,001430486 |
| EPHX4 | NA | NA | NA | 23,80479119 | 23,99239418 | 23,76617867 | 0,001421094 |
| HCCS | 21,75429872 | NA | NA | 23,31548018 | 24,12478836 | 22,86230591 | 0,002947384 |
| PDXDC1 | NA | NA | NA | 23,36051491 | 23,46201392 | 22,9480114 | 0,017312668 |
| LRRC59 | 29,64620906 | 29,62472444 | 29,74871213 | 31,82835506 | 32,24623678 | 31,86975143 | 6,96968E-05 |
| ST6GAL1 | 22,16081808 | NA | 22,455557 | 24,18865464 | 24,18010341 | 23,83180746 | 0,002140357 |
| BGN | NA | NA | NA | 23,80872384 | 23,77725933 | 23,68163906 | 0,001477996 |
| RHOT2 | 24,36105063 | 23,99351753 | 24,26026145 | 26,2535688 | 26,90035054 | 26,36897943 | 0,000711822 |
| AAAS | NA | 22,93493693 | 23,6314533 | 24,81382033 | 24,33953308 | 24,40373137 | 0,039814401 |
| EPG5 | 23,94190085 | 24,0924081 | 24,30772001 | 26,33406906 | 26,34915108 | 26,55939425 | 5,39198E-05 |
| PDHB | 22,77683781 | 22,45364943 | 22,51412457 | 24,42636816 | 25,30716411 | 24,91136529 | 0,001202764 |
| DNAJB2 | 20,99975783 | NA | NA | 22,70294711 | 23,22485076 | 23,08066531 | 0,002779591 |
| TTC27 | 23,14060076 | NA | NA | 23,52492234 | 23,63455849 | 23,46861609 | 0,019374947 |
| UTP6 | NA | NA | NA | 22,99875982 | 23,27454992 | 23,37371588 | 0,007733216 |
| RAB3D | 22,65086155 | NA | NA | 23,34272361 | 23,54651394 | 23,43071287 | 0,010402162 |
| AGPAT4 | NA | NA | NA | 22,83304372 | 23,22171278 | 22,96128647 | 0,000196131 |
| EBAG9 | NA | NA | NA | 22,96956569 | 23,36546275 | 23,20702266 | 0,001139823 |
| EMC10 | 22,93851283 | 22,97040856 | 23,16186382 | 25,15271097 | 25,43370819 | 25,28376211 | 0,000829919 |
| MGST3 | 24,35528119 | 23,98580929 | 24,29165187 | 26,17058465 | 26,79191067 | 26,46549006 | 0,002335585 |
| ZWILCH | NA | NA | NA | 22,33043879 | 22,38287201 | 22,40875369 | 0,003035272 |
| SDF2L1 | NA | NA | 23,54132353 | 23,3694621 | 23,81352679 | 23,68720394 | 0,040612674 |
| ADAM15 | NA | 23,41686658 | NA | 25,14293127 | 24,68367487 | 24,50712294 | 0,003880013 |
| TMEM9B | NA | NA | NA | 23,46238844 | 23,58817934 | 23,51869349 | 0,008960851 |
| MGST1 | NA | NA | NA | 23,05304821 | 23,74806588 | 23,29670116 | 0,000240116 |
| TNFRSF10C | NA | NA | NA | 23,72691993 | 23,39427532 | 23,11938381 | 0,002140357 |
| MYCT1 | 22,50489462 | 21,97098775 | 22,59846147 | 24,43440833 | 24,92347584 | 24,49106038 | 0,001173318 |
| STIM1 | 24,10301561 | 23,69720939 | 23,84230123 | 26,1392272 | 26,21324828 | 26,0489935 | 7,54399E-05 |
| SNX4 | 22,75625497 | NA | NA | 23,77223321 | 24,14047595 | 23,80420036 | 0,029559929 |
| APP | 22,88455437 | 22,86953435 | 23,16574796 | 25,26785196 | 25,14580997 | 25,23746615 | 8,34567E-05 |
| VIMP | 24,44170766 | 24,07781647 | 24,25205325 | 26,60865392 | 26,44671645 | 26,44097939 | 0,006381745 |
| COG6 | 24,29466954 | 23,778463 | 24,11941548 | 26,1720928 | 26,27891532 | 26,4312072 | 0,000112581 |
| MCU | 25,62354932 | 25,67823068 | 26,11967273 | 27,8334612 | 28,41086381 | 27,85180231 | 0,013211096 |
| MLST8 | 24,89245091 | 24,39434075 | 24,77409492 | 26,90391659 | 26,80279618 | 27,02612136 | 0,000454529 |
| STX3 | NA | NA | NA | 22,86453681 | 23,13113099 | 22,96287554 | 0,038872973 |
| BET1;DKFZp781C0425 | 23,41247992 | 23,9685993 | 23,65173757 | 25,64416435 | 26,27065249 | 25,76070777 | 0,000299756 |
| BAX | NA | NA | NA | NA | 23,92605778 | 23,36185384 | 0,030501486 |
| VAPB | 24,79728947 | 24,7905939 | 24,64273305 | 26,87126531 | 27,10319575 | 26,88460095 | 0,006890636 |
| GOSR2 | 25,529904 | 23,99877703 | 24,23637163 | 26,61133014 | 26,87712946 | 26,90115655 | 0,002210901 |
| WRB | 21,36677043 | NA | 21,61921159 | 22,42859326 | 23,61728451 | 23,2467435 | 0,008175036 |
| SLC35F6 | 23,67389822 | 23,47975856 | 23,8146028 | 25,54698487 | 25,62086773 | 26,40366637 | 0,000242602 |
| ARVCF | 22,41575827 | NA | NA | 23,76769469 | 23,39113134 | 23,28728965 | 0,004556996 |
| GIMAP1 | 26,35098948 | 26,47572935 | 26,04078285 | 28,29644737 | 28,62780587 | 28,52204334 | 0,038754646 |
| VEZT | NA | NA | NA | 23,26813761 | 23,66579044 | 23,46563017 | 0,02631263 |
| ECE1 | 22,82425102 | 22,7226533 | 23,1061188 | 25,44496456 | 24,29031646 | 25,47559021 | 0,000960049 |
| ILVBL | 25,00812766 | 25,09063258 | 25,21989021 | 27,17464732 | 27,33739079 | 27,35691038 | 0,003383823 |
| EMC1 | 24,39976086 | 24,38916286 | 24,68720394 | 26,69099076 | 26,66207267 | 26,65799111 | 0,000350077 |
| COX4I1 | NA | NA | NA | 22,47367801 | 22,8265609 | 22,58157483 | 0,02378839 |
| TTI2 | NA | NA | NA | 22,83888229 | 23,03101527 | 22,14895441 | 0,006215236 |
| ITGB4 | NA | NA | NA | 22,67292769 | 23,11619799 | 22,96560816 | 0,000521236 |
| SCCPDH | 25,95151506 | 25,66765926 | 25,72614039 | 27,73466642 | 28,28850466 | 27,80501268 | 0,000123747 |
| PPM1F | 25,45192795 | 25,51198544 | 25,52136201 | 27,42973964 | 28,04510239 | 27,47212286 | 0,046629198 |
| EMC7 | 23,89374739 | 23,50978273 | 24,09160132 | 25,8138937 | 26,33944813 | 25,79921798 | 0,00595173 |
| C19orf52 | 22,51188292 | 22,59281651 | 23,17867579 | 24,79039503 | 25,23117957 | 24,71764212 | 0,022136364 |

| | | | | | | | |
|---|---|---|---|---|---|---|---|
| CCDC132 | 24,20754404 | 23,27113357 | 23,95358121 | 25,83103426 | 25,9730087 | 26,07632877 | 0,003425418 |
| FADS1 | NA | NA | NA | 23,18980104 | 23,20016764 | 23,39309713 | 0,002841251 |
| NUP205 | 25,47475509 | 25,38446047 | 25,75052136 | 27,6434764 | 27,73292126 | 27,65758232 | 0,008532031 |
| DCBLD1 | NA | NA | NA | 23,08514695 | 22,88987303 | 23,1260339 | 0,017857376 |
| SLC25A51 | 23,04503571 | NA | NA | 24,05183756 | 23,87773796 | 23,84928673 | 0,025348721 |
| ATP5J2 | 25,16442116 | 25,30319713 | 25,27607754 | 27,25844742 | 27,60116232 | 27,30295318 | 0,016649925 |
| BZW2 | 24,65244894 | 25,2327172 | 25,06669296 | 26,97015839 | 27,07387989 | 27,31630916 | 0,000248739 |
| GLCE | 23,7691082 | 23,83277338 | 23,92732442 | 26,139559 | 26,07905844 | 25,71109096 | 0,003944726 |
| ARHGEF1 | 22,20131989 | 22,20111046 | 23,00005022 | 24,66887676 | 24,59080814 | 24,53984557 | 0,000875979 |
| HSPB1 | 29,83942064 | 29,36778039 | 29,54085814 | 31,47072738 | 31,97394615 | 31,68704038 | 0,00019131 |
| COG4 | 24,01384537 | 23,23734944 | 24,42348578 | 26,15913638 | 26,19277967 | 25,70495837 | 0,010769992 |
| SFXN4 | NA | NA | 22,77064131 | 23,63588725 | 24,29628106 | 23,66589884 | 0,014317191 |
| | 24,95978408 | 24,51044692 | 24,8766612 | 26,84433817 | 27,2680305 | 26,6073845 | 0,000711822 |
| FAM96B | 23,47778395 | NA | 23,62989819 | 24,61868526 | 24,8825035 | 24,409958 | 0,049382544 |
| SLC30A1 | NA | NA | NA | 22,91825254 | 23,07417806 | 23,20955332 | 0,000701702 |
| DCAF8 | 22,96227543 | 23,39073786 | NA | 24,22037006 | 24,33613114 | 24,65283185 | 0,028361384 |
| TBC1D20 | NA | NA | NA | 22,92453635 | 23,19224164 | 23,14622185 | 0,009662543 |
| TMEM63B | NA | NA | NA | 23,74888484 | 23,50008717 | 23,76151958 | 0,026438232 |
| RRAS2 | NA | NA | NA | 22,74833209 | 23,5149673 | 22,76510639 | 0,032105295 |
| SLC35E1 | 24,24829359 | 23,74868014 | 24,26184012 | 26,06747277 | 26,42527994 | 26,04301727 | 0,001677634 |
| STEAP3 | NA | NA | NA | 23,75358484 | 23,92397447 | 23,5553184 | 0,000731272 |
| JAG2 | 24,33851334 | 24,25031924 | 24,3457041 | 26,28029672 | 26,50166714 | 26,38824332 | 0,000116936 |
| NBEAL2 | 27,42494371 | 27,35892332 | 27,59633582 | 29,48229612 | 29,50468554 | 29,62186343 | 0,000504318 |
| CISD2 | 23,74693906 | 23,41312585 | 24,55561096 | 26,06663138 | 25,98943036 | 25,88550889 | 0,000911863 |
| NDUFA5 | 25,57441452 | 25,19955374 | 25,39809781 | 27,38977009 | 27,53482399 | 27,45732426 | 0,000512842 |
| GEMIN4 | NA | NA | NA | 22,3418554 | 22,81333106 | 22,95606556 | 0,005530631 |
| C2orf47 | 22,34564997 | NA | 22,88061785 | 24,19663034 | 24,31333645 | 24,24495955 | 0,002332152 |
| SEC61B | 26,51953356 | 26,62505553 | 26,96024822 | 28,78727802 | 28,78531477 | 28,71561444 | 0,007947095 |
| RAB14 | 24,36699715 | 23,9314784 | 24,15954063 | 25,96717049 | 26,46106159 | 26,21124415 | 0,000611397 |
| OSBPL5 | NA | 23,67292769 | 23,21294041 | 24,61386094 | 24,54916092 | 24,65195649 | 0,037438648 |
| RDH13 | NA | NA | NA | 23,10990078 | 22,9311357 | 23,11998532 | 0,008596158 |
| CYBA | NA | NA | NA | 23,25681105 | 23,58083912 | 23,0924081 | 0,012214 |
| CPT1A | 27,79792023 | 27,50112042 | 27,74188313 | 29,58471818 | 29,95143725 | 29,66817344 | 0,000483066 |
| RAB10 | 22,07933113 | NA | NA | 23,66795697 | 23,75215602 | 23,02621002 | 0,003754515 |
| DAAM1 | 22,5299576 | NA | NA | 24,2801551 | 23,66362064 | 23,62265601 | 0,002509307 |
| RAB18 | 22,45475412 | 22,98829081 | 22,99279177 | 24,4612646 | 25,20947895 | 24,87516176 | 0,000501791 |
| TMEM41B | 24,33572236 | 24,48080647 | 25,00748621 | 26,47628577 | 26,86372425 | 26,5803216 | 0,007143436 |
| EIF2B5 | 22,4888316 | NA | NA | 23,26298717 | 24,07724602 | 23,23520321 | 0,004523062 |
| VAMP3 | 22,02712172 | 21,99862211 | 21,58297622 | 24,23110631 | 23,60975318 | 23,84172557 | 0,000549953 |
| TELO2 | NA | NA | 22,78020653 | 23,417253 | 23,92469944 | 24,25371305 | 0,006466495 |
| MFN2 | 23,81860636 | 24,26363198 | 24,05150569 | 26,04972061 | 26,15855869 | 25,98498406 | 0,006438722 |
| ACSL3 | 27,47606941 | 27,00731511 | 27,15605267 | 29,11909385 | 29,31693489 | 29,2590763 | 0,000307615 |
| SCAMP1 | NA | NA | NA | 23,08857993 | 23,3027092 | 23,21708916 | 0,00060148 |
| PTPN2 | NA | NA | NA | 22,68786602 | 22,7005574 | 22,92012602 | 0,003122699 |
| EXOC7 | 27,80820829 | 27,89833355 | 27,89394407 | 29,87613423 | 29,80448965 | 29,95393638 | 0,004166123 |
| ALDH3A2 | NA | NA | NA | 22,86508456 | 23,0931177 | 23,1568435 | 0,00060148 |
| STOM | 27,64361402 | 27,50521576 | 27,64931353 | 29,30942979 | 29,64454258 | 29,85090865 | 0,000153132 |
| CYB5R3 | 22,07724602 | NA | 21,79910926 | 23,25183661 | 23,8422053 | 23,59046553 | 0,002764089 |
| TMEM126A | 24,56034216 | 24,44018737 | 24,48689393 | 26,19826143 | 26,85233824 | 26,42668807 | 0,011024986 |
| MGAT1 | 25,17047005 | 24,40529055 | 24,76004783 | 26,73298594 | 26,5579639 | 27,02231616 | 0,000705879 |
| TMX2 | NA | 23,38706019 | 23,59696134 | 24,66248019 | 25,05833498 | 24,41267373 | 0,023199817 |
| TSC2 | 22,97494807 | 23,31948249 | 23,48320763 | 25,13680467 | 25,52924871 | 25,06710344 | 0,006215236 |
| APOOL | 24,09087482 | 24,11973208 | 24,15344565 | 26,0393192 | 26,28536821 | 25,98276663 | 0,000270556 |
| VPS51 | 24,57594404 | 24,04568568 | 24,11925715 | 26,0775109 | 26,24103583 | 26,36449458 | 0,003627469 |
| SCARF1 | 24,72935977 | 24,77213251 | 24,66291476 | 26,66519407 | 26,68725735 | 26,75378884 | 0,008441992 |
| STX10 | NA | NA | NA | 22,6661373 | 23,14681227 | 22,92936678 | 0,004039838 |
| EXOC8 | 26,58405506 | 26,16342334 | 26,16417177 | 28,277905 | 28,1749806 | 28,35909094 | 0,000248739 |
| ABHD16A | 25,05916058 | 24,95619853 | 25,03052705 | 26,77994113 | 27,13428013 | 27,02390288 | 0,001845708 |
| SLC25A15 | NA | NA | NA | 21,77245473 | 22,30457637 | 22,34410624 | 0,021725262 |
| TCEA3 | NA | NA | NA | 24,02140566 | 24,09498679 | 23,96886293 | 0,014996883 |
| STUB1 | NA | NA | NA | 22,76813908 | 22,7963391 | 22,41263497 | 0,008960851 |
| COL13A1 | NA | NA | 21,9729167 | 22,79046464 | 22,80057133 | 22,63440339 | 0,010958864 |
| MAGT1 | 26,32678564 | 26,34326597 | 26,39394814 | 28,24049912 | 28,51720717 | 28,17688358 | 0,000623321 |
| MAP2K2 | 22,98194837 | 23,18895641 | NA | 24,76769469 | 24,21423077 | 24,7485266 | 0,009386002 |
| SEMA6B | 23,28559751 | 23,06941633 | NA | 24,43466284 | 24,45650983 | 24,43879236 | 0,026959684 |
| DHRSX | NA | NA | NA | 22,80177545 | 23,5798039 | 23,49350563 | 0,004452287 |
| WDFY3 | 26,65894451 | 26,76014938 | 26,90230721 | 28,7871223 | 28,62125212 | 28,74579853 | 0,000682569 |
| PDS5B | 22,49458023 | 22,94329677 | 23,17397291 | 24,85109935 | 24,78725933 | 24,78126765 | 0,00634236 |
| INF2 | 26,02932683 | 25,85193334 | 26,0305481 | 27,86171452 | 28,04233895 | 27,81341669 | 0,000188716 |
| UBXN4 | 21,78148779 | 21,61234299 | 22,04756728 | 23,81313531 | 23,73758322 | 23,683889 | 0,000467954 |
| CEPT1 | 24,57928601 | NA | 22,56975828 | 25,10561539 | 25,14763535 | 25,27884445 | 0,024938434 |
| PIGN | NA | NA | NA | 22,80335309 | 22,90903168 | 22,83992036 | 0,012214 |
| NGLY1 | NA | NA | NA | 22,51948557 | 22,6707254 | 22,57653232 | 0,002548793 |
| SPCS2 | 25,96730244 | 26,09162149 | 25,96655456 | 27,87882561 | 28,02527662 | 27,88814801 | 0,000460802 |
| TMCC1 | 22,42851659 | 22,44705396 | 21,83945909 | 25,15286567 | 23,60433333 | 23,72359096 | 0,04228797 |
| POMT1 | NA | NA | NA | 22,88390214 | 23,17575542 | 22,93358693 | 0,002874032 |
| MIA2 | NA | NA | NA | 23,03890493 | 22,6967426 | 22,31986834 | 0,002479503 |
| CTPS2 | 23,10149357 | NA | 23,29024614 | 24,36972832 | 24,64801081 | 24,39119691 | 0,010125568 |
| SLC35B2 | 23,94699477 | 23,09958475 | 23,25925593 | 25,26406169 | 25,26087159 | 25,48916246 | 0,000753252 |
| PLD1 | 24,45180222 | 23,73355493 | 24,8727688 | 26,17245528 | 26,45519325 | 26,13567015 | 0,001314272 |
| CLCC1 | NA | NA | NA | 22,84001643 | 22,4927236 | 22,55681568 | 0,000892548 |
| MGAT2 | 24,51225677 | 24,4604524 | 24,25385729 | 26,1891827 | 26,38205403 | 26,35292654 | 0,030727942 |
| SUCLA2 | 21,46811886 | NA | NA | 22,64456046 | 22,88935319 | 22,72980576 | 0,008435773 |
| VRK2 | 26,52224543 | 26,46009301 | 26,80193329 | 28,42842235 | 28,61922419 | 28,39856268 | 0,000750921 |
| ERGIC1 | 24,87586482 | 24,07895668 | 24,85452753 | 26,32159991 | 26,72663415 | 26,41165593 | 0,001666705 |
| ARMCX1 | 24,73164014 | 24,69991171 | 24,54008215 | 26,49612965 | 26,75149216 | 26,36472834 | 0,001400207 |
| CD46 | NA | NA | NA | 22,58968863 | 22,77404463 | 22,654865 | 0,000685815 |
| HIP1R | 27,08151887 | 26,94993568 | 27,06404259 | 28,94976875 | 28,90690069 | 28,8688541 | 0,000659095 |
| MAPRE2 | NA | NA | NA | 22,68799413 | 22,77719912 | 22,81936676 | 0,008532031 |
| PBXIP1 | 24,85771035 | 25,15603338 | 24,79114063 | 26,74406679 | 26,8854158 | 26,78206798 | 0,000398399 |
| TPP1 | 26,37845336 | 26,43766586 | 26,4348696 | 28,16825251 | 28,44805655 | 28,23644462 | 0,000546203 |
| TAP2 | 22,1154679 | 23,68238943 | 23,34299482 | 24,75938759 | 24,98328869 | 24,98997166 | 0,01435588 |
| HACD2 | NA | NA | NA | 22,9146625 | 23,44316311 | 22,50789721 | 0,002808754 |
| SLC39A11 | NA | NA | NA | 23,20282989 | 23,86069676 | 23,26613684 | 0,001539991 |
| ROBO1 | NA | NA | NA | 23,26713757 | 22,73450621 | 23,12815988 | 0,001821957 |
| DNAJC16 | 25,00440325 | 24,39270418 | 24,22736505 | 26,40259341 | 26,47756782 | 26,31390727 | 0,000623321 |
| TMEM39B | NA | NA | NA | 23,06370118 | 23,36091672 | 23,33456354 | 0,001194645 |
| EXOC6 | 26,45274497 | 26,31356135 | 26,75340632 | 28,40641017 | 28,31708589 | 28,33688025 | 0,006286855 |
| IKBIP | 22,48192751 | NA | NA | 23,14782164 | 23,18998196 | 23,413255 | 0,017445109 |
| USMG5 | 25,03858682 | 24,8029687 | 25,22136616 | 26,75544528 | 27,04082464 | 26,7912897 | 0,046629198 |
| TBC1D5 | 25,46007738 | 25,0116719 | 25,31014955 | 27,10099255 | 27,1447214 | 27,05641363 | 0,002024195 |
| TRABD | NA | NA | NA | 22,99548562 | 24,2365906 | 23,67335911 | 0,018355824 |
| LPGAT1 | 23,17949608 | 23,88147697 | 22,77739981 | 25,13899294 | 25,30138399 | 24,91415152 | 0,003834596 |

| | | | | | | | |
|---|---|---|---|---|---|---|---|
| TBC1D22B | 23,61706026 | 23,60783598 | 23,75154324 | 25,59063684 | 25,4592645 | 25,43784044 | 0,002201487 |
| RAB27A | NA | NA | 21,97683678 | 23,20594952 | 22,91672286 | 23,05928438 | 0,014854056 |
| NDUFAF2 | 23,35944288 | 22,93792046 | 23,19049447 | 24,9652999 | 25,2093302 | 24,80272224 | 0,000416304 |
| PYGB | 24,52898056 | 24,30250004 | 24,05308137 | 26,02985337 | 26,2925473 | 26,02930576 | 0,000806115 |
| HLTF | NA | NA | NA | 22,59149313 | 22,52571101 | 22,49194115 | 0,002159725 |
| SEC11A | 25,00782835 | 24,35709501 | 24,93355091 | 26,6768059 | 26,56024011 | 26,50954114 | 0,002102009 |
| SELK | 22,39827399 | 22,59482209 | 22,73032418 | 24,26148148 | 24,62153859 | 24,28234867 | 0,001737142 |
| ARL1 | NA | NA | NA | 22,28948648 | 22,57161101 | 22,86832898 | 0,010603097 |
| SPTLC1 | 24,33060291 | 24,50227436 | 24,54232768 | 26,27303847 | 26,20024624 | 26,3352794 | 0,000623321 |
| SYMPK | 23,32443549 | NA | NA | 24,28340888 | 23,61380475 | 23,64713252 | 0,025957979 |
| SMN1;SMN2 | 22,03365556 | 21,47105001 | 21,16177158 | 23,42828656 | 23,45035547 | 23,19801032 | 0,002799114 |
| NSDHL | 24,92029869 | 25,2507168 | 25,24811259 | 26,60215401 | 27,03485638 | 27,17540899 | 0,020866462 |
| TMED7-TICAM2;TMED7 | 24,62578018 | 24,79183616 | 25,11727657 | 25,87981898 | 27,23999862 | 26,7948876 | 0,01552091 |
| FRMD5 | NA | NA | 21,62283472 | 23,33429075 | 23,4947267 | 23,5553184 | 0,000938125 |
| TMEM120B | NA | NA | NA | 22,71355769 | 23,66394632 | 22,91201437 | 0,001540242 |
| OCIAD1 | 24,67260403 | 24,27859635 | 24,41918354 | 26,25747661 | 26,26653007 | 26,2203147 | 0,01619879 |
| OGFR | 22,95496588 | 22,57475519 | 22,30946969 | 24,17256973 | 24,65987006 | 24,37980879 | 0,032073028 |
| QSOX1 | 22,3953741 | 22,44760894 | NA | 23,83557089 | 23,49326129 | 23,35769911 | 0,026806268 |
| LEPREL4;P3H4 | NA | NA | NA | 22,7411888 | 22,99482989 | 22,83010587 | 0,028810693 |
| RAB29 | 24,08642717 | 23,55180306 | 23,90168597 | 25,44078935 | 25,74752819 | 25,70930388 | 0,001183378 |
| TMEM106B | NA | NA | NA | 22,97405532 | 23,44050423 | 23,08259932 | 0,020379684 |
| UQCRQ | 25,43186075 | 24,98858553 | 24,91533746 | 26,59875109 | 27,46302801 | 26,61400141 | 0,007610809 |
| ST6GALNAC4 | NA | NA | NA | 22,68671253 | 23,14458917 | 22,5022015 | 0,037424859 |
| IRAK1 | NA | NA | NA | 22,65896629 | 23,11313233 | 22,63205268 | 0,000906428 |
| SLC25A19 | 22,71852033 | 22,07564758 | 22,79546738 | 24,01750304 | 24,78461608 | 24,10501586 | 0,002224731 |
| UMPS | 22,55950219 | 22,52518528 | 22,74935553 | 24,33469992 | 24,34238452 | 24,46793236 | 0,001074128 |
| PTPRB | 25,02301453 | 25,04880646 | 25,11941548 | 26,92716615 | 26,72961908 | 26,83966091 | 0,005591359 |
| JUP | 24,58588953 | 24,4058099 | 24,83658244 | 26,34980913 | 26,23609786 | 26,54126444 | 0,041859482 |
| TMEM131 | 24,12020686 | 23,93012515 | 24,07716451 | 25,87902434 | 25,73528641 | 25,81198473 | 0,0002364 |
| EMC8 | 24,5579493 | 24,42162519 | 25,03875425 | 26,4826231 | 26,37120833 | 26,46001487 | 0,001907016 |
| CTDNEP1 | 22,91613969 | 22,50751013 | 22,54366152 | 23,88585795 | 25,4266241 | 23,94859965 | 0,009419542 |
| RHOT1 | 24,20471143 | 23,95171514 | 23,85148066 | 25,51761269 | 26,11086065 | 25,65452634 | 0,001375644 |
| DHCR24 | 22,83283131 | 22,44705396 | 23,3340179 | 24,45888918 | 24,55549394 | 24,87140633 | 0,004554412 |
| SEC63 | NA | NA | NA | 22,70364424 | 22,743656 | 22,43150373 | 0,002253125 |
| PPP2R5E | 22,87712009 | NA | 22,41201466 | 23,73954166 | 23,89411759 | 23,96815982 | 0,027980925 |
| PTPN1 | 24,52025323 | 24,19843005 | 24,44366901 | 25,89873718 | 26,383851 | 26,09359729 | 0,005218333 |
| RAB34 | 23,14250286 | 23,57830727 | NA | 24,30820625 | 24,55104027 | 24,45519325 | 0,047658873 |
| PAM16;CORO7-PAM16 | 24,52282909 | 25,160695 | 25,16350012 | 26,48718536 | 27,13408424 | 26,43447196 | 0,002166222 |
| VPS4A | 24,09675697 | 23,93678888 | 24,05639296 | 25,68303229 | 26,01586674 | 25,59602297 | 0,001515589 |
| EXOC2 | 25,92322646 | 25,72788079 | 25,78154282 | 27,50831743 | 27,49663256 | 27,6243584 | 0,001525715 |
| NUP35 | 21,83124693 | NA | 21,22670382 | 23,23363898 | 23,46861609 | 22,4245241 | 0,016649925 |
| GALNT6 | NA | NA | NA | 22,93140626 | 23,19276838 | 22,95631376 | 0,008070106 |
| FZD8 | NA | NA | NA | 22,97542048 | 22,77076228 | 22,66349035 | 0,001555445 |
| EXD2 | 24,54368512 | 24,18593585 | 24,83956481 | 26,09315398 | 26,41357783 | 26,22273006 | 0,0024656 |
| LPCAT4 | NA | NA | NA | 22,55688582 | 22,81372249 | 22,5711712 | 0,004139228 |
| DOCK4 | 30,04791253 | 30,72077614 | 30,64727322 | 32,20411088 | 32,09490376 | 32,25172038 | 0,00514457 |
| STAT2 | NA | 23,13517694 | 23,31755173 | 24,05498646 | 24,11664221 | 24,18495281 | 0,03855599 |
| TOMM40 | 22,69366221 | 22,97161933 | 22,53388291 | 24,12549798 | 24,6865095 | 24,50579121 | 0,003555585 |
| SLC25A3 | 26,83520944 | 26,33815626 | 27,06732915 | 28,45238366 | 28,76673473 | 28,13887579 | 0,030421219 |
| TUBB8 | 27,92422372 | 27,0155265 | 27,93038462 | 29,10119298 | 29,49830763 | 29,38337725 | 0,025194224 |
| CTNNB1 | 25,19027592 | 24,80346149 | 25,29302113 | 26,62142681 | 26,92660076 | 26,83050244 | 0,003194167 |
| NDUFB6 | NA | NA | NA | 22,58228718 | 23,17926827 | 22,3965243 | 0,005102965 |
| CRYBG3 | NA | NA | NA | 22,73022051 | 22,88427488 | 22,81911334 | 0,020574954 |
| SFXN3 | 28,27644144 | 28,3478172 | 27,57484757 | 29,56214514 | 30,05711119 | 29,63509153 | 0,003637798 |
| PEX5 | NA | NA | NA | 23,36479512 | 23,34921859 | 23,16793873 | 0,028926715 |
| METTL7A | 23,55403044 | 23,58359608 | 23,96013772 | 25,15911713 | 25,4252159 | 25,51809314 | 0,01882603 |
| VPS8 | NA | NA | NA | 22,6319195 | 22,59627895 | 22,52637984 | 0,008435773 |
| VAC14 | 25,93981789 | 26,02286642 | 26,09065277 | 27,74611899 | 27,5644184 | 27,708791 | 0,000367308 |
| YWHAQ | 24,1586935 | 24,089744 | 24,40249583 | 25,71379364 | 25,99425157 | 25,89296038 | 0,000472586 |
| TTC17 | 22,53692067 | 23,00691294 | NA | 23,69614827 | 24,13961755 | 23,74437481 | 0,031610035 |
| TUBB6 | 29,55380346 | 29,21576804 | 29,78492491 | 31,10774172 | 31,30012959 | 31,09074614 | 0,009601348 |
| CRIM1 | 22,949722 | 23,16810711 | 22,24222861 | 24,35890656 | 24,68645607 | 24,24966845 | 0,003397624 |
| ATP5A1 | 28,76430038 | 28,64571103 | 28,63346412 | 29,99010154 | 30,7139116 | 30,26360512 | 0,000777061 |
| TIMM10 | NA | NA | NA | 22,54729089 | 22,73287211 | 22,65110251 | 0,019374947 |
| ZFPL1 | 23,88753226 | 24,20224701 | 24,04460223 | 25,50124193 | 25,87452872 | 25,67473343 | 0,000820737 |
| TTC28 | 24,12793955 | 24,57016377 | 24,53575853 | 25,97705523 | 25,99931483 | 26,16697014 | 0,000711822 |
| PHKA1 | 23,21295525 | 22,97281155 | 23,5731146 | 24,79054418 | 24,86556602 | 25,00406029 | 0,003627469 |
| PECAM1 | 24,10629456 | 23,98320169 | 24,36793032 | 25,7645092 | 25,75322777 | 25,82708932 | 0,001153493 |
| FAM69B | 24,31920683 | 24,48885611 | 24,41254452 | 26,02827318 | 26,00643789 | 26,07175416 | 0,000493025 |
| CUL4A | NA | NA | 21,12949685 | 22,38094506 | 22,21646761 | 21,95928883 | 0,021383396 |
| EXOC6B | 25,85733068 | 25,75307471 | 25,84756742 | 27,42101517 | 27,43920466 | 27,47699644 | 0,001596805 |
| APOL2 | 22,78280789 | 23,12128245 | 23,53338392 | 24,62299107 | 24,76253371 | 24,92646504 | 0,00368695 |
| FARSA | 27,48200755 | 27,12435453 | 27,53675174 | 28,81787483 | 29,20347508 | 28,99482881 | 0,022802811 |
| TBC1D10A | NA | 22,77725933 | NA | 24,19828016 | 23,84814075 | 23,71732834 | 0,006144988 |
| EXOC4 | 26,33502378 | 26,13449557 | 26,14100244 | 27,76014938 | 27,88913499 | 27,81720394 | 0,006908778 |
| HSPA12B | NA | NA | NA | 22,66490122 | 22,94302843 | 22,67521279 | 0,001655102 |
| GNAQ | NA | NA | NA | 22,32113538 | 22,41160098 | 22,28717691 | 0,02282084 |
| CLCN7 | NA | NA | NA | 22,98072882 | 22,87578984 | 22,59896117 | 0,025343699 |
| TTC13 | 22,89459872 | 22,83980505 | 23,04383499 | 24,59428678 | 24,58290734 | 24,41286751 | 0,00033803 |
| B4GALT7 | NA | NA | NA | 22,6507301 | 23,41286751 | 22,97956056 | 0,002039808 |
| NCEH1 | NA | NA | 21,30791453 | 22,39516487 | 23,30131421 | 22,21445312 | 0,016775776 |
| NCLN | 25,30580821 | 25,23209502 | 25,23629863 | 26,68137097 | 26,95947457 | 26,93719312 | 0,002948997 |
| ATR | 25,14666469 | 24,84618093 | 24,93310062 | 26,59120776 | 26,41099315 | 26,72455403 | 0,000575523 |
| ZDHHC24 | NA | NA | NA | 22,6949164 | 23,48615742 | 23,02160898 | 0,016353668 |
| LRRC8C | 25,06262295 | 24,91255352 | 25,07496199 | 26,66302338 | 26,52049304 | 26,64402679 | 0,039657963 |
| SLC25A22 | 24,32855003 | 24,54096896 | 24,4373325 | 25,77816217 | 26,18072944 | 26,12192663 | 0,00748928 |
| SLC4A2 | NA | NA | NA | 23,00542307 | 23,00971724 | 23,18783954 | 0,009019741 |
| MGRN1 | NA | NA | NA | 22,48674666 | 22,2629585 | 21,99355209 | 0,012781607 |
| MIA3 | NA | NA | NA | 22,66431533 | 22,47447043 | 22,66912555 | 0,025348721 |
| SPCS3 | 24,29775088 | 24,36131842 | 24,51665129 | 25,93265019 | 26,16065654 | 25,83489612 | 0,000531651 |
| RETSAT | 22,86614168 | 22,20442786 | 23,85319532 | 24,66041423 | 24,50269926 | 24,49661732 | 0,010652086 |
| CNP | 26,79995902 | 26,77580356 | 26,65348801 | 28,13824109 | 28,51927109 | 28,29162552 | 0,000465699 |
| CSGALNACT1 | NA | NA | NA | 22,56353425 | 22,88879601 | 22,48853744 | 0,014038482 |
| RHOG | 26,31461616 | 26,54591033 | 26,33851334 | 27,84062159 | 28,10908777 | 27,94514249 | 0,012155628 |
| LTN1 | 23,63455849 | 23,12943396 | 23,92370251 | 24,84804521 | 25,12730986 | 25,38827617 | 0,002210901 |
| RAB13 | 25,76547065 | 25,54568944 | 25,53789234 | 27,2023778 | 27,17826167 | 27,13398629 | 0,003209812 |
| QSOX2 | NA | NA | NA | 22,44874346 | 23,05629372 | 22,96522062 | 0,020658564 |
| S100A16 | 23,55485018 | 23,35581886 | 23,38126199 | 24,76991531 | 25,23019023 | 24,93737274 | 0,028444564 |
| COPG2 | 22,88280199 | 23,42790308 | 22,81080381 | 24,78286787 | 24,58777889 | 24,39486405 | 0,001845402 |
| KNTC1 | 22,19002719 | NA | NA | 23,40600461 | 23,73252022 | 23,64504444 | 0,02378839 |
| ACADM | 23,68528006 | 23,20767808 | 23,27156105 | 24,5676131 | 25,39581207 | 24,8347997 | 0,003099117 |
| HSP90AB1 | 27,06465939 | 27,16288577 | 27,42822265 | 28,36582148 | 29,06090311 | 28,8459836 | 0,001400207 |
| RASIP1 | 29,03209951 | 28,99676926 | 28,89015028 | 30,49342928 | 30,59396992 | 30,44164039 | 0,00897884 |

| | | | | | | | |
|---|---|---|---|---|---|---|---|
| RAB6A | 24,56511603 | 23,91168529 | 24,61610682 | 26,13987122 | 25,83980505 | 25,69991171 | 0,025348721 |
| CDH5 | 24,99040455 | 25,26881581 | 25,26942235 | 26,57301344 | 26,85792387 | 26,67559508 | 0,022277449 |
| PIKFYVE | 21,63338374 | NA | NA | 22,14960573 | 22,56239337 | 22,14985377 | 0,021342803 |
| LRRC8A | 26,03088482 | 26,18427184 | 26,04597724 | 27,53689992 | 27,67155165 | 27,61372046 | 0,02378839 |
| MAN1B1 | 24,11052218 | 24,30848402 | 24,4684918 | 25,86747614 | 25,77021785 | 25,79213415 | 0,001314272 |
| PEX6 | 23,84517608 | 23,61313028 | 23,3754404 | 25,59662018 | 25,07806088 | 24,70054682 | 0,011842761 |
| TBRG4 | NA | NA | NA | 22,90350752 | 22,99232502 | 22,58497259 | 0,026478342 |
| SLC25A11 | 28,4410269 | 28,16442116 | 28,32932876 | 29,57342163 | 30,168659 | 29,69521252 | 0,001152601 |
| HEATR3 | 22,78626245 | 22,04536907 | 22,38498079 | 23,68431716 | 24,00923884 | 24,0232684 | 0,001756141 |
| S100A6 | 27,25502871 | 27,12563592 | 27,24181779 | 28,287228 | 28,91530326 | 28,89530277 | 0,001401403 |
| SGPL1 | 25,64366906 | 25,35857126 | 25,64647345 | 27,05786005 | 26,95404735 | 27,10159375 | 0,001244259 |
| GLS | NA | 22,3589602 | NA | 22,50579121 | 22,8681594 | 22,71305419 | 0,040721041 |
| MON2 | 24,66968786 | 24,34353708 | 24,63056487 | 25,865094 | 26,11402252 | 26,11094028 | 0,002148142 |
| CDC2;CDK1 | 25,00097 | 24,55613742 | 25,1215117 | 26,3578501 | 26,47812353 | 26,27120482 | 0,019116371 |
| DDX20 | 23,5149673 | 23,56523227 | 23,57265212 | 25,26985035 | 24,79168715 | 25,01864947 | 0,001148652 |
| LPCAT2 | 25,27873813 | 25,15147277 | 25,05378572 | 26,64044553 | 26,5862046 | 26,68177308 | 0,004970342 |
| IDH2 | 28,67023509 | 28,30630379 | 28,57582145 | 29,75542936 | 30,35237095 | 29,86696938 | 0,031448416 |
| RAB32 | 23,7509302 | 23,24353662 | 23,89652162 | 24,9788452 | 25,25082521 | 25,07802014 | 0,0278232 |
| RAB5A | 22,95881111 | 22,74546261 | 22,47409903 | 24,12959118 | 24,23000695 | 24,22736505 | 0,000778727 |
| ATP5H | 25,11814836 | 25,38715882 | 25,50020876 | 26,72832205 | 27,02495972 | 26,65812735 | 0,008310832 |
| ACSL1 | 21,88751367 | NA | 22,01984588 | 22,72750172 | 23,58680589 | 23,15741417 | 0,006947519 |
| EXOC1 | 25,54548324 | 25,36753046 | 25,54079164 | 26,92998976 | 26,88051977 | 27,00806353 | 0,003039528 |
| PPP2R5D | 24,31755173 | 24,54132353 | 24,77565288 | 25,92585411 | 26,01327036 | 26,05465532 | 0,029928407 |
| GPR56;ADGRG1 | NA | NA | NA | 22,70252444 | 22,9995686 | 22,66817344 | 0,032843387 |
| LRRC8D | 23,32072234 | 23,14293906 | 23,25209657 | 24,69365157 | 24,64157596 | 24,70567035 | 0,001285544 |
| ADAM10 | 24,24887264 | 24,54527701 | 24,6719565 | 25,98578757 | 26,02068532 | 25,77766066 | 0,024828045 |
| MTCH2 | 25,84809298 | 25,64226482 | 25,53418579 | 27,02538224 | 27,36669707 | 26,92546932 | 0,010610048 |
| ARF1;ARF3 | 25,17945812 | 24,65053291 | 25,2492706 | 26,48597324 | 26,48651038 | 26,37421355 | 0,013180716 |
| TAMM41 | 25,34523034 | 23,34921859 | 24,65605514 | 25,48683257 | 26,20635204 | 25,92456354 | 0,035972439 |
| COG7 | 25,381295 | 25,18457453 | 25,35043315 | 26,86171452 | 26,68805818 | 26,632147 | 0,001524978 |
| RAB11B;RAB11A | 23,40262593 | 23,37729527 | 23,51038655 | 24,66811933 | 25,42997902 | 24,44846621 | 0,011546131 |
| THEM6 | 26,89804518 | 25,61214611 | 25,54518862 | 27,44150189 | 27,4239826 | 27,44063096 | 0,033652924 |
| RNF123 | NA | NA | NA | 22,31307345 | 22,06193938 | 22,1388055 | 0,005476931 |
| STAT1 | 26,28790609 | 25,94911194 | 25,91779745 | 27,44237229 | 27,34908357 | 27,57996211 | 0,007610809 |
| MT-ATP8 | 24,76283781 | 24,40360137 | 24,34353708 | 25,70245046 | 26,08529285 | 25,93613738 | 0,002140357 |
| HSD17B12 | 27,85204053 | 27,38747111 | 27,7997738 | 28,7093499 | 29,60284425 | 28,93612614 | 0,007216199 |
| RANBP6 | NA | 22,42141974 | NA | 23,39073786 | 23,79143875 | 23,42123994 | 0,015952693 |
| DNAAF5 | 23,92886098 | 23,57045333 | 23,74057134 | 25,14802343 | 25,2127846 | 25,05250105 | 0,004554412 |
| RPTOR | 24,69083095 | 24,91141101 | 24,88338946 | 25,99660227 | 26,3579843 | 26,30340619 | 0,010450864 |
| RAB22A | NA | NA | NA | 23,07237994 | 22,94858183 | 23,05854143 | 0,002520668 |
| CUL4B | NA | NA | NA | 22,71584221 | 22,56523227 | 22,64934916 | 0,018071901 |
| FUNDC2 | 21,26985035 | NA | NA | 22,43659566 | 22,41361657 | 22,97573534 | 0,014951723 |
| EMC2 | 24,74658034 | 24,8624478 | 24,91852553 | 26,13322203 | 26,26801264 | 26,26968987 | 0,015821752 |
| GNB1 | 24,28933168 | 24,51382348 | 23,34854339 | 25,3146853 | 25,49237522 | 25,47540467 | 0,010952083 |
| BAG2 | NA | NA | 21,73036565 | 23,37901552 | 22,83535885 | 22,84682175 | 0,014994316 |
| NDUFC2 | 24,53498721 | 24,85709334 | 24,87286272 | 26,65662802 | 26,4239826 | 25,30155843 | 0,027616238 |
| TMEM33 | 24,88822932 | 24,80764343 | 25,33302838 | 26,15238218 | 26,50292683 | 26,48331528 | 0,002008383 |
| GCDH | 24,66302338 | 24,81567802 | 24,57675143 | 25,88550889 | 26,29084374 | 25,98378881 | 0,006672978 |
| RAC1;RAC3 | 24,21955792 | 24,14234705 | 24,33796918 | 25,58296474 | 25,51580954 | 25,70242404 | 0,030488169 |
| ELMO2 | 28,14345686 | 28,28034098 | 28,13828993 | 29,58478985 | 29,49321929 | 29,5650906 | 0,001691478 |
| PANK4 | 23,09971315 | 23,1305184 | 22,95659736 | 24,42059763 | 24,48049834 | 24,35944288 | 0,00454604 |
| ACTB;ACTG1 | 30,63875512 | 30,63961782 | 30,87554848 | 31,92351551 | 32,17352194 | 32,10842737 | 0,001375644 |
| EIF2AK3 | NA | NA | NA | 22,21859753 | 22,8464392 | 22,54222139 | 0,046629198 |
| WNK1 | 21,65796386 | NA | 22,97706396 | 23,21557921 | 23,49497079 | 23,39623684 | 0,026667557 |
| NOMO1 | NA | NA | NA | 23,31506551 | 23,06340496 | 23,07043218 | 0,004383602 |
| DCAKD | 24,39126247 | 24,25695498 | 24,25306379 | 25,57045333 | 25,86284977 | 25,48588113 | 0,007039273 |
| VDAC2 | 26,93044102 | 26,48758406 | 27,1270146 | 27,9699389 | 28,58373236 | 27,99614968 | 0,004383602 |
| FHOD1 | 25,59129337 | 25,84038148 | 25,9276409 | 27,12030576 | 27,09968907 | 27,10888844 | 0,006038569 |
| OSBPL8 | 24,99602034 | 25,19861739 | 24,70128742 | 26,28728965 | 26,2761308 | 26,28651433 | 0,006058234 |
| MTDH | 26,24056281 | 26,23469174 | 26,1614064 | 27,60703169 | 27,42686399 | 27,54951349 | 0,001030974 |
| ACAD9 | 24,7194189 | 24,58938004 | 24,70956682 | 25,72538644 | 26,35361651 | 25,87907109 | 0,038037952 |
| NFXL1 | 26,31195171 | 26,36659703 | 26,1477906 | 27,65601421 | 27,51487703 | 27,55800771 | 0,001825212 |
| STK25 | NA | 22,84441001 | 23,28728965 | 23,56092519 | 23,64460446 | 24,35655782 | 0,048572741 |
| C3orf58 | 24,6549633 | 24,80356003 | 24,56645221 | 25,94806489 | 26,00579568 | 25,97064115 | 0,011832855 |
| CLEC14A | 26,15410268 | 26,10860932 | 26,13684377 | 27,40584236 | 27,35447431 | 27,52857823 | 0,011891317 |
| NDUFA11 | 22,89659553 | 22,36236231 | 23,45199082 | 24,10693348 | 24,2898945 | 24,18237859 | 0,014854056 |
| NOS3 | 26,11424101 | 26,02572017 | 26,32333629 | 27,38212001 | 27,44986765 | 27,48998928 | 0,006488801 |
| CTPS1 | 26,08285916 | 25,84969238 | 26,20888387 | 27,19037013 | 27,57419795 | 27,225454 | 0,003407435 |
| CNTNAP3;CNTNAP3B | 24,12376273 | 23,70229192 | 23,75979393 | 25,16061807 | 25,11310848 | 25,14763535 | 0,003209812 |
| PARL | NA | NA | NA | 22,66116484 | 23,1730732 | 23,10932694 | 0,010881236 |
| TIMM44 | 25,40590726 | 25,66294191 | 25,75026577 | 26,91809328 | 27,22811775 | 26,48793667 | 0,003534119 |
| NF1 | 22,53328885 | 22,50110827 | 22,52382227 | 24,57398134 | 23,24176325 | 23,5406143 | 0,039744669 |
| LRRC41 | NA | NA | NA | 22,99337932 | 22,86819709 | 22,47778395 | 0,011822399 |
| SFXN1 | 27,98321256 | 27,53504656 | 28,09380882 | 28,93999719 | 29,68970849 | 28,72660167 | 0,009814723 |
| RNF213 | 27,92552592 | 27,98554874 | 27,95360341 | 29,16662543 | 29,16079597 | 29,26760643 | 0,022097707 |
| SLC25A13 | 27,61673784 | 27,64044553 | 27,88069491 | 28,69985216 | 29,18718732 | 28,96694504 | 0,00863886 |
| PI4KA | 25,69654629 | 25,57476097 | 25,61113312 | 26,70081136 | 27,02125738 | 26,84876161 | 0,003160893 |
| ATP5O | 26,68992505 | 26,63505692 | 26,55398651 | 27,64547055 | 28,05693038 | 27,82165041 | 0,010414429 |
| ARL8A;ARL8B | NA | 22,03429388 | 22,14886134 | 22,58927716 | 23,09740013 | 22,75344202 | 0,034959986 |
| VEPH1 | NA | NA | NA | 22,2776461 | 21,99458823 | 22,29686917 | 0,00821971 |
| ATF6 | NA | NA | NA | 22,78676098 | 22,17289014 | 22,43451014 | 0,035934548 |
| DHRS1 | 23,72961908 | 23,83913227 | 23,63843061 | 24,30716411 | 25,60221066 | 24,91515507 | 0,017342931 |
| NFIX | NA | NA | NA | 22,69981642 | 22,60584842 | 21,98919213 | 0,030673937 |
| ARMCX3 | 24,36979487 | 24,57871036 | 24,46138952 | 25,66370207 | 25,62301898 | 25,71313811 | 0,015740186 |
| ARF5 | 24,67093065 | 24,4019102 | 24,83050244 | 25,85031257 | 25,90398552 | 25,73130349 | 0,020922351 |
| PDS5A | 25,35940936 | 25,3086576 | 25,25778231 | 26,38332368 | 26,59149313 | 26,5145761 | 0,001991154 |
| ACSL4 | 23,85443242 | 24,00958057 | 24,13766474 | 25,11016371 | 25,40399135 | 25,03992575 | 0,004748446 |
| MMRN2 | 26,89087506 | 27,08648792 | 26,44837169 | 27,57217504 | 28,07065941 | 28,31630916 | 0,020658564 |
| COX6C | 24,4732073 | 24,83113093 | 24,95668598 | 25,65181967 | 26,54443707 | 25,59815475 | 0,044595692 |
| FMNL3 | 27,51435035 | 27,77479872 | 27,5744867 | 28,74970333 | 28,82089 | 28,79417493 | 0,004166123 |
| PNPLA6 | 25,17020259 | 25,07993329 | 24,91720561 | 26,36043119 | 26,18850373 | 26,11459846 | 0,047090649 |
| PEX1 | 24,19475325 | 24,09345625 | 24,22110797 | 25,30924763 | 25,3232332 | 25,36205457 | 0,010914623 |
| EHD4 | 29,75015075 | 29,85217451 | 29,69365157 | 30,86016097 | 31,03400362 | 30,88561944 | 0,00231509 |
| RAB8A | 23,76941092 | 23,73820196 | 24,28256077 | 25,16652966 | 24,96146312 | 25,14074897 | 0,00460468 |
| XPOT | 23,6027487 | 23,54002301 | 24,18820186 | 24,67432936 | 25,09852498 | 25,01427116 | 0,018218662 |
| KIAA0355 | 25,02369142 | 25,01013571 | 24,89161685 | 26,11024338 | 26,14367094 | 26,12336807 | 0,008435773 |
| MAGED1 | 25,102415 | 25,11921756 | 25,44135941 | 26,29904448 | 26,47804636 | 26,32808773 | 0,041395768 |
| DNAJB6 | 22,34196395 | 22,11762535 | 22,56404619 | 23,32882391 | 23,71931445 | 23,39060667 | 0,007216199 |
| STAB1 | 24,3778248 | 24,68741755 | 24,92016237 | 25,76554652 | 25,83968493 | 25,78616272 | 0,007743766 |
| DDOST | 24,26406169 | 25,36806359 | 24,51707198 | 25,42755147 | 25,8791646 | 26,23680954 | 0,037451441 |
| MTG2 | NA | NA | NA | 21,71452224 | 21,87150973 | 21,74377925 | 0,006963342 |
| TRIM4 | 25,15853943 | 25,33715256 | 25,19381379 | 26,34168915 | 26,35156244 | 26,3824169 | 0,009178643 |
| SLC25A29 | 24,15730622 | 24,15915563 | 23,98979847 | 25,1562263 | 25,26237791 | 25,25425388 | 0,004877392 |

| | | | | | | | |
|---|---|---|---|---|---|---|---|
| ELMO1 | 28,23196689 | 28,30708592 | 27,92682694 | 29,27659674 | 29,26228381 | 29,28786647 | 0,038459149 |
| COG8 | 23,77776098 | 23,5843992 | 23,24190869 | 24,18108976 | 24,90554713 | 24,87591167 | 0,049045448 |
| SFXN2 | 24,09192409 | 24,34840831 | 24,21044543 | 25,12206491 | 25,67656382 | 25,1669127 | 0,032313466 |
| CTNNA1 | 25,35039942 | 25,32433247 | 25,91179956 | 26,91239362 | 26,0535993 | 26,91182242 | 0,021385268 |
| HM13 | NA | NA | NA | 22,70186906 | 22,72014988 | 22,51542458 | 0,041040287 |
| HYOU1 | 22,96171039 | 22,40914229 | 22,58836237 | 23,31824158 | 23,97971755 | 23,92858994 | 0,016791221 |
| SLC25A12 | 27,04666425 | 26,97585339 | 26,83134842 | 27,68992505 | 28,39737982 | 28,02226324 | 0,036946623 |
| WDR61 | 24,33449535 | 24,17584677 | 24,29691117 | 25,47342409 | 25,44774134 | 25,14156772 | 0,003649987 |
| GALK1 | 24,80459427 | 24,70197479 | 24,46369848 | 25,54474658 | 25,95881111 | 25,6748681 | 0,029597961 |
| SH3BP4 | 26,11812855 | 26,00945243 | 26,1176729 | 27,18196174 | 27,17111937 | 27,07856993 | 0,005254074 |
| OXA1L | 24,8146517 | 25,60902043 | 25,37931305 | 26,28211886 | 26,46163931 | 26,23128945 | 0,028960271 |
| GEMIN5 | 25,56104177 | 25,70888307 | 25,79116547 | 26,88180367 | 26,62016858 | 26,71764212 | 0,014994316 |
| TIMM50 | 26,64718743 | 26,35368381 | 26,62044828 | 27,26865521 | 27,8754899 | 27,62373071 | 0,022436251 |
| NDUFAF4 | 25,98561388 | 24,91756985 | 25,20172371 | 26,24758756 | 26,70147251 | 26,27478094 | 0,033062078 |
| EIF4A1 | 28,7362159 | 28,782318 | 29,0589697 | 29,9433549 | 29,86973672 | 29,83018507 | 0,026806268 |
| THBS1 | 30,89489819 | 30,85630686 | 30,62400972 | 31,7779992 | 31,82937622 | 31,81989325 | 0,00469906 |
| ACBD5 | NA | NA | NA | 22,35987179 | 22,56818159 | 22,32077742 | 0,010883419 |
| UNC45A | 29,19387487 | 29,2420223 | 29,47875231 | 30,51163866 | 30,35877664 | 30,04557113 | 0,040459904 |

### Table S4. GO term enrichment analysis of SEC22B interactome hits.

| #term ID | ONTOLOGY | Description | Gene Ratio | Background Ratio | FDR | matching proteins in your network (gene ID) | Count |
|---|---|---|---|---|---|---|---|
| GO:0034220 | BP | ion transmembrane transport | 119/765 | 182/2446 | 1,13513E-06 | ATP11C/MAGT1/VDAC2/SFXN3/PEX1/TAP1/PARL/SLC33A1/TMEM120B/PEX5/ATP2C1/SLC30A6/ACSL1/WNK1/PHB2/SLC3A2/TRAM | 119 |
| GO:0055085 | BP | transmembrane transport | 129/765 | 210/2446 | 4,90579E-05 | ATP11C/MAGT1/VDAC2/SFXN3/PEX1/TAP1/RNF5/PARL/SLC33A1/TMEM120B/PEX5/ATP2C1/SLC30A6/ERLIN2/ACSL1/WNK1/PHB2/ | 129 |
| GO:0098656 | BP | anion transmembrane transport | 66/765 | 92/2446 | 5,885342028 | VDAC2/SFXN3/PEX1/TAP1/PEX5/ACSL1/SLC3A2/TRAM1/CLCN7/SLC25A11/SLC16A3/ABCC4/TIMM44/PRAF2/SLC25A12/TOMM40/S | 66 |
| GO:0010257 | BP | NADH dehydrogenase complex assembly | 33/765 | 35/2446 | 43,12245308 | NDUFB6/NDUFA6/NDUFV2/NDUFB5/TMEM126B/NDUFB8/NDUFV1/NDUFB10/OXA1L/NDUFS8/NDUFS4/NDUFA2/NDUFS5/NDUFS2 | 33 |
| GO:0032981 | BP | mitochondrial respiratory chain complex I assembly | 33/765 | 35/2446 | 43,12245308 | NDUFB6/NDUFA6/NDUFV2/NDUFB5/TMEM126B/NDUFB8/NDUFV1/NDUFB10/OXA1L/NDUFS8/NDUFS4/NDUFA2/NDUFS5/NDUFS2 | 33 |
| GO:0033108 | BP | mitochondrial respiratory chain complex assembly | 36/765 | 40/2446 | 88,86767114 | NDUFB6/NDUFA6/NDUFV2/NDUFB5/TMEM126B/NDUFB8/NDUFV1/NDUFB10/OXA1L/SCO1/NDUFS8/NDUFS4/NDUFA2/NDUFS5/N | 36 |
| GO:0061024 | BP | membrane organization | 121/765 | 224/2446 | 923,4742391 | ATP11C/APOOL/DOCK1/VDAC2/STX3/TAP1/GOSR2/STX12/VPS8/CHCHD3/STX18/REEP4/TMCC1/RFTN1/TRAM1/ABCA3/NUP93/RH | 121 |
| GO:0006812 | BP | cation transport | 75/765 | 122/2446 | 14562,96722 | MAGT1/SFXN3/PARL/CTNNB1/SLC33A1/ATP2C1/SLC30A6/WNK1/PHB2/SLC3A2/TMEM63B/COX4I1/NSF/SLC39A11/TMCO1/SNAP2 | 75 |
| GO:0007005 | BP | mitochondrion organization | 87/765 | 150/2446 | 21518,86147 | APOOL/NDUFB6/VDAC2/NDUFA6/PARL/TSC2/CHCHD3/NDUFV2/NDUFB5/TMEM126B/NDUFB8/PHB2/FAM162A/NDUFV1/NDUFB1 | 87 |
| GO:0009101 | BP | glycoprotein biosynthetic process | 44/765 | 59/2446 | 40703,22827 | MAGT1/ST6GALNAC3/DDOST/DPY19L1/CTNNB1/ST6GALNAC4/SOAT1/GALNT7/GLCE/MAN1A2/FAM20B/B3GAT3/RPN1/RPN2/ST6 | 44 |
| GO:1901137 | BP | carbohydrate derivative biosynthetic process | 72/765 | 118/2446 | 57656,63192 | MAGT1/ST6GALNAC3/DDOST/DPY19L1/PIGN/CTNNB1/CTPS1/ST6GALNAC4/SOAT1/PIGQ/ACSL1/PIGG/GALNT7/GLCE/DCAKD/MAN | 72 |
| GO:0015849 | BP | organic acid transport | 36/765 | 45/2446 | 99373,31572 | SFXN3/ACSL1/SLC3A2/SLC25A11/SLC16A3/ACSL4/PRAF2/SLC25A12/GLS/ACSL3/ITGB1/THBS1/SLC2A1/GJA1/ABCD3/ABCC1/ABCD1/ | 36 |
| GO:0009100 | BP | glycoprotein metabolic process | 49/765 | 70/2446 | 10718,45284 | MAGT1/ST6GALNAC3/NGLY1/DDOST/RNF5/DPY19L1/CTNNB1/ST6GALNAC4/SOAT1/GALNT7/GLCE/MAN1A2/FAM20B/B3GAT3/RP | 49 |
| GO:1903825 | BP | organic acid transmembrane transport | 29/765 | 34/2446 | 478420,8706 | SFXN3/ACSL1/SLC3A2/SLC25A11/SLC16A3/PRAF2/SLC25A12/ITGB1/THBS1/SLC2A1/ABCD3/ABCC1/ABCD1/SLC1A1/CPT1A/SLC25A1 | 29 |
| GO:1905039 | BP | carboxylic acid transmembrane transport | 29/765 | 34/2446 | 478420,8706 | SFXN3/ACSL1/SLC3A2/SLC25A11/SLC16A3/PRAF2/SLC25A12/ITGB1/THBS1/SLC2A1/ABCD3/ABCC1/ABCD1/SLC1A1/CPT1A/SLC25A1 | 29 |
| GO:0070085 | BP | glycosylation | 36/765 | 47/2446 | 749043,698 | MAGT1/ST6GALNAC3/DDOST/DPY19L1/ST6GALNAC4/GALNT7/MAN1A2/B3GAT3/RPN1/RPN2/ST6GAL1/MGAT1/STT3A/COG7/MG | 36 |
| GO:0045333 | BP | cellular respiration | 39/765 | 53/2446 | 957518,8435 | NDUFB6/NDUFA6/SUCLA2/NDUFV2/NDUFB5/NDUFB8/NDUFV1/COX4I1/NDUFB10/OXA1L/NDUFS8/UQCRQ/NDUFS4/NDUFA2/NDU | 39 |
| GO:0140352 | BP | export from cell | 119/765 | 239/2446 | 1185118,295 | SCAMP1/MAGT1/APOOL/RAB18/STX3/DDOST/EXOC6B/JAGN1/WLS/BRI3/SPARC/MGST1/PTPRB/MIA2/RIC1/METTL7A/RAB27A/RH | 119 |
| GO:0006486 | BP | protein glycosylation | 35/765 | 46/2446 | 1650621,963 | MAGT1/ST6GALNAC3/DDOST/DPY19L1/ST6GALNAC4/GALNT7/MAN1A2/B3GAT3/RPN1/RPN2/ST6GAL1/MGAT1/STT3A/COG7/MG | 35 |
| GO:0043413 | BP | macromolecule glycosylation | 35/765 | 46/2446 | 1650621,963 | MAGT1/ST6GALNAC3/DDOST/DPY19L1/ST6GALNAC4/GALNT7/MAN1A2/B3GAT3/RPN1/RPN2/ST6GAL1/MGAT1/STT3A/COG7/MG | 35 |
| GO:0098655 | BP | cation transmembrane transport | 60/765 | 98/2446 | 1942333,786 | MAGT1/SFXN3/PARL/SLC33A1/ATP2C1/SLC30A6/WNK1/PHB2/SLC3A2/TMEM63B/COX4I1/SLC39A11/TMCO1/LETM1/ATP1A1/APP/ | 60 |
| GO:0010876 | BP | lipid localization | 45/765 | 66/2446 | 2065016,411 | ATP11C/STX12/SOAT1/ACSL1/ABCA3/APOL2/ABCC4/ACSL4/PRAF2/ATP9A/SLC25A12/GLS/ACSL3/CDS2/ITGB1/THBS1/CLU/SLC2A1/ | 45 |
| GO:0006119 | BP | oxidative phosphorylation | 31/765 | 39/2446 | 2450621,208 | NDUFB6/NDUFA6/NDUFV2/NDUFB5/NDUFB8/NDUFV1/COX4I1/NDUFB10/NDUFS8/UQCRQ/NDUFS4/NDUFA2/NDUFS5/NDUFS2/N | 31 |
| GO:0046903 | BP | secretion | 120/765 | 244/2446 | 2459470,665 | SCAMP1/MAGT1/APOOL/RAB18/STX3/DDOST/EXOC6B/JAGN1/WLS/BRI3/SPARC/MGST1/PTPRB/MIA2/RIC1/METTL7A/RAB27A/RH | 120 |
| GO:0006120 | BP | mitochondrial electron transport, NADH to ubiquinone | 24/765 | 27/2446 | 2575744,266 | NDUFB6/NDUFA6/NDUFV2/NDUFB5/NDUFB8/NDUFV1/NDUFB10/NDUFS8/NDUFS4/NDUFA2/NDUFS5/NDUFS2/NDUFS6/NDUFS3/ | 24 |
| GO:0022904 | BP | respiratory electron transport chain | 32/765 | 41/2446 | 2799766,579 | NDUFB6/NDUFA6/NDUFV2/NDUFB5/NDUFB8/NDUFV1/COX4I1/NDUFB10/NDUFS8/UQCRQ/NDUFS4/NDUFA2/NDUFS5/NDUFS2/N | 32 |
| GO:0042773 | BP | ATP synthesis coupled electron transport | 28/765 | 34/2446 | 3440295,491 | NDUFB6/NDUFA6/NDUFV2/NDUFB5/NDUFB8/NDUFV1/COX4I1/NDUFB10/NDUFS8/UQCRQ/NDUFS4/NDUFA2/NDUFS5/NDUFS2/N | 28 |
| GO:0042775 | BP | mitochondrial ATP synthesis coupled electron transport | 28/765 | 34/2446 | 3440295,491 | NDUFB6/NDUFA6/NDUFV2/NDUFB5/NDUFB8/NDUFV1/COX4I1/NDUFB10/NDUFS8/UQCRQ/NDUFS4/NDUFA2/NDUFS5/NDUFS2/N | 28 |
| GO:0044255 | BP | cellular lipid metabolic process | 90/765 | 170/2446 | 3780209,776 | ST6GALNAC3/NCEH1/CEPT1/LPCAT4/ABHD16A/PON2/CLN6/PIGN/PNPLA6/ILVBL/ST6GALNAC4/SOAT1/ACBD5/ACADM/PIGQ/ERLIN | 90 |
| GO:0022900 | BP | electron transport chain | 35/765 | 47/2446 | 3846724,863 | NDUFB6/NDUFA6/NDUFV2/NDUFB5/NDUFB8/NDUFV1/COX4I1/NDUFB10/NDUFS8/UQCRQ/NDUFS4/NDUFA2/NDUFS5/NDUFS2/N | 35 |
| GO:0036503 | BP | ERAD pathway | 27/765 | 33/2446 | 8859069,762 | HM13/RNF5/ERLIN2/STUB1/ERLIN1/WFS1/DNAJB2/BCAP31/SEC61B/UBE2G2/CAV1/UBE4A/SYVN1/UBAC2/DNAJB14/STT3B/UBXN4 | 27 |
| GO:0032940 | BP | secretion by cell | 114/765 | 233/2446 | 9158850,054 | SCAMP1/MAGT1/APOOL/RAB18/STX3/DDOST/EXOC6B/JAGN1/WLS/BRI3/SPARC/MGST1/PTPRB/MIA2/RIC1/METTL7A/RAB27A/RH | 114 |
| GO:0034976 | BP | response to endoplasmic reticulum stress | 54/765 | 88/2446 | 10933645,31 | HM13/HYOU1/DDRGK1/RNF5/GOSR2/TPP1/SSR1/EIF2B5/ERLIN2/ERMP1/TBL2/STUB1/TMCO1/ERLIN1/WFS1/UFL1/VAPB/THBS1/C | 54 |
| GO:0006629 | BP | lipid metabolic process | 109/765 | 221/2446 | 1174956,819 | ST6GALNAC3/DHCR24/NCEH1/CEPT1/LPCAT4/ABHD16A/PON2/CLN6/PIGN/TPP1/PNPLA6/ILVBL/ST6GALNAC4/SOAT1/ACBD5/ACA | 109 |
| GO:0098660 | BP | inorganic ion transmembrane transport | 53/765 | 86/2446 | 1212055,17 | MAGT1/PARL/SLC33A1/ATP2C1/SLC30A6/WNK1/PHB2/SLC3A2/CLCN7/COX4I1/SLC25A11/SLC39A11/TMCO1/LETM1/ATP1A1/ITGB | 53 |
| GO:0015711 | BP | organic anion transport | 32/765 | 43/2446 | 19196799,68 | SFXN3/SLC33A1/SLC3A2/SLC25A11/SLC16A3/ABCC4/PRAF2/SLC25A12/GLS/SLC4A2/ITGB1/SLC2A1/GJA1/ABCC1/SLC1A1/SLC25A1/ | 32 |
| GO:0048284 | BP | organelle fusion | 36/765 | 51/2446 | 21814290,56 | STX3/TAP1/GOSR2/STX12/VPS8/CHCHD3/STX10/SNAP23/STX7/STX6/USO1/VAMP4/SEC22B/MFN2/SNAP29/TFRC/RAB13/STX17/R | 36 |
| GO:0006869 | BP | lipid transport | 40/765 | 60/2446 | 39796789,44 | ATP11C/STX12/SOAT1/ACSL1/ABCA3/APOL2/ABCC4/ACSL4/PRAF2/ATP9A/SLC25A12/GLS/ACSL3/ITGB1/THBS1/CLU/SLC2A1/GJA1/ | 40 |
| GO:0030433 | BP | ubiquitin-dependent ERAD pathway | 24/765 | 29/2446 | 39806217,91 | RNF5/ERLIN2/STUB1/ERLIN1/WFS1/DNAJB2/BCAP31/SEC61B/UBE2G2/CAV1/UBE4A/SYVN1/DNAJB14/STT3B/UBXN4/CCDC47/FAF2 | 24 |
| GO:1901135 | BP | carbohydrate derivative metabolic process | 96/765 | 193/2446 | 59986032,61 | MAGT1/ST6GALNAC3/NGLY1/DDOST/CLN6/RNF5/DPY19L1/PIGN/CTNNB1/SUCLA2/CTPS1/ST6GALNAC4/SOAT1/PIGQ/ACSL1/PIGG/ | 96 |
| GO:0046942 | BP | carboxylic acid transport | 26/765 | 33/2446 | 64685366,8 | SFXN3/SLC3A2/SLC25A11/SLC16A3/ABCC4/PRAF2/SLC25A12/GLS/ITGB1/GJA1/ABCC1/SLC1A1/SLC25A1/SLC16A1/SLC1A5/SLC27A4, | 26 |
| GO:0010256 | BP | endomembrane system organization | 74/765 | 139/2446 | 7852177,251 | RAB18/COG4/JAGN1/ARL1/COG2/STX18/REEP4/TMCC1/TRAM1/RAB27A/NUP93/SMPD4/CTDNEP1/EMC10/STK25/RAB29/ZW10/E | 74 |
| GO:0061025 | BP | membrane fusion | 36/765 | 53/2446 | 9768050,573 | STX3/TAP1/GOSR2/STX12/VPS8/STX18/NSF/STX10/SNAP23/STX7/STX6/USO1/VAMP4/SEC22B/SNAP29/TIE1/RAB13/NAPA/STX17/ | 36 |
| GO:0007029 | BP | endoplasmic reticulum organization | 24/765 | 30/2446 | 129158430,9 | RAB18/JAGN1/STX18/REEP4/TMCC1/TRAM1/SMPD4/EMC10/EMC8/VAPB/TMEM33/RAB10/LMAN2/BNIP1/EMC2/TRAM2/TMED2/ | 24 |
| GO:0006839 | BP | mitochondrial transport | 46/765 | 75/2446 | 141769652,2 | VDAC2/SFXN3/PARL/RHOT1/OXA1L/HAX1/TIMM44/HIP1R/MTX2/SLC25A12/MFN2/LETM1/TOMM40/CNP/SLC9A1/YWHAQ/CPT1A/ | 46 |
| GO:0098662 | BP | inorganic cation transmembrane transport | 47/765 | 79/2446 | 372779266,2 | MAGT1/PARL/SLC33A1/ATP2C1/SLC30A6/WNK1/PHB2/SLC3A2/COX4I1/SLC39A11/TMCO1/LETM1/ATP1A1/ITGB1/COX6C/ATP2A2/ | 47 |
| GO:0048278 | BP | vesicle docking | 24/765 | 31/2446 | 383411050,5 | STX3/EXOC6B/STX12/NSF/STX10/STX7/STX6/USO1/RAB3D/RALB/RAB13/STX17/RAB8A/RAB10/STX4/STX5/VAMP3/EXOC6/SCFD2/S | 24 |
| GO:0006865 | BP | amino acid transport | 20/765 | 24/2446 | 495815753,8 | SFXN3/SLC3A2/PRAF2/SLC25A12/GLS/ITGB1/GJA1/SLC1A1/SLC1A5/SFXN4/LRRC8D/LRRC8A/SLC25A29/LRRC8C/SFXN2/SLC25A22/S | 20 |
| GO:0003333 | BP | amino acid transmembrane transport | 17/765 | 19/2446 | 540021749,5 | SFXN3/SLC3A2/PRAF2/SLC25A12/ITGB1/SLC1A1/SLC1A5/LRRC8D/LRRC8A/SLC25A29/LRRC8C/SFXN2/SLC25A22/SFXN1/SLC38A10/S | 17 |
| GO:1902600 | BP | proton transmembrane transport | 18/765 | 21/2446 | 90771713,13 | SLC33A1/ATP2C1/PHB2/COX4I1/ATP1A1/COX6C/ATP2A2/SLC9A1/ATP6V1A/CLCN3/SLC25A3/NNT/TCIRG1/TMCO3/SLC30A5/DNAJC | 18 |
| GO:0006887 | BP | exocytosis | 88/765 | 182/2446 | 1120379074 | SCAMP1/MAGT1/APOOL/RAB18/STX3/DDOST/EXOC6B/JAGN1/BRI3/SPARC/MGST1/PTPRB/METTL7A/RAB27A/NSF/SNAP23/QSOX1 | 88 |
| GO:0015980 | BP | energy derivation by oxidation of organic compounds | 41/765 | 68/2446 | 1347773247 | NDUFB6/NDUFA6/SUCLA2/PHKA1/NDUFV2/NDUFB5/NDUFB8/NDUFV1/COX4I1/NDUFB10/OXA1L/NDUFS8/UQCRQ/NDUFS4/NDUF | 41 |
| GO:0048193 | BP | Golgi vesicle transport | 74/765 | 147/2446 | 1347773247 | SCAMP1/HYOU1/COG4/EXOC6B/GOSR2/MON2/NBAS/COG8/COG2/SEC22A/STX18/MIA2/NSF/SNAP23/SCAMP3/ZW10/STX6/USO1, | 74 |
| GO:0010232 | BP | vascular transport | 12/765 | 12/2446 | 204640061 | ABCC4/TFRC/SLC2A1/GJA1/ATP2B4/ABCC1/SLC1A1/SLC16A1/SLC1A5/SLC27A4/SLC27A1/SLC29A1 | 12 |
| GO:0007006 | BP | mitochondrial membrane organization | 28/765 | 41/2446 | 242173023,3 | APOOL/VDAC2/CHCHD3/RHOT1/OXA1L/CHCHD6/HIP1R/MTX2/MFN2/LETM1/CNP/TMEM11/SLC9A1/YWHAQ/ROMO1/TIMM10/BA | 28 |
| GO:0090174 | BP | organelle membrane fusion | 28/765 | 41/2446 | 242173023,3 | STX3/TAP1/GOSR2/STX12/VPS8/STX10/SNAP23/STX7/STX6/USO1/VAMP4/SEC22B/SNAP29/RAB13/STX17/RAB8A/STX4/BNIP1/STX | 28 |
| GO:0006906 | BP | vesicle fusion | 27/765 | 39/2446 | 2463883124 | STX3/TAP1/GOSR2/STX12/VPS8/STX10/SNAP23/STX7/STX6/USO1/VAMP4/SEC22B/SNAP29/RAB13/STX17/RAB8A/STX4/STX5/VAM | 27 |
| GO:0050801 | BP | ion homeostasis | 52/765 | 95/2446 | 2686472038 | TM9SF4/CLN6/ATP2C1/WNK1/SCO1/TMCO1/WFS1/GLS/LETM1/VAPB/TFRC/GNAI2/SLC4A2/ATP1A1/APP/HMOX1/ATP2A2/GJA1/S | 52 |
| GO:0030001 | BP | metal ion transport | 38/765 | 64/2446 | 5509878870 | MAGT1/PARL/CTNNB1/ATP2C1/SLC30A6/PHB2/SLC3A2/SLC39A11/TMCO1/ZMPSTE24/WFS1/LETM1/GNAI2/ATP1A1/ATP2A2/GJA1 | 38 |
| GO:0046471 | BP | phosphatidylglycerol metabolic process | 13/765 | 14/2446 | 5785407424 | LPCAT4/PHB2/ABCA3/HADHA/CDS2/HADHB/SLC27A1/LCLAT1/LPCAT1/LPGAT1/TAMM41/DNAJC19/STOML2 | 13 |
| GO:0150104 | BP | transport across blood-brain barrier | 11/765 | 11/2446 | 5966218253 | ABCC4/TFRC/SLC2A1/ATP2B4/ABCC1/SLC1A1/SLC16A1/SLC1A5/SLC27A4/SLC27A1/SLC29A1 | 11 |
| GO:0055065 | BP | metal ion homeostasis | 43/765 | 76/2446 | 6926524661 | ATP2C1/SCO1/TMCO1/WFS1/LETM1/VAPB/TFRC/GNAI2/ATP1A1/APP/HMOX1/ATP2A2/GJA1/SLC9A1/ATP2B1/ATP2B4/F2R/HMOX | 43 |
| GO:0055080 | BP | cation homeostasis | 48/765 | 89/2446 | 1,10344E+14 | TM9SF4/CLN6/ATP2C1/SCO1/TMCO1/WFS1/LETM1/VAPB/TFRC/GNAI2/SLC4A2/ATP1A1/APP/HMOX1/ATP2A2/GJA1/SLC9A1/ATP2 | 48 |
| GO:0098771 | BP | inorganic ion homeostasis | 48/765 | 89/2446 | 1,10344E+14 | TM9SF4/CLN6/ATP2C1/SCO1/TMCO1/WFS1/LETM1/VAPB/TFRC/GNAI2/SLC4A2/ATP1A1/APP/HMOX1/ATP2A2/GJA1/SLC9A1/ATP2 | 48 |
| GO:1990542 | BP | mitochondrial transmembrane transport | 23/765 | 33/2446 | 1,17913E+13 | SFXN3/TIMM44/SLC25A12/LETM1/TOMM40/CPT1A/SLC25A1/ROMO1/TIMM50/SFXN4/SLC25A29/MCU/DNAJC19/DNAJC30/SFXN2 | 23 |
| GO:0016192 | BP | vesicle-mediated transport | 186/765 | 466/2446 | 1,40855E+14 | SCAMP1/MAGT1/APOOL/HYOU1/RAB18/DOCK1/SCARF2/COG4/TM9SF4/STX3/DDOST/EXOC6B/TAP1/EVI5/GOSR2/MON2/TSC2/JA | 186 |
| GO:0035966 | BP | response to topologically incorrect protein | 41/765 | 73/2446 | 1,40855E+14 | HYOU1/DDRGK1/RNF5/GOSR2/TPP1/SSR1/ERMP1/TBL2/STUB1/RNF126/WFS1/UFL1/MFN2/VAPB/HSPB1/THBS1/HSP90AB1/CLU/P | 41 |
| GO:0072348 | BP | sulfur compound transport | 12/765 | 13/2446 | 1,5638E+14 | SLC33A1/SLC25A11/GJA1/ABCC1/ABCD1/SLC1A1/SLC27A1/LRRC8D/LRRC8A/SLC35B2/LRRC8C/SLC25A19 | 12 |
| GO:0046034 | BP | ATP metabolic process | 45/765 | 83/2446 | 1,73602E+14 | NDUFB6/NDUFA6/NDUFV2/NDUFB5/NDUFB8/AAAS/NDUFV1/COX4I1/NDUFB10/NUP93/NDUFS8/UQCRQ/NDUFS4/NDUFA2/NDUF | 45 |
| GO:1901698 | BP | response to nitrogen compound | 89/765 | 196/2446 | 2,08517E+14 | HM13/BSG/RNF5/CTNNB1/TSC2/EIF2B5/ERLIN2/VKORC1/SPARC/SLC3A2/MGST1/RFTN1/SOS1/STUB1/ABCC4/NDUFS4/USO1/ERLIN | 89 |
| GO:0006487 | BP | protein N-linked glycosylation | 17/765 | 22/2446 | 2,17293E+13 | MAGT1/DDOST/RPN1/RPN2/ST6GAL1/MGAT1/STT3A/MGAT2/STT3B/EXT2/ALG1/FUT8/ALG2/TMEM165/B4GALT7/UBE2J1/ALG5 | 17 |
| GO:0006875 | BP | cellular metal ion homeostasis | 40/765 | 72/2446 | 2,46468E+14 | ATP2C1/SCO1/TMCO1/WFS1/LETM1/VAPB/TFRC/ATP1A1/APP/HMOX1/ATP2A2/GJA1/SLC9A1/ATP2B1/ATP2B4/F2R/HMOX2/CDH5 | 40 |
| GO:0015908 | BP | fatty acid transport | 19/765 | 26/2446 | 2,48624E+14 | ACSL1/ACSL4/PRAF2/SLC25A12/GLS/ACSL3/ITGB1/THBS1/SLC2A1/GJA1/ABCD3/ABCD1/SLC1A1/CPT1A/SLC27A3/SLC27A4/SLC27A1 | 19 |
| GO:0007007 | BP | inner mitochondrial membrane organization | 13/765 | 15/2446 | 2,557E+13 | APOOL/CHCHD3/CHCHD6/MTX2/LETM1/TMEM11/ROMO1/TIMM10/MTX1/IMMT/DNAJC11/NDUFA13/AFG3L2 | 13 |
| GO:0008610 | BP | lipid biosynthetic process | 65/765 | 134/2446 | 2,557E+13 | ST6GALNAC3/DHCR24/CEPT1/LPCAT4/PIGN/ST6GALNAC4/PIGQ/ERLIN2/ACSL1/PIGG/PTDSS2/FADS3/ABCA3/SMPD4/CTDNEP1/ALD | 65 |
| GO:0006873 | BP | cellular ion homeostasis | 45/765 | 85/2446 | 3,65834E+14 | TM9SF4/CLN6/ATP2C1/SCO1/TMCO1/WFS1/LETM1/VAPB/TFRC/SLC4A2/ATP1A1/APP/HMOX1/ATP2A2/GJA1/SLC9A1/ATP2B1/ATP | 45 |
| GO:0030003 | BP | cellular cation homeostasis | 45/765 | 85/2446 | 3,65834E+14 | TM9SF4/CLN6/ATP2C1/SCO1/TMCO1/WFS1/LETM1/VAPB/TFRC/SLC4A2/ATP1A1/APP/HMOX1/ATP2A2/GJA1/SLC9A1/ATP2B1/ATP | 45 |
| GO:0003013 | BP | circulatory system process | 39/765 | 71/2446 | 4,30453E+14 | ECE1/ACVRL1/ABCC4/ZMPSTE24/CYB5R3/TFRC/GNAI2/ATP1A1/HMOX1/SLC2A1/JUP/ATP2A2/GJA1/SLC9A1/ATP2B1/FLNA/ATP2B4 | 39 |
| GO:0048878 | BP | chemical homeostasis | 67/765 | 141/2446 | 4,32756E+14 | TM9SF4/CLN6/JAGN1/SOAT1/ATP2C1/WNK1/ABCA3/SCO1/TMCO1/WFS1/GLS/LETM1/VAPB/TFRC/GNAI2/SLC4A2/ATP1A1/APP/H | 67 |
| GO:1905897 | BP | regulation of response to endoplasmic reticulum stress | 15/765 | 19/2446 | 4,379E+14 | DDRGK1/WFS1/UFL1/CLU/PTPN2/PTPN1/ATF6/BCAP31/TMEM33/CAV1/BAX/TMED2/DAB2IP/SYVN1/EIF2AK3 | 15 |
| GO:0007030 | BP | Golgi organization | 29/765 | 48/2446 | 4,42112E+14 | COG4/ARL1/COG2/STX18/STK25/RAB29/ZW10/STX6/USO1/VAMP4/MAP2K2/TMED10/STX17/RAB2A/COG7/LMAN2/STX5/TMED2/ | 29 |
| GO:0015748 | BP | organophosphate ester transport | 16/765 | 21/2446 | 4,93448E+14 | ATP11C/SLC33A1/ABCA3/ABCC4/ATP9A/GJA1/ABCC1/ABCD1/ANO6/SLC35B2/OSBPL8/OSBPL5/SLC25A51/SLC25A19/EPG5/TMEM3 | 16 |
| GO:0042147 | BP | retrograde transport, endosome to Golgi | 21/765 | 31/2446 | 5,08691E+14 | EVI5/GOSR2/ARL1/RIC1/STX10/RNF126/RAB29/STX6/RAB6A/STX5/VAMP3/TBC1D10B/TMEM87A/GBF1/TBC1D5/VTI1A/TMEM87B/ | 21 |
| GO:0051205 | BP | protein insertion into membrane | 17/765 | 23/2446 | 5,21519E+14 | TRAM1/OXA1L/EMC10/EMC8/YWHAQ/ROMO1/TIMM10/BAX/EMC2/TRAM2/EMC1/BAD/EMC7/TOMM22/EMC3/NDUFA13/BCS1L | 17 |
| GO:0016485 | BP | protein processing | 18/765 | 25/2446 | 5,32068E+14 | HM13/SPCS2/ECE1/SEC11A/ADAM10/ZMPSTE24/BAG2/THBS1/MMP14/SPCS3/ASPH/BAD/YME1L1/TBC1D10A/CTSZ/STOML2/AFG | 18 |
| GO:1903522 | BP | regulation of blood circulation | 18/765 | 25/2446 | 5,32068E+14 | ECE1/ZMPSTE24/ATP1A1/JUP/ATP2A2/GJA1/SLC9A1/ATP2B1/FLNA/ATP2B4/F2R/GJA5/SLC1A1/CAV1/ASPH/STIM1/DOCK4/DOCK5 | 18 |
| GO:0140029 | BP | exocytic process | 19/765 | 27/2446 | 5,32068E+14 | STX3/EXOC6B/SNAP23/SNX4/RAB3D/SNAP29/RALB/RAB13/NAPA/RAB8A/RAB10/STX4/VAMP3/EXOC6/SCFD2/SCFD1/EXOC4/VPS4 | 19 |
| GO:0018196 | BP | peptidyl-asparagine modification | 12/765 | 14/2446 | 6,50864E+14 | MAGT1/DDOST/RPN1/RPN2/ST6GAL1/MGAT1/STT3A/MGAT2/STT3B/FUT8/UBE2J1/ALG5 | 12 |
| GO:0018279 | BP | protein N-linked glycosylation via asparagine | 12/765 | 14/2446 | 6,50864E+14 | MAGT1/DDOST/RPN1/RPN2/ST6GAL1/MGAT1/STT3A/MGAT2/STT3B/FUT8/UBE2J1/ALG5 | 12 |
| GO:0072503 | BP | cellular divalent inorganic cation homeostasis | 33/765 | 58/2446 | 6,52692E+14 | ATP2C1/TMCO1/WFS1/LETM1/VAPB/APP/ATP2A2/GJA1/ATP2B1/ATP2B4/F2R/CDH5/SLC1A1/BCAP31/CXCR4/GNB1/CAV1/BAX/TC | 33 |
| GO:0072507 | BP | divalent inorganic cation homeostasis | 33/765 | 58/2446 | 6,52692E+14 | ATP2C1/TMCO1/WFS1/LETM1/VAPB/APP/ATP2A2/GJA1/ATP2B1/ATP2B4/F2R/CDH5/SLC1A1/BCAP31/CXCR4/GNB1/CAV1/BAX/TC | 33 |
| GO:0006091 | BP | generation of precursor metabolites and energy | 57/765 | 117/2446 | 6,77983E+13 | NDUFB6/NDUFA6/SUCLA2/PHKA1/NDUFV2/NDUFB5/NDUFB8/PHB2/AAAS/NDUFV1/COX4I1/NDUFB10/NUP93/OXA1L/NDUFS8/U | 57 |
| GO:0006631 | BP | fatty acid metabolic process | 32/765 | 56/2446 | 7,54071E+14 | PON2/ILVBL/ACBD5/ACADM/ERLIN2/ACSL1/FADS3/HADHA/ALDH3A2/FADS1/ACSL4/ERLIN1/SGPL1/ACSL3/FADS2/ABCD3/ABCD1/C | 32 |
| GO:0006643 | BP | membrane lipid metabolic process | 23/765 | 36/2446 | 8,25179E+14 | ST6GALNAC3/CLN6/PIGN/ST6GALNAC4/PIGQ/PIGG/FADS3/SMPD4/ALDH3A2/SPTLC1/VAPB/SGPL1/BAX/PDXDC1/HACD2/CERS6/SC | 23 |
| GO:0032527 | BP | protein exit from endoplasmic reticulum | 13/765 | 16/2446 | 8,41211E+14 | HM13/TM9SF4/BCAP31/SEC61B/UBE2G2/SYVN1/UBAC2/FAF2/DERL2/PREB/TMEM30A/UBE2J1/AUP1 | 13 |
| GO:0042157 | BP | lipoprotein metabolic process | 13/765 | 16/2446 | 8,41211E+14 | PIGN/PIGQ/PIGG/ZDHHC3/APOL2/ATM/RABL3/ZDHHC24/ZDHHC13/ABHD12/PIGK/PIGS/ZDHHC5 | 13 |
| GO:0010243 | BP | response to organonitrogen compound | 81/765 | 181/2446 | 8,42217E+13 | HM13/BSG/RNF5/CTNNB1/TSC2/EIF2B5/ERLIN2/VKORC1/SPARC/MGST1/SOS1/STUB1/ABCC4/NDUFS4/USO1/ERLIN1/WFS1/UFL1/ | 81 |
| GO:0098742 | BP | cell-cell adhesion via plasma-membrane adhesion molecules | 15/765 | 20/2446 | 1,08865E+14 | ROBO1/BSG/SCARF2/ATP2C1/ARVCF/PTPRM/WNK1/PTPRF/ITGB1/PECAM1/CDH5/ACVR1/PCDH1/ROBO4/ESAM | 15 |
| GO:0046467 | BP | membrane lipid biosynthetic process | 18/765 | 26/2446 | 1,12444E+14 | ST6GALNAC3/PIGN/ST6GALNAC4/PIGQ/PIGG/SMPD4/ALDH3A2/SPTLC1/VAPB/SGPL1/HACD2/CERS6/SCCPDH/PIGK/CERS2/PIGS/EL | 18 |
| GO:0015718 | BP | monocarboxylic acid transport | 17/765 | 24/2446 | 1,13612E+14 | SLC16A3/ABCC4/PRAF2/SLC25A12/GLS/ITGB1/GJA1/SLC1A1/SLC16A1/SLC1A5/SLC27A4/SLC27A1/LRRC8D/LRRC8A/LRRC8C/SLC25A | 17 |
| GO:0071806 | BP | protein transmembrane transport | 17/765 | 24/2446 | 1,13612E+14 | PEX1/PEX5/TRAM1/TIMM44/TOMM40/CLU/TMED10/SEC61G/SEC61B/ROMO1/PEX6/TRAM2/TIMM50/DNAJC19/TIMM21/TOMM2 | 17 |
| GO:0046486 | BP | glycerolipid metabolic process | 46/765 | 91/2446 | 1,14542E+14 | CEPT1/LPCAT4/ABHD16A/PIGN/PNPLA6/PIGQ/ACSL1/PIGG/PTDSS2/PHB2/ABCA3/HADHA/SMPD4/CTDNEP1/ACSL4/ACSL3/CDS2/R | 46 |
| GO:0006892 | BP | post-Golgi vesicle-mediated transport | 24/765 | 39/2446 | 1,27747E+14 | SCAMP1/EXOC6B/GOSR2/MON2/NSF/SNAP23/SCAMP3/ACSL3/RAB10/RAB14/STX4/VAMP3/EXOC8/EXOC6/SCFD1/GBF1/EXOC4/V | 24 |
| GO:0055082 | BP | cellular chemical homeostasis | 51/765 | 104/2446 | 1,29324E+14 | TM9SF4/CLN6/JAGN1/ATP2C1/SCO1/TMCO1/WFS1/LETM1/VAPB/TFRC/SLC4A2/ATP1A1/APP/HMOX1/ATP2A2/GJA1/HK1/SLC9A1/ | 51 |
| GO:0016050 | BP | vesicle organization | 52/765 | 107/2446 | 1,46948E+14 | STX3/TAP1/GOSR2/STX12/VPS8/TMCC1/RAB27A/NSF/STX10/SNAP23/RAB29/STX7/STX6/USO1/VAMP4/SEC22B/VAPB/SNAP29/F2R | 52 |
| GO:0015807 | BP | L-amino acid transport | 11/765 | 13/2446 | 1,60756E+14 | SLC3A2/PRAF2/SLC25A12/ITGB1/SLC1A1/SLC1A5/SLC25A29/SLC25A22/SFXN1/SLC25A13/SLC25A15 | 11 |
| GO:0006874 | BP | cellular calcium ion homeostasis | 29/765 | 51/2446 | 1,75044E+14 | ATP2C1/TMCO1/WFS1/LETM1/VAPB/APP/ATP2A2/GJA1/ATP2B1/ATP2B4/F2R/CDH5/BCAP31/CXCR4/GNB1/CAV1/BAX/TCIRG1/PK | 29 |
| GO:0055074 | BP | calcium ion homeostasis | 29/765 | 51/2446 | 1,75044E+14 | ATP2C1/TMCO1/WFS1/LETM1/VAPB/APP/ATP2A2/GJA1/ATP2B1/ATP2B4/F2R/CDH5/BCAP31/CXCR4/GNB1/CAV1/BAX/TCIRG1/PK | 29 |
| GO:0045055 | BP | regulated exocytosis | 66/765 | 145/2446 | 2,13423E+14 | SCAMP1/MAGT1/APOOL/RAB18/STX3/DDOST/BRI3/SPARC/MGST1/PTPRB/METTL7A/RAB27A/SNAP23/QSOX1/ADAM10/ABCC4/SN | 66 |
| GO:0006835 | BP | dicarboxylic acid transport | 13/765 | 17/2446 | 2,26947E+14 | SLC25A11/PRAF2/SLC25A12/GLS/ITGB1/GJA1/SLC1A1/SLC1A5/LRRC8D/LRRC8A/LRRC8C/SLC25A22/SLC25A13 | 13 |
| GO:0006984 | BP | ER-nucleus signaling pathway | 13/765 | 17/2446 | 2,26947E+14 | ERLIN2/TMCO1/ERLIN1/WFS1/ATP2A2/PTPN2/PTPN1/ATF6/TMEM33/TMED2/CCDC47/EIF2AK3/AMFR | 13 |
| GO:0098739 | BP | import across plasma membrane | 13/765 | 17/2446 | 2,26947E+14 | ACSL1/WNK1/SLC3A2/ATP1A1/ITGB1/THBS1/SLC2A1/SLC9A1/ATP2B4/SLC1A1/SLC1A5/SLC27A1/SLC30A1 | 13 |
| GO:1902001 | BP | fatty acid transmembrane transport | 13/765 | 17/2446 | 2,26947E+14 | ACSL1/PRAF2/SLC25A12/ITGB1/THBS1/SLC2A1/ABCD3/ABCD1/SLC1A1/CPT1A/SLC27A1/SLC25A22/SLC25A13 | 13 |
| GO:0006891 | BP | intra-Golgi vesicle-mediated transport | 17/765 | 25/2446 | 2,31311E+14 | GOSR2/COG8/COG2/NSF/RAB6A/NAPA/CUX1/GOLGA5/COG1/VTI1A/COG3/NAPG/RAB33B/COPG2/VTI1B/COG5/COG6 | 17 |

| | | | | | | | |
|---|---|---|---|---|---|---|---|
| GO:0003018 | BP | vascular process in circulatory system | 23/765 | 38/2446 | 2,38061E+14 | ECE1/ABCC4/TFRC/SLC2A1/GJA1/ATP2B1/ATP2B4/F2R/NOS3/CDH5/ABCC1/GJA5/SLC1A1/SLC16A1/TEK/CAV1/SLC1A5/SLC27A4/SL | 23 |
| GO:0015698 | BP | inorganic anion transport | 14/765 | 19/2446 | 2,38061E+14 | VDAC2/CLCN7/SLC25A11/SLC4A2/SLC1A1/CLCN3/SLC25A3/ANO6/LRRC8D/LRRC8A/LRRC8C/CLCC1/TTYH3/ANO10 | 14 |
| GO:0035967 | BP | cellular response to topologically incorrect protein | 32/765 | 59/2446 | 2,49382E+14 | HYOU1/DDRGK1/RNF5/GOSR2/TPP1/SSR1/ERMP1/TBL2/STUB1/RNF126/WFS1/UFL1/VAPB/PTPN2/PTPN1/ATF6/TMEM33/BAX/TM | 32 |
| GO:0051049 | BP | regulation of transport | 106/765 | 256/2446 | 2,64481E+13 | TM9SF4/PARL/CTNNB1/TSC2/JAGN1/STX18/ACSL1/WLS/WNK1/PHB2/AAAS/ABCA3/RIC1/RAB27A/RHOT1/NSF/HAX1/RAB29/ACSL4 | 106 |
| GO:0006986 | BP | response to unfolded protein | 34/765 | 64/2446 | 2,66663E+14 | HYOU1/DDRGK1/GOSR2/TPP1/SSR1/ERMP1/TBL2/STUB1/WFS1/UFL1/MFN2/VAPB/HSPB1/THBS1/HSP90AB1/PTPN2/PTPN1/ATF6/ | 34 |
| GO:1900101 | BP | regulation of endoplasmic reticulum unfolded protein response | 9/765 | 10/2446 | 2,74079E+14 | DDRGK1/WFS1/UFL1/PTPN2/PTPN1/ATF6/TMEM33/BAX/DAB2IP | 9 |
| GO:0032879 | BP | regulation of localization | 173/765 | 451/2446 | 3,24883E+14 | ROBO1/DOCK1/DDRGK1/TM9SF4/STX3/DNAJB6/PARL/CTNNB1/TSC2/JAGN1/FRMD5/STX18/ACSL1/PTPRM/RRAS2/WLS/WNK1/SPA | 173 |
| GO:0006816 | BP | calcium ion transport | 30/765 | 55/2446 | 3,35791E+14 | PARL/CTNNB1/ATP2C1/PHB2/SLC3A2/TMCO1/ZMPSTE24/WFS1/LETM1/GNAI2/ATP2A2/GJA1/SLC9A1/ATP2B1/ATP2B4/F2R/PHB/C | 30 |
| GO:0051046 | BP | regulation of secretion | 41/765 | 82/2446 | 3,35791E+14 | JAGN1/WLS/RIC1/RAB27A/RHOT1/NSF/ACSL4/HIP1R/SNX4/ACSL3/RAB3D/GNAI2/HMOX1/SLC2A1/RALA/ATP2A2/GJA1/RAB5A/F2R | 41 |
| GO:0030166 | BP | proteoglycan biosynthetic process | 10/765 | 12/2446 | 3,85661E+14 | CTNNB1/GLCE/FAM20B/B3GAT3/BGN/CHST14/CSGALNACT1/CANT1/EXT2/B4GALT7 | 10 |
| GO:0030970 | BP | retrograde protein transport, ER to cytosol | 10/765 | 12/2446 | 3,85661E+14 | HM13/BCAP31/SEC61B/UBE2G2/SYVN1/UBAC2/FAF2/DERL2/UBE2J1/AUP1 | 10 |
| GO:1902475 | BP | L-alpha-amino acid transmembrane transport | 10/765 | 12/2446 | 3,85661E+14 | SLC3A2/PRAF2/SLC25A12/ITGB1/SLC1A1/SLC1A5/SLC25A29/SLC25A22/SLC25A13/SLC25A15 | 10 |
| GO:1903513 | BP | endoplasmic reticulum to cytosol transport | 10/765 | 12/2446 | 3,85661E+14 | HM13/BCAP31/SEC61B/UBE2G2/SYVN1/UBAC2/FAF2/DERL2/UBE2J1/AUP1 | 10 |
| GO:0032787 | BP | monocarboxylic acid metabolic process | 49/765 | 103/2446 | 3,91474E+12 | BSG/PON2/ILVBL/ACBD5/ACADM/ERLIN2/ACSL1/FADS3/AAAS/HADHA/NUP93/ALDH3A2/SLC16A3/FADS1/ACSL4/ERLIN1/SLC25A12 | 49 |
| GO:0030968 | BP | endoplasmic reticulum unfolded protein response | 26/765 | 46/2446 | 3,95227E+14 | HYOU1/DDRGK1/GOSR2/TPP1/SSR1/ERMP1/TBL2/STUB1/WFS1/UFL1/VAPB/PTPN2/PTPN1/ATF6/TMEM33/BAX/TMED2/DAB2IP/SY | 26 |
| GO:0051604 | BP | protein maturation | 22/765 | 37/2446 | 4,34695E+14 | HM13/SPCS2/ECE1/SEC11A/STUB1/ADAM10/ZMPSTE24/WFS1/BAG2/THBS1/MMP14/SPCS3/GALNT2/ASPH/BAD/YME1L1/LMF2/TE | 22 |
| GO:0008217 | BP | regulation of blood pressure | 11/765 | 14/2446 | 4,62645E+14 | ECE1/ACVRL1/ATP1A1/HMOX1/GJA1/ATP2B1/F2R/NOS3/GJA5/EXT2/CTSZ | 11 |
| GO:0065002 | BP | intracellular protein transmembrane transport | 14/765 | 20/2446 | 5,09579E+14 | PEX1/PEX5/TRAM1/TIMM44/TOMM40/SEC61G/SEC61B/ROMO1/PEX6/TRAM2/TIMM50/DNAJC19/TIMM21/SEC63 | 14 |
| GO:0000041 | BP | transition metal ion transport | 13/765 | 18/2446 | 5,11009E+13 | ATP2C1/SLC30A6/SLC39A11/TFRC/ATP6V1A/TCIRG1/STEAP3/SLC30A7/SLC30A5/SLC39A7/TMEM165/ATP13A1/SLC30A1 | 13 |
| GO:0140056 | BP | organelle localization by membrane tethering | 27/765 | 49/2446 | 5,11009E+13 | STX3/EXOC6B/STX12/NSF/STX10/STX7/STX6/USO1/VAPB/RAB3D/SNAP29/RALB/RAB13/STX17/RAB8A/RAB10/TUBA4A/STX4/STX5/ | 27 |
| GO:1903530 | BP | regulation of secretion by cell | 39/765 | 79/2446 | 6,04817E+14 | JAGN1/WLS/RIC1/RAB27A/RHOT1/NSF/ACSL4/SNX4/RAB3D/GNAI2/HMOX1/SLC2A1/RALA/ATP2A2/GJA1/RAB5A/F2R/NOTCH1/IDH | 39 |
| GO:0006888 | BP | endoplasmic reticulum to Golgi vesicle-mediated transport | 42/765 | 87/2446 | 6,73633E+14 | HYOU1/COG4/GOSR2/COG8/COG2/SEC22A/STX18/MIA2/NSF/ZW10/USO1/SEC22B/VAPB/TMED10/BCAP31/NAPA/STX17/COG7/TM | 42 |
| GO:0016042 | BP | lipid catabolic process | 22/765 | 38/2446 | 6,8814E+14 | NCEH1/ABHD16A/PNPLA6/ILVBL/ACBD5/ACADM/HADHA/SMPD4/ALDH3A2/SGPL1/ABCD3/ABCD1/CPT1A/HADHB/CYP27A1/PLD1/ | 22 |
| GO:0048771 | BP | tissue remodeling | 18/765 | 29/2446 | 6,92519E+14 | TPP1/ACVRL1/RAB3D/TFRC/GJA1/FLNA/NF1/BGN/F2R/EPHA2/NOS3/TIE1/FLT4/MMP14/JAG1/CAV1/BAX/TCIRG1 | 18 |
| GO:0043269 | BP | regulation of ion transport | 73/765 | 171/2446 | 8,83884E+14 | TM9SF4/PARL/CTNNB1/JAGN1/ACSL1/WLS/WNK1/PHB2/ABCA3/HAX1/RAB29/ACSL4/ZMPSTE24/WFS1/B3GAT3/SNX4/ACSL3/GNA | 73 |
| GO:0002443 | BP | leukocyte mediated immunity | 56/765 | 125/2446 | 9,23083E+14 | SCAMP1/MAGT1/RAB18/DDOST/JAGN1/BRI3/MGST1/PTPRB/RFTN1/MLH1/METTL7A/RAB27A/SNAP23/QSOX1/ADAM10/STX7/SNX | 56 |
| GO:0006650 | BP | glycerophospholipid metabolic process | 40/765 | 83/2446 | 9,23083E+14 | CEPT1/LPCAT4/ABHD16A/PIGN/PNPLA6/PIGQ/PIGG/PTDSS2/PHB2/ABCA3/HADHA/SMPD4/ACSL3/CDS2/RAB5A/PI4KA/PTDSS1/HAI | 40 |
| GO:0006641 | BP | triglyceride metabolic process | 9/765 | 11/2446 | 9,23083E+14 | ACSL1/CTDNEP1/ACSL4/CPT1A/CAV1/SLC27A1/LPGAT1/PNPLA2/MBOAT7 | 9 |
| GO:0098661 | BP | inorganic anion transmembrane transport | 9/765 | 11/2446 | 9,23083E+14 | CLCN7/SLC25A11/SLC1A1/CLCN3/SLC25A3/ANO6/CLCC1/TTYH3/ANO10 | 9 |
| GO:0140354 | BP | lipid import into cell | 9/765 | 11/2446 | 9,23083E+14 | ACSL1/ACSL3/ITGB1/THBS1/SLC2A1/SLC1A1/SLC1A5/SLC27A4/SLC27A1 | 9 |
| GO:0006904 | BP | vesicle docking involved in exocytosis | 14/765 | 21/2446 | 9,8403E+13 | STX3/EXOC6B/RAB3D/RALB/RAB13/RAB8A/RAB10/VAMP3/EXOC6/SCFD2/SCFD1/EXOC4/VPS45/VTI1B | 14 |
| GO:0008016 | BP | regulation of heart contraction | 14/765 | 21/2446 | 9,8403E+13 | ZMPSTE24/ATP1A1/JUP/ATP2A2/GJA1/SLC9A1/ATP2B1/FLNA/ATP2B4/GJA5/SLC1A1/CAV1/ASPH/STIM1 | 14 |
| GO:0033866 | BP | nucleoside bisphosphate biosynthetic process | 14/765 | 21/2446 | 9,8403E+13 | ACSL1/DCAKD/ACSL4/ACSL3/PDHB/SLC25A1/HSD17B12/HACD2/SCD5/SLC35B2/GCDH/PANK4/ELOVL5/TECR | 14 |
| GO:0034030 | BP | ribonucleoside bisphosphate biosynthetic process | 14/765 | 21/2446 | 9,8403E+13 | ACSL1/DCAKD/ACSL4/ACSL3/PDHB/SLC25A1/HSD17B12/HACD2/SCD5/SLC35B2/GCDH/PANK4/ELOVL5/TECR | 14 |
| GO:0034033 | BP | purine nucleoside bisphosphate biosynthetic process | 14/765 | 21/2446 | 9,8403E+13 | ACSL1/DCAKD/ACSL4/ACSL3/PDHB/SLC25A1/HSD17B12/HACD2/SCD5/SLC35B2/GCDH/PANK4/ELOVL5/TECR | 14 |
| GO:0044272 | BP | sulfur compound biosynthetic process | 19/765 | 32/2446 | 1,00123E+14 | ACSL1/MGST1/GLCE/MGST3/ACSL4/ACSL3/PDHB/BGN/SLC25A1/HSD17B12/HACD2/SCD5/CHST14/SLC35B2/CSGALNACT1/GCDH/E) | 19 |
| GO:0006497 | BP | protein lipidation | 10/765 | 13/2446 | 1,02281E+13 | PIGN/PIGQ/PIGG/ZDHHC3/RABL3/ZDHHC24/ZDHHC13/PIGK/PIGS/ZDHHC5 | 10 |
| GO:0015909 | BP | long-chain fatty acid transport | 10/765 | 13/2446 | 1,02281E+13 | ACSL1/ACSL3/THBS1/SLC2A1/ABCD3/ABCD1/CPT1A/SLC27A3/SLC27A4/SLC27A1 | 10 |
| GO:0042158 | BP | lipoprotein biosynthetic process | 10/765 | 13/2446 | 1,02281E+13 | PIGN/PIGQ/PIGG/ZDHHC3/RABL3/ZDHHC24/ZDHHC13/PIGK/PIGS/ZDHHC5 | 10 |
| GO:0046949 | BP | fatty-acyl-CoA biosynthetic process | 10/765 | 13/2446 | 1,02281E+13 | ACSL1/ACSL4/ACSL3/SLC25A1/HSD17B12/HACD2/SCD5/GCDH/ELOVL5/TECR | 10 |
| GO:0061337 | BP | cardiac conduction | 13/765 | 19/2446 | 1,02281E+13 | ZMPSTE24/ATP1A1/JUP/ATP2A2/GJA1/SLC9A1/ATP2B1/FLNA/ATP2B4/GJA5/CAV1/ASPH/STIM1 | 13 |
| GO:0006937 | BP | regulation of muscle contraction | 11/765 | 15/2446 | 1,06272E+14 | ATP1A1/JUP/ATP2A2/SLC9A1/ATP2B1/F2R/GJA5/CAV1/CALCRL/DOCK4/DOCK5 | 11 |
| GO:0015800 | BP | acidic amino acid transport | 11/765 | 15/2446 | 1,06272E+14 | PRAF2/SLC25A12/GLS/ITGB1/GJA1/SLC1A1/LRRC8D/LRRC8A/LRRC8C/SLC25A22/SLC25A13 | 11 |
| GO:0044242 | BP | cellular lipid catabolic process | 18/765 | 30/2446 | 1,11456E+13 | ABHD16A/PNPLA6/ILVBL/ACBD5/ACADM/HADHA/SMPD4/ALDH3A2/SGPL1/ABCD3/ABCD1/CPT1A/HADHB/SLC27A4/PDXDC1/ABHD | 18 |
| GO:0006665 | BP | sphingolipid metabolic process | 17/765 | 28/2446 | 1,26616E+14 | ST6GALNAC3/CLN6/ST6GALNAC4/FADS3/SMPD4/ALDH3A2/SPTLC1/VAPB/SGPL1/BAX/PDXDC1/HACD2/CERS6/CERS2/ELOVL5/TECR | 17 |
| GO:0098657 | BP | import into cell | 17/765 | 28/2446 | 1,26616E+14 | ACSL1/WNK1/SLC3A2/ACSL3/ATP1A1/ITGB1/THBS1/SLC2A1/SLC9A1/ATP2B4/SLC1A1/SLC1A5/SLC27A4/SLC27A1/ATAD1/SLC29A1/ | 17 |
| GO:0022406 | BP | membrane docking | 28/765 | 54/2446 | 1,26964E+13 | STX3/EXOC6B/STX12/NSF/STX10/STX7/STX6/USO1/VAPB/RAB3D/SNAP29/RALB/RAB13/STX17/RAB8A/RAB10/TUBA4A/STX4/STX5/ | 28 |
| GO:0044281 | BP | small molecule metabolic process | 105/765 | 264/2446 | 1,30742E+14 | BSG/SFXN3/DHCR24/NCEH1/PON2/CLN6/SUCLA2/CTPS1/ILVBL/SOAT1/ACBD5/ACADM/ERLIN2/ACSL1/VKORC1/FADS3/AAAS/HADH | 105 |
| GO:0006644 | BP | phospholipid metabolic process | 42/765 | 90/2446 | 1,41183E+14 | CEPT1/LPCAT4/ABHD16A/PIGN/PNPLA6/PIGQ/PIGG/PTDSS2/PHB2/ABCA3/HADHA/SMPD4/FADS1/ACSL3/CDS2/RAB5A/PI4KA/PTDS | 42 |
| GO:0009260 | BP | ribonucleotide biosynthetic process | 22/765 | 40/2446 | 1,54581E+14 | CTPS1/ACSL1/DCAKD/ACSL4/SLC25A12/ACSL3/UMPS/PDHB/ATP6V1A/SLC25A1/HSD17B12/HACD2/SCD5/SLC35B2/GCDH/DNAJC30 | 22 |
| GO:0090407 | BP | organophosphate biosynthetic process | 53/765 | 120/2446 | 1,68016E+14 | CEPT1/LPCAT4/PIGN/CTPS1/PIGQ/ACSL1/PIGG/PTDSS2/DCAKD/FADS1/ACSL4/SLC25A12/ACSL3/CDS2/UMPS/PDHB/RAB5A/NOS3/A | 53 |
| GO:0001568 | BP | blood vessel development | 58/765 | 134/2446 | 1,82687E+14 | ROBO1/BSG/RNF213/CTNNB1/PTPRM/SPARC/PTPRB/ACVRL1/SOS1/EMC10/SGPL1/HSPB1/ITGB1/THBS1/HMOX1/RRAS/JUP/PECAN | 58 |
| GO:0043436 | BP | oxoacid metabolic process | 66/765 | 156/2446 | 1,82687E+14 | BSG/PON2/SUCLA2/CTPS1/ILVBL/ACBD5/ACADM/ERLIN2/ACSL1/FADS3/AAAS/HADHA/NUP93/ALDH3A2/MGST3/SLC16A3/FADS1/A | 66 |
| GO:0019637 | BP | organophosphate metabolic process | 81/765 | 198/2446 | 1,84289E+14 | CEPT1/LPCAT4/ABHD16A/PIGN/SUCLA2/PNPLA6/CTPS1/PIGQ/ACSL1/PIGG/PTDSS2/PHB2/AAAS/ABCA3/HADHA/NUP93/SMPD4/DC | 81 |
| GO:0035637 | BP | multicellular organismal signaling | 13/765 | 20/2446 | 1,95207E+14 | ZMPSTE24/ATP1A1/JUP/ATP2A2/GJA1/SLC9A1/ATP2B1/FLNA/ATP2B4/GJA5/CAV1/ASPH/STIM1 | 13 |
| GO:0006890 | BP | retrograde vesicle-mediated transport, Golgi to endoplasmic reticulum | 25/765 | 48/2446 | 2,02615E+14 | COG4/NBAS/STX18/NSF/ZW10/ATP9A/SEC22B/RAB6A/TMED10/NAPA/COG7/ARF5/TMEM115/LMAN2/BNIP1/TMED2/RINT1/SCFD: | 25 |
| GO:0048514 | BP | blood vessel morphogenesis | 53/765 | 121/2446 | 2,0421E+14 | ROBO1/BSG/RNF213/CTNNB1/PTPRM/SPARC/PTPRB/ACVRL1/SOS1/EMC10/SGPL1/HSPB1/ITGB1/THBS1/HMOX1/RRAS/JUP/FLNA/I | 53 |
| GO:0030148 | BP | sphingolipid biosynthetic process | 12/765 | 18/2446 | 2,09218E+14 | ST6GALNAC3/ST6GALNAC4/SMPD4/ALDH3A2/SPTLC1/VAPB/SGPL1/HACD2/CERS6/CERS2/ELOVL5/VAPA | 12 |
| GO:0006658 | BP | phosphatidylserine metabolic process | 8/765 | 10/2446 | 2,09218E+14 | LPCAT4/ABHD16A/PTDSS2/PTDSS1/SLC27A1/ABHD12/OSBPL8/OSBPL5 | 8 |
| GO:0008053 | BP | mitochondrial fusion | 8/765 | 10/2446 | 2,09218E+14 | CHCHD3/MFN2/TFRC/BAX/USP30/MFF/STOML2/AFG3L2 | 8 |
| GO:0009581 | BP | detection of external stimulus | 8/765 | 10/2446 | 2,09218E+14 | CTNNB1/JUP/GNA11/GNAQ/CXCR4/GNB1/PKD2/PIEZO1 | 8 |
| GO:0009582 | BP | detection of abiotic stimulus | 8/765 | 10/2446 | 2,09218E+14 | CTNNB1/JUP/GNA11/GNAQ/CXCR4/GNB1/PKD2/PIEZO1 | 8 |
| GO:0017004 | BP | cytochrome complex assembly | 8/765 | 10/2446 | 2,09218E+14 | OXA1L/SCO1/CYBA/HCCS/COX20/TIMM21/BCS1L/COA3 | 8 |
| GO:0019915 | BP | lipid storage | 8/765 | 10/2446 | 2,09218E+14 | SOAT1/CDS2/PTPN2/CPT1A/CAV1/PNPLA2/OSBPL8/AUP1 | 8 |
| GO:0086003 | BP | cardiac muscle cell contraction | 8/765 | 10/2446 | 2,09218E+14 | ATP1A1/JUP/ATP2A2/GJA1/SLC9A1/FLNA/GJA5/CAV1 | 8 |
| GO:1903725 | BP | regulation of phospholipid metabolic process | 8/765 | 10/2446 | 2,09218E+14 | PHB2/ABCA3/ACSL3/SLC27A1/LPCAT1/DNAJC19/CHP1/STOML2 | 8 |
| GO:0002274 | BP | myeloid leukocyte activation | 51/765 | 116/2446 | 2,15742E+14 | SCAMP1/MAGT1/RAB18/DDOST/BRI3/MGST1/PTPRB/METTL7A/RAB27A/SNAP23/QSOX1/ADAM10/SNX4/NDUFC2/RAB3D/SNAP29 | 51 |
| GO:0006638 | BP | neutral lipid metabolic process | 11/765 | 16/2446 | 2,15742E+14 | ABHD16A/ACSL1/CTDNEP1/ACSL4/CPT1A/CAV1/SLC27A1/ABHD12/LPGAT1/PNPLA2/MBOAT7 | 11 |
| GO:0006639 | BP | acylglycerol metabolic process | 11/765 | 16/2446 | 2,15742E+14 | ABHD16A/ACSL1/CTDNEP1/ACSL4/CPT1A/CAV1/SLC27A1/ABHD12/LPGAT1/PNPLA2/MBOAT7 | 11 |
| GO:0035384 | BP | thioester biosynthetic process | 11/765 | 16/2446 | 2,15742E+14 | ACSL1/ACSL4/ACSL3/PDHB/SLC25A1/HSD17B12/HACD2/SCD5/GCDH/ELOVL5/TECR | 11 |
| GO:0071616 | BP | acyl-CoA biosynthetic process | 11/765 | 16/2446 | 2,15742E+14 | ACSL1/ACSL4/ACSL3/PDHB/SLC25A1/HSD17B12/HACD2/SCD5/GCDH/ELOVL5/TECR | 11 |
| GO:0002444 | BP | myeloid leukocyte mediated immunity | 48/765 | 108/2446 | 2,16713E+14 | SCAMP1/MAGT1/RAB18/DDOST/JAGN1/BRI3/MGST1/PTPRB/METTL7A/RAB27A/SNAP23/QSOX1/ADAM10/SNX4/NDUFC2/RAB3D/S | 48 |
| GO:0001944 | BP | vasculature development | 59/765 | 138/2446 | 2,19864E+14 | ROBO1/BSG/RNF213/CTNNB1/PTPRM/SPARC/PTPRB/ACVRL1/SOS1/EMC10/ADAM10/SGPL1/HSPB1/ITGB1/THBS1/HMOX1/RRAS/JI | 59 |
| GO:0045017 | BP | glycerolipid biosynthetic process | 32/765 | 66/2446 | 2,19974E+14 | CEPT1/LPCAT4/PIGN/PIGQ/ACSL1/PIGG/PTDSS2/CTDNEP1/ACSL4/ACSL3/CDS2/RAB5A/PI4KA/PTDSS1/RAB14/VAC14/ATM/PLD1/SL | 32 |
| GO:0044743 | BP | protein transmembrane import into intracellular organelle | 9/765 | 12/2446 | 2,20034E+14 | PEX1/PEX5/TIMM44/TOMM40/ROMO1/PEX6/TIMM50/DNAJC19/TIMM21 | 9 |
| GO:0048661 | BP | positive regulation of smooth muscle cell proliferation | 10/765 | 14/2446 | 2,20683E+14 | IRAK1/ABCC4/MFN2/GNAI2/THBS1/HMOX1/GJA1/PHB/STAT1/CALCRL | 10 |
| GO:0034620 | BP | cellular response to unfolded protein | 26/765 | 51/2446 | 2,26858E+13 | HYOU1/DDRGK1/GOSR2/TPP1/SSR1/ERMP1/TBL2/STUB1/WFS1/UFL1/VAPB/PTPN2/PTPN1/ATF6/TMEM33/BAX/TMED2/DAB2IP/S\ | 26 |
| GO:0006082 | BP | organic acid metabolic process | 66/765 | 158/2446 | 2,43099E+14 | BSG/PON2/SUCLA2/CTPS1/ILVBL/ACBD5/ACADM/ERLIN2/ACSL1/FADS3/AAAS/HADHA/NUP93/ALDH3A2/MGST3/SLC16A3/FADS1/A | 66 |
| GO:0002576 | BP | platelet degranulation | 15/765 | 25/2446 | 2,5282E+14 | APOOL/SPARC/QSOX1/ABCC4/APP/THBS1/CLU/PECAM1/FLNA/TUBA4A/SCCPDH/TMX3/TOR4A/VTI1B/CYB5R1 | 15 |
| GO:0019725 | BP | cellular homeostasis | 59/765 | 139/2446 | 2,6083E+14 | TM9SF4/CLN6/JAGN1/ATP2C1/RHOT1/SCO1/TMCO1/WFS1/LETM1/VAPB/TFRC/SLC4A2/ATP1A1/APP/HMOX1/CYBA/ATP2A2/GJA1, | 59 |
| GO:0003015 | BP | heart process | 18/765 | 32/2446 | 2,61847E+14 | ZMPSTE24/GNAI2/ATP1A1/JUP/ATP2A2/GJA1/SLC9A1/ATP2B1/FLNA/ATP2B4/GJA5/SLC1A1/CXCR4/CAV1/ASPH/STIM1/EXT2/TREX | 18 |
| GO:0001525 | BP | angiogenesis | 48/765 | 109/2446 | 2,62286E+14 | ROBO1/BSG/RNF213/CTNNB1/PTPRM/SPARC/PTPRB/ACVRL1/EMC10/HSPB1/ITGB1/THBS1/HMOX1/RRAS/JUP/FLNA/NF1/ATP2B4/ | 48 |
| GO:0019752 | BP | carboxylic acid metabolic process | 64/765 | 153/2446 | 2,66023E+14 | BSG/PON2/SUCLA2/CTPS1/ILVBL/ACBD5/ACADM/ERLIN2/ACSL1/FADS3/AAAS/HADHA/NUP93/ALDH3A2/MGST3/SLC16A3/FADS1/A | 64 |
| GO:0033865 | BP | nucleoside bisphosphate metabolic process | 17/765 | 30/2446 | 3,01728E+14 | SUCLA2/ACSL1/DCAKD/ACSL4/TPST2/ACSL3/PDHB/ABCD1/SLC25A1/HSD17B12/HACD2/SCD5/SLC35B2/GCDH/PANK4/ELOVL5/TECR | 17 |
| GO:0033875 | BP | ribonucleoside bisphosphate metabolic process | 17/765 | 30/2446 | 3,01728E+14 | SUCLA2/ACSL1/DCAKD/ACSL4/TPST2/ACSL3/PDHB/ABCD1/SLC25A1/HSD17B12/HACD2/SCD5/SLC35B2/GCDH/PANK4/ELOVL5/TECR | 17 |
| GO:0034032 | BP | purine nucleoside bisphosphate metabolic process | 17/765 | 30/2446 | 3,01728E+14 | SUCLA2/ACSL1/DCAKD/ACSL4/TPST2/ACSL3/PDHB/ABCD1/SLC25A1/HSD17B12/HACD2/SCD5/SLC35B2/GCDH/PANK4/ELOVL5/TECR | 17 |
| GO:0070585 | BP | protein localization to mitochondrion | 22/765 | 42/2446 | 3,01728E+14 | PARL/OXA1L/HAX1/TIMM44/MFN2/TOMM40/HK1/YWHAQ/BCAP31/ROMO1/TIMM10/BAX/TIMM50/MGARP/BAD/DNAJC19/TIMN | 22 |
| GO:2001234 | BP | negative regulation of apoptotic signaling pathway | 22/765 | 42/2446 | 3,01728E+14 | HYOU1/VDAC2/TMEM161A/NDUFS3/HSPB1/GNAI2/THBS1/HMOX1/CLU/PTPN1/ITGA6/NOS3/CTNNA1/PTTG1IP/ACVR1/BAX/TRAD | 22 |
| GO:0009306 | BP | protein secretion | 28/765 | 57/2446 | 3,01728E+14 | JAGN1/WLS/MIA2/ACSL4/RAB3D/SLC2A1/GJA1/F2R/IDH2/TMED10/CPT1A/RAB13/SLC16A1/RAB8A/RAB10/STX4/ADAM9/TCIRG1/N | 28 |
| GO:0035592 | BP | establishment of protein localization to extracellular region | 28/765 | 57/2446 | 3,01728E+14 | JAGN1/WLS/MIA2/ACSL4/RAB3D/SLC2A1/GJA1/F2R/IDH2/TMED10/CPT1A/RAB13/SLC16A1/RAB8A/RAB10/STX4/ADAM9/TCIRG1/N | 28 |
| GO:0006893 | BP | Golgi to plasma membrane transport | 13/765 | 21/2446 | 3,18299E+14 | EXOC6B/NSF/ACSL3/RAB10/VAMP3/EXOC6/EXOC4/EXOC2/RAB34/OSBPL5/EXOC1/ARFGEF2 | 13 |
| GO:0051048 | BP | negative regulation of secretion | 13/765 | 21/2446 | 3,18299E+14 | JAGN1/GNAI2/HMOX1/GJA1/F2R/NOTCH1/IDH2/VAMP3/RHBDF2/PNKD/RHBDF1/RAB33B/SLC30A1 | 13 |
| GO:1903531 | BP | negative regulation of secretion by cell | 13/765 | 21/2446 | 3,18299E+14 | JAGN1/GNAI2/HMOX1/GJA1/F2R/NOTCH1/IDH2/VAMP3/RHBDF2/PNKD/RHBDF1/RAB33B/SLC30A1 | 13 |
| GO:0009152 | BP | purine ribonucleotide biosynthetic process | 19/765 | 35/2446 | 3,30976E+14 | ACSL1/DCAKD/ACSL4/SLC25A12/ACSL3/PDHB/ATP6V1A/SLC25A1/HSD17B12/HACD2/SCD5/SLC35B2/GCDH/DNAJC30/PANK4/ELOV | 19 |
| GO:1901568 | BP | fatty acid derivative metabolic process | 12/765 | 19/2446 | 3,56408E+14 | ABHD16A/ACSL1/ACSL4/ACSL3/ABCD1/SLC25A1/HSD17B12/HACD2/SCD5/GCDH/ELOVL5/TECR | 12 |
| GO:0006814 | BP | sodium ion transport | 11/765 | 17/2446 | 3,92662E+14 | WNK1/SLC3A2/ATP1A1/SLC9A1/ATP2B4/STOM/PKD2/NDUFA9/ANO6/CHP1/SLC38A10 | 11 |
| GO:0006998 | BP | nuclear envelope organization | 11/765 | 17/2446 | 3,92662E+14 | REEP4/NUP93/CTDNEP1/ZMPSTE24/EMD/ANKLE2/LEMD2/TMEM43/SUN2/VPS4A/LEMD3 | 11 |
| GO:0035337 | BP | fatty-acyl-CoA metabolic process | 11/765 | 17/2446 | 3,92662E+14 | ACSL1/ACSL4/ACSL3/ABCD1/SLC25A1/HSD17B12/HACD2/SCD5/GCDH/ELOVL5/TECR | 11 |
| GO:0048659 | BP | smooth muscle cell proliferation | 11/765 | 17/2446 | 3,92662E+14 | CTNNB1/IRAK1/ABCC4/MFN2/GNAI2/THBS1/HMOX1/GJA1/PHB/STAT1/CALCRL | 11 |
| GO:0048660 | BP | regulation of smooth muscle cell proliferation | 11/765 | 17/2446 | 3,92662E+14 | CTNNB1/IRAK1/ABCC4/MFN2/GNAI2/THBS1/HMOX1/GJA1/PHB/STAT1/CALCRL | 11 |
| GO:0040017 | BP | positive regulation of locomotion | 45/765 | 103/2446 | 3,94004E+14 | DOCK1/DDRGK1/STX3/RRAS2/WNK1/SPARC/EMC10/ADAM10/HSPB1/GNAI2/APP/ITGB1/THBS1/HMOX1/RRAS/PECAM1/FLNA/ITG/ | 45 |
| GO:0002790 | BP | peptide secretion | 28/765 | 58/2446 | 3,94741E+14 | JAGN1/WLS/MIA2/ACSL4/RAB3D/SLC2A1/GJA1/F2R/IDH2/TMED10/CPT1A/RAB13/SLC16A1/RAB8A/RAB10/STX4/ADAM9/TCIRG1/N | 28 |
| GO:0001894 | BP | tissue homeostasis | 20/765 | 38/2446 | 4,01595E+14 | TPP1/ABCA3/RAB3D/TFRC/HSPB1/ITGB1/SLC2A1/PECAM1/GJA1/NF1/F2R/NOS3/CDH5/SLC1A1/NOTCH1/CLCN3/BAX/TCIRG1/LPCA | 20 |
| GO:0002275 | BP | myeloid cell activation involved in immune response | 46/765 | 106/2446 | 4,10379E+14 | SCAMP1/MAGT1/RAB18/DDOST/BRI3/MGST1/PTPRB/METTL7A/RAB27A/SNAP23/QSOX1/ADAM10/SNX4/NDUFC2/RAB3D/SNAP29 | 46 |
| GO:0032880 | BP | regulation of protein localization | 76/765 | 190/2446 | 4,19395E+14 | DDRGK1/TM9SF4/STX3/DNAJB6/PARL/CTNNB1/JAGN1/WLS/ZDHHC3/STX10/HAX1/ADAM10/RAB29/STX7/STX6/ACSL4/B3GAT3/UF | 76 |
| GO:0006029 | BP | proteoglycan metabolic process | 10/765 | 15/2446 | 4,19395E+14 | CTNNB1/GLCE/FAM20B/B3GAT3/BGN/CHST14/CSGALNACT1/CANT1/EXT2/B4GALT7 | 10 |
| GO:0006493 | BP | protein O-linked glycosylation | 10/765 | 15/2446 | 4,19395E+14 | ST6GALNAC4/GALNT7/ST6GAL1/GALNT2/GALNT1/POMT1/TMTC3/XXYLT1/GALNT6/SDF2L1 | 10 |
| GO:1901570 | BP | fatty acid derivative biosynthetic process | 10/765 | 15/2446 | 4,19395E+14 | ACSL1/ACSL4/ACSL3/SLC25A1/HSD17B12/HACD2/SCD5/GCDH/ELOVL5/TECR | 10 |
| GO:0016197 | BP | endosomal transport | 34/765 | 74/2446 | 4,25615E+14 | EVI5/GOSR2/ARL1/TMCC1/RAB35/RIC1/STX10/RNF126/RAB29/STX6/VAMP4/SNX4/RAB6A/RAB13/RAB8A/RAB10/RAB14/STX5/VAN | 34 |
| GO:0019216 | BP | regulation of lipid metabolic process | 29/765 | 61/2446 | 4,31977E+13 | ACADM/ERLIN2/ACSL1/PHB2/ABCA3/STUB1/CTDNEP1/FADS1/ERLIN1/ZMPSTE24/ACSL3/ATP1A1/ABCD1/CPT1A/TEK/CAV1/PPP2R5 | 29 |
| GO:0034394 | BP | protein localization to cell surface | 9/765 | 13/2446 | 4,42631E+14 | TM9SF4/STX3/CTNNB1/HSP90AB1/FLNA/STX4/PTPRK/GBF1/PIGK | 9 |
| GO:0046470 | BP | phosphatidylcholine metabolic process | 9/765 | 13/2446 | 4,42631E+14 | CEPT1/LPCAT4/PNPLA6/ABCA3/ACSL3/SLC27A1/LPCAT2/LPCAT1/MBOAT7 | 9 |
| GO:0006851 | BP | mitochondrial calcium ion transmembrane transport | 8/765 | 11/2446 | 4,53968E+14 | PARL/PHB2/LETM1/PHB/MCU/YME1L1/STOML2/AFG3L2 | 8 |
| GO:0051788 | BP | response to misfolded protein | 8/765 | 11/2446 | 4,53968E+14 | RNF5/STUB1/RNF126/CLU/DNAJB14/SDF2L1/DNAJB12/AUP1 | 8 |
| GO:1901342 | BP | regulation of vasculature development | 25/765 | 51/2446 | 4,54126E+14 | CTNNB1/PTPRM/SPARC/EMC10/ADAM10/HSPB1/ITGB1/THBS1/HMOX1/RRAS/JUP/NF1/ATP2B4/EPHA2/NOS3/CDH5/TIE1/STAT1/C | 25 |

| | | | | | | | |
|---|---|---|---|---|---|---|---|
| GO:0043299 | BP | leukocyte degranulation | 45/765 | 104/2446 | 4,55116E+14 | SCAMP1/MAGT1/RAB18/DDOST/BRI3/MGST1/PTPRB/METTL7A/RAB27A/SNAP23/QSOX1/ADAM10/SNX4/NDUFC2/RAB3D/SNAP29 | 45 |
| GO:0072655 | BP | establishment of protein localization to mitochondrion | 21/765 | 41/2446 | 4,61483E+14 | PARL/OXA1L/HAX1/TIMM44/MFN2/TOMM40/HK1/YWHAQ/BCAP31/ROMO1/TIMM10/BAX/TIMM50/MGARP/BAD/DNAJC19/TIMM | 21 |
| GO:0008015 | BP | blood circulation | 28/765 | 59/2446 | 4,90801E+14 | ECE1/ACVRL1/ZMPSTE24/CYB5R3/ATP1A1/HMOX1/JUP/ATP2A2/GJA1/SLC9A1/ATP2B1/FLNA/ATP2B4/F2R/NOS3/CDH5/GJA5/STAT | 28 |
| GO:0060047 | BP | heart contraction | 16/765 | 29/2446 | 4,90801E+14 | ZMPSTE24/ATP1A1/JUP/ATP2A2/GJA1/SLC9A1/ATP2B1/FLNA/ATP2B4/GJA5/SLC1A1/CXCR4/CAV1/ASPH/STIM1/EXT2 | 16 |
| GO:0098876 | BP | vesicle-mediated transport to the plasma membrane | 16/765 | 29/2446 | 4,90801E+14 | STX3/EXOC6B/NSF/ACSL3/SLC1A1/RAB8A/RAB10/VAMP3/EXOC8/EXOC6/EXOC2/RAB34/OSBPL5/EXOC1/ARFGEF2 | 16 |
| GO:1903409 | BP | reactive oxygen species biosynthetic process | 13/765 | 22/2446 | 4,92599E+14 | RAB27A/CYB5R3/HSP90AB1/CLU/CYBA/ATP2B4/NOS3/ABCD1/CLCN3/CAV1/PKD2/GBF1/PIKFYVE | 13 |
| GO:0031984 | CC | organelle subcompartment | 303/765 | 440/2446 | 8,63494E-56 | ATP11C/HM13/SCAMP1/MAGT1/SPCS2/BSG/ST6GALNAC3/NOMO1/RAB18/DDRGK1/COG4/DHCR24/NCEH1/DHRS7B/DDOST/CEPT | 303 |
| GO:0042175 | CC | nuclear outer membrane-endoplasmic reticulum membrane network | 222/765 | 306/2446 | 2,90079E-40 | ATP11C/HM13/MAGT1/SPCS2/BSG/NOMO1/RAB18/DDRGK1/DHCR24/NCEH1/DHRS7B/DDOST/CEPT1/LPCAT4/CLN6/TAP1/RNF5/P | 222 |
| GO:0005789 | CC | endoplasmic reticulum membrane | 218/765 | 298/2446 | 2,90079E-40 | ATP11C/HM13/MAGT1/SPCS2/BSG/NOMO1/RAB18/DDRGK1/DHCR24/NCEH1/DHRS7B/DDOST/CEPT1/LPCAT4/CLN6/TAP1/RNF5/P | 218 |
| GO:0098827 | CC | endoplasmic reticulum subcompartment | 219/765 | 301/2446 | 5,73344E-40 | ATP11C/HM13/MAGT1/SPCS2/BSG/NOMO1/RAB18/DDRGK1/DHCR24/NCEH1/DHRS7B/DDOST/CEPT1/LPCAT4/CLN6/TAP1/RNF5/P | 219 |
| GO:0005783 | CC | endoplasmic reticulum | 273/765 | 444/2446 | 8,73614E-32 | ATP11C/HM13/MAGT1/SPCS2/BSG/HYOU1/NOMO1/RAB18/DDRGK1/DHCR24/NCEH1/DHRS7B/DDOST/CEPT1/LPCAT4/CLN6/TAP1/ | 273 |
| GO:0098588 | CC | bounding membrane of organelle | 270/765 | 459/2446 | 9,81286E-26 | ATP11C/ROBO1/SCAMP1/MAGT1/BSG/ST6GALNAC3/APOOL/RAB18/VDAC2/COG4/DHCR24/PEX1/STX3/DDOST/TAP1/GOSR2/SLC3 | 270 |
| GO:0031300 | CC | intrinsic component of organelle membrane | 96/765 | 123/2446 | 2,93544E-11 | HM13/APOOL/SFXN3/TAP1/ECE1/STX12/CHCHD3/PIGG/TBL2/WLS/RAB35/TRAM1/RHOT1/OXA1L/UNC50/SCO1/CHCHD6/TMCO1/S | 96 |
| GO:0031966 | CC | mitochondrial membrane | 136/765 | 204/2446 | 5,89861E-11 | APOOL/NDUFB6/VDAC2/SFXN3/NDUFA6/RNF5/PARL/ACADM/CHCHD3/ACSL1/NDUFV2/NDUFB5/TMEM126B/NDUFB8/PHB2/MGS | 136 |
| GO:0005740 | CC | mitochondrial envelope | 136/765 | 206/2446 | 2,38983E-10 | APOOL/NDUFB6/VDAC2/SFXN3/NDUFA6/RNF5/PARL/ACADM/CHCHD3/ACSL1/NDUFV2/NDUFB5/TMEM126B/NDUFB8/PHB2/MGS | 136 |
| GO:0031301 | CC | integral component of organelle membrane | 91/765 | 116/2446 | 2,90101E-10 | HM13/APOOL/SFXN3/TAP1/STX12/CHCHD3/PIGG/TBL2/WLS/TRAM1/RHOT1/OXA1L/UNC50/SCO1/CHCHD6/TMCO1/STX10/EMC10 | 91 |
| GO:0031967 | CC | organelle envelope | 191/765 | 335/2446 | 1,41222E-09 | APOOL/NDUFB6/VDAC2/SFXN3/NDUFA6/CEPT1/RNF5/DPY19L1/PARL/TMEM120B/ACADM/RNF123/CHCHD3/ACSL1/NDUFV2/NDU | 191 |
| GO:0031975 | CC | envelope | 191/765 | 335/2446 | 1,41222E-09 | APOOL/NDUFB6/VDAC2/SFXN3/NDUFA6/CEPT1/RNF5/DPY19L1/PARL/TMEM120B/ACADM/RNF123/CHCHD3/ACSL1/NDUFV2/NDU | 191 |
| GO:0098796 | CC | membrane protein complex | 156/765 | 257/2446 | 1,23962E-08 | HM13/MAGT1/SPCS2/APOOL/NDUFB6/VDAC2/STX3/NDUFA6/DDOST/TAP1/GOSR2/CTNNB1/COL13A1/STX12/PIGQ/CHCHD3/STX1 | 156 |
| GO:0005887 | CC | integral component of plasma membrane | 81/765 | 102/2446 | 3,31233E-08 | ATP11C/ROBO1/MAGT1/BSG/BCAM/SLC33A1/COL13A1/PTPRM/PTPRB/ACVRL1/PTPRF/SLC25A11/ICAM2/SLC16A3/FLRT2/MPZL1/ | 81 |
| GO:0031226 | CC | intrinsic component of plasma membrane | 82/765 | 105/2446 | 1,24957E-07 | ATP11C/ROBO1/MAGT1/BSG/BCAM/SLC33A1/COL13A1/PTPRM/PTPRB/ACVRL1/PTPRF/SLC25A11/ICAM2/SLC16A3/FLRT2/MPZL1/ | 82 |
| GO:0000139 | CC | Golgi membrane | 121/765 | 184/2446 | 2,24839E-07 | SCAMP1/BSG/ST6GALNAC3/COG4/DHCR24/GOSR2/SLC33A1/ST6GALNAC4/STX12/COG8/ARL1/ATP2C1/SLC30A6/COG2/STX18/GAL | 121 |
| GO:0098791 | CC | Golgi apparatus subcompartment | 132/765 | 211/2446 | 1,49717E-06 | ATP11C/SCAMP1/BSG/ST6GALNAC3/COG4/DHCR24/GOSR2/SLC33A1/ST6GALNAC4/STX12/COG8/ARL1/ATP2C1/SLC30A6/COG2/ST | 132 |
| GO:0019866 | CC | organelle inner membrane | 106/765 | 157/2446 | 6,85363E-06 | APOOL/NDUFB6/SFXN3/NDUFA6/DPY19L1/PARL/TMEM120B/CHCHD3/NDUFV2/NDUFB5/TMEM126B/NDUFB8/PHB2/NDUFV1/HA | 106 |
| GO:0005794 | CC | Golgi apparatus | 181/765 | 331/2446 | 1,8829E-05 | ATP11C/SCAMP1/BSG/ST6GALNAC3/RAB18/COG4/DHCR24/TM9SF4/CEPT1/UNC45A/GOSR2/TSC2/TPP1/SLC33A1/ST6GALNAC4/ST | 181 |
| GO:0005743 | CC | mitochondrial inner membrane | 95/765 | 141/2446 | 0,00136051 | APOOL/NDUFB6/SFXN3/NDUFA6/PARL/CHCHD3/NDUFV2/NDUFB5/TMEM126B/NDUFB8/PHB2/NDUFV1/HADHA/COX4I1/NDUFB1 | 95 |
| GO:0030176 | CC | integral component of endoplasmic reticulum membrane | 44/765 | 54/2446 | 191,55035 | HM13/TAP1/PIGG/TBL2/TRAM1/TMCO1/EMC10/EMC8/ZMPSTE24/WFS1/SGPL1/HLA-E/ATF6/BCAP31/TMEM33/BNIP1/PKD2/STIM | 44 |
| GO:0031227 | CC | intrinsic component of endoplasmic reticulum membrane | 45/765 | 56/2446 | 245,3941004 | HM13/TAP1/PIGG/TBL2/TRAM1/TMCO1/EMC10/EMC8/ZMPSTE24/WFS1/SGPL1/HLA-E/ATF6/DNAJB2/BCAP31/TMEM33/BNIP1/PK | 45 |
| GO:0098800 | CC | inner mitochondrial membrane protein complex | 44/765 | 55/2446 | 636,3763277 | APOOL/NDUFB6/NDUFA6/CHCHD3/NDUFV2/NDUFB5/NDUFB8/PHB2/NDUFV1/COX4I1/NDUFB10/CHCHD6/NDUFS8/UQCRQ/NDUF | 44 |
| GO:0005739 | CC | mitochondrion | 168/765 | 349/2446 | 682,608815 | BSG/APOOL/NDUFB6/VDAC2/SFXN3/NDUFA6/RNF5/PARL/SUCLA2/ACADM/CHCHD3/ACSL1/NDUFV2/NDUFB5/TMEM126B/NDUFB | 168 |
| GO:0031201 | CC | SNARE complex | 23/765 | 23/2446 | 1653,204893 | STX3/GOSR2/STX12/STX18/STX10/SNAP23/STX7/STX6/VAMP4/SEC22B/SNX4/SNAP29/NAPA/STX17/STX4/BNIP1/STX5/VAMP3/VTI1 | 23 |
| GO:0019867 | CC | outer membrane | 50/765 | 70/2446 | 22857,1383 | APOOL/VDAC2/CHCHD3/ACSL1/PHB2/MGST1/SMPD4/RHOT1/CHCHD6/HAX1/ACSL4/MTX2/MFN2/ACSL3/TOMM40/CYB5R3/CNP/ | 50 |
| GO:0031968 | CC | organelle outer membrane | 50/765 | 70/2446 | 22857,1383 | APOOL/VDAC2/CHCHD3/ACSL1/PHB2/MGST1/SMPD4/RHOT1/CHCHD6/HAX1/ACSL4/MTX2/MFN2/ACSL3/TOMM40/CYB5R3/CNP/ | 50 |
| GO:0140534 | CC | endoplasmic reticulum protein-containing complex | 41/765 | 53/2446 | 29646,57064 | HM13/MAGT1/SPCS2/HYOU1/DDOST/TAP1/NBAS/PIGQ/SEC11A/EMC10/SPTLC1/ZW10/EMC8/RPN1/RPN2/STT3A/SSR4/SEC61G/SE | 41 |
| GO:0030667 | CC | secretory granule membrane | 45/765 | 61/2446 | 42972,24959 | SCAMP1/MAGT1/BSG/RAB18/STX3/DDOST/BRI3/SPARC/MGST1/PTPRB/ABCA3/RAB27A/SNAP23/ADAM10/ABCC4/NDUFC2/RAB3D | 45 |
| GO:0005741 | CC | mitochondrial outer membrane | 44/765 | 61/2446 | 240059,9714 | APOOL/VDAC2/CHCHD3/ACSL1/PHB2/MGST1/RHOT1/CHCHD6/HAX1/ACSL4/MTX2/MFN2/ACSL3/TOMM40/CYB5R3/CNP/GJA1/HK | 44 |
| GO:0005747 | CC | mitochondrial respiratory chain complex I | 24/765 | 26/2446 | 537817,5524 | NDUFB6/NDUFA6/NDUFV2/NDUFB5/NDUFB8/NDUFV1/NDUFB10/NDUFS8/NDUFS4/NDUFA2/NDUFS5/NDUFS2/NDUFS6/NDUFS3/ | 24 |
| GO:0030964 | CC | NADH dehydrogenase complex | 24/765 | 26/2446 | 537817,5524 | NDUFB6/NDUFA6/NDUFV2/NDUFB5/NDUFB8/NDUFV1/NDUFB10/NDUFS8/NDUFS4/NDUFA2/NDUFS5/NDUFS2/NDUFS6/NDUFS3/ | 24 |
| GO:0045271 | CC | respiratory chain complex I | 24/765 | 26/2446 | 537817,5524 | NDUFB6/NDUFA6/NDUFV2/NDUFB5/NDUFB8/NDUFV1/NDUFB10/NDUFS8/NDUFS4/NDUFA2/NDUFS5/NDUFS2/NDUFS6/NDUFS3/ | 24 |
| GO:0005746 | CC | mitochondrial respirasome | 30/765 | 36/2446 | 623397,1734 | NDUFB6/NDUFA6/NDUFV2/NDUFB5/NDUFB8/NDUFV1/COX4I1/NDUFB10/OXA1L/NDUFS8/UQCRQ/NDUFS4/NDUFA2/NDUFS5/ND | 30 |
| GO:0070469 | CC | respirasome | 30/765 | 36/2446 | 623397,1734 | NDUFB6/NDUFA6/NDUFV2/NDUFB5/NDUFB8/NDUFV1/COX4I1/NDUFB10/OXA1L/NDUFS8/UQCRQ/NDUFS4/NDUFA2/NDUFS5/ND | 30 |
| GO:0009986 | CC | cell surface | 52/765 | 80/2446 | 1376053,753 | HM13/ROBO1/BCAM/ECE1/CAPN5/SPARC/PHB2/SLC3A2/ACVRL1/ADAM10/TNFRSF10C/ABCC4/VAMP4/MPZL1/CLPTM1/TFRC/APP | 52 |
| GO:0098803 | CC | respiratory chain complex | 28/765 | 34/2446 | 3440295,491 | NDUFB6/NDUFA6/NDUFV2/NDUFB5/NDUFB8/NDUFV1/COX4I1/NDUFB10/NDUFS8/UQCRQ/NDUFS4/NDUFA2/NDUFS5/NDUFS2/N | 28 |
| GO:1990204 | CC | oxidoreductase complex | 31/765 | 40/2446 | 6679556,338 | NDUFB6/NDUFA6/NDUFV2/NDUFB5/NDUFB8/NDUFV1/COX4I1/NDUFB10/NDUFS8/UQCRQ/NDUFS4/NDUFA2/NDUFS5/NDUFS2/NDUFS6/N | 31 |
| GO:0012506 | CC | vesicle membrane | 84/765 | 160/2446 | 21814290,56 | SCAMP1/MAGT1/BSG/RAB18/STX3/DDOST/TAP1/GOSR2/STX12/WLS/BRI3/SPARC/RAB35/MGST1/PTPRB/ABCA3/RAB27A/STX10/S | 84 |
| GO:0030659 | CC | cytoplasmic vesicle membrane | 82/765 | 156/2446 | 30431909,63 | SCAMP1/MAGT1/BSG/RAB18/STX3/DDOST/TAP1/GOSR2/STX12/WLS/BRI3/SPARC/RAB35/MGST1/PTPRB/ABCA3/RAB27A/STX10/S | 82 |
| GO:0005802 | CC | trans-Golgi network | 43/765 | 67/2446 | 61021592,05 | ATP11C/SCAMP1/COG4/COG8/ARL1/ATP2C1/COG2/WLS/RIC1/VAMP4/STX10/ADAM10/SCAMP3/RAB29/TGOLN2/STX6/ATP9A/VA | 43 |
| GO:0098573 | CC | intrinsic component of mitochondrial membrane | 29/765 | 39/2446 | 93966143,04 | APOOL/SFXN3/CHCHD3/RHOT1/OXA1L/SCO1/CHCHD6/MTX2/MFN2/TOMM40/TMEM11/CPT1A/SLC25A3/MTX1/IMMT/SFXN4/RH | 29 |
| GO:0032588 | CC | trans-Golgi network membrane | 24/765 | 30/2446 | 129158430,9 | SCAMP1/COG4/COG8/ARL1/COG2/RIC1/STX10/SCAMP3/STX6/VAMP4/APP/IGF2R/RAB6A/RAB8A/COG7/VAMP3/COG1/SCAMP4/V | 24 |
| GO:0099503 | CC | secretory vesicle | 86/765 | 171/2446 | 172498802,9 | SCAMP1/MAGT1/BSG/APOOL/RAB18/VDAC2/STX3/DDOST/ECE1/STX12/EBAG9/BRI3/SPARC/RAB35/MGST1/PTPRB/VEZT/ABCA3/M | 86 |
| GO:0032592 | CC | integral component of mitochondrial membrane | 28/765 | 38/2446 | 215410155,2 | APOOL/SFXN3/CHCHD3/RHOT1/OXA1L/SCO1/CHCHD6/MTX2/TOMM40/TMEM11/CPT1A/SLC25A3/MTX1/IMMT/SFXN4/RHOT2/M | 28 |
| GO:0031410 | CC | cytoplasmic vesicle | 187/765 | 454/2446 | 1212653992 | ATP11C/SCAMP1/MAGT1/BSG/APOOL/HYOU1/RAB18/VDAC2/TM9SF4/STX3/DDOST/CLN6/TAP1/GOSR2/TPP1/ECE1/STX12/VPS8/T | 187 |
| GO:0098798 | CC | mitochondrial protein-containing complex | 49/765 | 86/2446 | 1212653992 | APOOL/NDUFB6/NDUFA6/CHCHD3/NDUFV2/NDUFB5/NDUFB8/PHB2/NDUFV1/HADHA/COX4I1/NDUFB10/CHCHD6/NDUFS8/UQCR | 49 |
| GO:0097708 | CC | intracellular vesicle | 187/765 | 455/2446 | 1416216708 | ATP11C/SCAMP1/MAGT1/BSG/APOOL/HYOU1/RAB18/VDAC2/TM9SF4/STX3/DDOST/CLN6/TAP1/GOSR2/TPP1/ECE1/STX12/VPS8/T | 187 |
| GO:0008021 | CC | synaptic vesicle | 24/765 | 33/2446 | 2451992956 | SCAMP1/VDAC2/STX3/STX12/RAB35/STX10/ADAM10/STX7/STX6/VAMP4/SEC22B/RAB3D/APP/RAB5A/RAB13/CLCN3/RAB8A/RAB1 | 24 |
| GO:0070382 | CC | exocytic vesicle | 26/765 | 37/2446 | 2485650355 | SCAMP1/VDAC2/STX3/STX12/RAB35/RAB27A/STX10/ADAM10/STX7/STX6/VAMP4/SEC22B/RAB3D/APP/RAB5A/RAB6A/RAB13/CLC | 26 |
| GO:0030141 | CC | secretory granule | 75/765 | 152/2446 | 2862866358 | SCAMP1/MAGT1/BSG/APOOL/RAB18/VDAC2/STX3/DDOST/ECE1/EBAG9/BRI3/SPARC/MGST1/PTPRB/VEZT/ABCA3/METTL7A/RAB2 | 75 |
| GO:0035579 | CC | specific granule membrane | 13/765 | 14/2446 | 5785407424 | SCAMP1/PTPRB/SNAP23/ADAM10/CYBA/STOM/HMOX2/CKAP4/PLD1/ANO6/CD93/TMEM30A/CMTM6 | 13 |
| GO:0005773 | CC | vacuole | 72/765 | 148/2446 | 8738224294 | ATP11C/MAGT1/STX3/DDOST/TSC2/TPP1/ECE1/TMEM106B/LRBA/BRI3/SLC3A2/MGST1/CLCN7/RAB27A/NSF/SNAP23/RAB29/STX7 | 72 |
| GO:0033116 | CC | endoplasmic reticulum-Golgi intermediate compartment membrane | 18/765 | 23/2446 | 9433588405 | ROBO1/TAP1/GOSR2/SEC22B/TMED10/BCAP31/STX17/RAB2A/LMAN2/STX5/TMED2/YIF1B/PIEZO1/ERGIC1/TBC1D20/TMED9/CTSZ | 18 |
| GO:0035577 | CC | azurophil granule membrane | 14/765 | 16/2446 | 1,04558E+14 | MAGT1/DDOST/BRI3/MGST1/NDUFC2/RAB3D/SNAP29/STOM/RAB5C/CKAP4/LPCAT1/TMEM30A/CMTM6/VAPA | 14 |
| GO:0031304 | CC | intrinsic component of mitochondrial inner membrane | 19/765 | 25/2446 | 1,05693E+14 | APOOL/SFXN3/CHCHD3/OXA1L/SCO1/CHCHD6/MTX2/TMEM11/SLC25A3/MTX1/IMMT/SFXN4/MCU/SFXN2/SFXN1/SLC25A19/DNA | 19 |
| GO:0031305 | CC | integral component of mitochondrial inner membrane | 19/765 | 25/2446 | 1,05693E+14 | APOOL/SFXN3/CHCHD3/OXA1L/SCO1/CHCHD6/MTX2/TMEM11/SLC25A3/MTX1/IMMT/SFXN4/MCU/SFXN2/SFXN1/SLC25A19/DNA | 19 |
| GO:0005774 | CC | vacuolar membrane | 49/765 | 92/2446 | 1,38649E+14 | ATP11C/MAGT1/DDOST/ECE1/TMEM106B/BRI3/SLC3A2/MGST1/CLCN7/NSF/STX7/NDUFC2/RAB3D/SNAP29/RPN2/HSP90AB1/STO | 49 |
| GO:0005795 | CC | Golgi stack | 24/765 | 36/2446 | 2,27085E+14 | COG2/GOLIM4/NSF/USO1/ST6GAL1/BCAP31/RAB14/MGAT2/GALNT2/GALNT1/TMEM115/TMED2/RASIP1/LPCAT2/TMEM87A/GOL | 24 |
| GO:0009925 | CC | basal plasma membrane | 24/765 | 36/2446 | 2,27085E+14 | BSG/CTNNB1/SLC3A2/SLC16A3/ABCC4/TFRC/SLC4A2/ATP1A1/HSP90AB1/TACSTD2/SLC2A1/ITGB4/SLC9A1/ATP2B1/ATP2B4/ITGA3/ | 24 |
| GO:0030133 | CC | transport vesicle | 43/765 | 79/2446 | 2,35244E+14 | SCAMP1/VDAC2/STX3/GOSR2/STX12/YIPF3/RAB35/GOLIM4/RAB27A/STX10/ADAM10/STX7/TGOLN2/STX6/USO1/VAMP4/SEC22B/ | 43 |
| GO:0016324 | CC | apical plasma membrane | 30/765 | 50/2446 | 3,95686E+14 | RAB18/STX3/SLC3A2/RAB27A/HAX1/SLC16A3/ABCC4/HIP1R/SLC4A2/ATP1A1/HSP90AB1/SLC2A1/GJA1/SLC9A1/ATP2B1/ABCC1/ATI | 30 |
| GO:0005811 | CC | lipid droplet | 22/765 | 33/2446 | 4,90069E+14 | RAB18/IRAK1/METTL7A/CTDNEP1/ZW10/ACSL4/ACSL3/CYB5R3/RAB5C/BCAP31/UBE2G2/CAV1/CKAP4/NSDHL/LPCAT2/SCCPDH/LP | 22 |
| GO:0044232 | CC | organelle membrane contact site | 13/765 | 16/2446 | 8,41211E+14 | TMCC1/ACSL4/TOMM40/BCAP31/STX17/RAB32/TMEM41B/MBOAT7/CLCC1/RMDN3/OSBPL5/SACM1L/TMX2 | 13 |
| GO:0042581 | CC | specific granule | 21/765 | 32/2446 | 9,6041E+14 | SCAMP1/STX3/PTPRB/RAB27A/SNAP23/QSOX1/ADAM10/CYBA/JUP/STOM/HMOX2/CLCN3/CKAP4/STX4/PLD1/ANO6/CANT1/CD93, | 21 |
| GO:0009897 | CC | external side of plasma membrane | 14/765 | 18/2446 | 9,89461E+14 | BCAM/ECE1/ABCC4/CLPTM1/TFRC/THBS1/HLA-E/ITGA3/CDH5/MCAM/CLCN3/CXCR4/ADAM9/CLEC14A | 14 |
| GO:0045177 | CC | apical part of cell | 33/765 | 59/2446 | 9,89461E+14 | RAB18/STX3/SLC3A2/MGST1/RAB27A/HAX1/SLC16A3/ABCC4/HIP1R/SLC4A2/ATP1A1/APP/HSP90AB1/SLC2A1/GJA1/SLC9A1/ATP2B | 33 |
| GO:0005765 | CC | lysosomal membrane | 43/765 | 84/2446 | 1,32645E+14 | ATP11C/MAGT1/DDOST/ECE1/TMEM106B/BRI3/SLC3A2/MGST1/CLCN7/NSF/STX7/NDUFC2/RAB3D/SNAP29/HSP90AB1/STOM/GN | 43 |
| GO:0098852 | CC | lytic vacuole membrane | 43/765 | 84/2446 | 1,32645E+14 | ATP11C/MAGT1/DDOST/ECE1/TMEM106B/BRI3/SLC3A2/MGST1/CLCN7/NSF/STX7/NDUFC2/RAB3D/SNAP29/HSP90AB1/STOM/GN | 43 |
| GO:0000323 | CC | lytic vacuole | 62/765 | 134/2446 | 1,9324E+14 | ATP11C/MAGT1/STX3/DDOST/TSC2/TPP1/ECE1/TMEM106B/LRBA/BRI3/SLC3A2/MGST1/CLCN7/RAB27A/NSF/SNAP23/STX7/VAMP | 62 |
| GO:0005764 | CC | lysosome | 62/765 | 134/2446 | 1,9324E+14 | ATP11C/MAGT1/STX3/DDOST/TSC2/TPP1/ECE1/TMEM106B/LRBA/BRI3/SLC3A2/MGST1/CLCN7/RAB27A/NSF/SNAP23/STX7/VAMP | 62 |
| GO:0016323 | CC | basolateral plasma membrane | 19/765 | 29/2446 | 2,07739E+14 | BSG/CTNNB1/SLC3A2/SLC16A3/ABCC4/TFRC/SLC4A2/ATP1A1/HSP90AB1/SLC2A1/SLC9A1/ATP2B1/ATP2B4/ITGA3/ABCC1/SLC16A1, | 19 |
| GO:0031985 | CC | Golgi cisterna | 17/765 | 25/2446 | 2,31311E+14 | GOLIM4/ST6GAL1/BCAP31/GALNT2/GALNT1/TMEM115/TMED2/TMEM87A/GOLGA5/CSGALNACT1/SCFD1/CANT1/YIPF6/COG3/FU | 17 |
| GO:0030670 | CC | phagocytic vesicle membrane | 16/765 | 23/2446 | 2,38061E+14 | TAP1/SNAP23/SEC22B/CYBA/HLA-E/RAB5A/RAB8A/RAB10/STX4/TCIRG1/RAB32/VAMP3/RAB34/RAB22A/PIKFYVE/TAP2 | 16 |
| GO:0017119 | CC | Golgi transport complex | 9/765 | 10/2446 | 2,74079E+14 | COG4/COG8/COG2/COG7/TMEM115/COG1/COG3/COG5/COG6 | 9 |
| GO:0044233 | CC | mitochondria-associated endoplasmic reticulum membrane | 9/765 | 10/2446 | 2,74079E+14 | ACSL4/TOMM40/BCAP31/STX17/RAB32/TMEM41B/MBOAT7/CLCC1/TMX2 | 9 |
| GO:0048471 | CC | perinuclear region of cytoplasm | 72/765 | 163/2446 | 3,15906E+14 | DNAJB6/UNC45A/CTNNB1/TSC2/ECE1/PTPRM/BRI3/STX10/ADAM10/RAB29/STX7/STX6/USO1/ATP9A/HIP1R/ACSL3/TFRC/APP/ITGI | 72 |
| GO:0055037 | CC | recycling endosome | 27/765 | 48/2446 | 3,47061E+14 | ATP11C/SCAMP1/TPP1/STX12/RAB35/SCAMP3/RAB29/STX7/ATP9A/TFRC/APP/ITGB1/HLA-E/SLC1A1/RAB13/CLCN3/RAB8A/RAB10/ | 27 |
| GO:0043235 | CC | receptor complex | 21/765 | 35/2446 | 4,93846E+14 | PTPRB/ACVRL1/TFRC/APP/ITGB1/HSP90AB1/ITGB4/ITGA6/ITGA3/EPHA2/TIE1/FLT4/NOTCH1/TEK/ACVR1/TRADD/CALCRL/ABHD12/ | 21 |
| GO:0098797 | CC | plasma membrane protein complex | 32/765 | 61/2446 | 5,09579E+14 | CTNNB1/COL13A1/SLC3A2/ACVRL1/TFRC/GNAI2/ATP1A1/ITGB1/CYBA/HLA-E/JUP/ITGB4/GJA1/ITGA6/ITGA3/GNA11/CDH5/CTNNA | 32 |
| GO:0032580 | CC | Golgi cisterna membrane | 14/765 | 20/2446 | 5,09579E+14 | GOLIM4/ST6GAL1/BCAP31/GALNT2/GALNT1/TMEM115/TMED2/TMEM87A/CSGALNACT1/SCFD1/CANT1/COG3/FUT8/B4GALT7 | 14 |
| GO:0045178 | CC | basal part of cell | 24/765 | 42/2446 | 5,17781E+14 | BSG/CTNNB1/SLC3A2/SLC16A3/ABCC4/TFRC/SLC4A2/ATP1A1/HSP90AB1/TACSTD2/SLC2A1/ITGB4/SLC9A1/ATP2B1/ATP2B4/ITGA3/ | 24 |
| GO:0005768 | CC | endosome | 83/765 | 196/2446 | 5,32655E+14 | ATP11C/SCAMP1/BSG/TM9SF4/CLN6/GOSR2/TPP1/ECE1/STX12/VPS8/TMEM106B/IRAK1/WLS/RAB35/GOLIM4/RFTN1/GPR107/RA | 83 |
| GO:0031965 | CC | nuclear membrane | 41/765 | 84/2446 | 5,93816E+14 | CEPT1/DPY19L1/TMEM120B/RNF123/EI24/AAAS/NUP93/SMPD4/CTDNEP1/UNC50/HAX1/SLC16A3/ZMPSTE24/ITGB4/DNAJB2/CDH | 41 |
| GO:0005793 | CC | endoplasmic reticulum-Golgi intermediate compartment | 23/765 | 40/2446 | 5,97407E+14 | ROBO1/TAP1/GOSR2/SEC22B/PTPN2/TMED10/BCAP31/STX17/RAB2A/LMAN2/STX5/TMED2/YIF1B/TMED4/PIEZO1/GBF1/ERGIC1/T | 23 |
| GO:0001669 | CC | acrosomal vesicle | 9/765 | 11/2446 | 9,23083E+14 | BSG/VDAC2/VEZT/CD46/CTNNA1/NOTCH1/CAV1/ADAM15/POMT1 | 9 |
| GO:0030173 | CC | integral component of Golgi membrane | 9/765 | 11/2446 | 9,23083E+14 | UNC50/QSOX1/GALNT2/YIF1B/QSOX2/SLC35B2/CSGALNACT1/ERGIC1/TBC1D20 | 9 |
| GO:0031228 | CC | intrinsic component of Golgi membrane | 9/765 | 11/2446 | 9,23083E+14 | UNC50/QSOX1/GALNT2/YIF1B/QSOX2/SLC35B2/CSGALNACT1/ERGIC1/TBC1D20 | 9 |
| GO:0031306 | CC | intrinsic component of mitochondrial outer membrane | 9/765 | 11/2446 | 9,23083E+14 | RHOT1/MFN2/TOMM40/CPT1A/RHOT2/MGARP/FUNDC2/MFF/ARMCX3 | 9 |
| GO:0098799 | CC | outer mitochondrial membrane protein complex | 9/765 | 11/2446 | 9,23083E+14 | APOOL/CHCHD3/CHCHD6/MTX2/TOMM40/MTX1/IMMT/TOMM22/DNAJC11 | 9 |
| GO:0070821 | CC | tertiary granule membrane | 10/765 | 13/2446 | 1,02281E+13 | PTPRB/SNAP23/ADAM10/CYBA/STOM/RAB14/PLD1/ANO6/NBEAL2/CD93 | 10 |
| GO:0030666 | CC | endocytic vesicle membrane | 22/765 | 39/2446 | 1,02281E+13 | TAP1/WLS/SNAP23/SEC22B/CYBA/HLA-E/RAB5A/NOS3/RAB8A/RAB10/CAV1/STX4/TCIRG1/RAB32/SCARF1/VAMP3/TBC1D5/RAB34, | 22 |
| GO:0030139 | CC | endocytic vesicle | 40/765 | 84/2446 | 1,14776E+14 | HYOU1/TAP1/STX12/WLS/SPARC/RAB35/GOLIM4/SNAP23/STX7/STX6/SEC22B/RALA/IGF2R/CYBA/HLA-E/RAB5A/NOS3/RAB5C/RAB | 40 |
| GO:0099023 | CC | vesicle tethering complex | 23/765 | 42/2446 | 1,32174E+14 | COG4/EXOC6B/NBAS/COG8/COG2/VPS8/ZW10/STX17/RAB10/COG7/TMEM115/RINT1/EXOC8/EXOC6/COG1/EXOC4/COG3/EXOC2/ | 23 |
| GO:0005635 | CC | nuclear envelope | 59/765 | 136/2446 | 1,58321E+14 | CEPT1/DPY19L1/TMEM120B/RNF123/EI24/AAAS/NUP93/SMPD4/CTDNEP1/UNC50/HAX1/MGST3/SLC16A3/XPOT/ZMPSTE24/APP/S | 59 |
| GO:1990351 | CC | transporter complex | 14/765 | 22/2446 | 1,7895E+14 | ATP1A1/SLC9A1/TIMM10/PKD2/ANO6/LRRC8D/LRRC8A/ABHD12/MCU/LRRC8C/CLCC1/TTYH3/ABCB8/TMEM30A | 14 |
| GO:0030134 | CC | COPII-coated ER to Golgi transport vesicle | 18/765 | 31/2446 | 1,81245E+14 | GOSR2/USO1/SEC22B/APP/HLA-E/TMED10/STX17/LMAN2/STX5/TMED2/YIF1B/TMED4/ERGIC1/VTI1A/YIPF6/TMED9/CTSZ/VTI1B | 18 |
| GO:0098552 | CC | side of membrane | 33/765 | 68/2446 | 1,97981E+14 | HM13/BCAM/ECE1/ABCC4/CLPTM1/TFRC/GNAI2/S100A6/THBS1/HLA-E/JUP/ATP2A2/PTPN1/ATP2B1/RAB5A/ITGA3/GNA11/CDH5/ | 33 |
| GO:0031307 | CC | integral component of mitochondrial outer membrane | 8/765 | 10/2446 | 2,09218E+14 | RHOT1/TOMM40/CPT1A/RHOT2/MGARP/FUNDC2/MFF/ARMCX3 | 8 |
| GO:0005637 | CC | nuclear inner membrane | 11/765 | 16/2446 | 2,15742E+14 | DPY19L1/TMEM120B/UNC50/ZMPSTE24/LRPPRC/EMD/LBR/LEMD2/TMEM43/SUN2/LEMD3 | 11 |
| GO:0045121 | CC | membrane raft | 30/765 | 61/2446 | 2,2351E+14 | BSG/CLN6/TPP1/STX12/ERLIN2/RFTN1/GNAI2/ATP1A1/APP/ITGB1/HMOX1/SLC2A1/PECAM1/GJA1/HK1/SLC9A1/ATP2B1/RAB5A/A | 30 |
| GO:0098857 | CC | membrane microdomain | 30/765 | 61/2446 | 2,2351E+14 | BSG/CLN6/TPP1/STX12/ERLIN2/RFTN1/GNAI2/ATP1A1/APP/ITGB1/HMOX1/SLC2A1/PECAM1/GJA1/HK1/SLC9A1/ATP2B1/RAB5A/A | 30 |
| GO:0055038 | CC | recycling endosome membrane | 15/765 | 25/2446 | 2,5282E+14 | SCAMP1/STX12/RAB35/SCAMP3/HLA-E/SLC1A1/RAB13/RAB8A/RAB10/RAB14/VAMP3/SCAMP4/EHD4/CMTM6/VTI1B | 15 |
| GO:0045335 | CC | phagocytic vesicle | 22/765 | 42/2446 | 3,01728E+14 | TAP1/STX12/SNAP23/STX6/SEC22B/CYBA/HLA-E/RAB5A/CLCN3/RAB8A/RAB10/RAB14/STX4/TCIRG1/RAB32/VAMP3/RAB34/ZDHHC | 22 |
| GO:0030135 | CC | coated vesicle | 42/765 | 94/2446 | 3,29894E+14 | SCAMP1/GOSR2/ECE1/RAB35/GPR107/RAB27A/HAX1/ADAM10/TGOLN2/STX6/USO1/HIP1R/VAMP4/SEC22B/TFRC/APP/IGF2R/HLA | 42 |
| GO:1902495 | CC | transmembrane transporter complex | 12/765 | 19/2446 | 3,56408E+14 | ATP1A1/SLC9A1/PKD2/ANO6/LRRC8D/LRRC8A/ABHD12/MCU/LRRC8C/CLCC1/TTYH3/ABCB8 | 12 |
| GO:0098590 | CC | plasma membrane region | 70/765 | 172/2446 | 3,74559E+14 | BSG/RAB18/STX3/CTNNB1/COL13A1/WLS/SLC3A2/RAB35/VEZT/RAB27A/ICAM2/HAX1/ADAM10/SLC16A3/ABCC4/HIP1R/SNAP29/T | 70 |
| GO:0005766 | CC | primary lysosome | 20/765 | 38/2446 | 4,01595E+14 | MAGT1/STX3/DDOST/BRI3/MGST1/SNAP23/STX7/NDUFC2/RAB3D/SNAP29/CYB5R3/PYGB/STOM/RAB5C/CKAP4/LPCAT1/FAF2/TMI | 20 |
| GO:0042582 | CC | azurophil granule | 20/765 | 38/2446 | 4,01595E+14 | MAGT1/STX3/DDOST/BRI3/MGST1/SNAP23/STX7/NDUFC2/RAB3D/SNAP29/CYB5R3/PYGB/STOM/RAB5C/CKAP4/LPCAT1/FAF2/TMI | 20 |

| | | | | | | | |
|---|---|---|---|---|---|---|---|
| GO:0042470 | CC | melanosome | 24/765 | 48/2446 | 4,10258E+14 | BSG/STX3/TPP1/SLC3A2/RAB35/RAB27A/RAB29/SEC22B/TFRC/RPN1/ATP1A1/ITGB1/HSP90AB1/CNP/SLC2A1/RAB5A/STOM/TMED1 | 24 |
| GO:0048770 | CC | pigment granule | 24/765 | 48/2446 | 4,10258E+14 | BSG/STX3/TPP1/SLC3A2/RAB35/RAB27A/RAB29/SEC22B/TFRC/RPN1/ATP1A1/ITGB1/HSP90AB1/CNP/SLC2A1/RAB5A/STOM/TMED1 | 24 |
| GO:0005778 | CC | peroxisomal membrane | 14/765 | 24/2446 | 4,41353E+14 | PEX1/PEX5/ACBD5/ACSL1/MGST1/ALDH3A2/ACSL4/ACSL3/ABCD3/ABCD1/MAP2K2/PEX6/MAVS/ATAD1 | 14 |
| GO:0031903 | CC | microbody membrane | 14/765 | 24/2446 | 4,41353E+14 | PEX1/PEX5/ACBD5/ACSL1/MGST1/ALDH3A2/ACSL4/ACSL3/ABCD3/ABCD1/MAP2K2/PEX6/MAVS/ATAD1 | 14 |
| GO:0030672 | CC | synaptic vesicle membrane | 8/765 | 11/2446 | 4,53968E+14 | SCAMP1/STX12/RAB35/STX10/STX6/VAMP4/RAB5A/VAMP3 | 8 |
| GO:0099501 | CC | exocytic vesicle membrane | 8/765 | 11/2446 | 4,53968E+14 | SCAMP1/STX12/RAB35/STX10/STX6/VAMP4/RAB5A/VAMP3 | 8 |
| GO:0005769 | CC | early endosome | 42/765 | 96/2446 | 4,68937E+14 | TM9SF4/CLN6/STX12/VPS8/WLS/RFTN1/GPR107/MGRN1/RAB29/STX7/STX6/ATP9A/SNX4/TFRC/APP/IGF2R/HLA-E/GJA1/PTPN1/RA | 42 |

### Table S5. Differentially regulated proteins in shSEC22B and shSTX5 endothelial cells.

| shCTRL_A | shCTRL_B | shCTRL_C | shSEC22B_159152_A | shSEC22B_159152_B | shSEC22B_159152_C | shSEC22B_159288_A | shSEC22B_159288_B | shSEC22B_159288_C | shSTX5_59826_A | shSTX5_59826_B | shSTX5_59826_C | shSTX5_59827_A | shSTX5_59827_B | shSTX5_59827_C | Untransduced_A | Untransduced_B | Untransduced_C | Gene names | Correlation |
|---|---|---|---|---|---|---|---|---|---|---|---|---|---|---|---|---|---|---|---|
| 26,70754092 | 26,88750902 | 27,12119549 | NA | NA | NA | NA | NA | NA | NA | NA | NA | NA | 26,26815546 | 26,50738007 | 27,31311152 | 27,05248032 | 27,00560296 | COL12A1 | 3 |
| NA | 25,82301206 | NA | 26,28963071 | 26,34033137 | 26,70569671 | NA | 26,29045709 | 26,4103301 | 26,62282355 | 26,77492436 | 26,81127434 | 26,2950025 | 26,27765674 | 26,33568829 | 25,79109093 | 25,77256044 | 25,57039542 | NOL4L | 2 |
| NA | NA | NA | NA | 25,8499548 | NA | 28,73075304 | 28,73062349 | 28,78338006 | 27,14976851 | 27,78706001 | 26,16731477 | 27,21660082 | 26,72871128 | 26,57502075 | NA | NA | NA | COL6A3 | 2 |
| 26,20235912 | 26,19964734 | 26,10837003 | 25,1226574 | NA | 25,33459764 | NA | 25,07361434 | NA | 25,73704161 | 26,23870561 | 25,69726243 | 26,23673656 | 26,33778208 | 26,19341903 | 27,16086808 | 27,29386311 | 26,88599755 | STAU2 | 3 |
| 28,34870378 | 28,4703783 | 28,48515947 | NA | NA | NA | NA | NA | NA | 24,30612123 | 25,64696778 | 25,17230267 | 26,34382507 | 26,13082866 | 25,95659736 | 25,71298074 | NA | 26,06058363 | ANGPT2 | 3 |
| NA | NA | NA | NA | 25,87321485 | NA | NA | NA | NA | 25,28598547 | 25,63541679 | 25,89915221 | 26,10535562 | 25,97493495 | 26,24019884 | NA | 25,23611612 | NA | PDPK1;PDPK2P | 4 |
| 25,64839489 | 26,49596198 | 26,10860932 | 25,30949051 | NA | 25,56508697 | 25,38162506 | 24,90504209 | NA | NA | NA | NA | 25,58580359 | NA | NA | 25,57441452 | 25,59089378 | 25,83093758 | MTFR1L | 1 |
| NA | NA | NA | 25,83004299 | 25,25400152 | 25,86057837 | 25,80855201 | 25,70701424 | 25,60481403 | 26,01416472 | 25,87743374 | 26,40642639 | NA | NA | 25,07418622 | NA | NA | 25,58106907 | EIF1AD | 2 |
| 30,42802293 | 30,37160717 | 30,40334538 | 31,96612683 | 32,15587178 | 31,95750402 | 31,81459364 | 31,82209886 | 31,70775483 | 31,62583939 | 31,68633083 | 31,70148904 | 31,00564312 | 31,07067221 | 31,11153984 | 31,36833216 | 31,21974254 | 31,3516867 | NPEPPS | 2 |
| NA | NA | NA | 25,20445025 | 25,31267886 | 25,38221898 | 25,11890085 | 25,01852213 | 25,21793226 | 24,91725115 | 24,78136772 | 25,03523405 | 25,13363361 | 25,01792775 | 25,03724665 | 24,63173082 | 25,6901649 | NA | IFITM2;IFITM3;IFITM1 | 2 |
| 25,04335109 | 25,16959109 | 25,04339281 | NA | NA | NA | NA | NA | NA | NA | NA | NA | 24,64795593 | 24,64762662 | 24,88455437 | NA | 24,61745267 | 24,06907209 | ICOSLG | 3 |
| 27,34053512 | 27,24717103 | 27,51209096 | 27,24445152 | 27,47351699 | 27,09727956 | 26,48319225 | NA | 26,78144276 | NA | NA | 26,6499302 | NA | NA | NA | NA | NA | 26,75506319 | CD63 | 1 |
| 24,7280625 | 25,18744692 | 24,92261416 | NA | NA | NA | 25,53219515 | 25,85293349 | 25,46600375 | 26,62839703 | 26,67720928 | 26,17014528 | 26,0371838 | 26,0981836 | 26,08573859 | 25,6975806 | 25,60087886 | 25,26931534 | LMO7 | 4 |
| NA | NA | NA | 25,31996478 | 25,48830452 | 25,47410522 | 25,75325328 | 25,52650521 | 25,45923323 | NA | NA | NA | NA | NA | NA | 24,59696134 | NA | NA | SSSCA1 | 2 |
| 24,38922852 | 24,68560089 | 24,83537813 | 26,18374198 | 26,39188521 | 26,2807215 | 26,88378563 | 26,64965615 | 26,07306267 | 24,79585378 | 25,23187537 | 25,40008672 | 25,41112251 | 25,79178649 | 25,52912954 | NA | NA | NA | TTLL5 | 2 |
| 26,32890949 | 26,36668039 | 26,52169149 | NA | NA | NA | 27,4540324 | 27,2654395 | 27,57195813 | NA | NA | NA | 24,94837686 | NA | NA | 26,8062426 | 26,63159207 | 26,57369249 | LTBP2 | 1 |
| 25,17915435 | 25,17314184 | NA | 25,28764193 | 25,23534932 | 25,10129318 | 26,79773474 | 26,29848523 | 26,92241001 | 25,97381891 | 25,88040299 | 25,80653763 | 25,52163159 | 25,82488224 | 25,83708795 | 26,12561621 | 26,18504736 | 26,3215311 | ZNF512 | 2 |
| 31,73351938 | 31,73194273 | 31,68995837 | 29,93170945 | 29,97978295 | 29,97951041 | 30,7008279 | 30,69004165 | 30,54802552 | 29,21745623 | 29,3011223 | 29,18293262 | 31,83575157 | 31,96481948 | 32,03416765 | 31,19724221 | 31,02828372 | 31,00068263 | LAMB1 | 3 |
| NA | 23,58920857 | NA | 25,14020288 | NA | 24,89559747 | NA | 24,88683487 | NA | 24,62109139 | 25,16430606 | 25,0038459 | 24,94748534 | 24,74514456 | 24,20426366 | 24,10269532 | 24,85171893 | 24,6428432 | SOS1 | 2 |
| 25,95708468 | 26,37754351 | 26,17985671 | 25,11735584 | 24,88030957 | NA | 25,53255188 | NA | NA | NA | NA | NA | 24,96499157 | 25,22000096 | 24,92596727 | 25,95690749 | NA | 25,542859 | NEDD4 | 3 |
| 25,97633424 | 26,12687679 | 25,7285037 | 24,9693022 | 24,31699961 | 24,57282557 | 24,24365283 | 24,21623075 | 24,21504591 | NA | NA | 25,39345723 | 25,59212075 | 25,87963204 | 25,32831034 | 25,64586905 | 25,33237971 | 25,80358466 | CD99 | 3 |
| NA | NA | NA | NA | NA | NA | NA | NA | NA | NA | NA | NA | 27,28964829 | 27,1814879 | 27,24417929 | NA | NA | NA | MYCT1 | 4 |
| 32,06059394 | 31,88489202 | 32,08714579 | 29,3873273 | 29,12168954 | 29,58953006 | 30,37513378 | 30,52495969 | 30,34857716 | 29,64368282 | 29,75762442 | 29,65993129 | 32,41782539 | 32,28195531 | 32,28711238 | 31,02075419 | 30,90196211 | 30,97151573 | LAMA4 | 3 |
| 26,11036287 | 25,79832822 | NA | NA | NA | NA | NA | NA | NA | NA | NA | NA | NA | NA | NA | 25,9659824 | 26,49713528 | 26,3684633 | RBP1 | 3 |
| 25,14464362 | 25,12273638 | 25,55200836 | NA | NA | NA | NA | NA | NA | NA | 24,97823424 | NA | NA | 24,79709153 | NA | 25,18381769 | NA | 24,84186951 | SCLY | 3 |
| 26,07614524 | 26,14693654 | 26,19053968 | NA | NA | NA | NA | 25,02749299 | NA | 25,47395045 | 25,25789019 | 25,34499341 | 26,45309049 | 26,29307377 | 26,12599056 | 26,26703039 | 26,22624445 | 26,26137388 | MLLT4 | 3 |
| 26,74226872 | NA | NA | 27,30365005 | 27,29017582 | 27,08547521 | 27,76597641 | 27,5241751 | 27,50180378 | 26,84840345 | NA | 27,52065788 | NA | NA | NA | 27,13633532 | 27,14899318 | 26,69232178 | MIEN1 | 2 |
| 26,39875021 | 26,81714293 | 26,70173689 | NA | NA | NA | 25,13296719 | NA | NA | 25,03619877 | NA | NA | 26,6896585 | 26,50668421 | 26,48088349 | 26,36223855 | 26,74534976 | 26,59291914 | CDC42BPA | 3 |
| 26,15514558 | 26,08867293 | 26,34243539 | NA | NA | 25,23110631 | 24,76410419 | 24,3789494 | NA | 25,04113808 | 25,32639136 | 25,18449887 | 25,06254061 | 25,51144262 | 25,52208077 | 25,67441018 | 25,37311845 | 25,80503729 | FSD1 | 3 |
| 29,29653489 | 29,24204503 | 29,22267018 | 27,93100491 | 27,81017131 | 28,11137816 | 28,09073352 | 27,82183285 | 27,97207749 | 27,9780378 | 28,69025817 | 28,3550627 | 29,12620232 | 29,12067654 | 29,41052016 | 28,9379227 | 29,20673862 | 28,78016021 | ATOX1 | 3 |
| NA | 25,19163187 | NA | 26,91193667 | 27,13310442 | 26,6486966 | 27,58885129 | 28,00056176 | 27,51773282 | 26,11300909 | 26,26867306 | 26,57960252 | NA | NA | NA | NA | NA | NA | QSOX1 | 2 |
| 25,5524775 | NA | NA | 26,57960252 | 26,47201437 | 26,84541539 | 26,46199832 | 27,12524176 | 26,51273902 | 27,12868695 | 26,94937915 | 26,81470058 | 26,3968411 | 26,41902276 | 26,35845388 | 25,68161225 | 25,80331367 | 25,77296309 | SLC33A1 | 2 |
| 25,4271678 | 25,64262289 | 25,33292598 | NA | NA | NA | NA | NA | NA | NA | NA | NA | NA | 24,74924299 | 25,32961105 | 25,75072579 | 25,63599792 | 26,03615684 | EEF2K | 3 |
| 30,2724201 | 30,36191657 | 30,22054996 | 30,94118727 | 30,89113589 | 31,02207801 | 29,53473125 | 29,35205518 | 29,49594674 | 31,62675335 | 31,56696928 | 31,72024612 | 32,09282645 | 32,05694975 | 32,14961711 | 31,08853646 | 31,12896958 | 31,16883829 | IGF2BP3 | 4 |
| 25,29263506 | NA | 25,47707367 | 26,90138676 | 26,71135357 | 26,85174275 | 27,18876781 | 27,28859266 | 26,65526364 | 27,00517461 | 27,26445549 | 26,89642923 | 27,30286605 | 27,46699948 | 27,29911437 | 27,67485463 | 27,92223985 | 27,68618888 | BIN1 | 2 |
| NA | NA | NA | 26,69723591 | 26,08586013 | 26,29572074 | 25,08342739 | 25,79857543 | 24,72899665 | 25,32333629 | 25,97548171 | 25,54904338 | 25,35974447 | 24,99459686 | 25,85986782 | NA | NA | NA | VWA5A | 2 |
| 28,04072015 | 27,71423923 | 27,69849495 | NA | 25,70790948 | 26,04859862 | 27,16298178 | 26,13662868 | NA | 28,44407202 | 28,05211751 | 28,48577367 | 27,72351284 | 27,2164158 | 27,77052034 | 28,29469583 | 28,21771044 | 28,35068605 | SNRPC | 3 |
| NA | 25,04410191 | 24,91556541 | NA | NA | NA | 26,00868336 | 25,45854504 | NA | 26,60286194 | 26,59405893 | 26,7655971 | 25,92583148 | 25,29726111 | 26,13269269 | 26,00223696 | 25,61175225 | 25,97373134 | IRF6 | 2 |
| 25,40509575 | 24,67890226 | 25,50651776 | 27,7365385 | 27,93635083 | 27,87003081 | 28,12652237 | 28,43025824 | 28,06763688 | 27,76350913 | 27,9636143 | 28,15836607 | 26,91147958 | 26,60399391 | 26,45074875 | 26,00191496 | 24,47747518 | 26,06728811 | PLXNB2 | 2 |
| 26,43264142 | 26,3948477 | 25,66133884 | 25,43096803 | 25,81984911 | 25,75608187 | NA | NA | NA | NA | NA | NA | 24,40080336 | NA | NA | 25,94937915 | 26,04693481 | 25,7464522 | SYNM | 1 |
| NA | NA | NA | 27,48223841 | 26,82505213 | 27,15990628 | 27,08669037 | 27,10199442 | 26,46054614 | 25,92619355 | NA | NA | NA | NA | NA | NA | NA | 26,79004696 | LSM1 | 2 |
| 29,33287478 | 29,22953946 | 29,71337086 | 30,81596807 | 30,45032006 | 30,49018061 | 30,5995671 | 30,77462594 | 30,71947113 | 29,91456214 | 30,09055687 | 30,13050466 | 30,07495178 | 30,18364261 | 30,17819042 | 30,53296796 | 30,02194568 | 30,33031226 | ARPC3 | 2 |
| NA | NA | NA | 26,90907747 | 26,92524293 | 27,19215884 | 27,00378158 | 26,67006621 | 26,54755861 | 26,62839703 | 26,47068084 | 26,6063964 | 25,98235321 | 26,21467545 | 26,46468023 | 26,6434764 | 26,60766669 | 26,603711 | ZZEF1 | 2 |
| 26,46078046 | 26,58462859 | 26,41998717 | NA | NA | 25,74288546 | NA | NA | NA | 25,45992109 | NA | NA | 27,13369239 | 26,81677684 | 27,25069873 | NA | NA | 25,30619078 | SLC4A7 | 3 |
| 25,36359259 | 25,48726204 | 25,37471104 | 25,76759367 | 25,35444068 | 25,47385758 | 25,40447869 | 25,02791477 | 25,15093073 | NA | NA | 24,60036848 | 24,49692203 | 24,9685993 | NA | 24,92474474 | 25,04789173 | 24,9812429 | PIP5K1C | 1 |
| NA | NA | NA | NA | NA | NA | 25,40979619 | 25,72184538 | 25,61857325 | 25,66673869 | 25,94957953 | 25,80950909 | 26,26572563 | 26,08786405 | 26,2935825 | 26,30985475 | 26,58391165 | 26,38853895 | TBC1D4 | 2 |
| 25,6500124 | 25,9369461 | 25,83968493 | NA | 25,36125148 | NA | NA | NA | NA | 25,37716286 | NA | NA | 26,36396014 | 26,50430669 | 26,09866552 | 25,94147555 | 25,91179956 | 25,52737057 | DOCK9 | 3 |
| 25,33412022 | 25,18888098 | NA | 28,37442915 | 28,69272084 | 28,39791019 | 27,35396978 | 27,4308564 | 27,39132804 | 25,49390259 | 27,00142108 | 26,79041989 | 27,04113808 | 27,34401139 | 27,47753694 | 26,06260237 | 26,73866584 | 26,95803668 | NRP2 | 2 |
| 26,33548386 | 26,02774607 | 26,46780801 | 27,27666329 | 27,25457828 | 27,25394744 | 27,34375732 | 27,50301785 | 27,39974456 | 27,2793228 | 27,81799678 | 27,58956577 | 27,47343958 | 27,18158268 | 27,36636357 | 27,20853043 | 27,17169205 | 27,33330142 | NEMF | 2 |
| NA | 27,01286558 | NA | 28,08156966 | 28,55738698 | 28,37736975 | 28,02543505 | 28,42406272 | 27,87197027 | 27,48316149 | 27,22306162 | 27,57217504 | 27,67107918 | 27,50407933 | 27,31674073 | 26,35077034 | NA | 26,54489394 | PRAF2 | 2 |
| 24,39185244 | 24,64921758 | NA | 25,76817443 | 25,95251518 | 26,02780934 | 25,77210733 | 25,74783547 | 26,10795119 | 26,10281544 | 26,08302153 | 26,13891484 | 24,93678888 | 25,20873506 | 25,11525357 | 25,98997166 | 25,69763362 | 25,88778789 | BRD4 | 2 |
| 30,85549025 | 31,06618741 | 30,75771978 | 31,43966044 | 31,27169684 | 31,36416266 | 30,62121718 | 30,45454235 | 30,49667065 | 29,60015222 | 29,57434233 | 29,42564411 | 29,21045936 | 29,23735674 | 29,36959106 | 30,16734349 | 30,11458357 | 30,24547194 | SH3BGRL | 1 |
| 30,03773889 | 30,0889155 | 30,09358218 | 28,05004251 | 27,83376277 | 27,95858988 | 28,00083035 | 27,7518752 | 27,61736859 | 29,77541113 | 29,81596807 | 29,95088129 | 30,13723965 | 30,07418622 | 29,95449116 | 29,95670813 | 29,87597317 | 30,01916401 | SEC22B | 3 |
| NA | 25,91825254 | NA | 25,46145197 | 25,9514706 | 25,82614384 | 26,37471104 | 25,48833517 | 25,42934059 | 25,9978946 | 26,63173082 | 26,06000633 | NA | NA | NA | 23,93093725 | NA | NA | GIGYF1 | 2 |
| 26,49408577 | 26,60399391 | 26,57556902 | NA | NA | NA | NA | NA | NA | 27,61210391 | 27,91923051 | 27,76186193 | 26,23491096 | 26,56248362 | 26,03019025 | 25,39940232 | 26,21432342 | 26,23693723 | KHDRBS3 | 3 |
| 26,84170158 | 26,55651751 | 26,89781444 | 25,49771396 | 25,88126691 | 25,90788638 | 25,28785326 | NA | NA | 25,5088161 | 25,84009329 | 25,77836273 | 26,54518862 | 26,76357245 | 26,73582868 | 25,49548934 | 26,09946837 | 26,12405866 | PALM | 3 |
| 26,66261601 | 26,42447925 | 26,75951459 | 28,09898672 | 28,07183598 | 28,10559541 | 26,211764 | 26,45614946 | 26,01047723 | 28,42546404 | 28,40576122 | 28,28140266 | 28,81203983 | 28,26338125 | 28,65829763 | 26,71685755 | 26,44733147 | 26,56293482 | STK10 | 4 |
| 27,80501268 | 27,80519724 | 27,72325243 | 29,51990843 | 29,64018692 | 29,54826443 | 28,6836481 | 28,53036549 | 28,47332345 | 29,51996465 | 29,71728258 | 29,75776746 | 29,08324477 | 29,0356693 | 29,01286558 | 28,45755911 | 28,76856568 | 28,85409351 | NCEH1 | 2 |
| 28,9643031 | 28,90339948 | 28,59985083 | NA | NA | NA | 26,11457861 | 26,20267672 | NA | 26,99194026 | 27,27293172 | 26,92388382 | 26,33166242 | 26,31241921 | 25,45356153 | 27,45159787 | 27,59903497 | 26,90058088 | SORBS2 | 3 |
| 26,01869191 | 25,83270095 | 26,4143362 | NA | NA | 25,3414686 | NA | NA | NA | NA | NA | NA | 25,21400837 | NA | NA | 25,51095994 | 25,7164651 | 25,55420614 | NDUFA3 | 3 |
| 26,86702832 | 26,92965122 | 26,89284461 | 25,35991199 | 25,57925723 | 25,23950705 | 25,24343493 | 25,13887579 | 25,07879385 | 25,50470069 | 25,67962721 | 25,60628343 | 26,35863832 | 26,00770006 | 26,30943847 | 26,05633094 | 26,70648738 | 26,29049224 | NDUFB8 | 3 |
| 27,72194965 | 27,44994634 | 27,39124608 | NA | NA | NA | NA | NA | NA | 25,83932452 | NA | 26,20202276 | 27,11505508 | 26,97650906 | 26,83400398 | 27,77379318 | 27,84128168 | 27,68739085 | VAMP5 | 3 |
| 28,96826421 | 29,01147997 | 28,84664125 | 27,1379969 | 27,25781827 | 26,46243525 | 27,05393069 | 26,59405893 | 27,49289471 | 28,22742932 | 28,34608469 | 28,22632715 | 28,50419301 | 28,68875854 | 28,62791021 | 28,93190666 | 29,01315852 | 28,56772188 | PAPSS2 | 3 |
| NA | 25,03305057 | 25,27351875 | 26,93280785 | 26,93077938 | 27,09113721 | 26,28153532 | 26,28490951 | 26,5639095 | 26,03080064 | 26,25046382 | 25,87904772 | 26,34974165 | 26,37784134 | 26,46777692 | 24,99451055 | 25,31312882 | 25,15112434 | SGPL1 | 2 |
| 27,23224144 | 27,18177223 | 27,18215123 | 25,82070135 | 25,6082027 | 25,68517311 | 25,57827847 | NA | NA | 25,55121634 | 25,46534992 | NA | 26,65812735 | 26,36059863 | 26,41024922 | 26,68418337 | 26,89816054 | 26,69564397 | RAB3D | 3 |
| NA | NA | NA | 26,56759859 | 26,62728406 | 26,76900728 | 26,54277046 | 26,00975141 | NA | NA | NA | NA | 26,55878141 | 26,43370819 | NA | NA | NA | NA | PTBP3 | 2 |
| 26,36140209 | 26,04255814 | 26,19330622 | 26,68899192 | 26,90184706 | 26,32943997 | 27,43053741 | 26,9446958 | 27,0964755 | 27,41021686 | 27,4196497 | 27,73285659 | 27,34053512 | 27,26552892 | 27,4157067 | 27,08760107 | 26,78755827 | 26,98934374 | PEX11B | 2 |
| 35,47311898 | 35,10968873 | 35,34314576 | 33,92768893 | 33,50403196 | 33,59890191 | 36,90477264 | 36,94251184 | 37,17849469 | 34,10937738 | 33,64720888 | 33,79546444 | 33,65405594 | 33,27169684 | 33,24799153 | 35,1943608 | 35,14910981 | 35,08079967 | FN1 | 1 |
| 29,83421501 | 29,73280808 | 30,035512 | 25,17470446 | 26,10892831 | 25,0847659 | 26,03065333 | 25,53165988 | 26,02295106 | 28,31207293 | 28,38063465 | 28,4094887 | 29,38597069 | 29,07311376 | 29,38894945 | 29,26420938 | 29,61407164 | 29,19608154 | TFRC | 3 |
| 29,39065177 | 29,57535262 | 29,49300927 | 29,20263468 | 29,17776286 | 29,03456781 | 28,10529567 | 28,25435301 | 28,30029327 | 30,57235577 | 30,69029148 | 30,6192067 | 30,93880613 | 30,96842902 | 30,88467372 | 27,92320379 | 27,89775675 | 27,81268253 | MMP1 | 4 |
| 35,41808938 | 35,35635816 | 35,36169439 | 32,75787871 | 32,72419621 | 32,64254715 | 33,52647629 | 33,44418559 | 33,68217089 | 32,66250566 | 32,47961981 | 32,59705994 | 33,34942108 | 33,58127383 | 33,58138608 | 35,30198354 | 35,26027855 | 35,27648859 | VWF | 3 |
| 33,58542128 | 33,3353795 | 33,8202167 | 28,71315778 | 28,58401921 | 28,84628257 | 33,03467603 | 32,89008504 | 33,08679791 | 30,84183653 | 30,41588799 | 30,49980973 | 31,52439931 | 31,50847801 | 31,60262305 | 33,17328365 | 33,13781984 | 33,1682854 | THBS1 | 3 |
| 31,75585114 | 31,83492324 | 31,44997585 | 29,48275771 | 29,2657748 | 29,54333113 | 30,20314832 | 30,49609917 | 30,18742332 | 29,42728372 | 29,4349173 | 29,4617798 | 31,64873088 | 31,78019151 | 31,78460516 | 30,48941515 | 30,64581409 | 30,58190232 | SPARC | 3 |
| 26,77366744 | 26,73505394 | 26,4696566 | 29,66227645 | 29,93480536 | 29,63991102 | 28,69604212 | 29,54336801 | 29,32223196 | 32,10533938 | 32,18393834 | 32,26150502 | 27,2950025 | 26,74252573 | 26,49439101 | 30,05736946 | 29,97404876 | 29,86792678 | HMOX1 | 2 |
| 25,91754709 | 25,37607 | 25,55496725 | NA | NA | NA | NA | NA | NA | NA | 24,60783598 | NA | NA | NA | NA | 25,52886136 | 25,02808345 | 25,24796777 | TACSTD2 | 1 |
| 32,65731825 | 32,52318441 | 32,51451026 | 31,1016188 | 30,93368035 | 30,97240622 | 31,19876491 | 31,46372965 | 31,12028103 | 31,87000875 | 31,84404629 | 31,87070697 | 31,73202362 | 31,61333406 | 31,64710163 | 32,02785546 | 32,33306145 | 32,25141001 | UCHL1 | 3 |
| 32,25363755 | 32,1102023 | 32,15665548 | 30,53361784 | 30,59912368 | 30,53602914 | 30,83416979 | 30,90289657 | 30,88845288 | 30,3691126 | 30,41185389 | 30,30443363 | 32,50094953 | 32,51170934 | 32,50559437 | 31,39825686 | 31,22773915 | 31,28322561 | LAMC1 | 3 |
| 29,48924287 | 29,5113559 | 29,642375 | 26,27730202 | 26,07130408 | NA | NA | NA | NA | 26,34774963 | 25,61495624 | 26,5803216 | 30,0930784 | 30,07418622 | 30,02379715 | 30,36640526 | 30,74278269 | 30,56019637 | ACE | 3 |
| NA | NA | NA | NA | NA | NA | 26,90046571 | 27,42965984 | 27,38376862 | 27,2665122 | 27,44908049 | 27,48431451 | NA | 25,72249696 | 26,01145864 | NA | NA | 26,42273221 | MTHFD2 | 2 |
| 31,15206782 | 30,97923782 | 31,17486158 | 28,21752556 | 27,76490145 | 28,01674887 | 26,12553739 | 27,98065474 | 27,48362281 | 27,67982852 | 28,26999298 | 28,64797651 | 30,51483942 | 30,48989361 | 30,64572821 | 30,37129558 | 30,15822159 | 30,14843566 | NID1 | 3 |
| 29,81688363 | 29,74632726 | 29,78195859 | 28,90592579 | 28,63273639 | 28,38092359 | 28,65095739 | 28,65819546 | 28,45281565 | 28,44162061 | 28,49720382 | 28,37794888 | 29,7637149 | 29,6842001 | 29,79087039 | 29,7692911 | 29,42574414 | 29,73992144 | JUP | 3 |
| 25,88310974 | 25,77386862 | 26,07632877 | 25,88683487 | 26,07742941 | 25,77484898 | 27,44007645 | 26,82310927 | 26,81225409 | 24,29712114 | NA | 24,68056642 | NA | NA | NA | 26,5927766 | 26,74265421 | 26,28714872 | PYCR2 | 1 |
| 30,57452279 | 30,15605267 | 30,35520976 | 25,04922206 | NA | NA | NA | NA | NA | 27,36711383 | 26,61062635 | 26,81066165 | 30,85675209 | 31,02075419 | 30,50512109 | 30,57109017 | 31,06670066 | 30,7713235 | FABP4 | 3 |
| NA | NA | NA | 26,22809939 | 26,53741844 | 26,32445265 | NA | 25,9726582 | 25,79736369 | 25,54698487 | 25,95883323 | 25,21537925 | 24,9364744 | 25,14215225 | 25,10197439 | 27,39901109 | 27,42854215 | 27,06136674 | RPA2 | 2 |
| 29,88483381 | 30,01438291 | 29,9294537 | 28,60438282 | 28,92019646 | 28,75177945 | 27,65095739 | 27,79686881 | 27,97306346 | 30,10686861 | 30,11147766 | 30,22400524 | 30,01704103 | 29,81029391 | 29,68436733 | 29,34633836 | 29,37606586 | 29,38442753 | ITGA2 | 3 |
| 26,17611318 | 26,59149313 | NA | NA | NA | NA | NA | NA | NA | NA | NA | NA | 27,29631608 | 27,0986856 | 27,50294201 | 26,03900537 | NA | NA | IGFBP2 | 3 |
| 26,95803668 | 26,97169387 | 26,96433064 | 25,3069556 | NA | 24,89924443 | 25,70635563 | 25,21300719 | 25,4508903 | 26,14096345 | 25,91499546 | 25,82820371 | 26,1967804 | 26,4274236 | 26,35969421 | 27,07490076 | 26,54589561 | 26,70635563 | AGA | 3 |
| 28,06686743 | 27,78044185 | 27,92212641 | 26,53506139 | 26,82262315 | 26,31342295 | 28,29824049 | 27,91090802 | 28,18517972 | 26,76103762 | 26,53330965 | 26,44512248 | 26,8595835 | 26,79600237 | 26,62058811 | 27,43205197 | 28,01105335 | 27,66044143 | MAOA | 1 |
| 29,11073124 | 28,92334549 | 29,03480392 | NA | NA | NA | 28,52622908 | 28,04004075 | 28,27697382 | NA | NA | NA | 27,79532122 | 27,92167253 | 27,79655943 | 29,16240562 | 29,55870114 | 29,22333787 | BGN | 3 |
| 27,9281268 | 28,18584136 | 27,90850495 | 25,95945246 | 26,41475555 | 26,17495204 | 26,13676556 | 25,69147006 | NA | 27,91934419 | 27,81268253 | 27,51269381 | 26,90919195 | 27,09577157 | 26,29305622 | 28,42270014 | 27,8660261 | 27,91364948 | SDHB | 3 |
| 26,86242416 | 26,64581409 | 26,61708829 | 27,77536403 | 27,89937121 | 27,81090675 | 28,03322915 | 28,00608471 | 27,67835159 | 27,46295003 | 27,71934056 | 27,65184703 | 26,85293349 | 26,89145462 | 26,80906744 | 26,86242416 | 27,23617087 | 26,87244005 | MUT | 2 |
| 26,89284461 | 26,89284461 | 26,56411308 | 25,07614524 | NA | NA | NA | NA | NA | 26,34942108 | 26,35892332 | 26,34438396 | 28,17107163 | 27,59555355 | 27,79884731 | 28,23753909 | 27,8945224 | 27,73666752 | ITGA6 | 3 |
| 30,34857716 | 30,30519503 | 30,36650949 | 28,57762319 | 28,40944824 | 28,50627562 | 26,9375299 | 27,65533189 | 27,68518648 | 29,84297554 | 29,87428832 | 29,83199767 | 30,23637619 | 30,10249509 | 30,0555606 | 29,54907277 | 29,71392799 | 29,59930106 | ATP2B4 | 3 |
| 24,03800065 | NA | NA | 25,68731075 | 26,03363875 | 25,80963175 | 24,69386423 | 25,48732338 | 25,36074932 | 24,42636816 | 25,05622756 | 24,82522201 | NA | NA | 24,38225197 | NA | NA | NA | KDELR1;KDELR2 | 2 |
| NA | NA | NA | 27,62470701 | 27,8803446 | 27,90964977 | 25,86735831 | 26,2168783 | 26,41438459 | 26,69657282 | 25,91816154 | 26,17178748 | 26,51070345 | NA | 26,11862366 | 24,38561282 | NA | NA | FAS | 2 |

| | | | | | | | | | | | | | | | | | | | |
|---|---|---|---|---|---|---|---|---|---|---|---|---|---|---|---|---|---|---|---|
| 28,70833057 | 28,77099284 | 28,8819496 | 27,62331209 | 27,61960902 | 27,6921222 | 25,79872373 | 26,55398651 | 25,55584496 | 26,07508444 | 25,99767929 | 25,54813212 | 28,43718166 | 28,30020598 | 28,17940117 | 26,41028157 | 26,111 | 25,96492549 | AK4 | 3 |
| 28,81231531 | 28,4909839 | 28,36765543 | 28,08542456 | 26,57022169 | 27,02104553 | 27,03611491 | 27,3786518 | 27,59704662 | 26,98130823 | 26,92059401 | 27,32817335 | 26,34646517 | 26,40905163 | 26,5085139 | 28,67033641 | 28,40944824 | 28,58373236 | GRN | 1 |
| NA | NA | NA | 26,31480629 | 26,58477193 | 26,07979091 | 26,20306895 | 26,13961755 | 25,89010969 | 26,37456181 | 26,21756254 | 26,54574835 | NA | NA | NA | 26,14376824 | 26,24119953 | NA | GTF2E1 | 2 |
| 28,51054501 | 28,78012892 | 28,49876413 | 29,8483139 | 29,91541724 | 29,93382102 | 30,20862345 | 30,11768281 | 30,25282024 | 29,95282619 | 29,95629271 | 30,05969181 | 28,05568988 | 27,77950286 | 27,82468805 | 28,17331343 | 28,15339733 | 28,14850839 | EPHA2 | 2 |
| 25,90667136 | 26,1200684 | 26,08484698 | NA | NA | NA | NA | NA | 24,6159385 | 26,04017666 | 25,33183324 | 25,29530035 | 25,29291585 | 25,59192108 | 25,12652237 | 25,46232603 | 25,41231839 | 25,2649923 | EPHB2 | 3 |
| 25,17329437 | 24,92270489 | 24,75128784 | NA | NA | NA | NA | NA | NA | NA | 24,52880176 | 24,86031788 | 25,70733027 | 25,98973351 | 24,81928879 | 24,81181329 | NA | NA | DHRS7B | 3 |
| 26,55739428 | 26,45480118 | 26,37373247 | NA | NA | NA | NA | NA | NA | NA | NA | NA | 26,26309466 | 26,16731477 | 26,06847768 | 26,53555091 | 26,35665856 | 26,44384287 | PPIF | 3 |
| 26,21233934 | 25,93262766 | 26,18116561 | NA | NA | NA | NA | NA | NA | NA | NA | NA | NA | NA | NA | 26,37772553 | 26,50127231 | 26,03250418 | LTBP4 | 3 |
| 28,22999778 | 28,03773889 | 28,17944862 | 27,37285284 | 27,39607348 | 27,33602895 | 29,25745863 | 28,9760993 | 28,65263357 | 26,74906392 | 26,7261144 | 26,77743493 | 28,28042063 | 28,15678565 | 28,48199215 | 28,08050267 | 28,28330289 | 28,42930068 | PYCR1 | 1 |
| 26,14670353 | NA | 26,05262543 | 27,63961782 | 28,20443159 | 27,74983118 | 26,91182242 | 27,19403931 | 26,95038076 | 26,08127505 | 26,12563592 | 25,99051275 | 27,51359762 | 27,21918861 | 27,60907681 | 26,55996307 | 26,70990858 | 26,78656159 | GBP2 | 2 |
| 27,68705707 | 27,2780823 | 27,47599213 | 26,18757907 | 26,68230905 | 26,55920459 | 26,23222314 | 26,04310073 | NA | 26,40851724 | 26,34484107 | 26,52218556 | 27,65328299 | 27,84002124 | 27,68036521 | 27,64485197 | 27,97339197 | 27,86673362 | MPI | 3 |
| 32,65469619 | 32,69644016 | 32,60915609 | 31,34239088 | 31,50852523 | 31,27052774 | 31,37885847 | 31,43738012 | 31,360923 | 31,75640801 | 31,89677566 | 32,00135396 | 33,00816708 | 32,89500657 | 33,07515905 | 32,42589917 | 32,51516851 | 32,47050921 | CTNNA1 | 3 |
| 28,7430396 | 28,51337172 | 28,69919027 | NA | 23,4652565 | NA | 28,87730501 | 29,14513457 | 29,13589517 | 27,77680769 | 27,36151923 | 27,31734471 | 27,5798902 | 27,44868675 | 27,56637962 | 29,01363775 | 28,64879944 | 28,76670314 | FBN2 | 3 |
| 27,19826143 | 27,10209457 | 26,88471738 | 25,56563903 | 26,36721383 | 24,89642923 | NA | 24,38396632 | NA | 25,29827545 | 25,15584042 | 25,21981638 | 26,85138534 | 26,6056902 | 26,51132196 | 27,31094694 | 27,0215751 | 27,07551293 | TIE1 | 3 |
| 32,22937329 | 32,23680383 | 32,03642937 | 30,63460696 | 30,37036042 | 30,80532026 | 30,00597767 | 30,30497753 | 30,12361475 | 30,48529385 | 30,16770238 | 30,26840982 | 32,6910823 | 32,71128792 | 32,6910615 | 31,87693924 | 31,91972067 | 32,00175661 | MCAM | 3 |
| 32,09468358 | 32,08156966 | 32,07041631 | 30,7971472 | 30,83084996 | 31,04381258 | 30,60470094 | 30,88649186 | 30,78316147 | 32,12444697 | 32,22547124 | 32,0805535 | 32,59646024 | 32,50215957 | 32,59071714 | 31,85005916 | 31,63005621 | 31,54190137 | MAP1B | 3 |
| 24,67476037 | NA | NA | 26,26350662 | 25,52656491 | 25,77841286 | 25,72541244 | 25,83930049 | 25,70717227 | 25,50663882 | 25,77901434 | 25,47586848 | 26,03107419 | 26,05996509 | 25,67147066 | 25,25753057 | NA | NA | SYNJ1 | 2 |
| 28,1297189 | 27,3788998 | 27,63685536 | 26,55841651 | 26,30676443 | 26,24459669 | 26,06737018 | NA | NA | 26,39982604 | 26,7679977 | 26,28185364 | 27,47444567 | 27,57693879 | 27,33108149 | 28,43876857 | 28,39235208 | 28,46707725 | ENTPD1 | 3 |
| 28,12593146 | 28,02828372 | 28,31501367 | 26,08865271 | 26,63394907 | 26,75174753 | 26,95038076 | 27,02115146 | 27,44498036 | 27,90604052 | 27,85845753 | 27,84917934 | 28,10224478 | 28,10304564 | 28,24585724 | 27,90758845 | 27,78886537 | 27,41989076 | GNAQ | 3 |
| 30,54701429 | 30,64400959 | 30,42171748 | 31,8177605 | 31,55041287 | 31,7292301 | 31,22112642 | 32,1277096 | 31,87540201 | 31,88554671 | 31,89605384 | 32,0192303 | 30,32206 | 30,25900894 | 30,2997476 | 30,40202409 | 30,49228352 | 30,52729226 | HSPA1B;HSPA1A | 2 |
| 26,40282107 | 26,23730203 | 25,9538254 | NA | 25,4737647 | NA | NA | NA | NA | NA | NA | NA | 25,80432347 | 25,52112235 | 25,57224733 | 26,55733585 | 26,12601026 | 26,13772336 | METAP2 | 3 |
| 26,38819404 | NA | 26,51005448 | 27,20134981 | 27,33985585 | 27,28824061 | 27,09988969 | 26,96960961 | 27,24445152 | 27,0747987 | 27,31388998 | 27,66661681 | 27,41215684 | 27,026438 | 26,78294284 | 27,33048324 | 26,93517081 | 26,87981898 | NUDT2 | 2 |
| 25,2662441 | 25,62828577 | 25,55946719 | NA | NA | NA | NA | NA | NA | NA | NA | NA | NA | NA | NA | 24,95571092 | NA | 25,5995742 | IDH3G | 3 |
| 26,22171647 | 27,16432525 | 26,78556422 | 28,36185384 | 28,49121333 | 28,34726809 | NA | 24,60715871 | NA | 28,30669491 | 28,52622908 | 28,45998361 | 30,30888757 | 30,18210386 | 30,11445946 | 25,48646434 | 24,86240051 | 25,6164434 | SLC12A2 | 4 |
| NA | NA | NA | 25,51337172 | 24,8381225 | 25,33101313 | 25,06755483 | 26,08107183 | NA | NA | 25,07030114 | 24,79585378 | 25,33606302 | 25,14546016 | 25,29914931 | 27,23708316 | 26,96422046 | 26,82711356 | CDH11 | 2 |
| 30,23134897 | 30,22262412 | 30,210947 | 29,10971052 | 29,12240562 | 29,13447599 | 29,40261374 | 29,21909627 | 29,24499584 | 29,06332266 | 28,98212468 | 28,85922803 | 29,05058748 | 29,14243469 | 29,28460509 | 29,79771928 | 29,68899192 | 29,34470988 | CDH13 | 3 |
| 25,61506853 | 25,78977341 | 25,60320163 | NA | NA | 23,70893568 | 23,94154271 | NA | 24,21697079 | 25,33650574 | 24,99053439 | 25,23253423 | 26,22324579 | 25,97762302 | 26,30389388 | NA | NA | NA | BNIP3 | 4 |
| NA | NA | NA | 26,81885012 | 26,65020419 | 26,81738694 | 26,28943723 | 26,41617399 | 26,05080023 | 26,2740522 | 25,91615792 | 26,01307864 | NA | NA | NA | NA | NA | NA | SESN2 | 2 |
| 26,35326315 | 26,58663412 | 26,49321547 | 24,79892145 | NA | 24,98263609 | NA | NA | NA | 24,61487202 | NA | 25,15007853 | 26,26911912 | 26,97257056 | 26,75913358 | NA | NA | NA | CXCR4 | 3 |
| 27,0403021 | 27,22296953 | NA | 28,0401453 | 28,16906537 | 27,72468412 | 28,04379954 | 27,98250553 | 28,30247389 | 27,74631124 | 27,93438358 | 27,78973611 | 28,4921689 | 27,93584523 | 27,87314443 | 27,69305332 | 27,22296953 | 27,53207621 | RAP2B | 2 |
| NA | NA | NA | NA | NA | NA | 25,68664308 | 25,53299768 | 25,31931021 | 25,2005643 | 25,49191669 | 25,13625708 | NA | NA | NA | NA | NA | NA | RND3 | 2 |
| 25,84866611 | 25,59348915 | 25,62848047 | NA | NA | NA | NA | NA | NA | NA | NA | NA | 25,66779459 | 25,69784569 | 25,88609061 | 25,82230708 | 26,03693236 | 26,2399258 | SIRPA | 3 |
| 26,48083728 | 26,53734437 | 26,10835009 | 27,69218873 | 27,46037428 | 27,6681058 | 27,46995153 | 27,52163159 | 27,74611899 | 27,76357245 | 27,79054418 | 27,86242416 | 27,27213084 | 27,16959109 | 27,27132951 | 27,32748823 | 27,60116232 | 27,33790115 | EIF4G3 | 2 |
| 25,72148036 | 25,57065599 | 25,71866143 | 26,21262697 | 27,22315371 | 27,41675385 | 26,86495236 | 26,02591022 | 26,38756971 | 27,12435453 | 27,20527094 | 27,27630832 | 27,7667979 | 27,59327542 | 27,51141246 | 26,0173756 | 26,00705842 | 26,2335951 | EFNB1 | 2 |
| 28,52114482 | 28,3595099 | 28,24377085 | 27,07265389 | 27,06773944 | 27,27817094 | 25,9503585 | NA | NA | 26,72181931 | 26,75276855 | 27,14297021 | 27,97443475 | 27,71639969 | 27,78805635 | 27,09748051 | 27,34545032 | 27,40673454 | CDK17 | 3 |
| 32,65795705 | 32,57081882 | 32,22357379 | 31,43589102 | 31,07233445 | 31,21824183 | 29,89197602 | 29,79971205 | 29,73669977 | 31,24875504 | 31,0992376 | 31,28769037 | 33,22417779 | 32,80606971 | 33,12962986 | 31,9062341 | 31,65215486 | 31,83156739 | FABP5 | 3 |
| NA | NA | NA | 25,26004604 | 25,28227796 | 25,60978136 | 25,89015608 | 25,92526557 | NA | 25,98682931 | 26,15166631 | 26,16716161 | 25,62458152 | 25,89346967 | 25,66638656 | NA | NA | NA | OTUD4 | 2 |
| NA | 25,1832498 | NA | 26,07314441 | 26,59135045 | 26,00960193 | 26,13055375 | 26,53215055 | 26,61526502 | 25,28390337 | 26,50107729 | 25,47639394 | NA | NA | NA | 25,72689396 | 25,9583465 | 25,49265026 | XPC | 2 |
| 27,6621406 | 27,90397403 | 27,82656817 | 28,91866201 | 29,16199737 | 29,44251071 | 32,14628231 | 32,61787211 | 32,35129179 | NA | NA | NA | 25,79790786 | 25,28036753 | 25,58597547 | 26,24572126 | 26,99334477 | 26,90138676 | TAGLN | 2 |
| 32,24872677 | 32,24612349 | 32,16104834 | 30,92634627 | 30,99745588 | 31,24541527 | 30,53741844 | 31,217722 | 31,42947029 | 33,23534932 | 33,13412097 | 33,10016235 | 32,19075277 | 32,19727151 | 31,98522299 | 30,59894627 | 30,86348796 | 30,92500235 | AKAP12 | 4 |
| NA | NA | 25,63464158 | 26,71109096 | 26,83701575 | 26,79525928 | 26,78668621 | 26,62310273 | 26,81750893 | 25,93251503 | NA | 26,16311618 | 26,3854318 | 26,2789862 | NA | 27,01924356 | 27,07398201 | 27,28162375 | CSTF1 | 2 |
| NA | NA | NA | 25,7648888 | 25,88432146 | 25,80456965 | 25,9671265 | 26,00823455 | 25,66592594 | 25,42540801 | 25,5388101 | 25,32964526 | NA | NA | NA | 24,27795819 | NA | NA | PRKCD | 2 |
| 26,98630854 | 26,69019155 | 26,97136498 | 28,31630916 | 28,44264913 | 28,36314975 | 28,15913638 | 28,09692784 | 27,69172297 | 28,773636 | 29,02554066 | 28,72533443 | 27,75576361 | 27,43706258 | 27,5236518 | 28,05553474 | 28,28374445 | 28,17840416 | SRSF4 | 2 |
| 26,52534063 | 26,8312276 | 27,39018024 | NA | NA | NA | NA | NA | NA | NA | NA | 25,57390913 | 26,7224709 | 27,3972166 | NA | 26,67276587 | 26,71109096 | 26,94100532 | CAMSAP2 | 3 |
| NA | NA | NA | NA | 24,81811872 | NA | NA | NA | NA | NA | NA | NA | 26,19078455 | 26,4842223 | 26,53716661 | 27,652531 | 27,19413327 | 27,70464182 | CGNL1 | 4 |
| 30,26472392 | 30,32635278 | 30,26852137 | 27,14180156 | 26,81787483 | 27,3900162 | 31,36202112 | 30,90591145 | 31,1369342 | 29,27408776 | 28,72315476 | 28,84346983 | 27,71017141 | 27,55464529 | 27,62477672 | 30,74518945 | 31,03091638 | 30,77266111 | EFEMP1 | 1 |
| 25,61633121 | 25,45327892 | 25,38053558 | NA | NA | NA | 26,11193527 | 25,8540281 | 26,0548209 | 27,60978136 | 27,49982874 | 27,66756453 | 27,24099945 | 27,19168834 | 27,16317378 | 26,30789369 | 26,19844879 | 26,43273698 | FSTL1 | 4 |
| NA | 25,18570905 | 24,8811035 | NA | 25,87112428 | 25,81369803 | 25,55042388 | 25,59789911 | 25,09663635 | 26,01005032 | 26,20041463 | 26,06416597 | 25,73662881 | 25,74216591 | 25,5143955 | NA | 25,15915563 | NA | BNIP1 | 2 |
| 33,21956946 | 33,1421486 | 33,46319371 | 30,48471784 | 30,48682873 | 30,43589102 | 32,26932983 | 32,2681867 | 32,23552051 | 31,42342167 | 31,23089108 | 31,16740331 | 31,89511494 | 32,01021976 | 31,74538983 | 32,84927629 | 32,91929445 | 32,8720657 | MMRN1 | 3 |
| 25,68648279 | 25,06870318 | 25,31222876 | NA | NA | NA | NA | NA | NA | 24,71706681 | 24,67330519 | NA | 24,55859167 | 24,65272246 | 25,10952621 | 25,47044812 | 25,94628092 | 25,47206087 | SELENBP1 | 3 |
| 27,70450991 | 27,47204537 | 27,76268576 | NA | NA | NA | 25,92091197 | 25,64729724 | NA | 25,26745909 | 26,0288001 | NA | 26,62324231 | 26,42324533 | 26,62617022 | 27,32997025 | 27,728841 | 28,18697488 | PTK7 | 3 |
| NA | NA | NA | 27,91050778 | 27,92767481 | 27,86514121 | 28,67367586 | 28,48945343 | 28,53006777 | 27,63200829 | 27,76141813 | 27,60808987 | 27,41199527 | 27,25511878 | 27,51397403 | 27,44181846 | 27,49304746 | 27,47753694 | GOLGA4 | 2 |
| 27,92925615 | 28,06763688 | 28,05863432 | 28,88739281 | 29,28431827 | 29,07890071 | 29,29158161 | 29,21671644 | 29,27117365 | 29,20242451 | 29,06352839 | 28,96911552 | 28,83839905 | 28,90420379 | 28,88692787 | 29,36348398 | 29,52413773 | 29,22048077 | PDAP1 | 2 |
| 27,23863273 | 27,0383147 | 27,27497639 | 26,60978136 | 25,94737386 | 25,82381386 | 25,54495288 | 25,90062694 | 25,81124983 | 27,18186699 | 27,64003173 | 27,49258915 | 27,98652555 | 28,14150925 | 28,04004075 | 27,51239242 | 27,70391614 | 27,62470701 | ROCK1 | 3 |
| 26,40103131 | NA | NA | 26,65962512 | 27,17607512 | 26,67168661 | 25,88709062 | 25,69667893 | 26,65485407 | 26,84541539 | 27,06609757 | 26,72715371 | 26,6707416 | 26,47047915 | 26,56099805 | 26,24805829 | 26,10547552 | 26,66139321 | RIPK1 | 2 |
| NA | NA | NA | 27,7079358 | NA | 27,44427452 | 27,00955922 | 26,92592201 | NA | NA | NA | NA | 27,81860636 | 27,70028223 | 27,63519534 | NA | NA | NA | ATP6AP2 | 2 |
| 25,5669457 | 26,28894462 | 26,30811943 | NA | 25,09627441 | 24,67868739 | 25,37122495 | 25,21322973 | 24,36385991 | 25,20687366 | 25,60455956 | 25,46665728 | 26,7206457 | 26,77956548 | 26,1574026 | 26,42761541 | 26,41911923 | 26,40887352 | IDI1 | 3 |
| 27,73130349 | 28,15822159 | 27,96515676 | 28,94218059 | 29,25421783 | 28,78861649 | 29,31170491 | 29,34747931 | 29,53963854 | 28,2826403 | 28,225454 | 27,99139971 | 28,64705015 | 28,95615421 | 28,95751094 | 29,02427286 | 28,9054381 | 28,93691241 | CIRBP | 2 |
| 26,94648173 | 26,60356953 | 26,24517723 | 24,87272184 | 24,95264848 | 25,23699196 | 24,81308637 | 24,39976086 | NA | 25,49673921 | 26,10491591 | 25,40259341 | 25,9621695 | 25,80607048 | 25,42652813 | 27,15005915 | 27,18404478 | 27,01499471 | DAG1 | 3 |
| 28,1844232 | 28,02258074 | 28,25520885 | 26,61386094 | 27,30338877 | 26,70754092 | 26,53799599 | 27,09063258 | 26,25003003 | 26,28418587 | 26,58147139 | 25,96932416 | 27,81616649 | 27,96223571 | 28,03433167 | 27,32825897 | 27,38187256 | 27,01403699 | ASAH1 | 3 |
| 29,30014051 | 28,99023682 | 29,12045408 | 28,15450834 | 28,04207798 | 28,01179984 | 30,03249367 | 30,39130755 | 30,17080907 | 28,08182359 | 27,45810693 | 27,80372014 | 29,52241001 | 29,5929655 | 26,15348431 | 28,47908009 | 28,69252132 | 28,46349581 | LTBP1 | 1 |
| 27,21632328 | 27,61680794 | 27,57845124 | 28,47529643 | 28,71571262 | 28,51765774 | 28,97555277 | 28,9749787 | 28,83584191 | 27,27701817 | 27,72253605 | 27,3762522 | 27,76097419 | 28,07841723 | 27,67808289 | 28,3248303 | 28,11679091 | 28,28343537 | GOLGB1 | 2 |
| 25,73407201 | 25,78451624 | NA | 27,74599082 | 28,04967907 | 28,1125717 | 25,51851341 | 26,92942548 | 26,64265043 | 26,85543084 | 27,22112642 | 27,37102549 | 25,98830382 | 26,1777296 | 26,26740551 | 25,96382373 | 26,1125717 | 26,11816817 | PSME4 | 2 |
| NA | NA | 28,02078068 | 28,84469734 | 28,76891267 | 28,85655917 | 28,08253435 | 28,48050604 | 28,39750223 | 27,92648766 | 28,05687871 | 27,91900314 | 28,62693608 | 28,47324602 | 28,38590899 | 28,45430701 | 28,54932987 | 28,16700844 | RABEP1 | 2 |
| 28,66444826 | 28,23260742 | 28,54333113 | NA | 25,89746826 | 26,36396014 | NA | 27,21604569 | 26,59576694 | 26,83291821 | 27,30147122 | 26,77718407 | 28,24513188 | 28,4086792 | 28,4688258 | 28,46995153 | 28,61999374 | 28,05765351 | VAMP3 | 3 |
| 26,32697417 | 26,46151442 | 26,2247551 | NA | 24,49234465 | 31,94363436 | 31,93266004 | 31,99742223 | 24,84517608 | 25,24869171 | 24,02791477 | NA | NA | NA | NA | 27,22370611 | 27,71279712 | 27,80636554 | IGFBP7 | 1 |
| 29,03915183 | 28,95986144 | 28,94198478 | 27,61456315 | 27,59042269 | 27,67667142 | 27,06074853 | 27,35194991 | 26,76268576 | 27,25024694 | 27,36335865 | 27,46559904 | 28,86705778 | 28,92323213 | 28,97262533 | 28,81729544 | 28,73489249 | 28,71230517 | TST | 3 |
| 24,9791942 | NA | NA | 25,87602881 | 25,78591337 | 26,15749898 | 24,43879236 | NA | 24,46687505 | 25,87368423 | 26,03975845 | 25,90664842 | 27,47521911 | 27,395665 | 27,35623877 | 25,67591806 | 25,88000589 | 25,84484098 | RALGAPA2 | 4 |
| 25,8362935 | 25,96778615 | 26,37999052 | NA | 24,23622563 | 24,02943215 | 25,22195611 | 25,43205197 | NA | 24,94315366 | 25,17729239 | 25,27113357 | 25,50497339 | 25,47685743 | 25,41267373 | 25,40691291 | 25,96941199 | 26,33596083 | UAP1L1 | 3 |
| NA | NA | NA | 26,10391606 | 25,77235908 | 26,05019824 | 26,16215588 | 25,85095633 | 25,96932416 | 25,66755099 | 25,65023158 | 25,83869959 | 25,99714088 | 25,4354579 | 24,99761469 | 25,68984509 | NA | 25,41796116 | RABL6 | 2 |
| 25,76339515 | 26,300878 | 26,4017963 | 27,87443491 | 27,93145585 | 27,73214496 | 27,3390063 | 27,40786924 | 27,28920854 | 26,91968515 | 27,1270146 | 27,21029678 | 27,61280698 | 27,2972786 | 27,10109277 | 26,46240404 | 26,78369229 | 26,55436717 | VPS26B | 2 |
| 25,24212682 | 25,29949871 | 24,75414577 | 27,02538224 | 27,09093538 | 26,70951424 | 26,73673202 | 26,4526036 | 25,84071762 | 25,52554972 | NA | 24,639811 | 27,04510239 | 27,11832661 | 27,01637696 | NA | NA | NA | GXYLT1 | 2 |
| NA | NA | 25,44059928 | 26,65116274 | 26,8793516 | 26,68431716 | 26,42051732 | 26,34207928 | 26,20234043 | 26,69657282 | 26,75506319 | 26,66343063 | 26,58362477 | 25,8736373 | 26,32630563 | 26,39886436 | 26,00077664 | 26,18601144 | GRIPAP1 | 2 |
| 29,19413327 | 29,18084798 | 29,06237077 | 30,60893586 | 30,72427754 | 30,75724291 | 30,22101115 | 30,14126561 | 30,17200216 | 28,93308936 | 28,94413724 | 29,24633306 | 29,26948032 | 29,02411431 | 28,95781534 | 28,70698132 | 28,53274508 | 28,56260007 | TP53I3 | 2 |
| 25,77743493 | 25,40778822 | 25,31202098 | NA | 24,39519103 | NA | NA | NA | NA | 24,16652966 | 24,65075201 | NA | 25,04067835 | 24,85922803 | 24,70919869 | 24,85500303 | 25,81558031 | 25,75271751 | HSPA12B | 3 |
| 27,43125503 | 27,06927701 | 27,08233131 | 28,38195505 | 29,09861031 | 28,69617481 | 28,10294555 | 28,37127481 | 28,4806216 | 28,79878552 | 28,32156551 | 28,54067342 | 28,35455838 | 28,46337888 | 28,58082475 | 27,48769139 | 26,99841121 | 27,31484085 | MYCBP | 2 |
| 25,63588725 | 26,24241762 | 26,01775788 | 27,79841474 | 27,89261304 | 28,1841394 | 27,82784042 | 27,38681358 | 27,325774 | 28,08658915 | 27,9538254 | 27,92614829 | 27,19713674 | 27,20181717 | 27,06167575 | 27,88046138 | 27,49381099 | 27,69025817 | FKBP15 | 2 |
| 25,84496067 | 26,17420918 | 25,62731189 | 25,36101716 | NA | 25,54008215 | 26,02455821 | 26,32993604 | 25,83414869 | NA | NA | 24,88920463 | NA | NA | NA | 26,02312032 | 26,56383577 | 25,66955271 | RIF1 | 1 |
| 24,91779745 | 25,24517723 | 24,91574775 | NA | NA | NA | NA | NA | NA | 25,36416058 | 25,50821163 | 25,41118718 | 24,35056803 | 24,42284447 | 24,55344462 | 25,32844731 | 25,42882964 | 25,10149357 | NAA35 | 3 |
| 25,21018529 | 26,13686332 | 25,38564573 | 26,55904409 | 26,81445612 | 26,57744312 | 27,01616439 | 26,99194026 | 26,56955549 | NA | NA | NA | 26,38236742 | 25,8146028 | 25,97101407 | 25,71581079 | 25,63161982 | 25,10593504 | KDELC1 | 2 |
| 29,9969847 | 30,02498613 | 30,12213405 | 28,79776565 | 29,04385168 | 28,59576694 | 27,91826392 | 28,10184418 | 27,88331953 | 26,42095094 | 24,76668418 | 26,58233313 | 30,27464322 | 30,17080907 | 30,02670182 | 30,01118668 | 30,21662394 | 30,19483777 | CD109 | 3 |
| 24,92474474 | 25,52880176 | 25,68461145 | NA | NA | NA | NA | 24,79436708 | NA | NA | NA | NA | NA | NA | NA | 24,54073253 | 24,93701347 | NA | QSOX2 | 1 |
| NA | NA | NA | 27,44931668 | 27,57801928 | 27,25898648 | 27,19055852 | 27,55105495 | 27,43999722 | NA | NA | NA | 27,75633642 | 28,26163392 | NA | NA | NA | NA | TMED4 | 2 |
| NA | NA | NA | 25,84988324 | NA | 25,68554742 | 26,56140601 | 26,64141059 | 26,2570989 | 25,93077938 | NA | 25,65400727 | 26,08768199 | 26,3747442 | 25,63184181 | NA | NA | NA | RALGAPB | 2 |
| 24,18071047 | NA | NA | 24,81382033 | 24,83773764 | 24,74309098 | 24,92750527 | 24,96075639 | NA | 24,69678504 | 24,4754356 | 24,68683006 | 24,29775088 | 24,15073709 | 24,58978005 | 25,28160606 | 25,21504591 | 24,99049111 | SPATS2 | 2 |
| 26,11676117 | 25,79585378 | 25,4652565 | NA | NA | NA | 25,14117789 | 24,88604408 | NA | 25,71573225 | 25,45353013 | 25,09924763 | 25,59576694 | 25,63870679 | 25,78196797 | 25,7456575 | 26,86997199 | 26,39445524 | TRIM22 | 2 |
| 30,6462434 | 30,61745617 | 30,58530935 | 31,7876828 | 31,73976857 | 31,81130496 | 32,01733976 | 31,98243753 | 31,83424516 | 32,04430128 | 31,64864517 | 32,06384336 | 30,91748164 | 30,72906799 | 30,99886847 | 30,46626053 | 30,67109605 | 30,48577367 | CD9 | 2 |
| NA | NA | NA | 26,52157169 | NA | NA | NA | NA | NA | 26,57830727 | NA | 26,27369659 | 26,37666621 | 25,89977455 | 26,21748859 | NA | NA | NA | OXR1 | 4 |
| 25,76448389 | NA | 26,64278813 | 27,37840375 | 27,66600724 | 27,34638063 | 27,44521722 | 26,92297704 | 26,87114778 | NA | NA | NA | 26,41660891 | 26,65102584 | 26,86549523 | 26,70358616 | 26,44414313 | 26,82264746 | PDPR | 2 |
| NA | NA | NA | NA | 25,99392777 | NA | 25,53198701 | 25,60002813 | 24,66394632 | 24,458326 | 26,97213228 | 26,74393844 | 25,74855219 | 25,84110168 | 25,26305883 | 25,18196174 | NA | 25,80769256 | APPL2 | 2 |
| NA | NA | NA | 26,12242044 | 27,08222978 | 26,15230481 | 26,04462308 | 26,06778047 | 26,06422766 | 26,00725094 | 25,72941164 | 25,80163732 | NA | NA | NA | 25,51890355 | 25,79434229 | 25,79498672 | TSPAN14 | 2 |
| 24,51713207 | 24,69173628 | NA | 25,96745636 | 26,10739254 | 25,69468796 | 25,38010616 | 25,66866039 | 25,48495979 | NA | NA | 24,42194616 | 24,66356636 | 24,54226863 | NA | 25,57427014 | 25,28725442 | 24,55121634 | SLC35B2 | 2 |
| 27,39140999 | 25,81999524 | 26,15216939 | 29,04609177 | 29,97541611 | 29,14011997 | 28,96375208 | 28,51634321 | 28,90592579 | 28,06948189 | 28,40835527 | 28,43785631 | 28,01307864 | 28,29215232 | 27,83972097 | 26,60978136 | 26,30975069 | 26,63878964 | BICD2 | 2 |
| NA | 25,40288611 | 25,44256213 | 26,67572966 | 27,0454149 | 26,81592228 | 26,16530327 | 26,28985933 | 25,920753 | 25,86422035 | 26,07728678 | 25,9712334 | 26,25147548 | 26,29330184 | 26,26282592 | 26,7810675 | 27,00838417 | 26,96939003 | NEK9 | 2 |
| 24,84244511 | 25,12234144 | 25,25328024 | 24,53082684 | NA | NA | 24,05266688 | NA | NA | NA | NA | NA | NA | NA | 24,18268168 | 24,98115578 | 25,10593504 | 24,92397447 | MICAL1 | 3 |
| NA | NA | NA | 26,78094239 | 26,4322591 | 26,52602754 | NA | NA | NA | NA | NA | NA | 26,67586424 | 26,45621214 | 26,19084106 | NA | NA | NA | AFAP1L1 | 2 |

| | | | | | | | | | | | | | | | | | | | |
|---|---|---|---|---|---|---|---|---|---|---|---|---|---|---|---|---|---|---|---|
| 26,19236582 | 26,36436099 | 26,50266891 | 25,65329666 | NA | NA | NA | NA | NA | 24,9652999 | 25,24778673 | 24,99851881 | 25,96115397 | 26,21313701 | 25,32202991 | 26,60583147 | 26,86041261 | 26,24159961 | GIPC2 | 3 |
| NA | NA | 25,08998639 | 25,85621483 | 25,64825773 | 25,85619108 | 25,44865525 | 25,48049834 | 25,64303594 | 25,97318392 | 26,01318515 | 26,04168121 | 26,39114773 | 26,23072163 | 25,84033345 | 24,49661732 | NA | NA | TGFBRAP1 | 2 |
| NA | NA | NA | 25,93303306 | NA | 26,36404366 | 25,2128588 | 25,43837994 | 25,412383 | 25,82825214 | 25,33036356 | 25,94467346 | NA | 25,28121692 | NA | 25,67976142 | 24,94154271 | NA | TRAPPC12;CGI-87 | 2 |
| NA | 24,69948816 | 25,07058777 | 25,51282942 | 25,24328964 | 25,6960952 | NA | 25,25234205 | 25,8723461 | 25,93280785 | 26,05386856 | 26,08294035 | 25,59032275 | 25,85067025 | 25,66207267 | 26,41612565 | 26,08031968 | 26,17706423 | SREK1 | 2 |
| 25,31962031 | 25,57732786 | 25,83958883 | 24,69492702 | NA | 24,79704204 | NA | NA | NA | 25,15877054 | NA | 25,16886459 | 25,4556636 | 25,11684047 | 25,27735523 | 26,48916246 | 26,36733049 | 25,95816947 | DDB2 | 3 |
| NA | NA | NA | 24,94176656 | 24,74545235 | 24,99386301 | NA | NA | NA | 24,64421937 | 24,9910969 | 24,6243863 | NA | NA | NA | 24,12257842 | NA | NA | TSC2 | 2 |
| 31,67934197 | 31,75186722 | 31,8073947 | 30,43807451 | 30,51493345 | 30,46246255 | 30,53417465 | 30,72468412 | 30,41749848 | 28,67047148 | 28,44047255 | 28,36823847 | 31,89048373 | 31,83428284 | 31,90540941 | 30,44233274 | 30,66186887 | 30,63157472 | PXDN | 3 |
| 25,01945567 | NA | 24,87988907 | 26,34160433 | 25,68330007 | 26,31848296 | 25,57192921 | 26,24775052 | 26,4804367 | 26,87290967 | 26,59662018 | 27,13682422 | 24,76309117 | 25,544098 | 25,85740188 | 25,44123275 | 25,92992206 | 25,24706234 | SMARCC1 | 2 |
| 28,77426462 | 28,77737222 | 28,93851171 | 27,33261873 | 27,08851119 | 27,57701085 | 28,01945567 | 27,76186193 | 27,48116074 | NA | 25,53928355 | 25,65337867 | 28,87114778 | 28,62850133 | 29,00971938 | 28,77407606 | 28,97314559 | 28,84212136 | LPP | 3 |
| NA | NA | NA | 26,72663415 | 26,74033973 | 26,83110676 | 25,57701085 | 26,64705015 | 26,05635161 | 26,78917641 | 26,98381055 | 27,05475881 | 26,86950141 | 26,39948382 | 26,55775945 | 26,33058581 | 26,03485638 | NA | SYAP1 | 2 |
| 29,61243791 | 29,64113492 | 29,47554769 | 28,30699904 | 28,48273848 | 28,456502 | 28,39766543 | 28,23480135 | 28,5862404 | 29,47303307 | 29,43251001 | 29,52941629 | 29,40950893 | 29,20671533 | 29,29725673 | 29,46750487 | 29,30486877 | 29,20897222 | FERMT2 | 3 |
| 28,38619688 | 28,37418037 | 28,31919821 | 26,4230369 | 26,47529643 | NA | NA | NA | NA | 26,86561322 | NA | 26,94681634 | 28,81424218 | 28,56310944 | 28,81182553 | 28,79745646 | 28,14331088 | 28,22015784 | ESAM | 3 |
| 28,49963869 | 28,61400141 | 28,49922049 | 31,04596162 | 30,92309042 | 31,06721373 | 30,15786033 | 30,23728835 | 30,06368267 | 30,30486877 | 30,0604651 | 30,05258397 | 30,39212685 | 30,14637336 | 30,38329484 | 28,87214645 | 28,43447991 | 28,42129613 | PBXIP1 | 2 |
| 25,45236794 | 25,44937966 | 25,38962241 | NA | NA | NA | NA | NA | NA | 25,39891327 | 25,14109992 | 25,54986597 | 25,28647908 | NA | NA | 25,9286577 | 25,49619062 | 25,65916234 | FLYWCH2 | 3 |
| 26,68618888 | 27,13858289 | 27,43356494 | NA | NA | NA | NA | NA | NA | NA | NA | NA | 27,10239497 | 27,59220631 | 27,12839197 | 26,0898248 | 24,87965541 | 25,8159467 | ZC3HAV1L | 3 |
| 27,60272038 | 28,04723652 | 27,77511281 | NA | NA | NA | NA | NA | NA | 25,09329505 | 24,78481574 | NA | 28,70553194 | 28,94793117 | 28,48146873 | 27,60597272 | 27,38755328 | 27,55288787 | DCHS1 | 3 |
| 27,46629943 | 27,52663952 | 27,9290868 | 25,72507434 | NA | 26,03120042 | NA | NA | NA | 27,30565168 | 27,30626032 | 27,07081293 | 26,44047255 | 26,55825592 | 26,53143679 | 27,29570322 | 27,37815565 | 27,48684791 | TRIM47 | 3 |
| 25,14328169 | 25,22460792 | 24,97897608 | 26,48721603 | 26,63823725 | 26,50800001 | 26,42322929 | 26,67976142 | 26,26527852 | 26,00536738 | 26,05494507 | 26,13447599 | 26,08669037 | 26,0020867 | 25,94453943 | 25,84040549 | 25,60772312 | 25,42940444 | DHRS1 | 2 |
| 27,74451596 | 27,76319251 | 27,80851519 | NA | NA | NA | NA | 26,35314534 | NA | 26,19078455 | 26,82893002 | 26,52671414 | 27,51254312 | 26,90253723 | 27,09526856 | 28,42498374 | 28,33150867 | 28,21488848 | PUS7 | 3 |
| 25,68691018 | 25,76146886 | 25,16177158 | NA | NA | NA | NA | NA | NA | NA | NA | 25,07071059 | 25,46957897 | 25,75177307 | 25,46385436 | NA | 25,09139955 | 25,00615963 | GPX7 | 3 |
| 27,4809297 | 27,74541388 | 27,49579429 | 25,19227174 | NA | NA | 26,14129485 | 26,44737877 | 26,14267813 | 27,2295853 | 27,14326222 | 27,20555061 | 26,91136529 | 26,82529481 | 26,76597641 | 27,16624232 | 27,36101716 | 27,0558967 | PLIN2 | 3 |
| 26,89122282 | 27,42934059 | 27,5555232 | NA | NA | NA | 25,03132664 | 25,68217507 | 25,79382157 | 26,38825975 | 26,53058875 | 26,34394364 | 26,73414956 | 26,92987692 | 26,77504999 | 27,0820267 | 27,41804162 | 27,1284903 | TSNAX;DISC1 | 3 |
| NA | NA | NA | NA | 26,38066768 | 26,95925345 | 25,7665831 | 25,74995901 | 26,04393509 | NA | NA | NA | 25,90403148 | NA | NA | NA | NA | NA | AGPAT1 | 2 |
| 28,05346466 | 28,42642415 | 28,21618449 | 27,9686762 | 27,5896372 | 27,65724157 | 27,75206668 | 27,81200922 | 28,13775267 | 26,61428231 | 26,62811887 | 26,4426254 | 27,00345992 | 26,8075943 | 26,96675257 | 27,69869365 | 27,3711086 | 27,59077959 | NIPSNAP1 | 1 |
| 24,6880048 | 24,91328425 | 24,62834141 | 25,76068239 | 25,87696558 | 25,68086147 | 26,04743415 | 26,11084074 | 25,9789979 | 24,33360852 | 24,67076861 | NA | 25,37792406 | 25,34346931 | 25,67968089 | 25,64812055 | 25,54848493 | 25,78186794 | MSTO1 | 2 |
| NA | NA | NA | 28,05393069 | 28,23329338 | 28,12602996 | NA | 28,17426634 | 28,15995439 | NA | NA | 27,66431262 | NA | NA | NA | NA | NA | NA | ELOVL1 | 2 |
| NA | NA | NA | 24,97316202 | 25,76372439 | 25,14339847 | 25,70812004 | 25,6002834 | 26,16175237 | 25,79275475 | NA | 25,28079229 | NA | NA | NA | 24,3116746 | NA | NA | LSM14B | 2 |
| 26,53109466 | 26,81689888 | 26,28741296 | NA | NA | NA | 25,37305205 | NA | NA | 27,22536206 | 27,14121688 | 26,89781444 | NA | 25,26381104 | 25,03535992 | 26,58362477 | 27,23817715 | 26,84050154 | MAK16 | 3 |
| NA | NA | NA | 25,84677872 | 25,73127759 | 25,79317661 | 25,76564768 | 25,68955187 | NA | NA | NA | NA | NA | 25,56302213 | 25,74116308 | 25,3757718 | NA | NA | CHMP4A | 2 |
| 25,27141857 | NA | NA | 25,68760443 | 25,86360611 | 25,48980559 | 25,42454333 | 25,73184727 | 25,65228481 | 25,76749264 | 25,6042202 | 25,86740544 | 25,4278072 | 25,35632274 | 25,4373325 | 25,88187367 | 25,70139319 | 25,53644051 | WDR33 | 2 |
| 26,11107962 | 25,91424278 | NA | 27,12376273 | 27,36794698 | 27,38492152 | 26,58075288 | 27,24181779 | 26,00778559 | 26,98663404 | 27,13025915 | 27,18933354 | 25,76418014 | 25,41622232 | 25,39555061 | 26,85863538 | 26,24427004 | 26,57758718 | PITHD1 | 2 |
| 26,91615792 | 27,23899709 | 27,17959099 | 24,92297704 | 25,66976894 | 26,38160856 | 25,13010201 | 25,37600374 | 23,10872097 | 25,69922337 | 25,74409247 | 26,11137816 | 27,07918055 | 27,03663897 | 26,87255747 | 29,5262664 | 29,91541724 | 29,22068833 | ABLIM1 | 3 |
| NA | NA | NA | NA | NA | NA | NA | 24,84928673 | 25,06212884 | NA | 24,65943458 | 24,98254905 | NA | NA | NA | 24,02284526 | 24,32209869 | 24,30354554 | POLR1E | 2 |
| NA | NA | NA | 25,40054281 | 25,57963129 | 25,72009768 | 26,63034268 | 26,62407946 | 26,52620669 | NA | NA | NA | NA | 24,97254865 | 25,04443548 | NA | NA | 25,00594555 | TWSG1 | 2 |
| NA | 26,1006718 | NA | 25,88502007 | 26,6945817 | 25,91816154 | 27,36151923 | 27,62477672 | 27,4291011 | NA | NA | NA | 27,12149194 | 27,80568927 | 27,3004242 | NA | NA | NA | SIL1 | 2 |
| NA | NA | NA | 24,74051988 | 25,29014066 | NA | 25,3967268 | 25,41425554 | 25,20638931 | NA | 25,02445254 | 25,01780035 | NA | NA | NA | 24,66237153 | NA | NA | IRF2BPL | 2 |
| 25,88562526 | 26,14384607 | 25,83675096 | NA | NA | NA | NA | NA | NA | NA | NA | NA | 25,73502811 | 25,66722613 | 25,67392517 | 25,96811586 | 26,12608906 | 25,73311527 | SEMA6B | 3 |
| 26,36297424 | 26,51800307 | 26,46998257 | 27,82092042 | 28,13584625 | 27,59027991 | 27,82231924 | 27,52723633 | 27,37807295 | 26,3631581 | 26,58649096 | 26,45255647 | 27,30365005 | 27,37260379 | 27,19516642 | 26,70226549 | 26,14230809 | 26,53185319 | NT5DC2 | 2 |
| 30,81191738 | 30,76864455 | 30,75827593 | 29,33212785 | 29,32425091 | 29,63709729 | 30,41780025 | 30,39519511 | 30,40446245 | 28,40271538 | 28,26552892 | 28,11470767 | 29,78455056 | 29,92153067 | 29,5615626 | 31,10017801 | 30,88554671 | 30,80470504 | MMRN2 | 1 |
| 25,52751972 | 26,2959484 | NA | 26,52047805 | 26,67949299 | 26,55477701 | 26,25190883 | 26,61189293 | 26,55186172 | 25,95504572 | 25,92281829 | 25,60005649 | 26,54924907 | 26,75620915 | 26,59192108 | 26,29818804 | 26,31522103 | 26,20571839 | SDF2L1 | 2 |
| NA | NA | NA | 26,76875495 | 27,12238094 | 27,02432571 | 26,60031176 | 26,84577429 | 26,46153003 | 26,37626876 | 26,33601192 | 26,39906 | 26,26203734 | 26,69869365 | 26,43644316 | 25,87645044 | 26,44382707 | 26,25037346 | RIC8A | 2 |
| 28,00099148 | 27,57036647 | 27,34798611 | 25,29498498 | 24,79585378 | 25,52109239 | NA | NA | NA | 28,97571675 | 28,88334867 | 28,80513572 | 28,37438769 | 28,41118718 | 28,56499978 | NA | NA | NA | ESM1 | 4 |
| NA | NA | NA | 26,77781113 | 26,91113667 | 26,41552941 | 25,08253435 | 25,60178575 | NA | NA | NA | NA | NA | NA | NA | NA | NA | NA | XPNPEP3 | 2 |
| 26,63394907 | NA | NA | 26,54467289 | 26,72974872 | 26,3595099 | 26,62072793 | 26,53497237 | NA | 27,07592091 | 26,78431653 | 26,95227077 | 26,64209952 | 26,81909384 | 27,27346539 | 26,93966099 | 26,44540668 | 26,80020595 | EIF2B3 | 2 |
| NA | NA | 25,50112042 | 26,74995901 | 27,19760546 | 26,81115182 | 25,72798463 | 26,26047682 | 26,21066837 | 26,83376277 | 26,48383804 | 26,448671 | 26,0845429 | 26,0428712 | 26,43901438 | NA | NA | NA | RAB12 | 2 |
| 25,15112434 | 25,25641517 | 24,6623172 | NA | NA | NA | NA | 24,05581398 | NA | NA | 24,99217804 | 24,39447159 | 25,23290014 | NA | 24,91624905 | NA | 25,38758615 | NA | BLOC1S4 | 3 |
| NA | NA | NA | NA | NA | NA | NA | NA | NA | 25,7344597 | 25,61739662 | 25,67128168 | 25,42043701 | NA | NA | 25,90841333 | 26,03172627 | 25,93618232 | WDR70 | 2 |
| 26,3833072 | 26,60470094 | 25,9673904 | NA | NA | NA | NA | NA | NA | 26,46774583 | 26,91067932 | 25,92915455 | 26,53235866 | 26,91205092 | 26,61568598 | 25,77600444 | 25,68024447 | 26,03288247 | CMTM6 | 3 |
| 30,23546345 | 30,38277968 | 30,33564571 | 31,49609917 | 31,540424 | 31,58866367 | 31,47716054 | 31,4889844 | 31,62714488 | 31,57145188 | 31,83684274 | 31,92991218 | 31,25979469 | 31,19940864 | 31,14187463 | 30,14807194 | 30,10086727 | 30,17605134 | IGF2BP1 | 2 |
| 32,67598198 | 32,61280698 | 32,65454682 | 31,36332732 | 31,26013131 | 31,29575795 | 30,76982705 | 31,15200736 | 30,9511593 | 32,32820011 | 32,32873512 | 32,45132766 | 32,07179123 | 31,93361 | 32,01581226 | 32,7197771 | 32,89989337 | 32,80593522 | EHD2 | 3 |
| 25,65856322 | 25,27923423 | 25,64600643 | 24,45870148 | NA | NA | 24,1662998 | 24,31437414 | NA | 24,57334578 | 24,69705027 | 24,79029559 | 25,2419814 | 24,94051255 | 25,92698525 | 25,83802629 | 25,23651761 | 25,05564851 | GSKIP | 3 |
| 27,00560296 | 26,82359523 | 27,7881186 | 28,70572967 | 28,78353618 | 28,73918109 | 28,3948477 | 28,02701834 | 28,45151927 | 28,4329679 | 28,12853947 | 28,41373115 | 27,78506527 | 28,13232008 | 28,05030205 | 27,58427016 | 26,98836883 | 27,77994113 | VPS18 | 2 |
| 25,85095633 | 25,90007409 | 25,91702346 | NA | NA | 25,3496573 | NA | 25,15865499 | NA | 25,62463729 | 26,01573916 | 25,92154769 | 25,832049 | 25,62658801 | 25,67728995 | 25,58723524 | 25,89407132 | 25,5276092 | DIP2B | 3 |
| 25,20254595 | 25,53747768 | 25,61647144 | 25,4567918 | 25,40801507 | 25,412383 | 25,85357608 | 25,84302048 | 26,11628526 | 24,99373346 | NA | 24,64817543 | NA | NA | NA | 25,96974133 | 26,03309259 | 26,09105648 | PCDH10 | 1 |
| NA | 26,19880471 | 25,69181613 | 26,69338571 | 26,59733084 | 26,62784065 | 26,69391738 | 26,87044242 | 26,47639394 | 26,91991242 | 27,44616428 | 26,98793533 | 26,344858 | 26,1586935 | 25,97644351 | 26,42675205 | 26,63311763 | 26,54942535 | MBD2 | 2 |
| 25,5920637 | 26,02751408 | 26,00947379 | NA | NA | NA | NA | NA | NA | NA | NA | NA | 26,09186357 | 26,02373371 | 26,50277512 | 26,96675257 | 27,30059876 | 27,22866826 | MRC2 | 3 |
| NA | NA | NA | 26,59960257 | 26,79476368 | 26,62811887 | 27,25078907 | 26,62156654 | 26,65403459 | 26,54942535 | 26,39767359 | 26,78531477 | NA | NA | NA | NA | NA | 26,31207293 | UQCR10 | 2 |
| NA | NA | NA | 28,49533685 | 28,3600125 | 26,85174275 | 27,19131183 | 28,62968284 | 28,07423727 | 27,6707416 | 27,67539318 | 27,5832661 | 28,25826769 | 28,72686148 | 29,09627441 | 28,0403021 | 28,63554134 | 27,57036647 | CFDP1 | 2 |
| NA | NA | NA | 26,91718284 | 26,83665467 | NA | 26,68083465 | 26,66600724 | 26,78081727 | 26,45057572 | 26,6500672 | 26,49057084 | 26,65499061 | 26,81665479 | 26,75289612 | 26,15792296 | 26,28305556 | 26,46273166 | VTI1B | 2 |
| 26,59804114 | 25,98800036 | 26,39573037 | 27,998734 | 28,04197358 | 27,62191582 | 27,76236896 | 27,78113005 | 27,62268393 | 26,57065599 | 26,77228356 | 26,51939858 | 28,25826769 | 27,95626501 | 28,2617684 | 26,94815403 | 26,76711373 | 26,90023536 | TMEM2 | 2 |
| 28,84828405 | 29,13560166 | 28,97254317 | 29,85196609 | 30,14297021 | 30,05452595 | 30,16086808 | 30,306608 | 30,09471504 | 29,17138188 | 29,61029194 | 29,39220876 | 29,5156667 | 29,36026373 | 29,32197401 | 28,94519831 | 29,29756286 | 29,11063169 | AK3 | 2 |
| 25,81124983 | NA | NA | 26,14084646 | 27,02717657 | 26,36779705 | 26,14295074 | 26,03307158 | 26,12238094 | 26,55831432 | 26,68204109 | 26,59291914 | 26,17272232 | 26,15435381 | 25,1668361 | 26,82541613 | 26,45455019 | 26,22387179 | MSRA | 2 |
| 25,78394202 | 26,04178563 | 25,98474509 | NA | NA | NA | NA | NA | NA | NA | NA | NA | NA | NA | 25,00225843 | 26,08832921 | NA | NA | MEMO1 | 3 |
| 27,60279116 | NA | NA | 27,19338143 | 27,77906445 | 27,22094198 | 27,30269177 | 27,62533428 | 27,84326015 | 27,42750352 | 28,09963892 | 27,82614384 | 27,41264143 | 27,43753887 | 27,59917689 | 28,04280859 | 27,77448456 | 28,02422001 | SF3B6 | 2 |
| 25,83723235 | 26,13410383 | 25,8707011 | NA | NA | NA | 23,84364354 | 23,75521604 | NA | NA | NA | NA | 25,5954255 | 25,51929359 | 25,5431541 | 24,5167114 | 24,65310529 | 25,07602288 | BACE2 | 3 |
| 24,89536634 | 25,24027164 | 25,5249821 | 26,97454115 | 27,59462849 | 27,4144975 | 26,79563087 | 26,80808552 | 26,28404463 | 26,33306251 | 26,28409759 | 25,4506701 | 26,40184511 | 26,4021217 | 26,05854143 | 25,27625506 | 25,24008963 | 26,06960481 | NFS1 | 2 |
| NA | NA | NA | 27,86354704 | 27,44340027 | 27,76154494 | 27,23662709 | 27,46388553 | 27,56884552 | 26,73492478 | 27,26981469 | 27,27755034 | NA | 26,17373279 | 26,40116155 | 27,10149357 | 27,32671708 | 26,57758718 | EPN1 | 2 |
| 26,14442969 | 26,28718395 | 26,1998158 | NA | 25,13402547 | NA | NA | NA | NA | NA | NA | NA | 25,99422998 | 26,58448523 | 26,61960902 | 26,00154993 | 26,35548284 | 26,6957767 | ENPP4 | 3 |
| 27,87689535 | 27,6953121 | 28,08527259 | NA | NA | 25,6278963 | 26,089845 | 26,01627068 | 26,69232178 | 26,78469095 | 27,08598165 | 27,47745974 | 27,09154079 | 26,53179371 | 27,1191186 | 27,16269373 | 27,76894421 | 27,75059802 | SRSF6 | 3 |
| 30,23008943 | 30,38555934 | 30,14479433 | 29,57287977 | 29,94893377 | 29,6503754 | 30,24524524 | 30,21754868 | 29,94251621 | 28,33014127 | 28,4172369 | 28,53274508 | 29,37509234 | 29,08126489 | 29,26286623 | 30,2067619 | 29,97951041 | 29,99293976 | ACAT1 | 1 |
| NA | NA | NA | 24,89495021 | NA | 25,07030114 | 25,8099506 | 26,04038572 | 25,97596269 | NA | NA | NA | NA | NA | NA | 24,60308841 | 25,69866716 | 24,72951536 | NF2 | 2 |
| 26,60667878 | 26,72987835 | 26,29108974 | 28,09270045 | 28,38968805 | 28,00190422 | 27,96046919 | 28,39296616 | 28,17212142 | 28,45959284 | 28,71881818 | 28,47374922 | 27,71581079 | 27,60385246 | 27,63020379 | 27,58262026 | 27,74104733 | 27,55545006 | DNAJC7 | 2 |
| 25,30758106 | 25,03006393 | 25,10976531 | NA | NA | NA | NA | NA | NA | NA | NA | NA | NA | NA | NA | 25,88201365 | 25,30260462 | 25,95599907 | NTN4 | 1 |
| NA | NA | NA | 26,71161615 | 26,80636554 | 26,5567806 | 25,79012155 | 26,05355788 | NA | 25,75946379 | 25,72431983 | 25,69999112 | 26,24061739 | 26,09502705 | 25,82106645 | NA | NA | NA | AGO1 | 2 |
| 24,76855306 | 23,86315708 | 25,49246691 | 26,62198566 | 26,90035054 | 26,65157336 | 27,25979469 | 27,13320243 | 26,83364215 | 28,01132 | 28,36431924 | 28,33998323 | 26,42938848 | 26,61104867 | 26,4410269 | 25,64735214 | 25,10952621 | 25,77738476 | KIAA1598 | 2 |
| NA | NA | NA | 28,24213591 | 28,71943849 | 28,75579544 | 27,52065788 | 27,64951911 | 27,64485197 | 27,16748706 | NA | NA | 27,53200188 | 27,86135957 | 27,81860636 | NA | NA | NA | CNIH4 | 2 |
| 26,72390338 | 26,67841875 | 26,43088829 | 25,98139534 | 25,15065963 | NA | NA | NA | NA | 25,83446218 | 25,07630838 | 25,3098721 | 27,24798588 | 26,99258867 | 26,99744242 | 25,15332967 | 25,59482779 | 25,56110005 | DIAPH2 | 3 |
| 23,15123275 | NA | 23,33442715 | NA | NA | NA | NA | 24,41235069 | NA | 23,4266241 | 23,78066711 | 23,60828731 | NA | NA | NA | NA | NA | 23,17586199 | PIP5K1A | 2 |
| 26,27923423 | 26,03795877 | 25,25880682 | 27,02886332 | 27,89029527 | 27,54752919 | 27,3378161 | 27,3069556 | 27,97185829 | 26,65129963 | 25,89305299 | 26,72194965 | 27,84637225 | 27,78631231 | 27,88988927 | NA | 26,30293576 | 26,39067227 | APOC3 | 2 |
| 26,46881804 | 26,18144999 | 25,85635732 | NA | NA | NA | 25,35669214 | 26,12856896 | 25,46911314 | 24,37676555 | NA | 24,74237153 | 24,59747292 | 24,88822932 | 24,63782282 | 27,24308985 | 27,21595314 | 26,70530122 | BCAM | 3 |
| 26,39914152 | 26,93977307 | 26,73182138 | 25,05386856 | 25,32909775 | NA | 25,10805092 | NA | 24,89744518 | 25,89635994 | 25,81906947 | 25,70120809 | 26,29659615 | 26,54663168 | 26,38743824 | 26,34340153 | 26,61372046 | 26,3202403 | SEPT6 | 3 |
| 26,30535596 | 26,15830828 | 26,39275331 | 27,72292684 | 27,76407888 | 27,70272786 | 27,74406679 | 27,73918109 | 28,31704275 | 28,2673162 | 27,96911552 | 28,15754717 | 27,0437474 | 26,94960179 | 26,77818724 | 26,71332169 | 27,56005056 | 27,39737982 | CBX1 | 2 |
| 28,50453401 | 28,51972101 | 28,54572626 | 27,41981041 | 26,96554211 | 27,9714198 | 26,36937889 | 25,76329383 | 26,22324579 | 26,71842627 | 27,95160399 | 27,89186017 | 27,77548962 | 28,16336575 | 28,35401184 | 28,74762422 | 28,7884298 | 28,64003173 | ERG | 3 |
| 27,91461917 | 27,04447717 | 27,77925236 | 28,76739793 | 28,85549025 | 28,49933455 | 28,70833057 | 28,94066936 | 29,22124168 | 28,330013 | 27,89567836 | 28,15286567 | 27,27550931 | 27,32422945 | 27,68872519 | 27,87249876 | 27,92013965 | 27,75665455 | CUTA | 2 |
| NA | 26,29331939 | NA | 26,69059121 | 26,00002442 | 26,05856207 | 26,48314611 | 26,41494905 | 26,19172599 | 26,86525922 | 25,71915775 | 26,65444439 | 26,62254431 | 26,15954063 | 26,57701085 | 26,06406316 | 26,35756489 | 26,12742795 | RPL37A | 2 |
| 25,47540467 | 25,61837721 | 25,84340393 | NA | NA | NA | NA | NA | NA | NA | NA | NA | NA | NA | NA | 26,05465532 | 25,48105293 | 26,18012238 | CREB1 | 1 |
| 25,35019706 | 25,51761269 | 25,632147 | NA | NA | NA | NA | NA | NA | 25,0638575 | 25,17790064 | 25,54889644 | 25,24183597 | 24,79545748 | NA | 25,88725335 | 26,11263135 | 25,83958883 | PSPH | 3 |
| 27,21076125 | 27,3011223 | 27,34028043 | 28,60883014 | 28,89177328 | 28,61375558 | 28,56598033 | 28,66003333 | 28,49205426 | 27,85138534 | 28,13496552 | 28,07933316 | 28,060697 | 27,8787087 | 27,87097148 | 28,28784446 | 28,25304575 | 28,27905707 | UGP2 | 2 |
| 27,33185032 | 27,33116694 | 27,81616649 | 25,31858639 | 25,24190869 | 25,3874218 | NA | 25,24890882 | NA | 25,58434185 | 25,48203833 | NA | NA | NA | 26,31565292 | 26,54113148 | 26,95548922 | 26,66438044 | ARVCF | 3 |
| 26,31361324 | 26,385547 | 26,24407039 | 25,09976932 | NA | NA | NA | NA | NA | NA | 25,68445094 | NA | 26,19921672 | 26,34771584 | NA | NA | NA | 26,12121525 | NDUFB3 | 3 |
| NA | NA | 24,57871036 | 24,67986878 | NA | NA | NA | NA | NA | 25,16729563 | 25,16315458 | 25,28503302 | 24,86641528 | 25,49310856 | 25,05440692 | NA | NA | NA | FARP1 | 4 |
| 26,36924575 | 26,28778282 | 26,23737497 | NA | NA | NA | NA | 25,45910813 | NA | 25,98104687 | 26,11703872 | 26,06570735 | 25,98793533 | NA | 26,07993329 | 26,20045205 | 26,05343358 | 26,33068838 | RAPH1 | 3 |
